# Engineering fluoroacetate dehalogenase by growth-based selections to degrade non-natural organofluorides

**DOI:** 10.1101/2025.07.16.665192

**Authors:** Suzanne C. Jansen, Pauline van Beers, Clemens Mayer

## Abstract

The widespread use of organofluorides in modern society has inadvertently led to the bioaccumulation of harmful pollutants, most prominently per- and polyfluorinated alkyl substances (PFAS). In principle, tailored biocatalysts able to cleave C—F bonds represent an attractive strategy to combat this (emerging) environmental crisis. However, Nature is largely impartial to C—F bonds, with fluoroacetate dehalogenases (FAcDs) standing out as a notable exception, catalyzing the hydrolysis of single C—F bonds in fluoroacetate at high turnover rates. To expand the substrate scope of FAcDs and harness its catalytic prowess for non-natural organofluorides, we designed and applied a robust growth-based selection strategy for large-scale FAcD engineering. Specifically, we demonstrate that FAcD-catalyzed C—F bond cleavage of (natural and) synthetic organofluorides generates metabolizable carbon sources for bacteria, enabling in vivo enrichment of active FAcD variants. By forcing populations expressing diverse FAcD-libraries to utilize various organofluorides as sole carbon source, we elicited a broad panel of FAcD variants that displayed improved activities and drastically altered substrate profiles. In these efforts, we also identified a previously overlooked inhibition pathway, which largely impedes the conversion of gem-difluoride compounds. Overall, our study presents the first large-scale engineering campaign of FAcDs and introduces an operationally simple selection platform that paves the way toward adapting these enzymes for the sustainable degradation of contaminating organofluorides.

## Introduction

Organofluorides have become indispensable in modern society, as they have found widespread applications as pharmaceuticals, agrochemicals, oil/water-repelling materials, or surfactants.^1-4^ The unique physiochemical properties and inherent stability of C—F bonds make them ideal to tailor the properties of molecules and materials alike.^5,6^ However, the stability of C—F bonds also has unintended consequences, such as organofluorides persisting and accumulating in the environment.^7,8^ Particularly, the contamination of soil and water by per-and polyfluorinated alkyl substances (PFAS) is detrimental to ecosystems and human health.^9,10^ Consequently, sustainable and efficient strategies to degrade PFAS and other organofluoride species are in urgent demand.

Bioremediation with the help of specialized biocatalysts and/or bacteria is an attractive strategy to seek and destroy organofluorides. However, Nature is largely agnostic to C—F bonds. Indeed, only a single natural enzyme class is known to selectively install a C—F bond^11^ and only about a dozen natural products have been identified to contain fluoride atoms.^12^ Among these compounds, fluoroacetate (FA) is particularly notable, as it is produced by ∼40 plant species as a defense mechanism against herbivores.^13,14^ Specifically, FA functions as a metabolic poison that can enter the citric acid cycle as fluoroacetyl-CoA, where it is subsequently converted to 2-fluorocitrate (**Fig. 1A**).^15^ The latter compound is a potent aconitase inhibitor that leads to the accumulation of citrate in the blood and, ultimately, shuts down the citric acid cycle.

**Figure 1:**
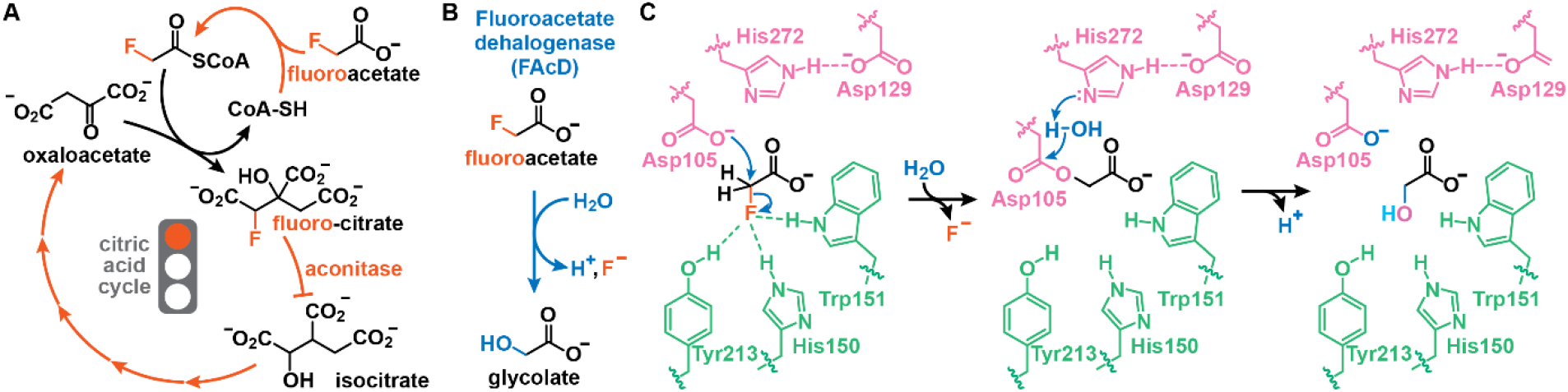
The role and degradation of fluoroacetate in nature. **A:** The toxicity of fluoroacetate (FA) derives from its ability to enter the citric acid cycle as fluoroacetyl-CoA. Following condensation with oxaloacetate, a competitive aconitase inhibitor (2-fluorocitrate) is produced that stalls the citric acid cycle, resulting in citrate accumulation and ATP depletion. **B:** Fluoroacetate dehalogenases degrade FA by hydrolyzing the C—F bond to yield glycolate, a proton, and fluoride. **C:** The FAcD-catalyzed defluorination is generally accepted to proceed through an S_N_2 mechanism, in which Asp105 performs a nucleophilic attack. Concomitant C—F cleavage is facilitated through a specialized fluoride-binding pocket. The resulting acyl-enzyme intermediate is cleaved by water (activated by His272) to release glycolate with a subsequent proton transfer regenerating the nucleophile.

In response to the (general) toxicity of FA, soil bacteria and those residing in cattle rumen have evolved fluoroacetate dehalogenases (FAcDs, EC 3.8.1.3), which neutralize this metabolic poison by promoting its hydrolysis to glycolate (**Fig. 1B**).^16-19^ To do so, FAcDs employ an Asp-His-Asp catalytic triad that facilitates C—F bond cleavage in a two-step mechanism, which is supported by both experimental and computational efforts (**Fig. 1C**).^20-25^ Aided by a specific fluoride binding pocket, a catalytic aspartate first substitutes the fluoride to yield an acyl-enzyme intermediate, which, with the help of a histidine, is subsequently hydrolyzed to release glycolate.^20,22,26,27^

Somewhat surprisingly, although characterized FAcDs efficiently hydrolyze FA with rates >10 s^-1^, examples to engineer these enzymes to accept alternative non-natural fluorinated substrates or improve their activity are scarce. Indeed, we could only identify a single prior report in which amino acid substitutions were introduced in a FAcD to accept bulkier substrates.^28^ We ascribe the scarcity of FAcD-engineering studies to the difficulties associated with assaying enzymatic activity – FAcDs neither act on UV/Vis-active substrates/products nor feature an assayable cofactor. While proton release can be coupled to a pH indicator as readout,^29^ established assays are not readily adaptable to (high-throughput) enzyme-engineering efforts, nor monitor reactions inside host organisms.

To overcome this challenge, we present a technically simple growth-based selection systems to engineer FAcDs. Specifically, we demonstrate that (non-)natural organofluorides can be converted to carbon sources by active defluorinases, enabling us to faithfully elicit improved FAcD variants from diverse populations by serial passaging. Notably, kinetic characterization of selected variants revealed FAcDs with increased activity and/or drastically altered substrate selectivity. In these efforts, we also identified a previously unknown inhibition pathway that prevents the conversion of gem-difluoride compounds. By having established a straightforward means to enable the large-scale engineering of FAcDs, we anticipate that our approach will prove useful for tailoring the activities of these defluorinases toward emerging and harmful organofluorides that persist in our environment.

## Results and Discussion

### Evaluating the substrate scope of FAcD-H1

From the set of previously characterized defluorinases, we chose FAcD-H1 (=H1) from *Delftia acidovorans* strain B (formerly *Moraxella* sp. strain B) for our studies.^17,22^ To evaluate its substrate scope, we produced and purified H1 and incubated it with FA as well as non-natural substrates featuring an additional methyl group and/or a second C—F bond. To assess defluorination activity, we employed ^19^F-NMR spectroscopy, which enables us to monitor both fluoride (F^-^) release and substrate consumption in complex mixtures (see **Supporting Information** for details).

Consistent with prior reports,^22^ H1 (0.5 μM) rapidly converted FA (10 mM) with an apparent initial rate (*v*_0,app_) of 32 ± 4.5 s^-1^ (**Fig. 2A** and **Supporting Table S1**). The more sterically hindered substrate 2-fluoropropionate (FP, racemic) was also readily accepted, albeit with a 17-fold lower initial rate of 1.9 ± 0.4 s^-1^. Conversely, the defluorination of gem-difluorinated substrates proved significantly more challenging (**Fig. 2A** and **Supporting Table S1**): the activity of H1 for 2,2,-difluoroacetate (F_2_A) was ∼9,000-fold lower than for FA, while for 2,2-difluoropropionate (F_2_P) no significant fluoride formation could be observed, even after prolonged reaction times (6 days) and in presence of higher enzyme concentrations (20 μM).

**Figure 2:**
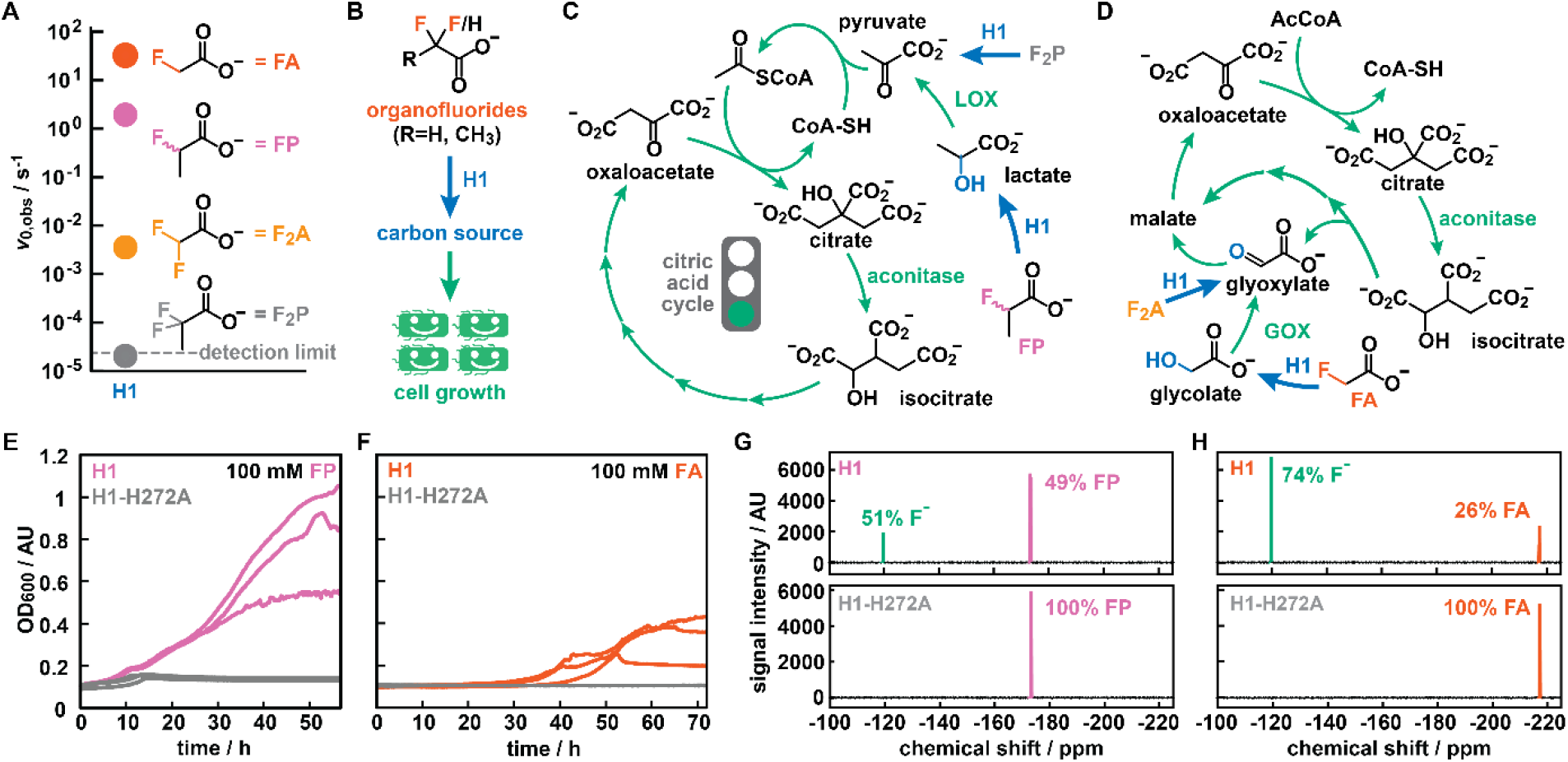
Design and validation of a growth-based selection strategy for FAcD engineering. **A:** Initial rates of H1 on four organofluorides. Rates were determined by ^19^F-NMR and represent the average of 2-3 measurements from biological duplicates. **B:** In growth-based selections, the enzymatic defluorination of either substrate by H1 yields a suitable carbon source that promotes *E. coli* growth in minimal media. **C:** FP or F_2_P can enter the citric acid cycle once successfully converted to lactate or pyruvate, respectively. LOX = lactate oxidase, GOX = glycolate oxidase. **D:** FA or F_2_A can enter the citric acid cycle via the glyoxylate shunt once successfully converted to glycolate or glyoxylate, respectively. **E-F:** Growth curves (in triplicate) of H1-or H1-H272A-producing *E. coli* in presence of 100 mM FP (**E**) or 100 mM FA (**F**) as sole carbon source. Activity of H1 enables growth under these conditions, whereas a host producing inactive enzyme variant H1-H272A cannot proliferate. **G-H:** ^19^F-NMR spectra of culture supernatants from **E-F**, showing the successful conversion of FP (**G**) or FA (**H**) to F^-^ for cells producing H1, but no F^-^ release for cells producing H1-H272A.

### Design and validation of growth-based selections

While ^19^F-NMR provides a convenient means to characterize purified FAcD variants, it lacks the necessary throughput for large-scale directed evolution campaigns. To allow for the engineering of H1 toward non-natural organofluorides, we next explored the feasibility of in vivo growth-based selection strategies, which are easy-to-use, low-cost, and effective to assay millions of enzyme variants in parallel.^30,31^ To create the necessary link between enzymatic activity and host survival, we designed and evaluated simple growth-based selections, in which the conversion of fluorinated substrates provides the sole carbon source for growth of the host organism, *Escherichia coli* (**Fig. 2B**). In principle, all four substrates (FA, FP, F_2_A, and F_2_P) yield potential carbon sources upon successful defluorination. Specifically, defluorination of FP and F_2_P by H1 yields lactate and pyruvate, respectively, which are C3 carbon sources that can directly fuel the citric acid cycle (**Fig. 2C**). Conversely, FA and F_2_A give glycolate and glyoxylate, respectively, which are C2 carbon sources that can be metabolized via the glyoxylate shunt (**Fig. 2D**). Additionally, while F_2_A, FP, and F_2_P at concentrations of <100 mM are innocuous compounds for *E. coli*, FA is inherently toxic and thus adds an additional selective pressure.

To validate whether this approach can differentiate between active and inactive defluorinases, we constructed H1-H272A, in which the essential histidine from the catalytic triad is substituted for an alanine. Using an inactive H1 variant as a negative control is ideal for growth-based selections, as it mimics the burden bestowed on the host cell upon induction, which also affects fitness. Following production and purification of H1-H272A, we confirmed that it is produced at levels comparable to the wildtype and it lacks any appreciable activity for any of the four substrates (**Supporting Table S1**).

Next, we monitored bacterial growth in a defined minimal medium at 30 °C under varying conditions by measuring the optical density at 600 nm (OD_600_) in a plate reader (see **Supporting Information** for details). As positive controls, we provided *E. coli* expressing H1 with 100 mM glycolate, 50 mM lactate, or 50 mM pyruvate as sole carbon sources, which gave a robust growth phenotype for the C3 carbon sources pyruvate and lactate, and a more moderate growth phenotype for the C2 carbon source glycolate (**Fig. S1**). Gratifyingly, when substituting lactate for 100 mM FP, cultures producing H1 could proliferate over a period of 48 hours (OD_600_ from ≈ 0.5–1), whereas cultures featuring H1-H272A failed to grow under otherwise identical conditions (**Fig. 2E**). When supplementing minimal media with 100 mM FA as a sole carbon source, H1 producers were again the only ones that could proliferate, but displayed extended lag times (∼36 hours) and reached lower cell densities (end OD_600_ ≈ 0.3–0.4, **Fig. 2F**). Together with the subsequent stationary phase, this less robust growth phenotype likely results from both the inherent toxicity of FA and glycolate being a relatively poor carbon source.^32^ Lastly, we also attempted to grow *E. coli* expressing H1-variants on F_2_A and F_2_P as sole carbon sources. However, these attempts proved fruitless, consistent with the low activities H1 displays for these substrates (**Fig. S1**).

To further attest that the observed growth phenotypes correlated with enzymatic activity, we subjected cells’ supernatants to ^19^F-NMR analyses. H1-producing cells displayed significant conversion of racemic FP (51%, consistent with the high enantioselectivity of FAcDs) and substantial fluoride formation (74%) when grown in FA (**Figs. 2G-H**). Conversely, none of the substrates were converted by bacteria producing H1-H272A under otherwise identical conditions (**Figs. 2G-H**). For bacteria that failed to grow in the presence of F_2_A or F_2_P as sole carbon source, ^19^F-NMR did not detect any fluoride release over a period of 72 hours (**Fig. S1**).

Combined, these experiments demonstrate that FAcD activity can be linked to host fitness by employing either FA or FP as the sole carbon source. While both approaches are suitable for growth-based selections, we predominantly employed FP in our engineering efforts described below, as: (1) the growth phenotype provided by the carbon source (lactate) is more pronounced, and (2) engineering FAcD-H1 for non-natural and more sterically demanding organofluorides aligns with our goal to, ultimately, degrade persistent pollutants.

### Design and generation of H1-libraries

To apply our growth-based selections, we constructed H1 libraries by randomizing residues lining the active site of this enzyme. Toward this end, we first constructed an *AlphaFold2^33,34^* model of H1 and then submitted the model to the online webserver *HotSpot Wizard*,^35^ which identified Gln245 and Ser147 as targets for randomization (**Fig. 3A**). To probe potential epistatic interactions, we included Met246, whose side chain points toward the active site. Finally, we also chose Trp180, as substitution for smaller residues at this position facilitated turnover of sterically-demanding α-fluorocarboxylic acids in a homologous FAcD (RPA1163 from *Rhodopseudomonas palustris* CGA009).^28^

**Figure 3:**
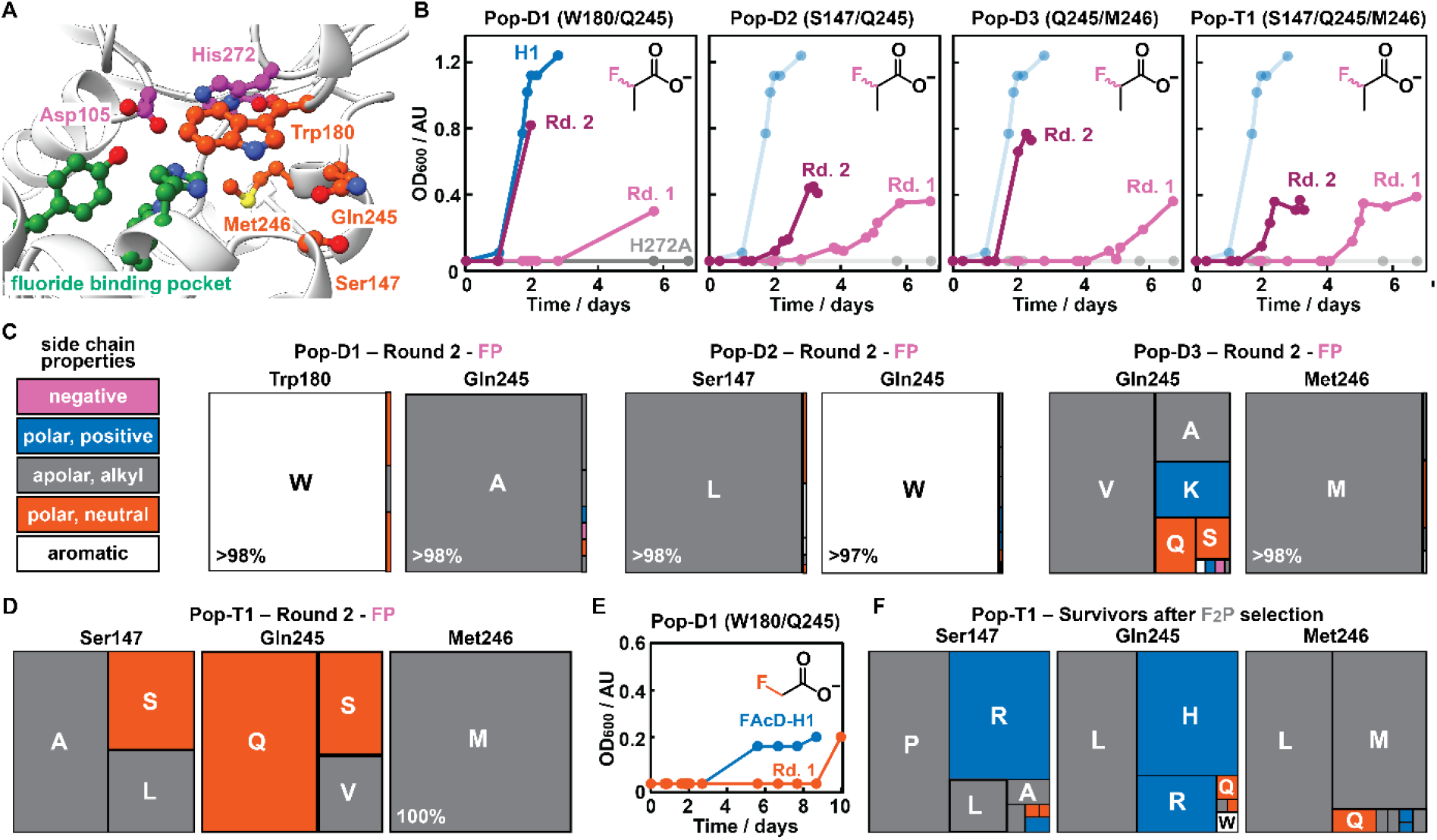
Selecting better FAcDs. **A:** Close-up of the active site of H1, showing the catalytic triad in purple (Asp105 and His272), the fluoride binding pocket in green (His150, Trp151 and Tyr213), and residues targeted for site-directed mutagenesis in orange (Ser147, Trp180, Gln245, and Met246). **B:** OD_600_ measurements over time of two consecutive serial passages (pink and purple) for all populations using FP as sole carbon source. Controls (H1 and H1-H272A) were measured in parallel with Pop-D1, and are shown in blue and grey, respectively. The same control traces are included in the other populations with lower opacity for comparison. Note that the lag period decreases from the initial to the second passage. **C-D:** Treemap charts depicting the enrichment of amino acids across randomized positions in populations after two passages on FP as sole carbon source. On the left, a legend for the color-code of the properties of amino acid side chains. **E:** OD_600_ measurements over time showing growth of Pop-D1 (orange) or H1 (blue) with FA as sole carbon source. **F:** Treemap chart depicting the enrichment of amino acids across randomized positions for survivors of Pop-T1 after prolonged incubation with F_2_P as sole carbon source.

Based on these considerations, we constructed a total of four libraries using degenerate NNK codons (see **Supporting Information** for details): three libraries featuring two randomized positions giving Lib-D1 (Trp180/Gln245), Lib-D2 (Ser147/Gln245), and Lib-D3 (Gln245/Met246); and one library in which three positions were randomized, Lib-T1 (Ser147/Gln245/Met246). Following transformation into a cloning *E. coli* strain, the number of transformants exceeded the theoretical genetic diversity of each library by a factor of 3.9 – 20 (see **Supporting Table S2**). Next, we transformed each library into the appropriate expression strain and applied whole-plasmid sequencing (WPS) of pooled plasmid samples to assess the diversity of populations Pops-D1-D3 and Pop-T1 prior to selection. Mapping obtained reads to the reference plasmid and quantifying the amino acid distribution across randomized positions attested on the overall high degree of amino acid diversity for each position in every library (**Supporting Figure S2**).

Lastly, to provide an estimate of active H1-variants in the transformed libraries, we determined the growth characteristics in presence of FP as sole carbon source of 282 individual colonies across all four populations. While for Pop-D2 more than 50% of all inspected colonies were able to utilize FP as a sole carbon source, Pop-D1, Pop-D3, and Pop-T1 fared significantly worse, with only <10% of all inspected variants being able to grow within 48 hours (**Supporting Fig. S3**). These results suggest that the latter three libraries contain a significant number of variants with deleterious mutations, while substitutions at Ser147 & Gln245 (in Pop-D2) are generally more permissive.

### Autonomous selection of H1-variants

With all populations featuring variants with growth phenotypes, we next performed a series of selection campaigns in which individual populations were challenged to utilize different organofluorides as sole carbon sources. Specifically, we subjected all four populations to two consecutive serial passages in the presence of FP. Monitoring the OD_600_ over time revealed that growth phenotypes for the initial passage required 3–5 days, significantly longer than H1-producing cells (2 days, **Fig. 3B**). This extended lag period suggests that indeed many cells produced inactive variants that could not proliferate, while the few cells with *active* variants required longer lag periods to reach detectable cell densities. Interestingly, Pop-D2 featured a somewhat shorter lag period than the other populations in the initial passage, again suggesting that substitutions of Ser147 and Gln245 are more permissive. Strikingly, all four populations displayed significantly shorter lag periods (1-2 days) during the following passage, indicative of the successful enrichment of active defluorinases (**Fig. 3B**).

To track the enrichment of individual variants throughout these selection campaigns, we sequenced the remaining genotypic diversity after each passage by performing WPS on pooled plasmids and by Sanger sequencing 4–18 random colonies per passage (**Figs. 3C-D, Supporting Fig. S4** and **Supporting Table S3**). Gratifyingly, these data attested on the robust enrichment of a small number of variants for all selections and revealed notable trends across populations: (1) Trp180 and Met246 appear critical for defluorination activity, since FP-based selections almost exclusively enriched variants retaining these wildtype residues after randomization. This result is consistent with the outcome of our plate reader experiments, in which growth phenotypes for populations that targeted one of these residues were scarce (<10% for Pop-D1, Pop-D3, and Pop-T1). (2) Ser147 and Gln245 are more permissive to substitutions, yet small apolar residues appear favorable, such as Ala/Leu(/Ser) for position Ser147 and Ala/Val for Gln245. (3) Growth-based selections with FP as sole carbon source are robust and do not encourage the enrichment of *escape variants*. Indeed, only in Pop-T1 we detected *hitchhikers*, marked by the excision of the H1 coding sequence in about half of the cells left in the population. These passengers likely originated from a spontaneous recombination event (the host *E. coli* BL21(DE3) possesses a functional recombination gene, *recA*), which provides increased fitness by lowering the gene expression / protein production burden.

Based on all sequencing results from these FP-based selections, we chose nine H1-variants for further characterization as they (1) were strongly enriched after two passages, (2) were present in populations after the initial passage, or (3) performed well in our initial plate reader experiment (see **Supporting Table S4** for a detailed summary). These nine variants are Q245**A** and Q245**G** from Pop-D1; S147**L**-Q245**W** and S147**V**-Q245**W** from Pop-D2; Q245**V**, Q245**M**-M246**L**, and Q245**F**-M246**I** from Pop-D3; and S147**L**-Q245**V** and S147**R**-Q245**P**-M246**L** from Pop-T1. For clarity, shorthand names are used throughout the manuscript that only list the substituted residues (i.e. H1-A, -G, -LW, -VW, -V, -ML, -FI, -LV, and -RPL).

Besides FP-based selections, we also performed a selection campaign utilizing FA as sole carbon source for Pop-D1. While the H1 wildtype control started to proliferate slowly after approximately 5 days, Pop-D1 required 10 days to reach a measurable OD_600_ ≈ 0.2 (**Fig. 3E**). Because the growth conditions were highly stringent, we did not perform a second passage but immediately analyzed the remaining diversity in the population by Sanger sequencing. Interestingly, we found that all 6 tested colonies harbored the same variant: Q245**R** (and W180W, consistent with previous findings) (**Supporting Table S3**). As we did not identify this H1 variant in any of the populations in the FP-based selection, we added H1-R to the panel for further characterization (**Supporting Table S4**).

Lastly, we performed selection campaigns utilizing F_2_P as potential carbon source for Pop-D2, Pop-D3, and Pop-T1. Unfortunately, none of these populations were able to grow to a measurable OD_600_ even after prolonged incubation (14 days). Curious to see whether cells remained viable after two weeks, we plated a fraction of the selection cultures on LB agar plates. Interestingly, Pop-T1 gave rise to ∼10-fold more colonies than Pop-D3, and about ∼100-fold more colonies than Pop-D2 (**Supporting Fig. S5**). Intrigued by this stark difference, we subjected surviving colonies from Pop-T1 to WPS sequencing, which revealed a marked enrichment of H1-variants distinct from our previous efforts (**Fig. 3F**). Notably, Met246, which was deemed critical for defluorinase activity on FP, had been substituted with Leu in 48% of reads. Additionally, Ser147 and Gln245 showed enrichment of positively charged residues (Arg, His) rather than small, apolar ones as observed before. While the observed *survival phenotype* could be unrelated to defluorination activity, we added another three representative H1-variants of this group for further characterization, namely S147**R-**Q245**L** (H1-RL), S147**R**-Q245**H**-M246**L** (H1-RHL), and S147**P**-Q245**L** (H1-PL, **Supporting Table S4**).

### Validation of growth phenotypes of selected H1-variants

Prior to the kinetic characterization of all 13 H1-variants we identified during our selections, we evaluated their growth phenotypes when challenged with the organofluorides of interest. To exclude unintended fitness adaptations resulting from mutations in the vector backbone or within the host’s genotype, we cloned the H1-coding sequences anew and transformed them into fresh *E. coli* hosts.

From the recorded growth curves on (non-)natural organofluorides as sole carbon sources (**Fig. 4A** and **Supporting Fig. S6**), a number of interesting trends across the groups of H1-variants are readily apparent: (1) Hosts producing strongly enriched H1-variants from FP-(or FA)-based selections (H1-A, -LW, -V, -LV(, -R)) displayed strong growth on FP as carbon source (**Fig. 4A**), attesting on the robustness of our growth-based selection strategy. (2) H1-variants chosen from the initial 96-well plate screen (H1-G and -VW) demonstrated repeated growth on FP. (3) Results for variants that featured substitutions to the conserved Met246 varied, with H1-FI displaying robust growth on FP, while H1-ML and -RPL largely failed to proliferate. (4) Survivors from F_2_P-based selections (H1-RL, -PL, and -RHL) could not utilize FP, but as all other tested H1-variants, they also failed to proliferate in presence of F_2_P. Lastly, (5) growth curves on FA and F_2_A did not show consistent proliferation (**Supporting Fig. S6**), a fact that we ascribe to cells possibly residing in stationary phase, excessive stringency by FA, and/or carbon source (=acetate) carryover from the inoculum (see **Supporting Discussion** for details).

**Figure 4:**
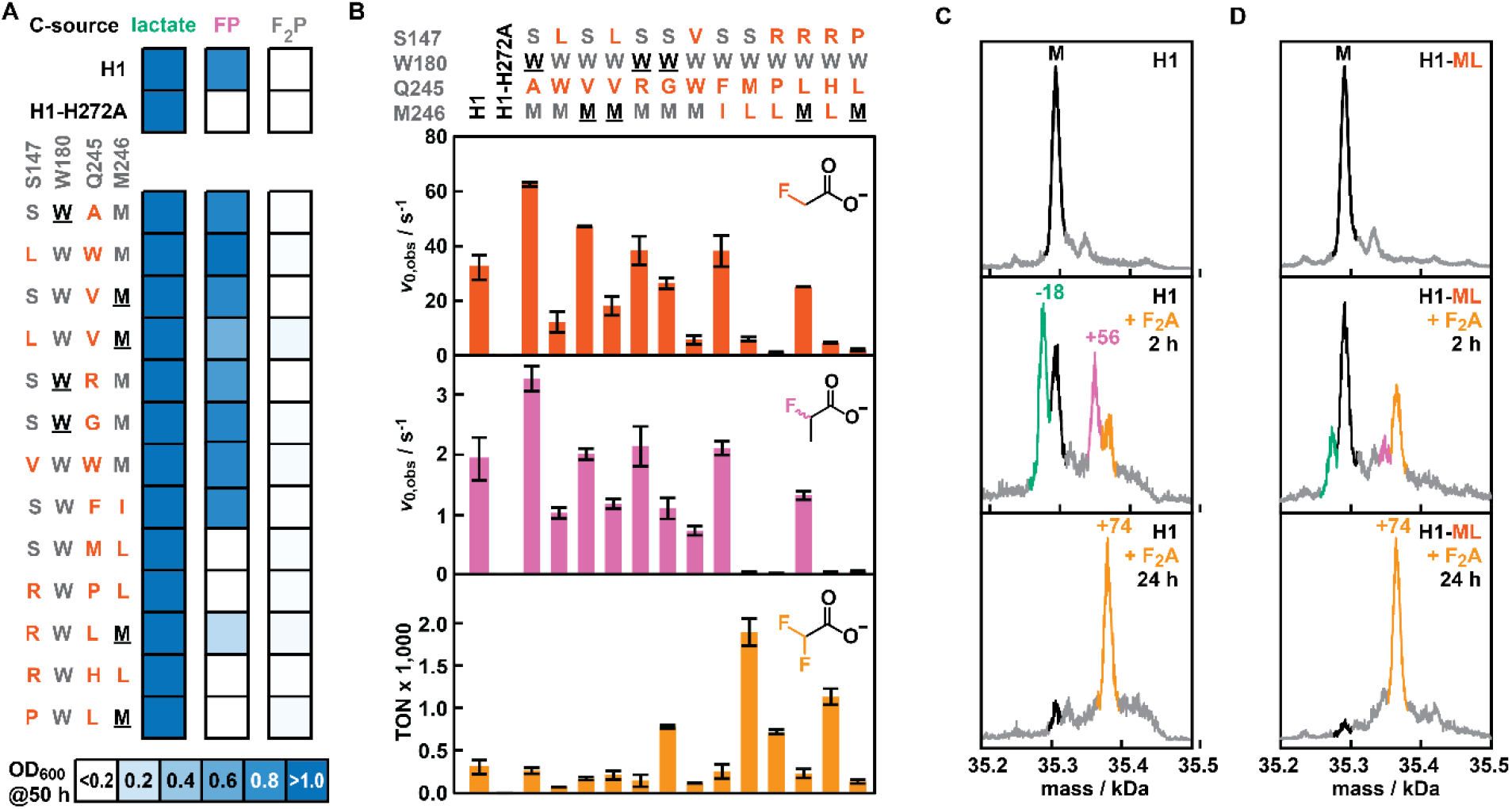
Characterization of selected H1-variants. **A:** Growth properties that H1-variants bestow on their host when challenged to utilize lactate, FP, or F_2_P. OD_600_ values represent the average of 3 measurements from 3 single colonies after 50-hour incubation. The corresponding growth curves can be found in **Fig. S6. B:** Kinetic parameters of H1-variants on FA, FP and F_2_A. Values represent the average of 2-3 measurements from technical duplicates (except for H1, in which biological duplicates were used). Standard deviations are given. **C-D:** Deconvoluted masses from UPLC-MS spectra for H1 (**C**) or H1-ML (**D**) before and during F_2_A incubation. The observed mass (M) for unmodified FAcDs is marked in black, and three mass species are labeled in green (M-18), pink (M+56) and orange (M+74). The corresponding raw spectra can be found in **Fig. S8**.

### Kinetic characterization of H1-variants

To verify that the observed growth phenotypes correlate with the enrichment of active defluorinases, we produced and purified all 13 H1-variants (**Supporting Fig. S7** and **Supporting Table S5**) and determined their kinetic parameters for organofluoride hydrolysis by ^19^F-NMR spectroscopy. Gratifyingly, strongly enriched variants from FP-(or FA)-based selections (H1-A, -LW, -V, -LV (, -R)) all demonstrated activity on both FA and FP (**Fig. 4B** and **Supporting Table S6**). In fact, three of these variants (H1-A, -V, -R) exceeded wildtype activity for both substrates, with H1-A being the most active and displaying an apparent initial rate (*v*_0,app_) of 63 ± 0.82 s^-1^ for FA (2-fold higher than wildtype) and 3.2 ± 0.21 for FP (1.7-fold higher than wildtype). Variants chosen from the initial 96-well plate screen (H1-G, H1-VW) were active on FA and FP but performed (slightly) worse than the wildtype FAcD. Furthermore, the inclusion of variants from the initial serial passage that featured Met246 mutations yielded notable results. Specifically, H1-FI displayed high catalytic activities with a *v*_0,app_ of 38 ± 5.8 s^-1^ for FA (1.2-fold higher than H1) and 2.1 ± 0.11 for FP (1.1-fold higher than H1), demonstrating that Met246 is not essential for defluorination activity. Conversely, variants carrying the Met246Leu substitution performed significantly worse: H1-ML and H1-RPL retained only 20% and 4% of wildtype activity, respectively, in presence of FA, and just 2% and 1% residual activity with FP as the substrate (**Fig. 4B** and **Supporting Table S6**). Lastly, variants originating from F_2_P selections overall performed poorly on FA and FP, with the exception of H1-RL, whose activity on these substrates were comparable to the wildtype FAcD.

To gain insight into the substrate promiscuity for all H1-variants, we further determined their activity with the gem-difluorinated substrates F_2_A and F_2_P (**Fig. 4B** and **Supporting Table S6**). While all variants were able to hydrolyze F_2_A, none of them yielded appreciable conversions for F_2_P (<1%), even at elevated enzyme concentrations (20 μM) and extended reaction times (up to 7 days). Notably, for none of the F_2_A samples we could identify a mono-fluorinated product by ^19^F-NMR, indicating that upon breaking the first C—F bond and hydrolysis of the acyl-enzyme intermediate, the monofluorinated product either undergoes spontaneous defluorination or is rapidly hydrolyzed. Instead, the enzymes seem to undergo catalysis-dependent inactivation, leading to covalent modification of the H1-variants (*vide infra*).

Because of this inactivation pathway, FAcD variants were compared based on their turnover numbers (TONs) after 24 hours rather than their apparent initial rates. This analysis revealed a strong correlation: H1-variants that showed poor activity on mono-fluorinated substrates often exhibited significantly increased TON when tested with F_2_A (**Fig. 4B**). For example, H1-ML and -RHL displayed TONs of 1,900 ± 161 and 1,140 ± 95, thus exceeding the wildtype enzyme (TON = 307 ± 81) by a factor of 6.1 and 3.7, respectively. Moreover, three of the four variants with the highest F_2_A TONs possess a Met246Leu substitution (**Fig. 4B**), a residue that was highly conserved in selections with mono-fluorinated substrates (*cf.* **Fig. 3D-E**). The sole exception in this trend is the *generalist* H1-G (featuring a Gln245Gly substitution) which displays high activities for FA (*v*_0,app_ = 26 ± 2.1 s^-1^), FP (*v*_0,app_ = 1.1 ± 0.17 s^-1^), and F_2_A (TON= 783 ± 21).

### Elucidation of F_2_A-dependent inactivation pathway of FAcDs

As indicated before, we observed F_2_A-dependent inactivation of H1-variants during our kinetic characterization efforts. Notably, while prior works on F_2_A conversion by homologous FAcDs have consistently described low conversions over prolonged incubation periods,^21,36,37^ none of them have pinpointed a catalysis-dependent inactivation pathway as the reason for this poor performance.

To elucidate this pathway, we incubated H1 with an excess of F_2_A and periodically analyzed samples by mass spectrometry (UPLC-MS). Following 2-hour incubation, we identified masses of H1 (M) and three distinct species, namely M-18 Da, M+56 Da, and M+74 Da (**Fig. 4C** and **Fig. S8**). While the former two species are transient, the M+74 Da species becomes the dominant species after 24-hour incubation. Based on prior mechanistic investigations,^21,36,37^ we tentatively assign the three species as follows (**Fig. S9**): (1) M-18 corresponds to the fragmentation of the Asp105 acyl-enzyme intermediate that is formed after the initial defluorination reaction; (2) M+56 is consistent with the formation of an imine intermediate that results from the condensation of glyoxylate and a nucleophilic lysine residue; (3) M+74 matches the mass of an enzyme-glycolate covalent intermediate (Enz—C_2_H_2_O_3_) rather than an enzyme-FA intermediate (Enz—C_2_HO_2_F) with a mass difference of M+76.

To identify potential causes for the increased TONs observed for selected H1-variants, we performed analogous MS time-course studies with H1-ML, which provides a 6.1-fold increase in conversion when compared to H1 (**Fig. 4D**). While this improved variant still suffers from inactivation after 24 hours, we observe a marked decrease in signals corresponding to the transient species M-18 and M+56 after two hours. Given these observations, higher TONs are likely the result of overall slower inactivation, possibly by virtue of slower activity. While the Met246Leu substitution negatively impacts initial rates for the mono-fluorinated substrates FA and FP, this setback is likely critical for the beneficial conversion of F_2_A.

In an effort to further pinpoint the residue that is modified in the M+74 species, we digested (inactivated) H1-variants using (chymo)trypsin and subjected the resulting peptides to LC-MS/MS analysis (see **Supporting Information** for details). Unfortunately, we were unable to detect significant differences between samples of H1-variants in presence and absence of F_2_A, indicating that the observed modification is not stable throughout the proteomics workflow. Drawing on prior mechanistic investigations, we suspect that hydrolysis of a second acyl-enzyme intermediate is hindered, making nucleophilic residues in the active site or catalytic triad likely sites of modification (**Fig. S9**).

## Conclusion

The continuous and widespread use of organofluorides in modern society necessitate sustainable degradation strategies for the harmful, persistent pollutants that arise from fluorinated compounds and materials. While C—F bonds are inherently stable and largely absent from Nature, engineering existing biocatalysts to recognize harmful and persistent pollutants represent an attractive strategy. Toward this goal, FAcDs are particularly promising starting points, as they can hydrolyze C—F bonds in FA multiple times per seconds. However, their engineering toward non-natural organofluorides has remained virtually unexplored.

In this work, we developed and applied a novel growth-based selection platform that enables high-throughput engineering of FAcDs for the degradation of organofluorides. Our cost-effective and technically simple strategy relies on the growth advantage conferred by active biocatalysts that can utilize fluorinated (non-)natural substrates as sole carbon source. By using bacterial growth as a proxy for C—F bond cleavage activity, consecutive serial passages elicited active H1-variants from diverse starting populations. Attesting on the robustness of this strategy, we did not observe *escape variants*, with only one population partially harboring *hitchhikers* which had excised the H1 coding sequence to reduce expression burden. Selected H1-variants displayed up to 2-fold higher activities for FA and 1.7-fold increased activity for FP. Moreover, by including survivors on F_2_P and variants that went extinct after an initial passage, we identified H1-variants with drastically altered substrate selectivity. Notably, large increases in F_2_A turnover required a sharp loss in defluorination activity for FA and FP. A Met246Leu substitution proved key to retard a previously unknown F_2_A-dependent inactivation pathway, which we characterized by time-course UPLC-MS studies. With the degradation of gem-difluorinated carboxylic acids being a critical steppingstone for the degradation of PFAS and related compounds, this inactivation pathway warrants further investigation in the future.

Lastly, the realization of technically simple, growth-based selections based on the catabolism of organofluorides provides ample opportunities for the discovery and engineering of FAcDs. For example, it offers a straightforward means to elicit novel defluorinases from PFAS-contaminated soil samples. Additionally, it is conceivable that other short-chain (gem-di)fluorinated molecules can be used as sole carbon sources, enabling the engineering of FAcDs toward more PFAS-like molecules. Lastly, interfacing this platform with targeted in vivo mutagenesis strategies in the future should allow for the continuous and autonomous exploration of long evolutionary pathways.^38^ As such, we anticipate that the operationally simple, high-throughput selection strategy we developed will prove valuable in making defluorinases fit for the degradation of persistent and harmful organofluorides in our environment.

## Supporting information

19F-NMR spectra

## Acknowledgements

C.M. and S.C.J. are thankful to J. Hekelaar and F. Y. Ho for analytical support. We thank L. Longwitz for valuable feedback and helpful suggestions regarding ^19^F-NMR, A.R. Oliveira for valuable discussions, and J.L. Maehara Said dos Reis for assistance in the preparation of H1-variants. C.M. acknowledges the NWO (OCENW.M20.278 and OCENW.XS24-4.245) and the ERC (ERC-2024-COG 101125957) for funding.

## Author contributions

C.M. and S.C.J. conceived the project; P.v.B. collected preliminary data; S.C.J. designed, performed, and analyzed experiments; C.M. supervised research; S.C.J. and C.M. wrote, edited, and revised the manuscript. All authors approved the final version of the manuscript.

## ASSOCIATED CONTENT

The manuscript is accompanied by a Supporting Information file, which includes Supporting Tables and Figures, a Supporting Discussion, experimental procedures, and sequences of H1-variants used in the study. Additionally, a supporting document is provided, which contains all raw ^19^F-NMR spectra for kinetic characterization of H1-variants and determination of defluorination activity in the supernatant of cultures.

## Notes

### Competing Interest Statement

The authors have declared no competing interest.

