## Supporting Information for "Engineering fluoroacetate dehalogenase by growth-based selections to degrade non-natural organofluorides"

#### Table of contents

1. Supporting Figures S2
2. Supporting Tables S7
3. Supporting Discussion S13
4. Experimental S15
5. Sequences S33
6. Supporting References S37

### 1. Supporting Figures

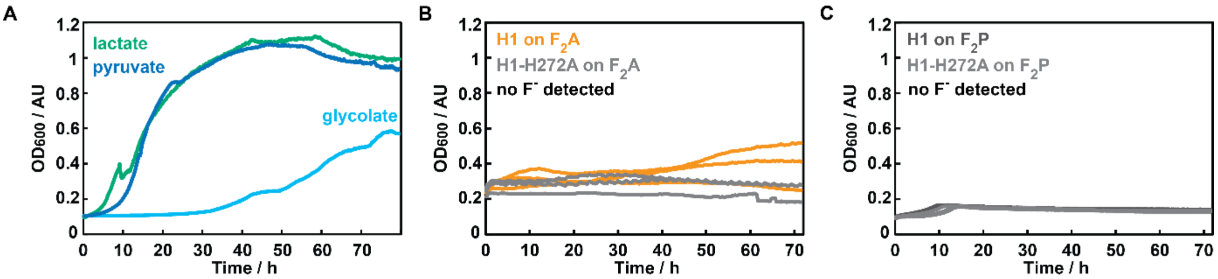


**Figure S1: A:** Positive control growth phenotypes of cells producing H1, when utilizing lactate (green), pyruvate (dark blue) or glycolate (light blue) as carbon sources. **B-C:** Growth phenotype of H1- or H1-H272A-producing cells failing to utilize $F_2A$ (**B**) or $F_2P$ (**C**) as sole carbon source. No $F^-$ was detected in culture supernatant by $^{19}$F-NMR.

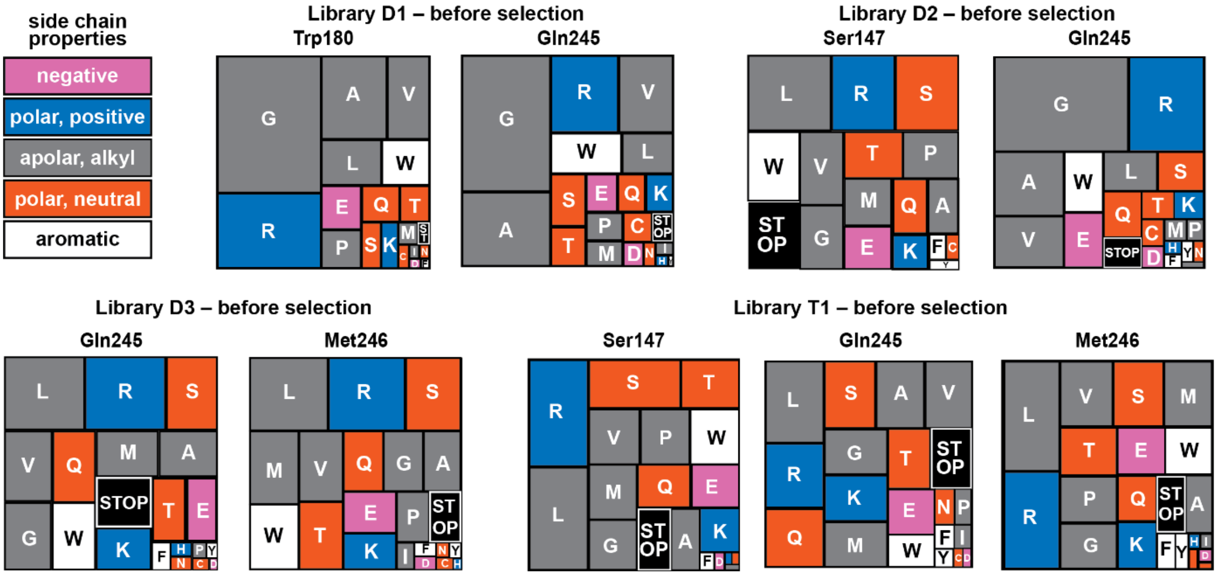


**Figure S2:** Treemap charts depicting the amino acid distribution across randomized positions for all populations prior to selection. Top left, a legend for the color-code of the properties of amino acid side chains. The data is based on raw reads from whole-plasmid sequencing samples (see *Experimental*).

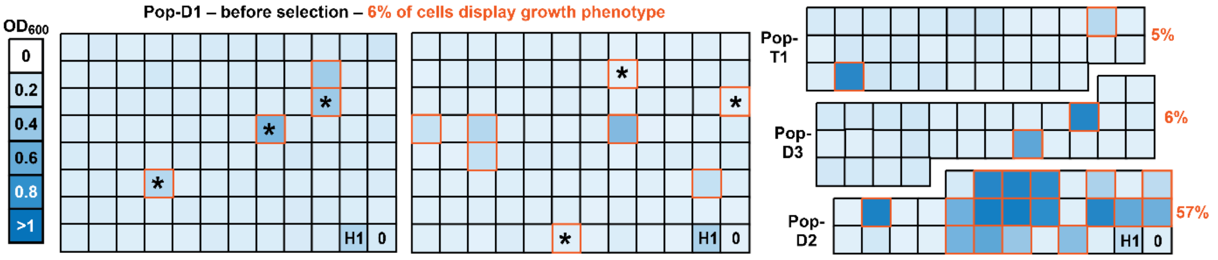


**Figure S1:** Distribution of H1-variants displaying a growth phenotype across populations when utilizing FP. Each block depicts the $OD_{600}$ for a host producing a randomized H1-variant after 50 hours of growth on FP. Values represent single measurements. Blocks marked with orange lines were considered to have a positive growth phenotype. Blocks marked with asterisks showed a positive growth phenotype within 24 hours of growth, and were subjected to $^{19}$F-NMR analysis.

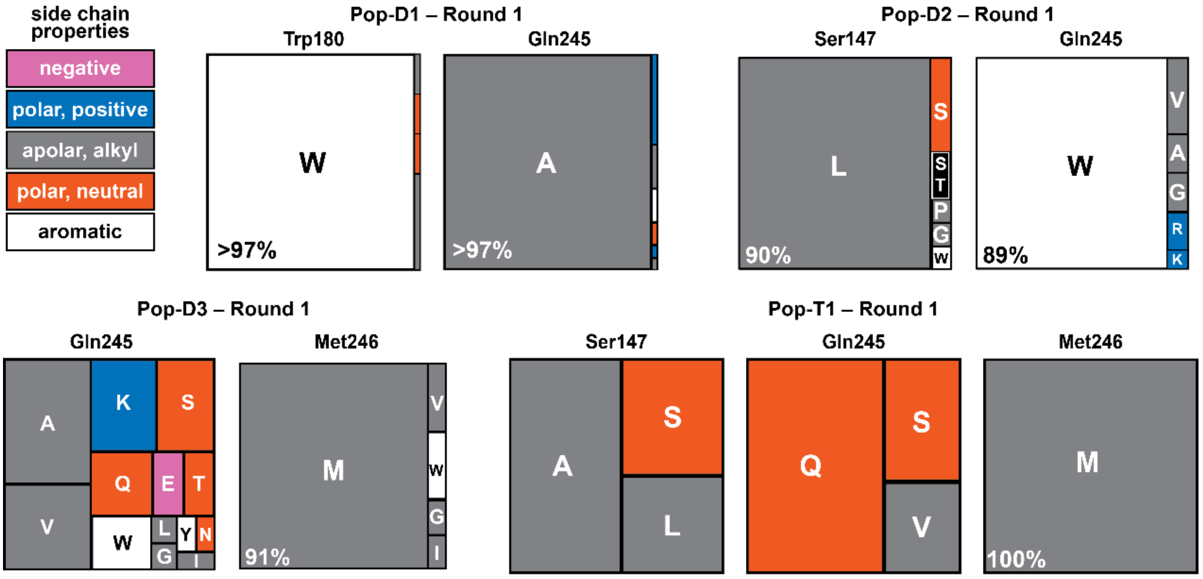


**Figure S2:** Treemap charts depicting the enrichment of amino acids across randomized positions in populations after one passage on FP as sole carbon source. Top left, a legend for the color-code of the properties of amino acid side chains. The data is based on raw reads from whole-plasmid sequencing samples (see *Experimental*).

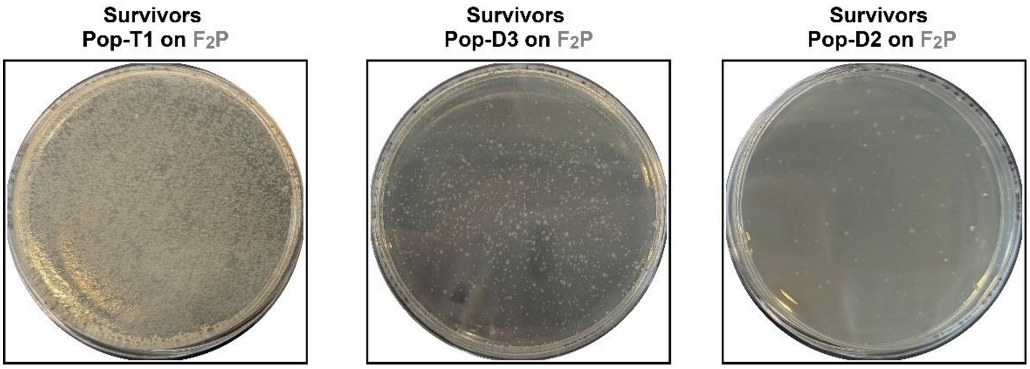


**Figure S3:** Comparison of surviving bacteria following a 14-day incubation with $F_2P$ as the sole carbon source.

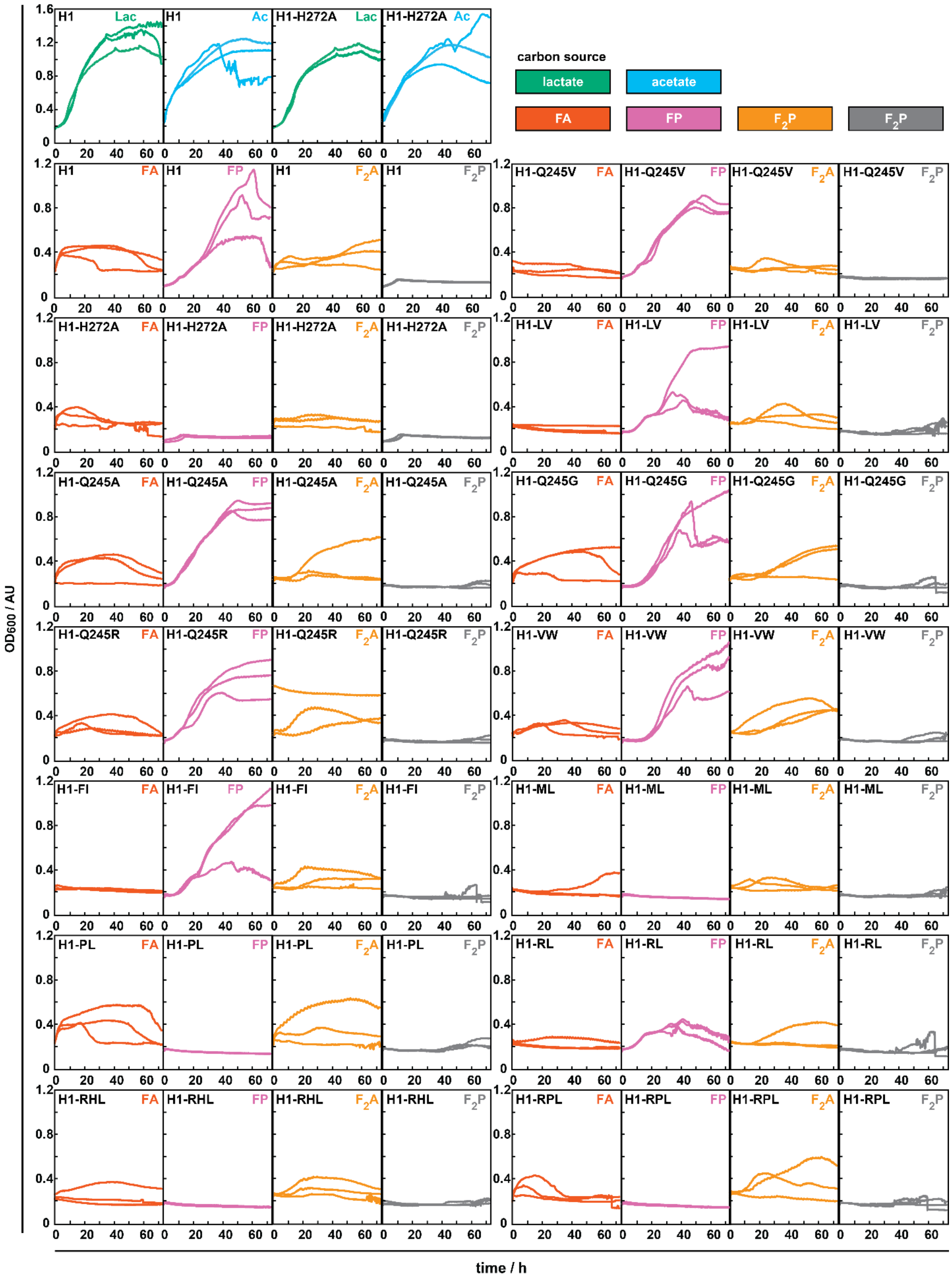


**Figure S4:** Growth curves (in triplicate) of H1-producing *E. coli* in presence of 100 mM FA (red), FP (purple), $F_2A$ (orange), and $F_2P$ (grey). Additionally, the growth curves for H1 and H1-H272A for lactate (green) and acetate (blue) are shown at the top. These growth curves are representative for all H1-variants. Top right, a legend for the color-code of the carbon sources used.

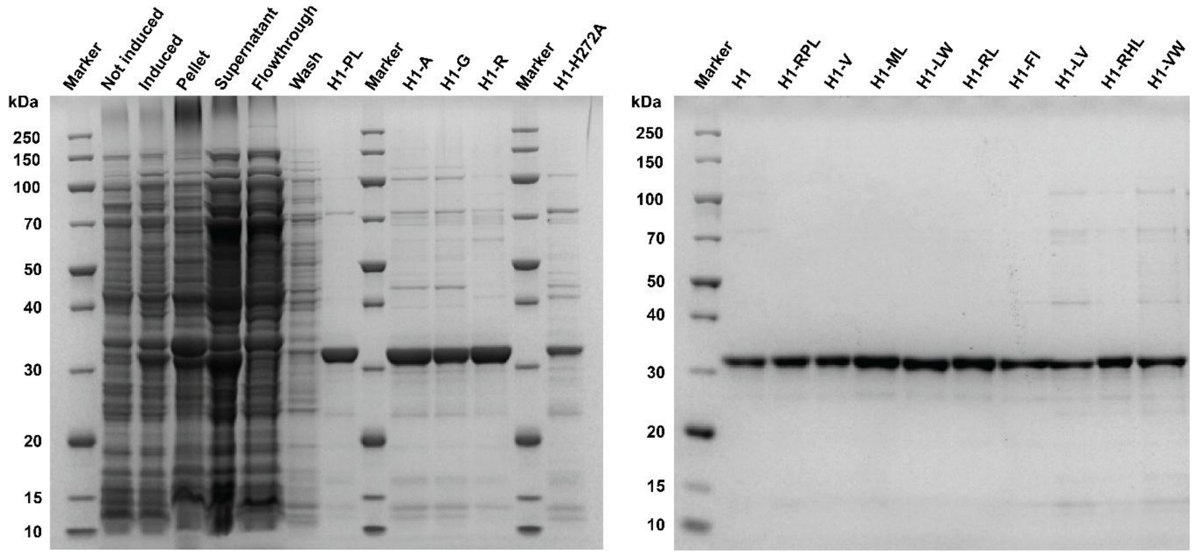


**Figure S5:** SDS-PAGE analyses of production and purification of H1-variants**.** The gels depict representative fractions throughout the production and purification process for H1-PL and the elution fractions of all H1-variants after purification.

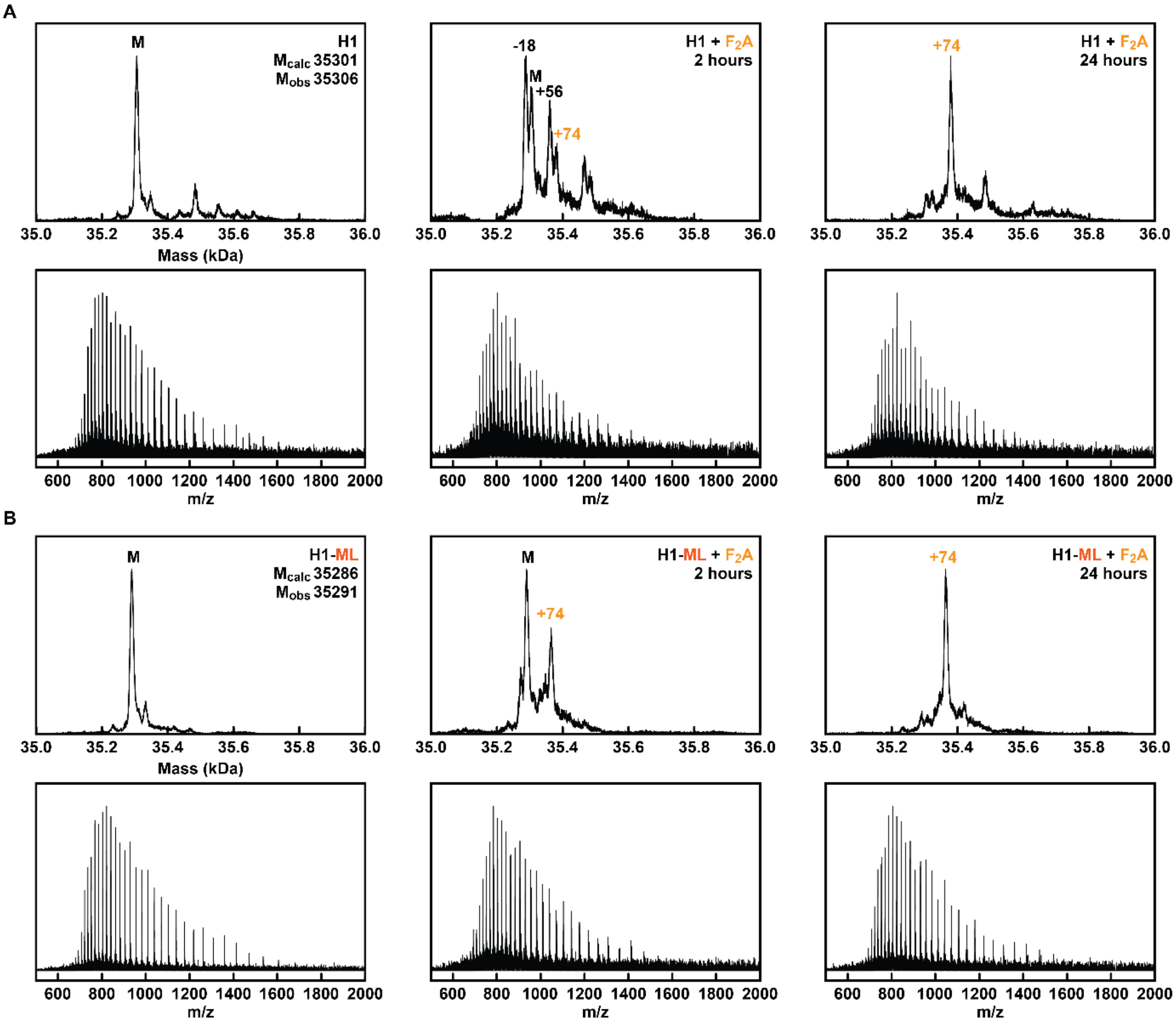


**Figure S8:** Raw and deconvoluted masses from UPLC-MS spectra for H1 (**A**) or H1-ML (**B**) before and during $F_2A$ incubation.

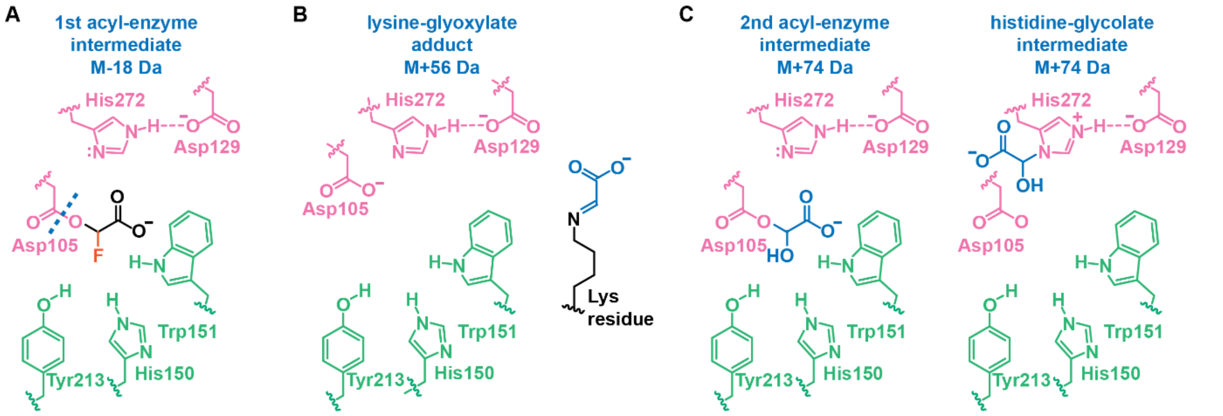


**Figure S9: A-C:** Potential structures of tentatively assigned species observed in UPLC-MS studies following the incubation of H1-variants with $F_2A$. **A:** The M-18 Da species arises from the fragmentation of the ester bond in the initial acyl-enzyme intermediate. **B:** The M+56 Da species results from the condensation of an unknown lysine residue (or N-terminal amine) with glyoxylate. **C:** The M+74 species could indicate the presence of the second acyl-enzyme intermediate or the glycolation of a nucleophilic residue in the active site. As representative example, the structure for the glycolated His272 is depicted.

### 2. Supporting Tables

**Supporting Table S1:** Apparent initial rates for FAcD-H1 and H1-H272A. Numbers corresponding to **Fig. 2A** are given here. Wildtype H1 rates were determined from biological duplicates. Rates are based on $F^-$ release measured by $^{19}$F-NMR spectroscopy (see *Experimental* for details). Standard deviations are given. Traces of $F^-$ (<1%) detected in $F_2P$ assays are ascribed to substrate or buffer impurities. Note: n.d. stands for 'not detected'.

| Enzyme | FA | FP | $F_2A$ | $F_2P$ |
|---|---|---|---|---|
| | $v_{0,app}$ ($s^{-1}$) | $v_{0,app}$ ($s^{-1}$) | $v_{0,app}$ ($s^{-1}$) | $v_{0,app}$ ($s^{-1}$) |
| **H1** | 32 ± 4.5 | 1.9 ± 0.36 | $3.5 \times 10^{-3} \pm 9.4 \times 10^{-4}$ | n.d. |
| **H1-H272A** | n.d. | n.d. | n.d. | n.d. |

**Supporting Table S2:** The transformation efficiency was based on the colony-forming units (cfus) and the total culture volume. For transformations into NEB 10-beta, a negative control was included in the Golden Gate restriction-ligations which contained no insert DNA. This control gave an estimate of background, i.e. colonies harboring an empty vector. Note: n.a. stands for 'not applicable'.

| | Host | Method | # cfus on diluted plate | # transformants (corrected for dilution) | Coverage of theoretical genetic diversity | Background |
|---|---|---|---|---|---|---|
| **Lib-D1** | NEB 10-beta | Chemical transformation | 50 | $2.0 \times 10^4$ | $2.0 \times 10^4 / 32^2 =$ 20 | $4.0 \times 10^2$ (2%) |
| **Lib-D2** | NEB 10-beta | Chemical transformation | 36 | $4.0 \times 10^3$ | $4.0 \times 10^3 / 32^2 =$ 3.9 | $1.1 \times 10^2$ (3%) |
| **Lib-D3** | NEB 10-beta | Chemical transformation | 71 | $7.9 \times 10^3$ | $7.9 \times 10^3 / 32^2 =$ 7.7 | $1.1 \times 10^2$ (1%) |
| **Lib-T1** | NEB 10-beta | Electroporation | 392 | $4.4 \times 10^5$ | $4.4 \times 10^5 / 32^3 =$ 13 | $4.4 \times 10^2$ (0.1%) |
| **Pop-D1** | BL21(DE3) | Chemical transformation | 17 | $3.4 \times 10^2$ | n.a. | n.a. |
| **Pop-D2** | BL21(DE3) | Electroporation | 42 | $4.2 \times 10^6$ | n.a. | n.a. |
| **Pop-D3** | BL21(DE3) | Electroporation | 143 | $1.4 \times 10^7$ | n.a. | n.a. |
| **Pop-T1** | BL21(DE3) | Electroporation | 79 | $7.9 \times 10^6$ | n.a. | n.a. |

**Supporting Table S3**: Sanger sequencing results of 101 random single colonies throughout the selection campaigns. Variants chosen for characterization are marked in green. Codons that were not randomized are blocked off in gray. See also the results from whole-plasmid sequencing (**Fig. 3C-D** and **Supporting Fig. S4**).

| Population | Clone no. | S147 codon | S147 | W180 codon | W180 | Q245 codon | Q245 | M246 codon | M246 | Notes, additional substitutions/mutations |
|---|---|---|---|---|---|---|---|---|---|---|
| **Pop-D1** FP plate reader screening, random clones with growth phenotype | 1 | | | TGG | W | AGT | S | | | |
| | 2 | | | TGG | W | GCG | A | | | |
| | 3 | | | TGG | W | GCG | A | | | R230H (C**G**C->C**A**C), F83F (TTC->TT**T**) |
| | 4-5 | | | TGG | **W** | GGG | **G** | | | |
| | 6 | | | TGG | W | TGG | W | | | A146T (**G**CA->**A**CA), Q191K (**C**AG->**A**AG), A280A (GC**T**->GC**C**) |
| **Pop-D2** FP plate reader screening, random clones with growth phenotype | 1 | TGG | W | | | GCG | A | | | |
| | 2 | TCG | S | | | GCG | A | | | |
| | 3 | GTG | **V** | | | TGG | **W** | | | |
| | 4 | TCG | S | | | GAG | E | | | |
| **Pop-D1 Round 2** FP selection | 1-6 | | | TGG | **W** | GCG | **A** | | | |
| **Pop-D1 Round 1** FA selection | 1-6 | | | TGG | **W** | CGG | **R** | | | |
| **Pop-D2 Round 1** FP selection | 1-11 | TTG | L | | | TGG | W | | | |
| | 12 | TCG | S | | | AAG | K | | | |
| | 13 | GGG | G | | | TGT | C | | | |
| | 2 clones are removed due to ambiguous or failed sequencing results | | | | | | | | | |
| **Pop-D2 Round 2** FP selection | 1-7 | TTG | **L** | | | TGG | **W** | | | |
| | 3 clones are removed due to ambiguous or failed sequencing results | | | | | | | | | |
| **Pop-D3 Round 1** FP selection | 1-3 | | | | | CAG | Q | ATG | M | |
| | 4 | | | | | ACG | T | ATG | M | |
| | 5 | | | | | AAG | K | ATG | M | |
| | 6-7 | | | | | GTG | V | ATG | M | V232V (GT**T**-> GT**G**) |
| | 8 | | | | | AGG | R | ATG | M | C176C (TG**C**->TG**T**), S79S (TC**A**->TC**C**) |
| | 9 | | | | | GCG | A | ATG | M | Q229Q (CAG-> CA**A**) |
| | 10 | | | | | GAG | E | ATG | M | |
| | 11 | | | | | ATG | **M** | CTG | **L** | |
| | 12 | | | | | TTT | **F** | ATT | **I** | |
| | 13 | | | | | TTG | L | TTG | L | Poor seq. quality |
| | 14 | | | | | ACG | T | GGG | G | |
| | 1 clone is removed due to ambiguous or failed sequencing results | | | | | | | | | |
| **Pop-D3 Round 2** FP selection | 1-10 | | | | | GTG | **V** | ATG | **M** | V232V (GT**T**-> GTG) |
| **Pop-T1 Round 1** FP selection | 1 | TCG | S | | | AGT | S | ATG | M | |
| | 2 | CCG | P | | | TTG | L | ATG | M | |
| | 3 | CGG | **R** | | | CCG | **P** | CTG | **L** | A46V (G**C**C -> G**T**C) |
| | 4 | TAG | STOP | | | CCG | P | TTG | L | R81H (C**G**C->C**A**C) |
| | 5 | TAG | STOP | | | TTG | L | CTG | L | Base (G) insertion at G247, causing frameshift |
| | 6 | GCG | A | | | CAG | Q | ATG | M | Poor seq. quality |
| | 7-13 | Partial (2 out of 7) or complete (5 out of 7) gene deletion | | | | | | | | |
| | 7 clones are removed due to ambiguous or failed sequencing results | | | | | | | | | |
| **Pop-T1 Round 2** FP selection | 1-5 | TTG | **L** | | | GTG | **V** | ATG | **M** | |
| | 6 | GCG | A | | | CAG | Q | ATG | M | Poor seq. quality |
| | 7-18 | Gene deletion | | | | | | | | |
| **Pop-T1 Round 1** $F_2$P selection | 1 | CCG | **P** | | | TTG | **L** | ATG | **M** | |
| | 2-3 | CGG | **R** | | | CAT | **H** | CTG | **L** | |
| | 4 | CGG | **R** | | | TTG | **L** | ATG | **M** | |

**Supporting Table S4:** Panel of H1 variants chosen for characterization and their origins. The carbon source that was used during selections is given in brackets**.**

| Defluorinase shorthand | Substitutions in H1 enzyme | Library source and rationale |
|---|---|---|
| **H1-A** | Q245A | Pop-D1 round 2 (FP), most common variant. Also found in initial plate reader assessment. |
| **H1-LW** | S147L-Q245W | Pop-D2 round 1 and 2, most common variant. |
| **H1-V** | Q245V | Pop-D3 round 2 (FP), most common variant. |
| **H1-LV** | S147L-Q245V | Pop-T1 round 2 (FP), most common variant after wildtype. |
| **H1-R** | Q245R | Pop-D1 round 1 (FA), sole surviving variant after FA selection. |
| **H1-G** | Q245G | Pop-D1 (FP), found twice in initial plate reader assessment with growth phenotype. |
| **H1-VW** | S147V-Q245W | Pop-D2 (FP), found in initial plate reader assessment with growth phenotype. |
| **H1-FI** | Q245F-M246I | Single colony from Pop-D3 round 1 (FP) that had an uncommon Met246 substitution. |
| **H1-ML** | Q245M-M246L | Single colony from Pop-D3 round 1 (FP) that had a Met246 substitution to Leu, a substitution that we also identified in the $F_2P$ selection. |
| **H1-RPL (+A46V)** | A46V-S147R-Q245P-M246L | Pop-T1 round 1 (FP), survivor with similar substitutions to those identified in $F_2P$ selections. |
| **H1-RL** | S147R-Q245L | Pop-T1 round 1 ($F_2P$) survivor. |
| **H1-RHL** | S147R-Q245H-M246L | Pop-T1 round 1 ($F_2P$) survivor. |
| **H1-PL** | S147P-Q245L | Pop-T1 round 1 ($F_2P$) survivor. Also present in Pop-T1 round 1 (FP). |

**Supporting Table S5:** Mass analysis of purified enzymes**.** Q-TOF MS results confirm the identity of several purified H1 variants in 50 mM $Na_2HPO_4$ buffer and/or MilliQ. The H272A and PL proteins are present with the expected mass, but also reveal a peak with a mass difference of +176 to 181 Da. We ascribe this to a possible N-terminal gluconoylation of the His-tag[1], although this was not investigated in detail. It is worth noting that these two variants were produced at 18 ˚C instead of the usual 37 ˚C.

| Defluorinase variant | Expected (M-Met, Da) | Observed (M-Met, Da) |
|---|---|---|
| **H1** | 35300.75 | **35305.2** |
| **H1-H272A** | 35234.69 | **35240.6**, 35417.0 (+176) |
| **H1-A** | 35243.70 | **35247.0** |
| **H1-LW** | 35384.91 | **35387.6** |
| **H1-V** | 35271.75 | **35275.4** |
| **H1-LV** | 35297.83 | **35302.7** |
| **H1-R** | 35328.80 | **35335.0** |
| **H1-G** | 35229.67 | **35233.4** |
| **H1-VW** | 35370.89 | **35374.1** |
| **H1-FI** | 35301.76 | **35306.4** |
| **H1-ML** | 35285.78 | **35288.7** |
| **H1-RPL (+A46V)** | 35348.86 | **35353.5** |
| **H1-RL** | 35354.89 | **35359.1** |
| **H1-RHL** | 35360.83 | **35366.1** |
| **H1-PL** | 35295.82 | **35301.2**, 35482.0 (+181) |

**Supporting Table S6:** Apparent initial rates and turnover numbers (TON) of purified FAcD-H1 variants. Numbers corresponding to **Fig. 4B** are given here. Wildtype H1 parameters were determined from biological duplicates. H1 variant parameters were determined from technical duplicates unless indicated otherwise. All parameters are based on $F^-$ release measured by $^{19}F$-NMR spectroscopy (see *Experimental* for details). Standard deviations are given. Note: n.d. stands for 'not detected'.

| | FA | FP | $F_2A$ |
|---|---|---|---|
| **Enzyme** | $v_{0,app}$ ($s^{-1}$) | $v_{0,app}$ ($s^{-1}$) | **TON** |
| **H1** | 32 ± 4.5 | 1.9 ± 0.36 | 307 ± 81 (bio. dup. with tech. replicate) |
| **H1-H272A** | n.d. | n.d. | n.d. |
| **H1-A** | 63 ± 0.82 | 3.2 ± 0.21 | 267 ± 37 |
| **H1-LW** | 12 ± 3.8 | 1.0 ± 0.090 | 75 ± 2 |
| **H1-V** | 47 ± 0.16 | 2.0 ± 0.093 | 174 ± 19 |
| **H1-LV** | 18 ± 3.5 | 1.2 ± 0.080 | 213 ± 50 |
| **H1-R** | 38 ± 5.3 | 2.1 ± 0.33 | 147 ± 71 |
| **H1-G** | 26 ± 2.1 | 1.1 ± 0.17 | 783 ± 21 |
| **H1-VW** | 5.7 ± 1.6 | 0.72 ± 0.079 | 116 ± 2 |
| **H1-FI** | 38 ± 5.8 | 2.1 ± 0.11 | 256 ± 86 |
| **H1-ML** | 5.9 ± 0.76 | 0.047 ± 0.0029 | 1900 ± 161 (tech. tripl) |
| **H1-RPL (+A46V)** | 1.3 ± 0.20 | 0.022 ± 0.00028 | 723 ± 27 |
| **H1-RL** | 25 ± 0.05 | 1.3 ± 0.071 | 232 ± 53 |
| **H1-RHL** | 4.7 ± 0.25 | 0.039 ± 0.0044 | 1140 ± 95 |
| **H1-PL** | 2.1 ± 0.54 | 0.052 ± 0.0065 | 134 ± 22 (tech. tripl) |

**Supporting Table S7:** Contents of Minimal Medium with Vitamins (MMV) used for growth-based screening and selection. The salt solution (adapted from literature[2]) was autoclaved for 30 min at 121 ˚C. Prior to use, filtered vitamin solution (of 1000× stock), filtered trace metals solution (of 200× stock) and various (fluorinated) carbon sources were added.

| **Salt solution (1×)** | | |
|---|---|---|
| **Compound** | **g/L** | **mM** |
| $Na_2HPO_4 \cdot 7\ H_2O$ | 4.0 | 14.8 |
| $KH_2PO_4$ | 1.4 | 10.3 |
| $MgSO_4 \cdot 7\ H_2O$ | 0.4 | 1.7 |
| $(NH_4)_2SO_4$ | 1.0 | 7.6 |
| **Trace metals solution (200×)**[3] | | |
| **Compound** | **g/L** | **mM** |
| $Ca(NO_3)_2$ | 780 | 4.75 |
| $FeSO_4 \cdot 7\ H_2O$ | 200 | 0.72 |
| $ZnSO_4 \cdot 7\ H_2O$ | 10 | 0.035 |
| $H_3BO_4$ | 10 | 0.16 |
| $CoCl_2 \cdot 6\ H_2O$ | 10 | 0.042 |
| $CuSO_4 \cdot 5\ H_2O$ | 10 | 0.040 |
| $MnSO_4 \cdot 1\ H_2O$ | 4 | 0.024 |
| $Na_2MoO_4 \cdot 2\ H_2O$ | 3 | 0.012 |
| $NiCl_2 \cdot 6\ H_2O$ | 2 | 0.008 |
| $Na_2WO_4 \cdot 2\ H_2O$ | 2 | 0.006 |
| **Vitamin solution (1000×)** in 50/50 ethanol/demi water[2] | | |
| **Compound** | **g/L** | **mM** |
| Biotin | 2.2 | 0.01 |
| Folic acid | 2.2 | 0.005 |
| p-aminobenzoic acid | 200 | 1.46 |
| Riboflavin | 220 | 0.58 |
| Panthothenic acid | 440 | 2.01 |
| Niacinamide | 440 | 3.60 |
| Pyridoxine · HCl | 440 | 2.14 |
| Thiamine · HCl | 440 | 1.30 |
| **Carbon sources** (1M - 4M stocks, neutralized to pH ≈ 7 – 8 with NaOH when needed) | | |
| **Compound** | **Final concentration** | **Sterilization** |
| Glucose | 11 mM (= 0.2 w/v %) | Filtered |
| Sodium acetate | 75 mM | Filtered |
| Sodium fluoroacetate | 100 mM | None |
| Sodium DL-lactate | 50 mM (racemic) | Autoclaved |
| Sodium pyruvate | 50 mM | None |
| Sodium glycolate | 100 mM | Autoclaved |
| Sodium 2-fluoropropionate | 100 mM (racemic) | None |
| Sodium 2,2-difluoropropionate | 100 mM | None |

**Supporting Table S8**: DNA fragments required for generating H1 NNK libraries via oePCR. Templates and primer pairs are indicated.

| **Lib-D1** | **Template(s)** | **Primer pair** | **Fragment size** |
|---|---|---|---|
| Fragment D1-A | pACYC_dehH1 | *dehH1_GG_for_BsaI* + *dehH1_W180_rev* | 551 bp |
| Fragment D1-B | pACYC_dehH1 | *dehH1_W180NNK_for* + *dehH1_Q245_rev* | 217 bp |
| Fragment D1-C | pACYC_dehH1 | *dehH1_Q245NNK_for* + *dehH1_GG_rev_BsaI* | 189 bp |
| Intermediate fragment D1-B-C | D1-B and D1-C | *dehH1_W180NNK_for* + *dehH1_GG_rev_BsaI* | 384 bp |
| Full-length insert Lib-D1 | D1-A and D1-B-C | *dehH1_GG_for_BsaI* + *dehH1_GG_rev_BsaI* | 913 bp |
| **Lib-D2** | **Template(s)** | **Primer pair** | **Fragment size** |
| Fragment D2-A | dehH1 synthesized gene | *dehH1_GG_for_BsaI* + *dehH1_S147_rev* | 452 bp |
| Fragment D2-B | dehH1 synthesized gene | *dehH1_S147NNK_for* + *dehH1_Q245_rev* | 313 bp |
| Fragment D2-C | is the same as fragment D1-C | | |
| Intermediate fragment D2-A-B | D2-A and D2-B | *dehH1_GG_for_BsaI* + *dehH1_Q245_rev* | 746 bp |
| Full-length insert Lib-D2 | D2-A-B and D2-C | *dehH1_GG_for_BsaI* + *dehH1_GG_rev_BsaI* | 913 bp |
| **Lib-D3** | **Template(s)** | **Primer pair** | **Fragment size** |
| Fragment D3-A | dehH1 synthesized gene | *dehH1_GG_for_BsaI* + *dehH1_Q245_rev* | 746 bp |
| Fragment D3-B | dehH1 synthesized gene | *dehH1_Q245NNK_M246NNK_for* *dehH1_GG_rev_BsaI* | 189 bp |
| Full-length insert Lib-D3 | D3-A and D3-B | *dehH1_GG_for_BsaI* + *dehH1_GG_rev_BsaI* | 913 bp |
| **Lib-T1** | **Template(s)** | **Primer pair** | **Fragment size** |
| Fragment T1-A | is the same as fragment D2-A | | |
| Fragment T1-B | is the same as fragment D2-B | | |
| Fragment T1-C | is the same as fragment D3-B | | |
| Intermediate fragment T1-B-C | T1-B and T1-C | *dehH1_S147NNK_for* + *dehH1_GG_rev_BsaI* | 480 bp |
| Full-length insert Lib-T1 | T1-A and T1-B-C | *dehH1_GG_for_BsaI* + *dehH1_GG_rev_BsaI* | 913 bp |

#### 3. Supporting Discussion

During growth-based screening to validate FAcD-H1 variants, we observed inconsistent growth curves when we utilized FA and $F_2A$ as sole carbon source. In general, growing precultures of bacteria on the poor carbon source acetate resulted in less predictable growth speeds in these experiments. We found this inconsistency only when pre/main cultures were cultivated in 96-well plates with MMV and acetate, while culturing those in tubes (5 mL) under otherwise similar conditions gave the expected phenotype that correlated well with enzyme activity (see *Experimental*). This inconsistency is illustrated well when comparing growth behavior on FA of hosts producing wildtype (H1) or inactive (H1-H272A) variants (**Supporting Fig. S10** below). We also found significant differences in $F^-$ release in culture supernatant of H1-producing hosts depending on which pre/main culturing conditions were applied. This result is relevant, since growth should provide consistent readout of enzyme activity. We did not observe this inconsistency between protocols when pre/main cultures were grown on MMV with glucose (and subsequently, on FP or $F_2P$ in the final plate).

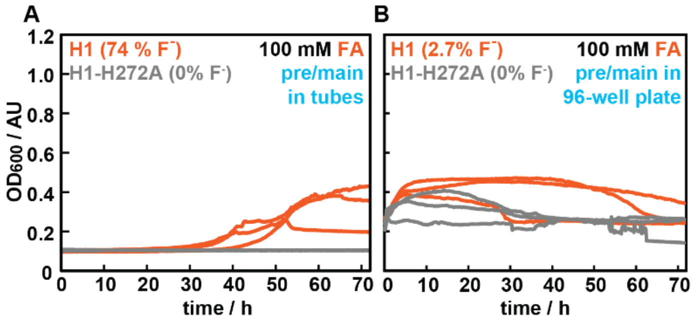


**Figure S10: Growth curves of cells producing H1 or H1-H272A when utilizing FA as sole carbon source, with $F^-$ release indicated. A:** Growth phenotype when pre/main cultures were grown in tubes. **B:** Growth phenotype when pre/main cultures were grown in 96-well plates.

With these considerations in mind, the growth curves on FA/$F_2A$ observed after cultivating the pre/main cultures in 96-well plates could have the following explanations: (1) The inducer medium added to the 96-well main culture contains 75 mM acetate. Carryover of this carbon source could explain ‘growth’ on $F_2A$, since no $F_2A$ consumption was detected ($F^-$

release, see also **Supporting Fig. S10**). (2) The $OD_{600}$ of the main cultures from MMV with acetate is likely higher than we anticipate (>0.6) upon inoculating the final plate, causing a higher starting $OD_{600}$ at t = 0 h. Due to the low culture volume in the 96-well plates used for the main culture (500 µL), measuring $OD_{600}$ is not straightforward whereas in tubes (5 mL) this is easily monitored using a portable cell density meter. On occasion we, by eye, observed variable $OD_{600}$ across different wells of the pre/main culture. (3) High FA toxicity kills the high number of stationary cells, but the turbidity of dying cells can obscure (poor) growth from survivors. Low $F^-$ release further suggests that cells are no longer in exponential phase, and are struggling to produce sufficient FAcDs.

Due to these issues, growth curves on FA and $F_2A$ for engineered FAcD-H1 variants are left largely out of consideration, and we decided to assess them based on more reliable in vitro characterization instead of optimizing the pre/main culturing conditions.

#### 4. Experimental

**Safety Statement:** No unexpected or unusually high safety hazards were encountered. Sodium fluoroacetate was handled with extra attention to safety (often in dilute, pH-neutralized aqueous solutions) and stored away when unused.

**Materials:** Chemicals were purchased from *abcr*, *BLD-Pharm*, *TCI Europe*, and *Sigma-Aldrich*, and used without purification unless noted otherwise. In particular, sodium monofluoroacetate (99%) (FA), sodium 2,2-difluoroacetate (97%) ($F_2A$), and sodium 2,2-difluoropropionate (95%) ($F_2P$) were purchased from ABCR, and 2-fluoropropionic acid (97%) (FP) was purchased from BLD-Pharm. *Escherichia coli* strains NEB 10-beta and BL21(DE3) (*New England Biolabs*) were used for cloning and expression/selection experiments, respectively. Bacteria were cultured in Lysogeny broth (LB) medium during non-selective conditions, or Super Optimal Broth (with Catabolite repression) (SOB, SOC) medium during bacterial cell transformation. During selections, bacteria were cultured in a chemically defined Minimal Medium with Vitamins (MMV), supplemented with trace metals, vitamin solution, and various (fluorinated) carbon sources (**Supporting Table S7**). Standard cloning primers and sequencing primers were synthesized by *Eurofins Genomics* (Germany). Primers with degenerate codons (NNK) were synthesized by *Biolegio* (The Netherlands) and *Eurofins Genomics* (Germany). The synthetic gene for FAcD-H1 (dehH1) was ordered from *Twist Bioscience* (USA). Plasmid isolation kits (QIAprep Spin Miniprep Kits) and PCR and gel clean-up kits (QIAquick PCR Purification Kit, QIAquick Gel Extraction Kit) were purchased from *QIAGEN* (Germany) and used according to the manufacturer's instructions. Sanger sequencing of plasmids and PCR products as well as whole plasmid sequencing (WPS, by *Oxford Nanopore* technology) was carried out by *Eurofins Genomics* (Germany). In silico cloning and sequence analysis was performed using *Snapgene* (V1.1.3) and *Geneious* (V8.1.9) software. Phusion® High Fidelity DNA Polymerase, GC buffer, deoxynucleotide (dNTP) solution mix, Hi-T4 DNA ligase, T4 DNA ligase reaction buffer, dimethyl sulfoxide (DMSO), BsaI-HFv2, DpnI and CutSmart® buffer were purchased from *New England Biolabs*. Ni-NTA resin (Ni Sepharose 6 Fast Flow) and PD MiniTrap™ G-25 columns were purchased from *Cytiva* (Germany). Precast SDS PAGE gels (SurePAGE™, Bis-Tris, 10x8, 12%, 15 wells) and Tris-MOPS-SDS Running Buffer Powder were purchased from *GenScript* (USA). InstantBlue® Coomassie Protein Stain was purchased from *Abcam* (United Kingdom). Electroporation cuvettes (2 mm) were purchased from *Fisher Scientific* (The Netherlands).

**Methods**: The concentration of DNA in solutions was determined based on the absorption at 260 nm on a Thermo Scientific Nanodrop 2000 UV-Vis spectrophotometer. The concentration of protein in solutions was determined based on the absorption at 280 nm on the same apparatus. Molar extinction coefficients at 280 nm and molecular weight of proteins were calculated using the ProtParam Expasy web server (https://web.expasy.org/protparam/).

Cellular density by means of optical density at 600 nm ($OD_{600}$) was measured on an Ultrospec 10 Cell Density Meter (*Biochrom*) in cuvettes or directly from 5 mL cultures. Plate reader growth assays were recorded on a Synergy H1 microplate reader (*BioTek*). $^{19}$F-NMR spectra were recorded on a Bruker 400 in 10% $D_2O$ (400 MHz). $^{19}$F-NMR spectra were processed and integrated using MestReNova (V12.0.0) software.

UPLC-MS analysis was performed on an Acquity UPLC system (*Waters*) coupled to a quadrupole/time-of-flight (QToF) mass spectrometer (*Waters*) equipped with a PDA detector. The protein samples were injected on a reversed phase ACQUITY UPLC BEH300 C4 1.7 µm column (2.1 mm×150 mm) and the eluent system employed a combination of 0.1% formic acid in MilliQ (A) and 0.1% formic acid in acetonitrile (B) at a flow rate of 0.3 mL/min. The gradient started with 90% A and 10% B for 2 minutes, then varied linearly from 10 to 50% B (v/v) from 2-10 minutes, 50 to 95% B from 10-11 minutes, kept at 95% B from 11-13 minutes, returning to 5% B from 13-13.1 min, re-equilibration to 5% B from 13.1-20 minutes. The sample injection volume was 3 µL of ca. 3 µM protein. Mass spectra were obtained in the ESI-positive ion mode over a mass range between 500 to 2000 Da.

LC-MS/MS analysis was performed on digested, reconstituted peptides on an Ultimate 3000 RSLC chromatography system (*Thermo Scientific*) coupled to an Exploris 480 Mass Analyser (*Thermo Scientific*). Peptides were loaded into a C18 Trap cartridge (Acclaim Pepmap C18 Reversed Phase Trap Cartridge, 5 µm, 0.3 mm I.D. × 5 mm L., *Thermo Scientific*) and then separated on a reversed phase 1.9 µm nano-LC column (PepSep C18 Repro-Sil AQ 75 µm I.D.× 40 cm, *Bruker*). The eluent system employed a combination of 0.1% formic acid in MilliQ (A) and 0.1% formic acid in 80% acetonitrile (B). The gradient started from 95% A and 5% B, to 35% solvent B over 60 minutes at a flow rate of 0.3 mL/min. Sample injection volume was 2 µL. Eluted peptides were ionized by online nano-electrospray. Data dependent acquisition (DDA) mode was used to obtain high resolution master scans from 385 m/z to 1540 m/z or 250 m/z to 1000 m/z, at a resolution of 120,000 at m/z 200. The ions at +2 to +6 charge states were then selected according to their abundance and fragmented by HCD at 30% normalized collision energy. The resulting fragment ions were measured by data-dependent (dd) MS2 scans at a resolution of 15,000.

LC-MS/MS data was analyzed using PEAKS Studio 12.5 (*Bioinformatics Solutions*) with a MS1 error tolerance of 10 ppm for precursor ions and a fragment ion error tolerance of 0.02 Da. Charge density spectra from UPLC-MS were obtained using MassLynx (V4.1) software and subsequently deconvoluted using MagTran (V1.03)[4] software, using a mass range of 35,000-36,000 (35,200-35,500 when zoomed in), a charge range of 1-100, a S/N threshold of 5 and a max. no of species of 5.

**Competent cells:** Chemically competent cells were made using the Inoue method[5,6]. Electrocompetent cells were made using glycerol/mannitol density step centrifugation[7] and were electroporated on the same day to maximize transformation efficiency.

**Assembly of pACYC_dehH1:** All cloning and growth experiments were performed using the selection plasmid pACYC_GG, our in-house expression plasmid based on the commercially available pACYCDuet-1 whose design and construction has been described in our earlier work.[8] In brief, this vector harbors a p15A origin of replication, encodes the chloramphenicol resistance gene *cat*, and features two multiple cloning sites (MCSs) specific for type IIS restriction enzymes BsaI or Esp3I, enabling the modular exchange of two target genes via Golden Gate Assembly. Both positions are under IPTG-inducible promoters (T7), and the vector also introduces an N-terminal His-tag on the gene cloned into MCS1. In this work, MCS1 was used for all dehH1 variants and libraries, and MCS2 was left empty.

The amino acid sequence for our target enzyme, FAcD-H1 (DehH1 as given on Uniprot Q01398), was converted to a DNA sequence and codon optimized for *E. coli* using the online Codon Optimization Tool from Integrated DNA Technologies. The sequence was flanked by BsaI recognition sites and was purchased as a synthetic gene (see *Sequences*). This *dehH1* gene was then cloned into MCS1 of pACYC_GG by Golden Gate Assembly with Hi-T4 DNA ligase and BsaI-HFv2. The following thermocycler program was used for the assembly: (1) 30 cycles alternating between 37 ˚C and 16 ˚C for 5 and 10 min respectively, (2) a final digestion step at 55 ˚C for 20 min, and (3) an enzyme inactivation step at 65 ˚C for 20 min. The assembly reactions were transformed into chemically competent *E. coli* NEB10-beta cells. A single colony was picked from LB plates containing chloramphenicol (34 µg/mL) and used to inoculate 5 mL LB medium with chloramphenicol (34 µg/mL). Bacteria were grown overnight, plasmids isolated, and successful assembly of pACYC_dehH1 was confirmed by Sanger sequencing with *MCS1_Up* and *DuetDOWN1*. For protein expression and/or growth-based selections (described later), the plasmid was transformed into chemically competent *E. coli* BL21(DE3).

**Site-directed mutagenesis for making pACYC_dehH1_H272A:** Starting from plasmid pACYC_dehH1, mutagenic primers *dehH1_H272A_fw* and *dehH1_H272_rv* were used to generate a control plasmid that would encode an inactive H1 variant in which the histidine base of the catalytic triad was substituted by an alanine (H272A). The PCR reaction was performed in MilliQ water with Phusion-HF DNA polymerase, 200 μM dNTPs, 0.5 μM forward and 0.5 μM reverse primer, 1X GC buffer, 3% DMSO, and ~1 ng template DNA in a 50 μL reaction. The following thermocycler settings for this QuikChange were used: (1) initial denaturation at 95 ˚C for 3 min, (2) 16 cycles of denaturation at 95 ˚C for 30 s, annealing at 63 ˚C for 30 s, and extension at 72 ˚C for 1:30 min; (3) a final extension at 72 ˚C for 10 min. PCR product formation was confirmed by agarose gel electrophoresis, and the resulting PCR product was digested with DpnI for 1 hour at 37 ˚C to remove remaining template DNA. Next, the DNA was transformed into chemically competent *E. coli* NEB10-beta cells. A single colony was picked from LB plates containing chloramphenicol (34 μg/mL) and used to inoculate 5 mL LB medium with chloramphenicol (34 μg/mL). Bacteria were grown overnight, plasmids isolated, and successful generation of pACYC_dehH1-H272A was confirmed by Sanger sequencing with *DuetDOWN1*. For protein expression and/or growth-based selections (described later), the plasmid was transformed into chemically competent *E. coli* BL21(DE3).

**AlphaFold model and residue selection mutagenesis:** Currently, no crystal structure is available for H1. To allow visual inspection of the enzyme structure, an AlphaFold2-Multimer model[9,10] was generated of the wildtype sequence of H1, which encodes a homodimer. To do so, the open source ColabFold platform was used[11]. In total, four residues were targeted for NNK randomization. Trp180 was inspired by a work from the Wang group where, in a homologous fluoroacetate dehalogenase (RPA1163 from *Rhodopseudomonas palustris* CGA009), its substitution for smaller residues made room for bulkier α-fluorocarboxylic acids[12]. After submitting our model to the online webserver HotSpot Wizard[13], Gln245 and Ser147 were identified as functional hot spots (i.e. mutable residues not involved in catalysis, but positioned near the catalytic pocket) and thus also chosen for mutagenesis. Met246 was chosen following visual inspection of the model, as it was pointing towards the active site and was conveniently positioned next to Gln245. With epistatic interactions in mind, it was only randomized in combination with Gln245. All libraries were generated as double or triple NNK libraries, in order to search for potential epistatic effects and increase library size. The combinations of randomized residues were Lib-D1 (W180NNK + Q245NNK), Lib-D2 (S147NNK + Q245NNK), Lib-D3 (Q245NNK + M246NNK), and Lib-T1 (S147NNK + Q245NNK+ M246NNK).

**Generation of NNK libraries:** Overlap extension PCR (oePCR) with primers bearing degenerate NNK codons was employed to randomize the targeted positions. Starting from pACYC_dehH1 or the ordered dehH1 gene, two or three DNA fragments with partially overlapping ends were generated using PCR with various *NNK_for* mutagenic primers in combination with their respective *_rev* primers (see **Supporting Table S8)**. The partially overlapping fragments were subsequently used as templates and were merged together by oePCR. Some fragments were interchangeable between libraries as they randomized the same parts of the sequence. In our hands, coupling of more than two fragments by oePCR directly did not produce sufficient product. Therefore, an intermediate fragment was generated that merged two fragments together when required. In the final oePCR, amplification of an (intermediate) fragment together with its complementary partially overlapping fragment in the presence of primers *dehH1_GG_for_BsaI* and *dehH1_GG_rev_BsaI* yielded the full-length 913 bp construct (**Supporting Table S8**).

In total, three (oe)PCR reactions were required per library to create the full-length insert. The following PCR protocol was used: (1) initial denaturation at 95 ˚C for 3 min, (2) (5+)25 cycles of denaturation at 95 ˚C for 30 s, annealing at 64˚C for 30 s and extension at 72 ˚C for 15 s; (3) a final extension at 72 ˚C for 10 min. For oePCR reactions, the primers were added to the PCR reaction mixture after 5 cycles. All PCR products were separated on a 0.8 % agarose gel and excised from the gel to remove unspecific amplification products and remove template DNA. All PCR reactions were performed in MilliQ water with Phusion-HF DNA polymerase, 200 μM dNTPs, 0.5 μM forward and 0.5 μM reverse primer, 1X GC buffer, 3% DMSO, and ~1 ng template DNA in 50 μL reactions. In case of poor DNA yields, up to 8 reactions of 50 μL were pooled together prior to gel extraction, rather than increasing the number of cycles so as not to introduce PCR bias.

To assemble the selection plasmids, the partially randomized CDS was cloned into pACYC_GG using Golden Gate Assembly with Hi-T4 DNA ligase and BsaI-HFv2 as described earlier. A negative control was included that did not contain insert DNA but solely the target vector, to determine the number of background transformants which harbored empty vectors, which was generally very low (≤3%, see **Supporting Table S2**). The entirety of the restriction-ligation reactions was transformed into NEB 10-beta cells, either by heat shock into chemically competent cells for the double libraries Lib-D1, Lib-D2 and Lib-D3, or by electroporation for the triple library Lib-T1. The number of transformants was calculated based on the number of colony-forming units and final culture volume (**Supporting Table S2**). All colonies were scraped from large selective LB agar plates (~200 mL) containing chloramphenicol (34 μg/ml). Library plasmid DNA was isolated and sequenced with *DuetDOWN1*, *MCS1_Up* and whole-plasmid sequencing to verify library quality and confirm successful randomization (see also the section on *Sequencing analysis*). The isolated plasmids were stored at -20 ˚C until the start of a selection or screening experiment.

**Growth-based screening of *E. coli* producing H1 or H272A in the plate reader with preculturing in tubes:** This protocol had some minor adaptations based on the carbon source used (*vide infra*) and whether or not the pre- and main cultures were incubated in tubes or in 96-deep well plates (see next section). The overall workflow is the same, however.

Precultures of 5 mL Minimal Medium with Vitamins (MMV, see **Supporting Table S7**) with 34 µg/mL chloramphenicol and 11 mM glucose (= 0.2 w/v %) as sole carbon source were inoculated with single colonies from fresh agar plates with *E. coli* BL21(DE3) cells harboring pACYC_dehH1 or pACYC_dehH1-H272A. The precultures were incubated at 37 ˚C with moderate shaking (135 rpm) for ≈ 20 h. The following morning, main cultures of 5 mL MMV with 17 µg/mL chloramphenicol and 11 mM glucose were inoculated with 50 µL of the corresponding preculture and grown at 37 ˚C, 135 rpm, until the optical density at 600 nm ($OD_{600}$) was approximately 0.3. Then, expression of the dehH1 or dehH1-H272A coding sequence was induced by addition of IPTG (final concentration 1 mM), and expression was allowed for 3 hours at 37 ˚C, 135 rpm. During this step, the final plate assay plate was prepared, i.e. a flat-bottom 96-well plate suitable for a plate reader. In each well was added: (1) 180 µL of MMV containing 17 µg/mL chloramphenicol, 1 mM IPTG, and a variable (fluorinated) carbon source (e.g. 111 mM FP, making the final concentration 100 mM) and (2) 20 µL of the induced main culture ($OD_{600}$ ≈ 0.6). The following final concentrations were used for each tested carbon source: 100 mM FA, 100 mM racemic FP, 100 mM $F_2A$, 100 mM $F_2P$, 100 mM glycolate, 50 mM pyruvate, 50 mM racemic lactate, 75 mM acetate. Assay plates were closed with a transparent plastic lid and transferred into a Synergy H1 microplate reader that had been preheated to 30 °C. While continuously shaking (double orbital, 425 c.p.m.) at 30 °C, growth was monitored by measuring $OD_{600}$ every 10 minutes from the bottom of the wells for a period up to 110 hours (generally around 72 hours was sufficient).

Depending on the carbon source under study in the plate reader, the proceedings regarding the preculture and main culture are slightly different. When growth on C3 carbon sources (FP, $F_2P$, pyruvate, or lactate) was assessed, precultures and main cultures were supplemented with 11 mM glucose as sole carbon source, and followed the proceedings above. However, when growth on C2 carbon sources (FA, $F_2A$, glycolate, or acetate) was assessed, precultures and main cultures were supplied with 75 mM acetate as sole carbon source instead, in order already upregulate the expression of genes involved in the glyoxylate shunt.[14] In this case, incubation times also had to be increased due to slower growth rates: the precultures were incubated for at least 26 hours (up to 48 hours), the main culture was incubated overnight (≈ 16 hours) after inoculation, and the induced main culture was allowed to express for 3 – 4 hours. Additionally, the volume used from the preculture to inoculate the main culture was increased from 50 µL to 100 µL if necessary.

**Growth-based screening of *E. coli* producing FAcD-H1 variants in the plate reader with preculturing in 96-well plates:** When investigating the growth phenotype of the panel with enriched FAcD-H1 variants after selections, the abovementioned screening protocol was also applied, with some modifications. To exclude any changes to host fitness that could have been the result of spontaneous background mutations in the *E. coli* genome or the selection plasmid backbone, the H1 CDS of the hits was recloned prior to growth screening. Specifically, they were amplified by routine PCR with primers *dehH1_GG_for_BsaI* and *dehH1_GG_rev_BsaI*, gel extracted, and cloned into fresh pACYC_GG vectors by Golden Gate Assembly (as described earlier). These products were transformed into NEB10-beta, isolated plasmids were confirmed by Sanger sequencing with *DuetDOWN1* or whole-plasmid sequencing, and subsequently transformed into BL21(DE3). Then, recloned variants were cultured in 96-well plates (instead of tubes) and assessed in the plate reader for their growth on 75 mM acetate, 100 mM FA, 100 mM $F_2A$, 50 mM racemic lactate, 100 mM racemic FP, and 100 mM $F_2P$.

The following growth-screening protocol was applied to check the growth characteristics of single colonies harboring FAcD-H1 variants before and after selection. A 96-deep well plate (pre-plate) filled with 500 µL Minimal Medium with Vitamins (MMV, **Supporting Table S7**) containing 34 µg/mL chloramphenicol and 11 mM glucose (= 0.2 w/v %) was inoculated with single colonies from freshly streaked or transformed *E. coli* BL21(DE3) cells, harboring a selection plasmid (pACYC_dehH1) or a mutated gene variant thereof. For FAcD-H1 variants after selection, colonies were picked in triplicate (technical replicates). The 96-deep well plate was incubated overnight (≈ 18 hours) at 37 °C while shaking at 750 rpm (*Titramax 1000 & Incubator 1000, Heidolph*). The next morning, 25 µL of the densely grown overnight cultures was used to inoculate a new 96-deep well plate (main plate) containing 500 µL MMV, 17 µg/mL chloramphenicol and 11 mM glucose. After incubating the main culture for 3 hours at 37 °C, 750 rpm, expression was induced by addition of 16.5 µL MMV with 17 µg/mL chloramphenicol, 11 mM glucose, and 30 mM IPTG (final concentration 1 mM IPTG). The plate was incubated at 37 °C, 750 rpm for another 3 hours to allow expression. Then, transparent 96-well assay plates (flat-bottom) were set up as before, by adding 20 µL of the induced main culture (OD ≈ 0.6) to 180 µL of MMV containing 17 µg/mL chloramphenicol, 1 mM IPTG, and a variable (fluorinated) carbon source. Growth was monitored in the plate reader as described earlier.

Also in this case, the protocol has a number of adaptations when growth on C2 carbon sources (FA, $F_2A$, or acetate) was assessed. In those cases, the pre-plate and main plate were supplied with 75 mM acetate as sole carbon source instead of glucose. The pre-plate was incubated for 40 – 48 hours, the main culture was incubated overnight (≈ 16h) after inoculation, and the induced main culture (induced by addition of 16.5 µL MMV with 17 µg/mL chloramphenicol, 75 mM acetate, and 30 mM IPTG, giving a final concentration of 1 mM IPTG) was allowed to express for 4 hours.

**Growth phenotype categorization:** To depict the growth characteristics of each variant in a comprehensive manner (i.e. a single figure), $OD_{600}$ after 50 hours of growth was depicted for each well with a white ($OD_{600} = 0$) to blue ($OD_{600} \geq 1$) color gradient. Cells were considered to have a positive growth phenotype on FP when $OD_{600} > 0.2$ at this time, and double-checked by visual inspection of each growth curve relative to its corresponding wildtype and inactive (H1-H272A) control. It should be noted that some clones in **Supporting Figure S3** showed a growth phenotype early on and were subjected to $^{19}$F-NMR analysis after 24 hours of growth ($F^-$ release ranged between 14-30%); therefore, these clones did not have an $OD_{600}$ value after 50 hours of growth. Nevertheless, these clones were counted towards the number of clones with a positive growth phenotype, and their last measured $OD_{600}$ value (after 24 hours) was depicted with an asterisk.

$OD_{600}$ values depicted in **Supporting Figure S3** are from single measurements; values in **Fig. 4A** are averaged from technical triplicates (corresponding growth curves are given in **Supporting Figure S6**).

**Preparation of $^{19}$F-NMR samples from cultures:** During culturing, fluoride release from successful fluorinated substrate conversion was periodically measured by $^{19}$F-NMR spectroscopy. For selection cultures, 110 µL of a 5 mL culture was removed by pipetting. For wells of interest in a 96-well plate, the plate reader was paused after 24, 72, or 110 hours of incubation, 110 – 200 µL was removed from wells of interest by pipetting, and the plate reader was resumed. To prepare the samples for $^{19}$F-NMR, cells were spun down by centrifugation (10 min, 13,000 rpm) and 100 µL of supernatant was mixed with 350 µL MilliQ and 50 µL $D_2O$. The mixture was transferred to an NMR tube and $F^-$ release was determined by $^{19}$F-NMR (see *$^{19}$F-NMR analysis*).

**Growth-based selections of FAcD-H1 variants from libraries in liquid media:** Chemically competent or freshly prepared electrocompetent *E. coli* BL21(DE3) were transformed with library plasmids of Lib-D1, Lib-D2, Lib-D3, or Lib-T1. Following recovery, a fraction of the cells was plated on LB agar plates with 34 µg/mL chloramphenicol to calculate the number of transformants (**Supporting Table S2**). All transformants were grown directly from the recovering cells by topping the medium up to 4 mL SOB, adding 34 µg/mL chloramphenicol to kill cells devoid of a plasmid, and by growing the cultures for ~20h at 30 °C, 135 rpm. Precultures of controls BL21(DE3) pACYC_dehH1 or pACYC_dehH1-H272A were inoculated in 4 mL LB with 34 µg/mL chloramphenicol and grown overnight at 37 °C, 135 rpm. These controls were grown alongside the selection cultures under the same conditions. Following incubation of the precultures, main cultures of 5 mL MMV and 17 µg/mL chloramphenicol, supplemented either with 11 mM glucose (Glc) or with 50 mM acetate (Ac) as carbon source, were inoculated with 50 µL – 200 µL of the corresponding preculture. Main cultures were incubated at 37 °C, 135 rpm until an OD ≈ 0.2 was reached. For the Glc cultures, this was after approximately 3 – 4 hours; for the Ac cultures, this required approximately 7 – 8 hours. Gene expression was induced by addition of 1 mM IPTG and main cultures were incubated for 3 more hours at 37 °C, 135 rpm. During this incubation step, selection medium was freshly prepared, containing 5 mL MMV, 1 mM IPTG, 17 µg/mL chloramphenicol, and a variable fluorinated carbon source: 100 mM FA, 100 mM $F_2A$, 100 mM FP, or 100 mM $F_2P$.

To start the selections, the induced cultures were diluted 1:100 in selection medium with FP or $F_2P$ from Glc main cultures, and were diluted 1:50 in selection medium with FA or $F_2A$ from Ac main cultures. Selection cultures were grown at 30 °C (for Lib-D2, Lib-D3, or Lib-T1) or 37 °C (for Lib-D1), 135 rpm while routinely measuring $OD_{600}$ using a portable cell density meter. Since the carbon sources are poor, growth of libraries required at least 3–4 days for FP and >1 week for FA. Therefore, $OD_{600}$ values of 0.3–0.4 were generally considered to be sufficient to continue to the next passage, which would be started by diluting 1:100 in fresh selection medium. In this way, one (on FA) or two (on FP) selection rounds were performed.

During the selections, $^{19}$F-NMR samples of the supernatant would occasionally be measured (as described earlier) to check either if traces of defluorination could be detected prior to observable measurable cell density, or to confirm that observable growth was the result of active defluorination. Furthermore, selection survivors were routinely plated from the selection cultures on LB agar plates with 34 µg/mL chloramphenicol using appropriate dilutions. A 25% glycerol stock was made of the mixed populations prior to selection and after passage, to enable long-term storage.

For the selections on $F_2P$, no measurable growth was observed after 14 days incubation, so 200 µL of supernatant was plated without dilution on LB agar plates containing 34 µg/mL chloramphenicol to assess cell viability. Surviving colonies of the most promising library were collected, pooled plasmids were isolated, and sent for whole plasmid sequencing (WPS).

**Sequencing analysis throughout selections:** The change in composition of the population during the selections was monitored by Sanger sequencing (with *DuetDOWN1* or *MCS1_Up*) of single colonies, and by whole plasmid sequencing (WPS) of pooled plasmids from mixed populations. To increase plasmid yields, LB medium with 34 µg/mL chloramphenicol was used to start cultures from mixed populations, prior to selection and after each passage. After overnight growth their plasmids were isolated and sent for sequencing.

To analyze WPS data, raw reads were assembled to the reference plasmid (pACYC_dehH1) in *Geneious*, and mutated codons across randomized positions were extracted. Before quantification, invalid data was removed: as some reads sequenced the area outside of the CDS, these invalid codons ('---') were trimmed from the dataset using an Excel script. Furthermore, during one Pop-T1 selection campaign, hitchhiker cells had excised the dehH1 CDS, in which case the reads missing the CDS (49% of reads) were removed manually. After curating the data, all extracted NNK codons were translated to the respective amino acids and their ratios were determined using another Excel script. Once the composition of each population did not change significantly anymore, they were considered fully enriched.

**Protein production and purification:** Flasks containing 250 mL or 500 mL LB with 34 µg/mL chloramphenicol were inoculated with 250 µL or 500 µL, respectively, of a densely grown overnight culture of *E. coli* BL21(DE3) cells harboring the appropriate pACYC_dehH1 (or a variant thereof) plasmid. Cells were grown at 37 °C, 135 rpm until an OD of ~0.3 was reached, and gene expression was induced by adding 1 mM IPTG. Enzymes were produced overnight (~20h) at 37 °C (with the exception of H1-H272A and H1-PL, which were produced at 18 °C), 135 rpm, after which the cells were harvested by centrifugation (3,700 rpm for 20 min, 4 °C). When needed, cell pellets were stored at -20 °C until purification. Next, cell pellets were resuspended in buffer (20 mL, 50 mM $Na_2HPO_4$, pH 8, containing 1 mg/mL egg white lysozyme). The cells were then lysed by sonication for 10 min, with 5 s pulse and 5 s pause cycles at 70% amplitude, and cellular debris was removed by centrifugation (12,000 rcf for 45 min, 4 °C). The supernatant was loaded onto a Ni-NTA resin and purified according to the manufacturer's specifications. Elution fractions containing protein were pooled, concentrated, and finally stored in 50 mM $Na_2HPO_4$, pH 8 with 5% glycerol at -20 °C. The purity and identity of dehH1 variants was confirmed by SDS-PAGE (**Supporting Fig. S7**) and mass spectrometry, respectively (Q-ToF UPLC-MS, **Supporting Table S5**).

**Reaction assays for in vitro characterization of defluorination activity:** To investigate enzyme activity on FA, FP and $F_2A$, standard enzymatic reactions were performed in 1.5 mL Eppendorf tubes at 25 ˚C without shaking with a volume of 450 µL in 50 mM $Na_2HPO_4$ buffer (pH 8.0) with 10 mM substrate. Reactions with wildtype H1 were performed as biological duplicates, all other reactions were performed as technical duplicates (or in a few indicated cases, technical triplicates). Substrates were prepared as 20× stocks and adjusted to pH ~8 by addition of NaOH. 20 µM stock solutions of the FAcD-H1 variants were prepared in the same buffer. To initiate the reactions, enzyme was added: the final concentration was 0.5 µM enzyme for reactions with FA, 0.5 µM or 5 µM enzyme for reactions with FP (5 µM for poorly performing enzymes) and 1 µM enzyme for reactions with $F_2A$. Samples of 100 µL were taken at several time points (for $F_2A$ after 24 hours only) and quenched by addition to 1 volume of ice cold MeCN. Next, 250 µL MilliQ was added to increase the final volume and suppress noise, and 50 µL $D_2O$ was added for shimming. For a number of samples, instead of pure $D_2O$ 50 µL of a freshly prepared 10× stock of $NaBF_4$ in $D_2O$ was added as a qualitative internal standard (0.5 mM final concentration). Samples were mixed by vortexing, transferred to an NMR tube and analyzed by $^{19}$F-NMR immediately.

For assaying activity on $F_2P$, 450 µL reactions were set up in 50 mM $Na_2HPO_4$ buffer (pH 8.0), containing 10 mM $F_2P$, 45 µL $D_2O$, and 20 µM enzyme. Samples were mixed briefly by pipetting, transferred to an NMR tube, and incubated at 25 ˚C in a non-shaking incubator. Reaction progress was monitored by $^{19}$F-NMR, measuring after at least 24h incubation.

**$^{19}$F-NMR analysis**: Using $^{19}$F-NMR, substrate (FA, $F_2A$, FP, or $F_2P$) consumption and product ($F^-$) formation could be monitored simultaneously, both in cellular supernatant and in reaction assays. The apparent initial rate of each enzyme was calculated based on the formation of $F^-$ after a suitable timepoint, at which 5-10% conversion had occurred. $F^-$ concentration was determined based on the quantitative analysis of $F^-$ and substrate signal. The same ppm ranges were used for signal integration to prevent bias: -119.400 to -120.200 ppm for $F^-$; -216.100 to -217.000 ppm for FA; -123.500 to -124.800 ppm for $F_2A$; -172.600 to -173.400 ppm for FP; -96.500 to -98.900 ppm for $F_2P$. The spectra were referenced to the $F^-$ signal at -119.800 ppm or to the internal standard $NaBF_4$ signal at -150.200 ppm if used. Auto baseline correction (Whittaker Smoother) and auto phase correction were applied when necessary. Controls containing no enzyme or an inactive enzyme variant (FAcD-H1-H272A) provided insight into background trace levels of $F^-$ that sometimes followed from impurities in the buffer, internal standard, or substrate stocks. Corrections based on these controls were applied when necessary.

An example of calculating apparent initial rate ($v_{0,app}$) is given. The processed $^{19}$F-NMR spectrum below (**Supporting Fig. S11**) indicates the conversion of FP (10 mM) with enzyme variant VW (0.5 μM) after 30 minutes. Signals for $F^-$ at -119.80 ppm and FP at -172.95 ppm are both integrated, and total integrals are set to 100. From these values, we find 6.02% $F^-$ release and 93.98% FP, corresponding to 0.602 mM $F^-$ and 9.398 mM FP. The following formula was applied to determine the apparent initial rate: $v_{0,app}$ = [$F^-$ formed in mM] / ([enzyme concentration in mM] × time in seconds) = 0.602 mM / (0.0005 mM × 1800 seconds) = 0.7 $s^{-1}$. Duplicates were used to calculate an average and standard deviation for each entry. Turnover number (TON) corresponds to the number of moles of a substrate that a mole of catalyst can convert before becoming inactivated, so to determine those values, the following formula was applied: TON = (product in moles) / (catalyst in moles). The product in this case was $F^-$ and was determined in the same way as described above. Duplicates (or in a few indicated entries, triplicates) were used to calculate an average TON and standard deviation for each entry. All TONs were determined based on 24-hour time samples.

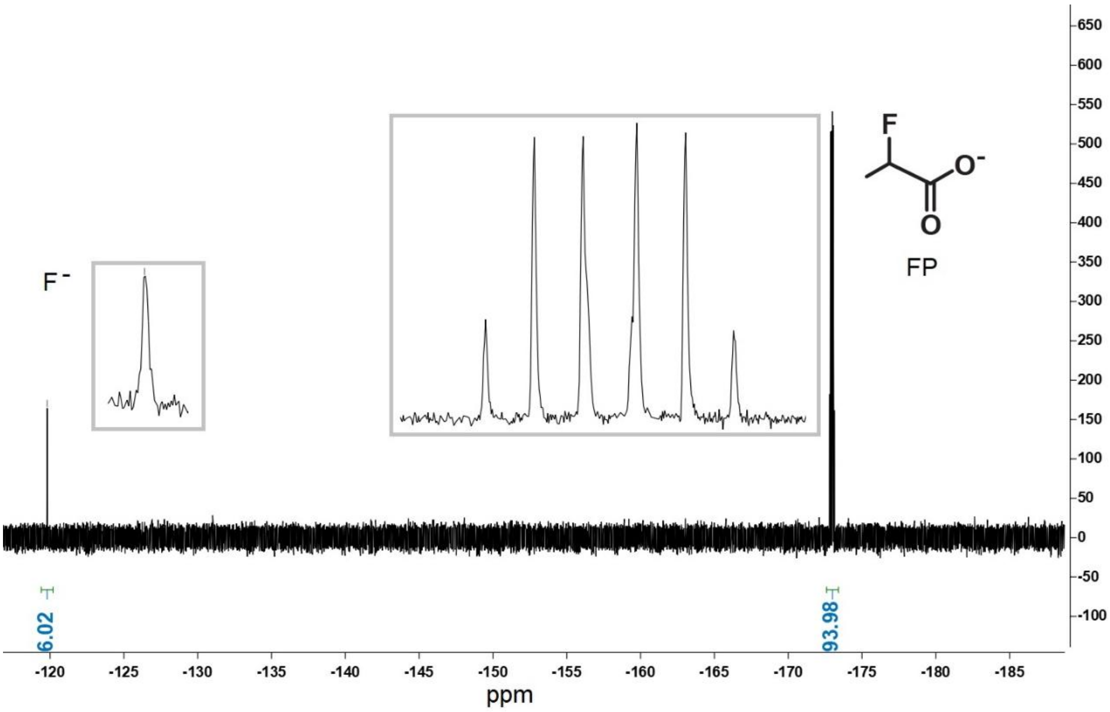


**Figure S11: $^{19}$F-NMR spectrum of enzyme variant H1-VW (0.5 μM) converting FP (10 mM).** The reaction was quenched after 30 minutes. Close-ups of the signals are given in grey boxes; they show the expected splitting.

**Enzyme modification studies upon $F_2A$ incubation:** Please refer to *Methods* for a detailed description of mass analyzer equipment. A 9 μM H1, H1-ML, and H1-H272A protein sample in 50 mM $Na_2HPO_4$ buffer (pH 8.0) were incubated with 10 mM $F_2A$ at room temperature without shaking. Negative controls included the same proteins and buffer, but were incubated without substrate. Aliquots were taken at t = 2h and t = 24h, diluted with MilliQ to 3 μM protein, and immediately subjected to UPLC-MS analysis. Aliquots of the 9 μM samples were also used for LC-MS/MS analysis by (chymo)trypsin digest, described below.

**(Chymo)trypsin digest:** Solutions with 9 μM enzyme, incubated with and without $F_2A$ for 24 hours, were denatured with 2 M urea and reduced with 10 mM TCEP for 1 h at 37 °C, and then alkylated with 20 mM iodoacetamide at room temperature in the dark for 45 minutes. Samples were diluted with 67 mM $NH_4HCO_3$ until a urea concentration of 1 M was reached. These diluted samples were digested with 1:25 (w/w) sequencing grade modified trypsin (*Promega*) at 37 °C overnight with agitation. Alternatively, they were supplemented with 10 mM $CaCl_2$ and digested with 1:20 (w/w) sequencing grade chymotrypsin (*Promega*) and incubated at 25 °C overnight. After digestion, trifluoroacetic acid (TFA) was added to reach 1% (v/v) final concentration and any precipitated detergent was removed by centrifugation. To purify the sample by solid phase extraction, the digested peptides were cleaned with C18 SPE spin tips (*Pierce*) according to the manufacturer's instructions, dried by centrifugation under vacuum, and reconstituted in 20 µL 2% MeCN and 0.1% formic acid for injection. Digested peptides were measured on an Ultimate 3000 RSLC chromatography system coupled to an Exploris 480 Mass Analyser (MS1 resolution of 120,000 and MS2 resolution of 15,000, see *Methods*). Using PEAKS Studio 12.5 software, raw files were analyzed by searching against the forward and reverse peptide sequences of the *E. coli* proteome and the target dehH1 sequences. Identified proteins were reported if the false discovery rate of the protein identifications were less than 1%. Carbamidomethylation at cysteines was set as a fixed modification; variable modifications included acetylation at N-term, deamination, and modification of Asp/His/Tyr by $C_2H_1O_3$ (+73), $C_2H_2O_3$ (+74), and $C_2H_3O_3$ (+75).

#### 5. Sequences

**Primers:** Sequences of primers used in this work, with mutations or randomizations shown underlined and bold.

| Name | Sequence (5’ → 3’) |
|---|---|
| *DuetDOWN1* | GATTATGCGGCCGTGTACAA |
| *MCS1_Up* | GGAGATATACCATGGGCAGC |
| *dehH1_H272A_fw* | TTACCTGGTGGA**<u>GC</u>**CTTCTTCGTGGACCAGTTTCC |
| *dehH1_H272_rv* | TCCACCAGGTAAGGATGCGTTCGTTG |
| *dehH1_GG_for_BsaI* | ACCGGTCTCCTGGTATGGACTTTCCGGGGT |
| *dehH1_GG_rev_BsaI* | GTTGGTCTCCCAAGTCACCCATTGCGTGCTAAG |
| *dehH1_S147NNK_for* | GAATCGCTTAGTCGCCGCA**<u>NNK</u>**TATTGGCATTGGTATTTC |
| *dehH1_S147_rev* | TGCGGCGACTAAGCGATTCGTGTTCATAAAC |
| *dehH1_W180NNK_for* | TTATGAAACATGCTTGTTTGGT**<u>NNK</u>**GGAGCCACTAAAGTGTCTG |
| *dehH1_W180_rev* | ACCAAACAAGCATGTTTCATAAAAAAAGTCGGGG |
| *dehH1_Q245NNK_for* | GGTATTTTATGGGTCTAAGGGC**<u>NNK</u>**ATGGGCCAGCTTTTCGATA |
| *dehH1_Q245_rev* | GCCCTTAGACCCATAAAATACCAATGTAGGAC |
| *dehH1_Q245NNK_M246NNK_fw* | GGTATTTTATGGGTCTAAGGGC**<u>NNKNNK</u>**GGCCAGCTTTTCGATA |

**DNA Sequence (5’ →3’) of dehH1 synthetic gene (CDS flanked by BsaI sites, codon optimized for *E. coli*)**

```
CTAACCGGTCTCCTGGTATGGACTTTCCGGGGTTCAAGAACTCTACGGTGACAGTTGACGGCGTAGATATTGCCT
ATACAGTCTCTGGAGAGGGGCCACCTGTGTTAATGCTGCATGGTTTCCCTCAGAACCGTGCGATGTGGGCGCGTG
TGGCCCCACAACTTGCTGAGCACCACACTGTCGTTTGCGCAGATCTTCGCGGCTACGGTGACAGCGACAAGCCGA
AGTGTTTACCAGATCGCTCAAATTACTCATTTCGCACATTCGCCCACGACCAATTGTGTGTGATGCGTCACCTTG
GGTTCGAGCGTTTTCATCTTGTCGGACATGATCGCGGCGGTCGCACCGGGCATCGCATGGCCTTGGACCACCCCG
AGGCCGTACTGTCCTTGACCGTCATGGATATTGTACCGACTTATGCCATGTTTATGAACACGAATCGCTTAGTCG
CCGCATCATATTGGCATTGGTATTTCTTACAACAACCTGAGCCATTTCCAGAACACATGATTGGGCAAGACCCCG
ACTTTTTTTATGAAACATGCTTGTTTGGTTGGGGAGCCACTAAAGTGTCTGATTTTGACCAACAGATGCTTAACG
CTTACCGCGAATCGTGGCGTAATCCAGCAATGATCCACGGTAGTTGCAGCGATTATCGTGCCGCCGCAACAATCG
ACTTGGAGCATGATTCGGCTGACATTCAGCGCAAGGTTGAATGTCCTACATTGGTATTTTATGGGTCTAAGGGCC
AGATGGGCCAGCTTTTCGATATTCCTGCAGAGTGGGCCAAACGCTGCAACAATACAACGAACGCATCCTTACCTG
GTGGACACTTCTTCGTGGACCAGTTTCCAGCTGAGACTTCGGAGATCTTGCTGAAGTTCTTAGCACGCAATGGGT
GACTTGGGAGACCAACTA
```

**Protein sequences:** Protein sequences of the wildtype FAcD-H1 (DehH1) and the variants further characterized in this work are given below. For clarity, deviations from the wildtype sequence are marked in bold red. The N-terminal His-tag and corresponding linker is included in the sequence and underlined. Residue numbering starts at the Met after this linker.

**H1**

MGSSHHHHHHGSGLVPRGSAGMDFPGFKNSTVTVDGVDIAYTVSGEGPPVLMLHGFPQNRAMWARVAPQLAEHHT
VVCADLRGYGDSDKPKCLPDRSNYSFRTFAHDQLCVMRHLGFERFHLVGHDRGGRTGHRMALDHPEAVLSLTVMD
IVPTYAMFMNTNRLVAASYWHWYFLQQPEPFPEHMIGQDPDFFYETCLFGWGATKVSDFDQQMLNAYRESWRNPA
MIHGSCSDYRAAATIDLEHDSADIQRKVECPTLVFYGSKGQMGQLFDIPAEWAKRCNNTTNASLPGGHFFVDQFP
AETSEILLKFLARNG*

**H1-H272A**

MGSSHHHHHHGSGLVPRGSAGMDFPGFKNSTVTVDGVDIAYTVSGEGPPVLMLHGFPQNRAMWARVAPQLAEHHT
VVCADLRGYGDSDKPKCLPDRSNYSFRTFAHDQLCVMRHLGFERFHLVGHDRGGRTGHRMALDHPEAVLSLTVMD
IVPTYAMFMNTNRLVAASYWHWYFLQQPEPFPEHMIGQDPDFFYETCLFGWGATKVSDFDQQMLNAYRESWRNPA
MIHGSCSDYRAAATIDLEHDSADIQRKVECPTLVFYGSKGQMGQLFDIPAEWAKRCNNTTNASLPGG**A**FFVDQFP
AETSEILLKFLARNG*

**H1-A (Q245A)**

MGSSHHHHHHGSGLVPRGSAGMDFPGFKNSTVTVDGVDIAYTVSGEGPPVLMLHGFPQNRAMWARVAPQLAEHHT
VVCADLRGYGDSDKPKCLPDRSNYSFRTFAHDQLCVMRHLGFERFHLVGHDRGGRTGHRMALDHPEAVLSLTVMD
IVPTYAMFMNTNRLVAASYWHWYFLQQPEPFPEHMIGQDPDFFYETCLFGWGATKVSDFDQQMLNAYRESWRNPA
MIHGSCSDYRAAATIDLEHDSADIQRKVECPTLVFYGSKG**A**MGQLFDIPAEWAKRCNNTTNASLPGGHFFVDQFP
AETSEILLKFLARNG*

**H1-LW (S147L-Q245W)**

MGSSHHHHHHGSGLVPRGSAGMDFPGFKNSTVTVDGVDIAYTVSGEGPPVLMLHGFPQNRAMWARVAPQLAEHHT
VVCADLRGYGDSDKPKCLPDRSNYSFRTFAHDQLCVMRHLGFERFHLVGHDRGGRTGHRMALDHPEAVLSLTVMD
IVPTYAMFMNTNRLVAA**L**YWHWYFLQQPEPFPEHMIGQDPDFFYETCLFGWGATKVSDFDQQMLNAYRESWRNPA
MIHGSCSDYRAAATIDLEHDSADIQRKVECPTLVFYGSKG**W**MGQLFDIPAEWAKRCNNTTNASLPGGHFFVDQFP
AETSEILLKFLARNG*

**H1-V (Q245V)**

MGSSHHHHHHGSGLVPRGSAGMDFPGFKNSTVTVDGVDIAYTVSGEGPPVLMLHGFPQNRAMWARVAPQLAEHHT
VVCADLRGYGDSDKPKCLPDRSNYSFRTFAHDQLCVMRHLGFERFHLVGHDRGGRTGHRMALDHPEAVLSLTVMD
IVPTYAMFMNTNRLVAASYWHWYFLQQPEPFPEHMIGQDPDFFYETCLFGWGATKVSDFDQQMLNAYRESWRNPA
MIHGSCSDYRAAATIDLEHDSADIQRKVECPTLVFYGSKG**V**MGQLFDIPAEWAKRCNNTTNASLPGGHFFVDQFP
AETSEILLKFLARNG*

**H1-LV (S147L-Q245V)**

MGSSHHHHHHGSGLVPRGSAGMDFPGFKNSTVTVDGVDIAYTVSGEGPPVLMLHGFPQNRAMWARVAPQLAEHHT
VVCADLRGYGDSDKPKCLPDRSNYSFRTFAHDQLCVMRHLGFERFHLVGHDRGGRTGHRMALDHPEAVLSLTVMD
IVPTYAMFMNTNRLVAA**L**YWHWYFLQQPEPFPEHMIGQDPDFFYETCLFGWGATKVSDFDQQMLNAYRESWRNPA
MIHGSCSDYRAAATIDLEHDSADIQRKVECPTLVFYGSKG**V**MGQLFDIPAEWAKRCNNTTNASLPGGHFFVDQFP
AETSEILLKFLARNG*

**H1-R (Q245R)**

MGSSHHHHHHSGLVPRGSAGMDFPGFKNSTVTVDGVDIAYTVSGEGPPVLMLHGFPQNRAMWARVAPQLAEHHT
VVCADLRGYGDSDKPKCLPDRSNYSFRTFAHDQLCVMRHLGFERFHLVGHDRGGRTGHRMALDHPEAVLSLTVMD
IVPTYAMFMNTNRLVAASYWHWYFLQQPEPFPEHMIGQDPDFFYETCLFGWGATKVSDFDQQMLNAYRESWRNPA
MIHGSCSDYRAAATIDLEHDSADIQRKVECPTLVFYGSKG**R**MGQLFDIPAEWAKRCNNTTNASLPGGHFFVDQFP
AETSEILLKFLARNG*

**H1-G (Q245G)**

MGSSHHHHHHSGLVPRGSAGMDFPGFKNSTVTVDGVDIAYTVSGEGPPVLMLHGFPQNRAMWARVAPQLAEHHT
VVCADLRGYGDSDKPKCLPDRSNYSFRTFAHDQLCVMRHLGFERFHLVGHDRGGRTGHRMALDHPEAVLSLTVMD
IVPTYAMFMNTNRLVAASYWHWYFLQQPEPFPEHMIGQDPDFFYETCLFGWGATKVSDFDQQMLNAYRESWRNPA
MIHGSCSDYRAAATIDLEHDSADIQRKVECPTLVFYGSKG**G**MGQLFDIPAEWAKRCNNTTNASLPGGHFFVDQFP
AETSEILLKFLARNG*

**H1-VW (S147V-Q245W)**

MGSSHHHHHHSGLVPRGSAGMDFPGFKNSTVTVDGVDIAYTVSGEGPPVLMLHGFPQNRAMWARVAPQLAEHHT
VVCADLRGYGDSDKPKCLPDRSNYSFRTFAHDQLCVMRHLGFERFHLVGHDRGGRTGHRMALDHPEAVLSLTVMD
IVPTYAMFMNTNRLVAA**V**YWHWYFLQQPEPFPEHMIGQDPDFFYETCLFGWGATKVSDFDQQMLNAYRESWRNPA
MIHGSCSDYRAAATIDLEHDSADIQRKVECPTLVFYGSKG**W**MGQLFDIPAEWAKRCNNTTNASLPGGHFFVDQFP
AETSEILLKFLARNG*

**H1-FI (Q245F-M246I)**

MGSSHHHHHHSGLVPRGSAGMDFPGFKNSTVTVDGVDIAYTVSGEGPPVLMLHGFPQNRAMWARVAPQLAEHHT
VVCADLRGYGDSDKPKCLPDRSNYSFRTFAHDQLCVMRHLGFERFHLVGHDRGGRTGHRMALDHPEAVLSLTVMD
IVPTYAMFMNTNRLVAASYWHWYFLQQPEPFPEHMIGQDPDFFYETCLFGWGATKVSDFDQQMLNAYRESWRNPA
MIHGSCSDYRAAATIDLEHDSADIQRKVECPTLVFYGSKG**FI**GQLFDIPAEWAKRCNNTTNASLPGGHFFVDQFP
AETSEILLKFLARNG*

**H1-ML (Q245M-M246L)**

MGSSHHHHHHSGLVPRGSAGMDFPGFKNSTVTVDGVDIAYTVSGEGPPVLMLHGFPQNRAMWARVAPQLAEHHT
VVCADLRGYGDSDKPKCLPDRSNYSFRTFAHDQLCVMRHLGFERFHLVGHDRGGRTGHRMALDHPEAVLSLTVMD
IVPTYAMFMNTNRLVAASYWHWYFLQQPEPFPEHMIGQDPDFFYETCLFGWGATKVSDFDQQMLNAYRESWRNPA
MIHGSCSDYRAAATIDLEHDSADIQRKVECPTLVFYGSKG**ML**GQLFDIPAEWAKRCNNTTNASLPGGHFFVDQFP
AETSEILLKFLARNG*

**H1-RPL (A46V-S147R-Q245P-M246L)**

MGSSHHHHHHSGLVPRGSAGMDFPGFKNSTVTVDGVDIAYTVSGEGPPVLMLHGFPQNRAMWARV**V**PQLAEHHT
VVCADLRGYGDSDKPKCLPDRSNYSFRTFAHDQLCVMRHLGFERFHLVGHDRGGRTGHRMALDHPEAVLSLTVMD
IVPTYAMFMNTNRLVAA**R**YWHWYFLQQPEPFPEHMIGQDPDFFYETCLFGWGATKVSDFDQQMLNAYRESWRNPA
MIHGSCSDYRAAATIDLEHDSADIQRKVECPTLVFYGSKG**PL**GQLFDIPAEWAKRCNNTTNASLPGGHFFVDQFP
AETSEILLKFLARNG*

**H1-RL (S147R-Q245L)**

MGSSHHHHHHSGLVPRGSAGMDFPGFKNSTVTVDGVDIAYTVSGEGPPVLMLHGFPQNRAMWARVAPQLAEHHT
VVCADLRGYGDSDKPKCLPDRSNYSFRTFAHDQLCVMRHLGFERFHLVGHDRGGRTGHRMALDHPEAVLSLTVMD
IVPTYAMFMNTNRLVAA**R**YWHWYFLQQPEPFPEHMIGQDPDFFYETCLFGWGATKVSDFDQQMLNAYRESWRNPA
MIHGSCSDYRAAATIDLEHDSADIQRKVECPTLVFYGSKG**L**MGQLFDIPAEWAKRCNNTTNASLPGGHFFVDQFP
AETSEILLKFLARNG*

**H1-RHL (S147R-Q245H-M246L)**

MGSSHHHHHHSGLVPRGSAGMDFPGFKNSTVTVDGVDIAYTVSGEGPPVLMLHGFPQNRAMWARVAPQLAEHHT
VVCADLRGYGDSDKPKCLPDRSNYSFRTFAHDQLCVMRHLGFERFHLVGHDRGGRTGHRMALDHPEAVLSLTVMD
IVPTYAMFMNTNRLVAA**R**YWHWYFLQQPEPFPEHMIGQDPDFFYETCLFGWGATKVSDFDQQMLNAYRESWRNPA

MIHGSCSDYRAAATIDLEHDSADIQRKVECPTLVFYGSKG**HL**GQLFDIPAEWAKRCNNTTNASLPGGHFFVDQFP
AETSEILLKFLARNG*

**H1-PL (S147P-Q245L)**

<u>MGSSHHHHHHGSGLVPRGSAG</u>MDFPGFKNSTVTVDGVDIAYTVSGEGPPVLMLHGFPQNRAMWARVAPQLAEHHT
VVCADLRGYGDSDKPKCLPDRSNYSFRTFAHDQLCVMRHLGFERFHLVGHDRGGRTGHRMALDHPEAVLSLTVMD
IVPTYAMFMNTNRLVAA**P**YWHWYFLQQPEPFPEHMIGQDPDFFYETCLFGWGATKVSDFDQQMLNAYRESWRNPA
MIHGSCSDYRAAATIDLEHDSADIQRKVECPTLVFYGSKG**L**MGQLFDIPAEWAKRCNNTTNASLPGGHFFVDQFP
AETSEILLKFLARNG*
