## Supplementary material for "Engineering fluoroacetate dehalogenase by growth-based selections to degrade non-natural organofluorides": 19F-NMR spectra

P39_E1_WT1_100mMFA_20240924120145
STANDARD FLUORINE PARAMETERS

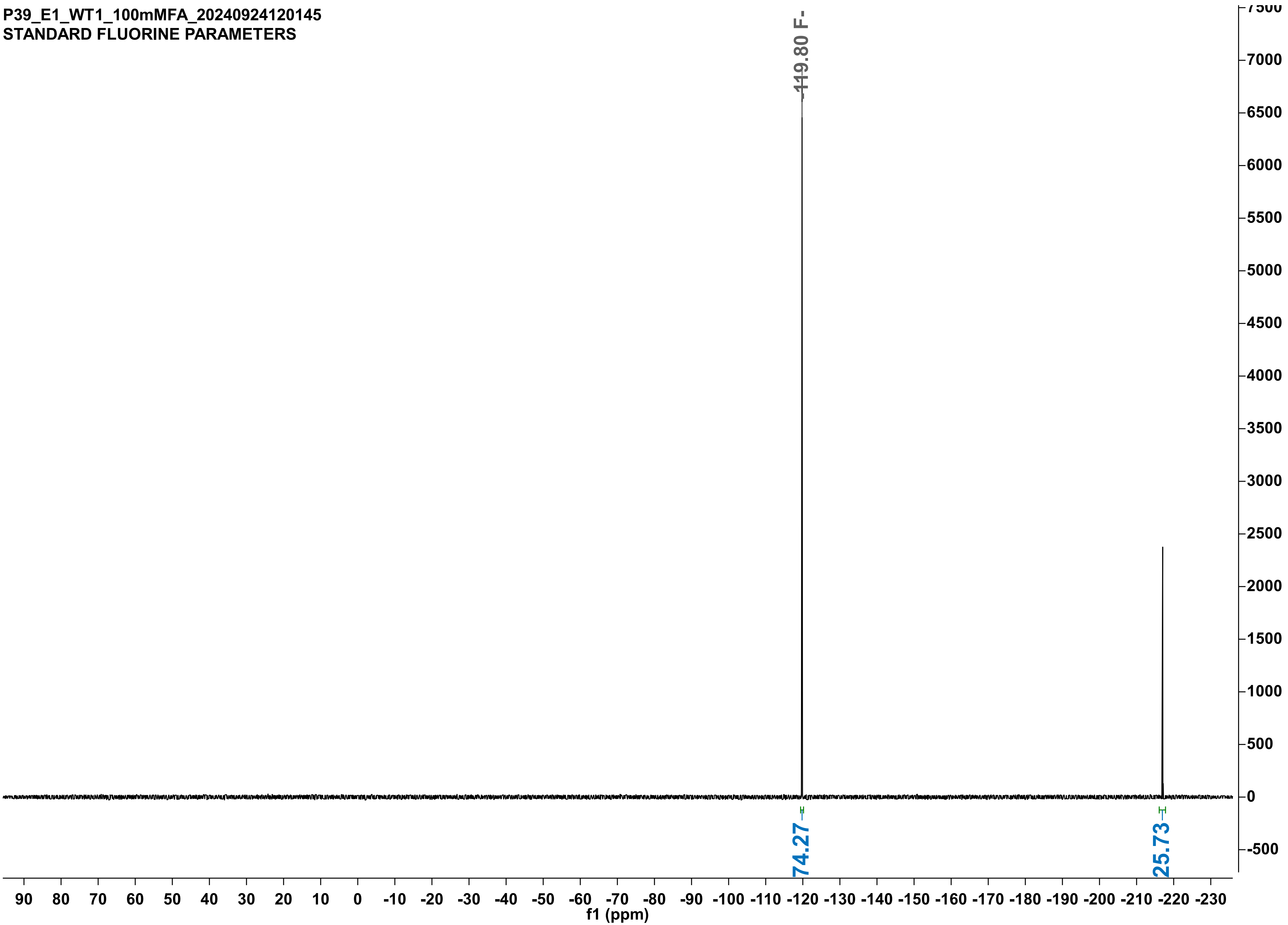

P39_E4_H272A_100mMFA_20240924120239
STANDARD FLUORINE PARAMETERS

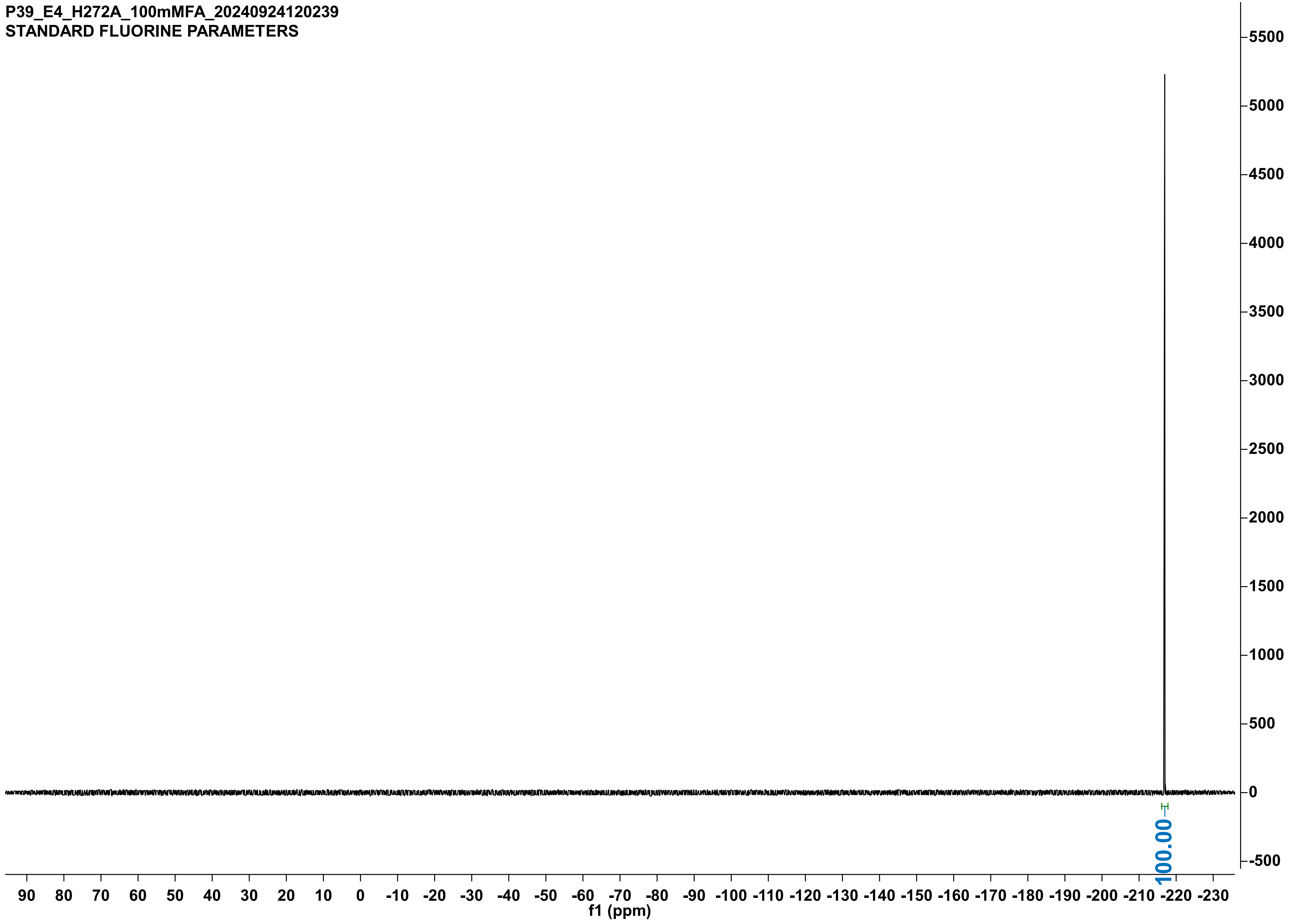

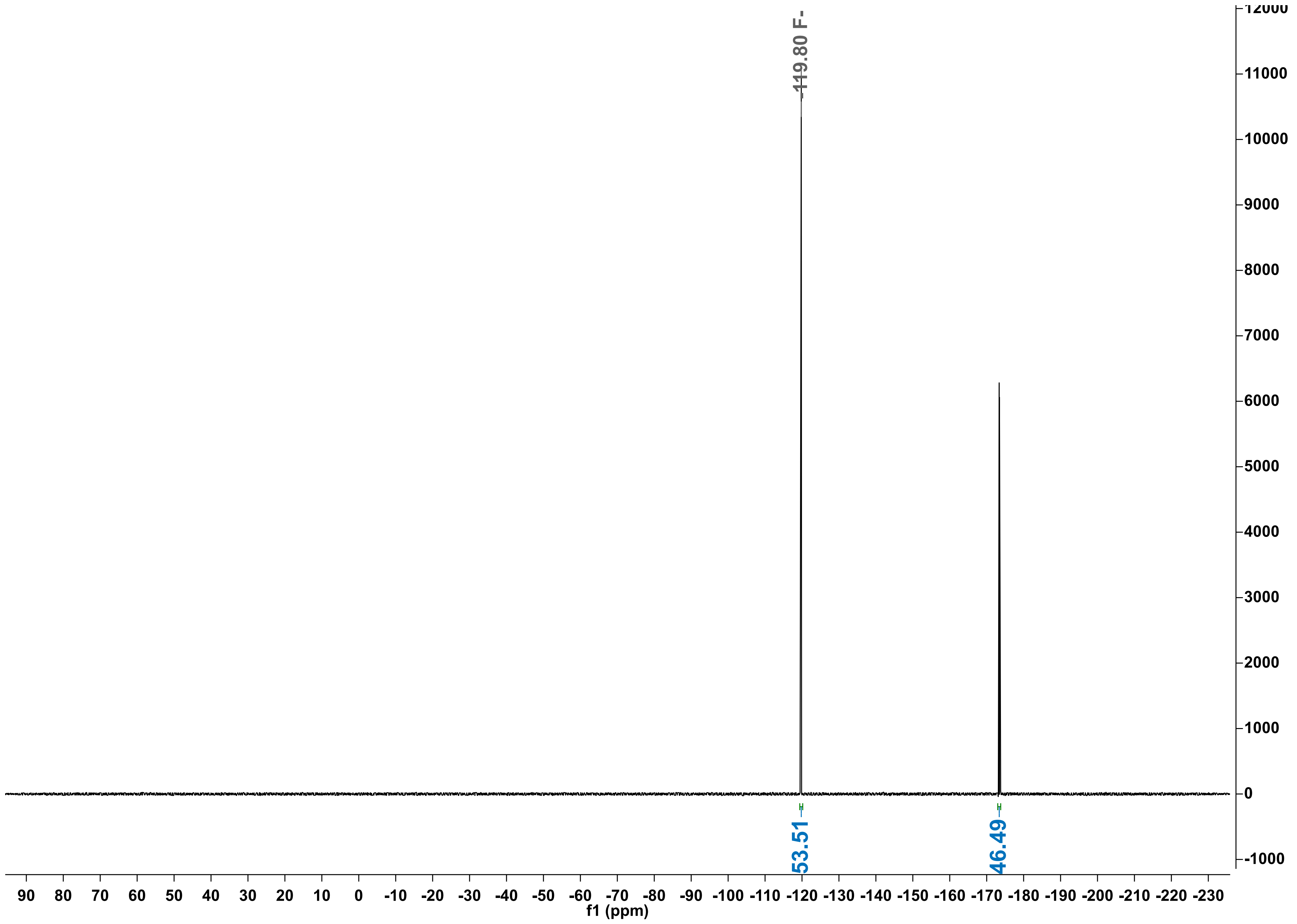

-119.80 F-
53.51
46.49
f1 (ppm)

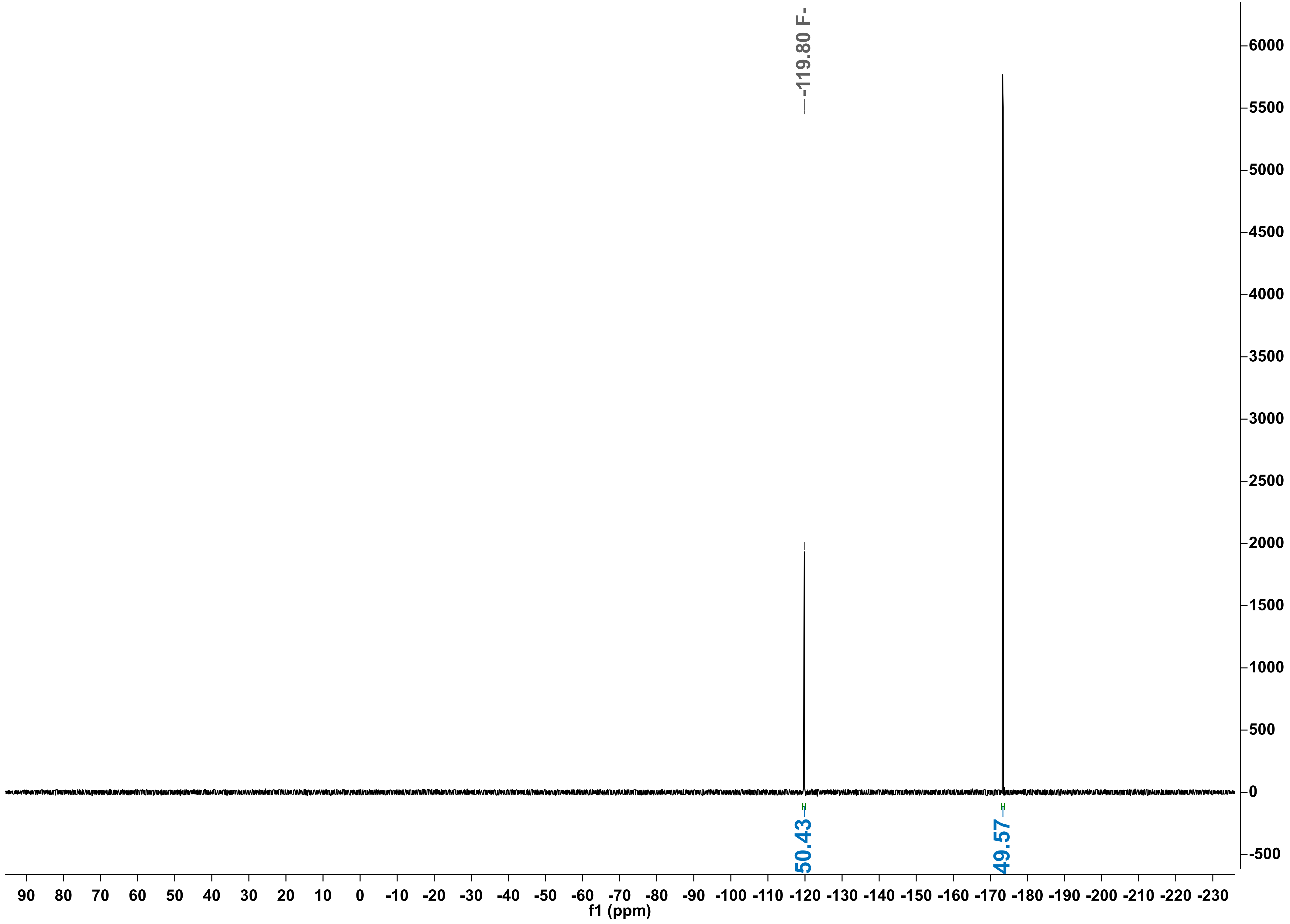

-119.80 F-
50.43
49.57
6000
5500
5000
4500
4000
3500
3000
2500
2000
1500
1000
500
0
-500
90 80 70 60 50 40 30 20 10 0 -10 -20 -30 -40 -50 -60 -70 -80 -90 -100 -110 -120 -130 -140 -150 -160 -170 -180 -190 -200 -210 -220 -230
f1 (ppm)

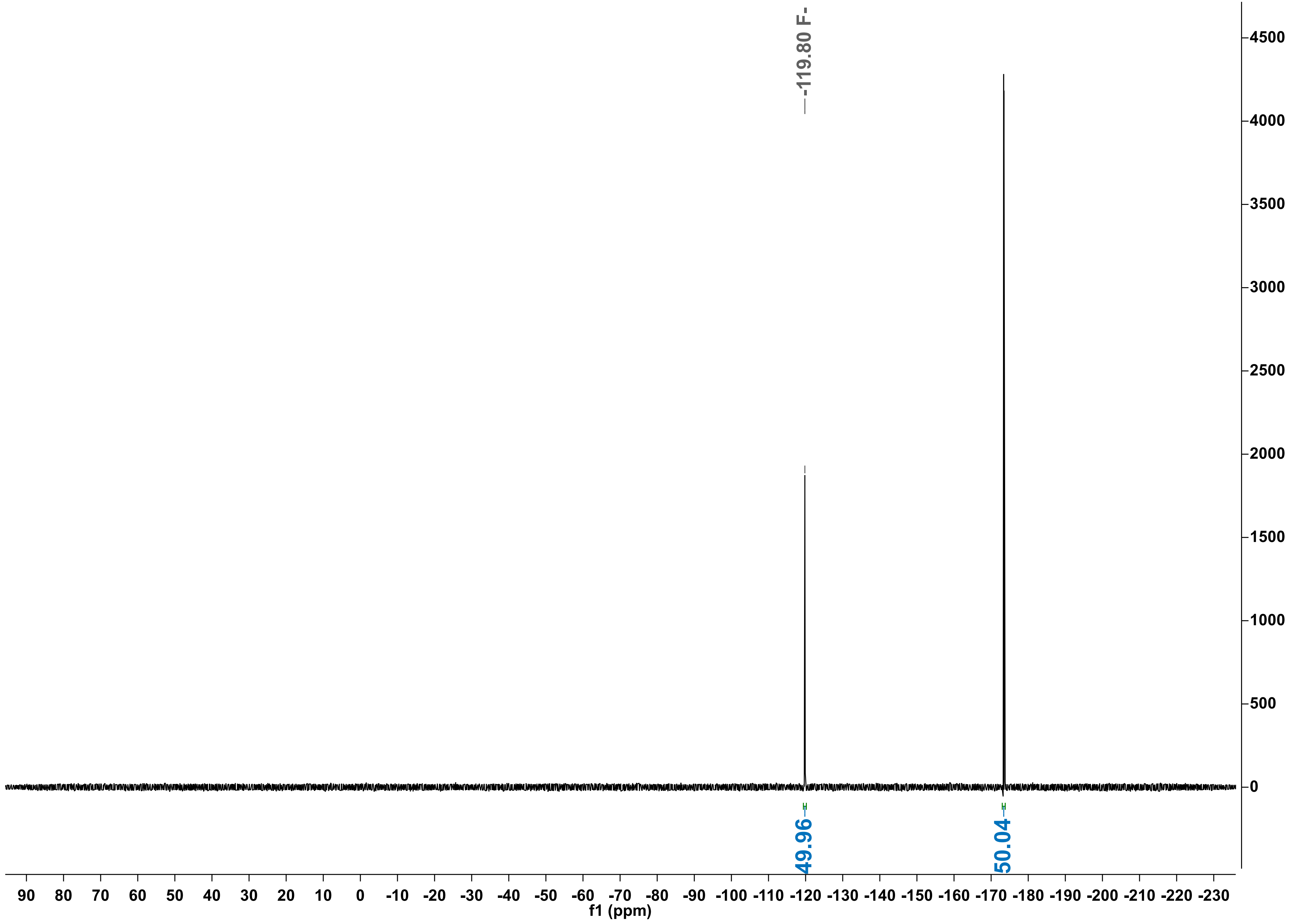

-119.80 F-
4500
4000
3500
3000
2500
2000
1500
1000
500
0
49.96
50.04
90 80 70 60 50 40 30 20 10 0 -10 -20 -30 -40 -50 -60 -70 -80 -90 -100 -110 -120 -130 -140 -150 -160 -170 -180 -190 -200 -210 -220 -230
f1 (ppm)

P50_sample19_null1_FPA_72h_20250511164508
STANDARD FLUORINE PARAMETERS

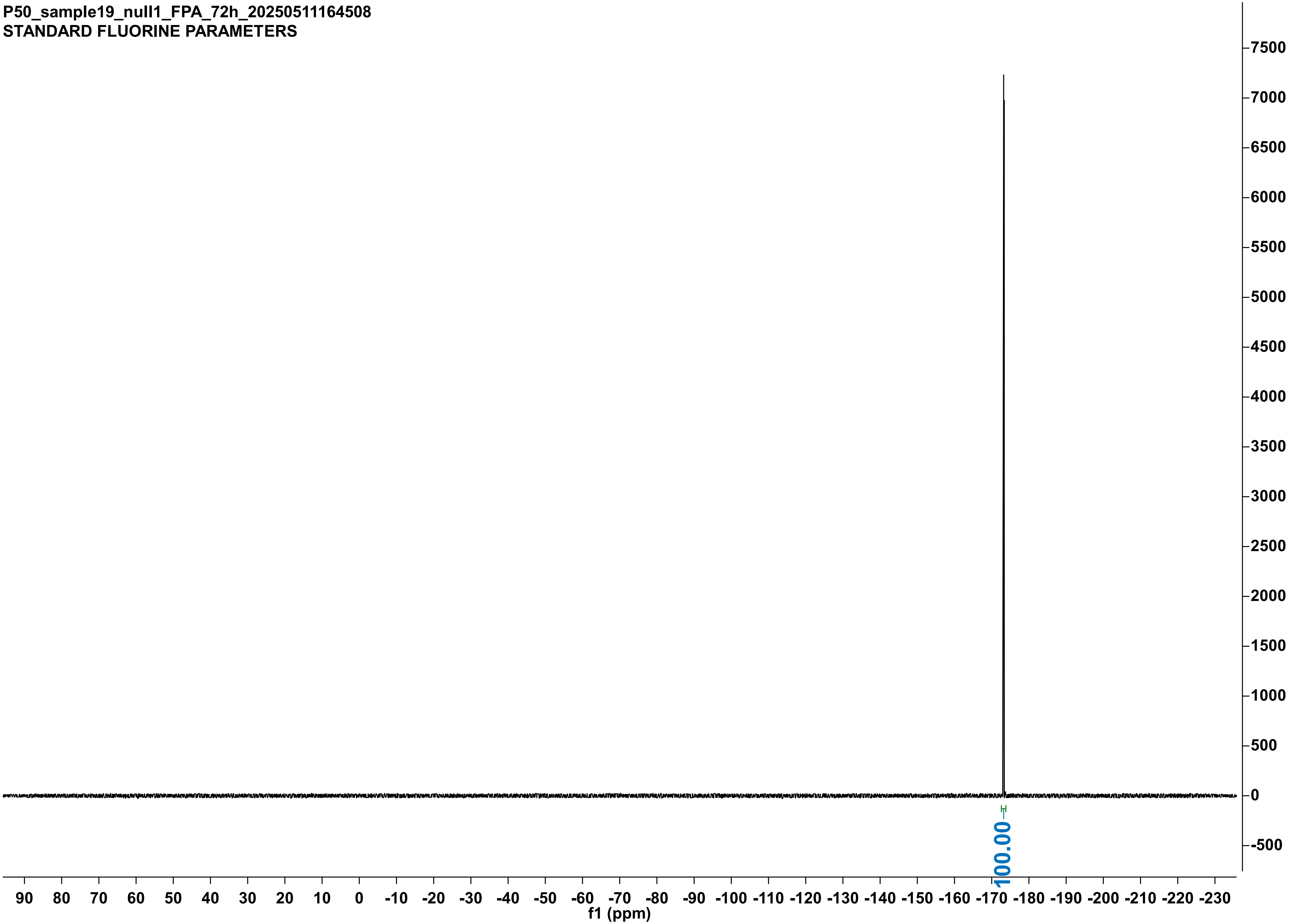

P50_sample20_null2_FPA_72h_20250511164527
STANDARD FLUORINE PARAMETERS

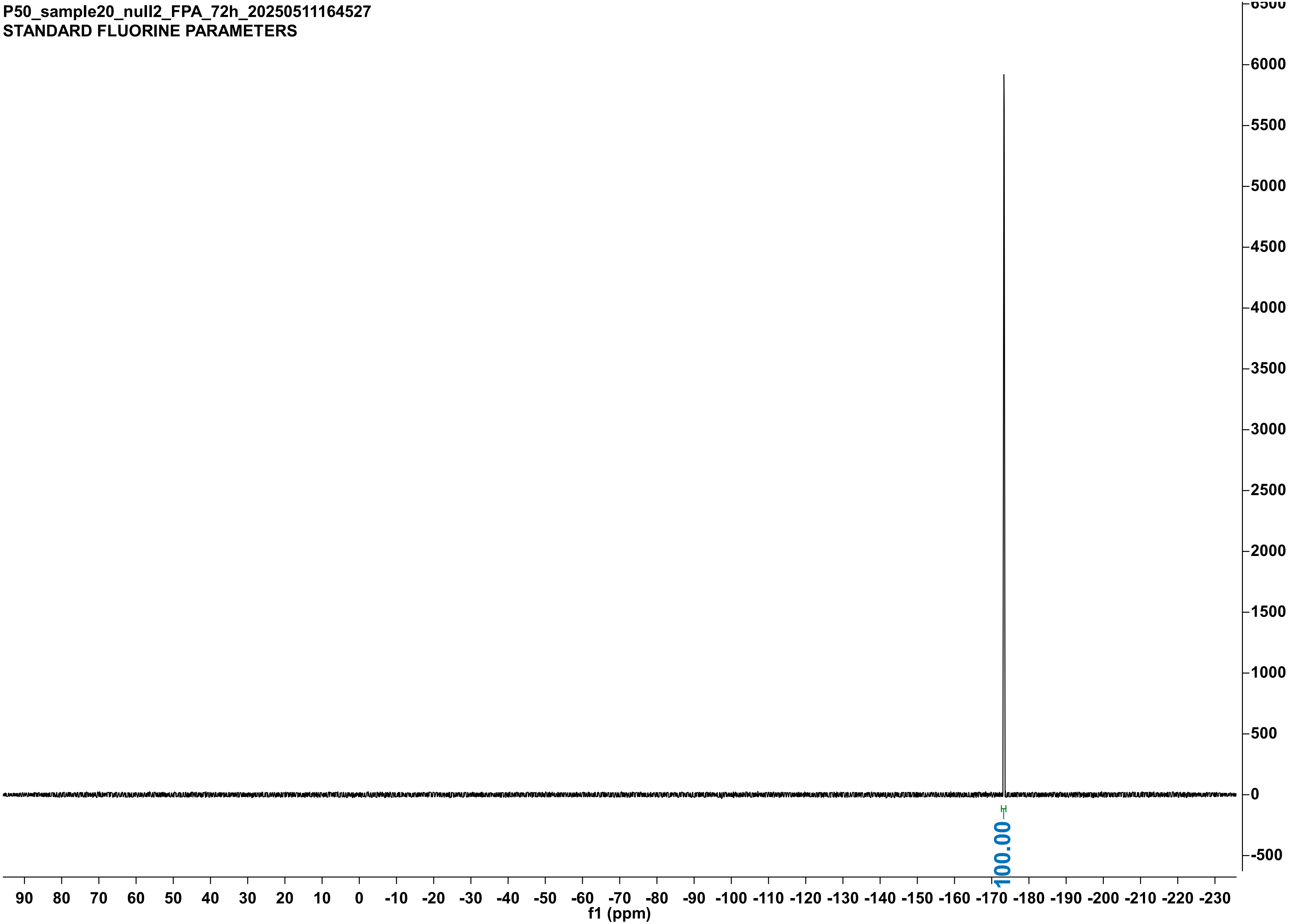

P50_sample21_null3_FPA_72h_20250511164540
STANDARD FLUORINE PARAMETERS

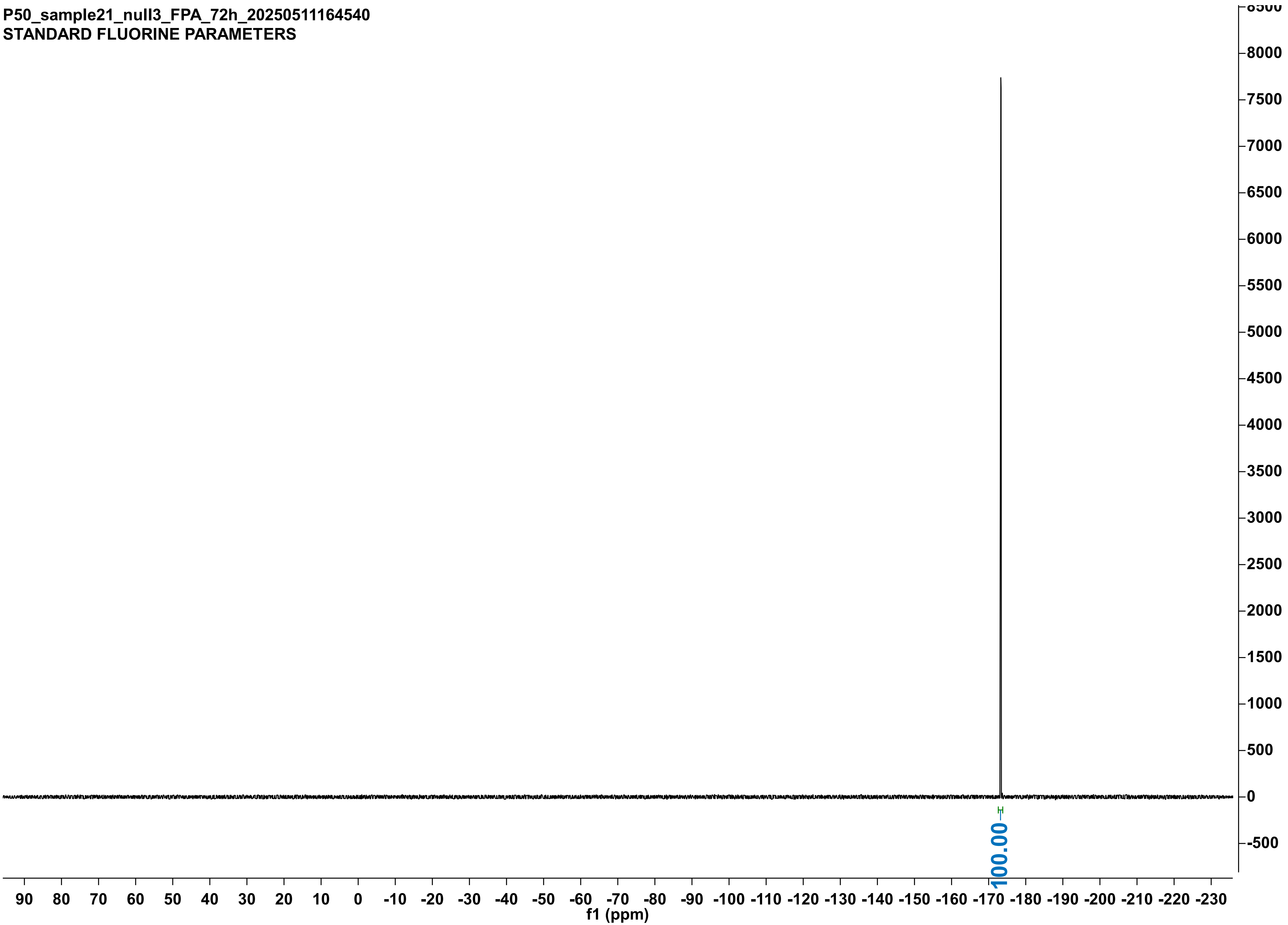

P50_sample10_WT1_2FAc_72h_20250511160442
STANDARD FLUORINE PARAMETERS

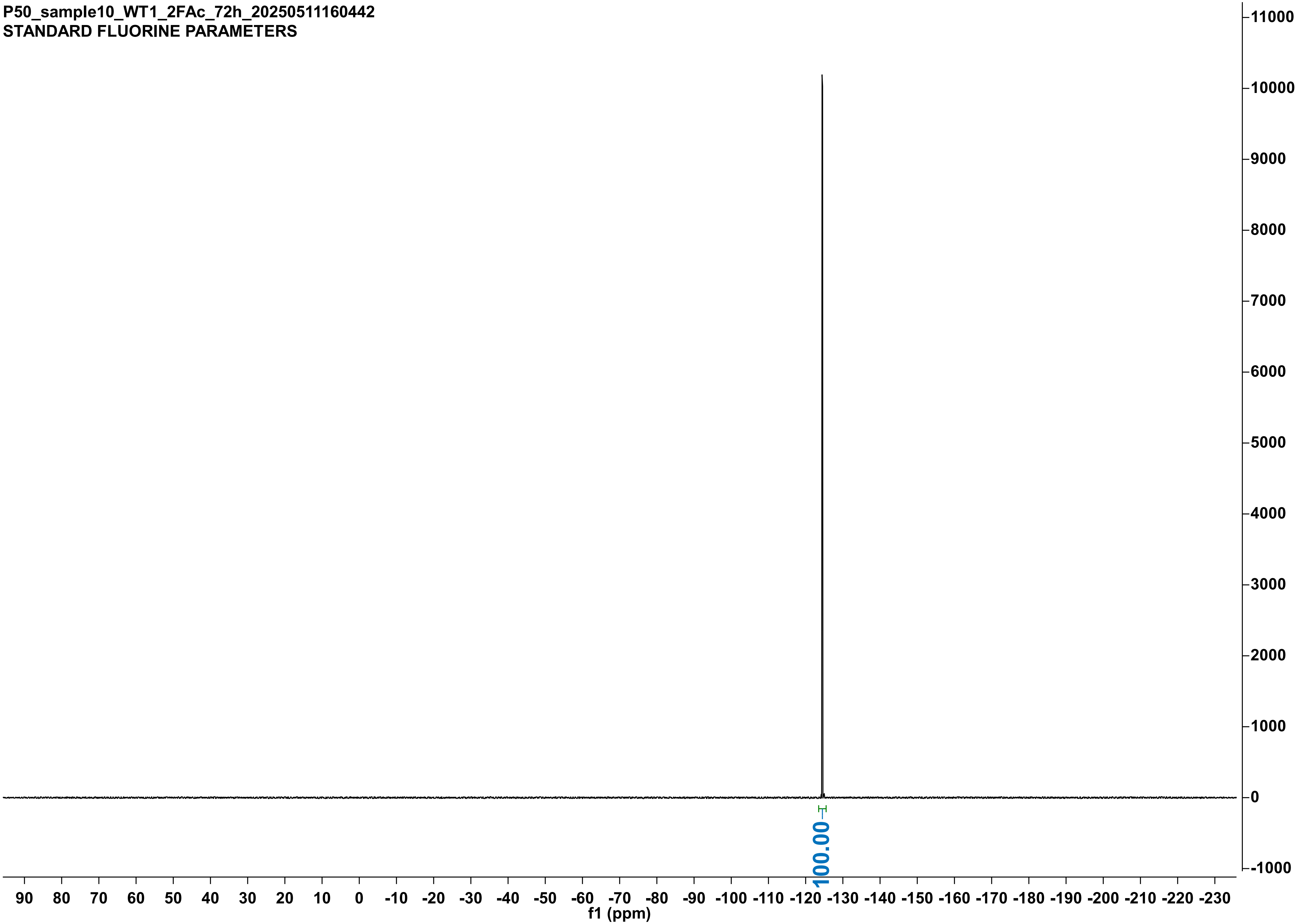

P50_sample11_WT2_2FAc_72h_20250511160453
STANDARD FLUORINE PARAMETERS

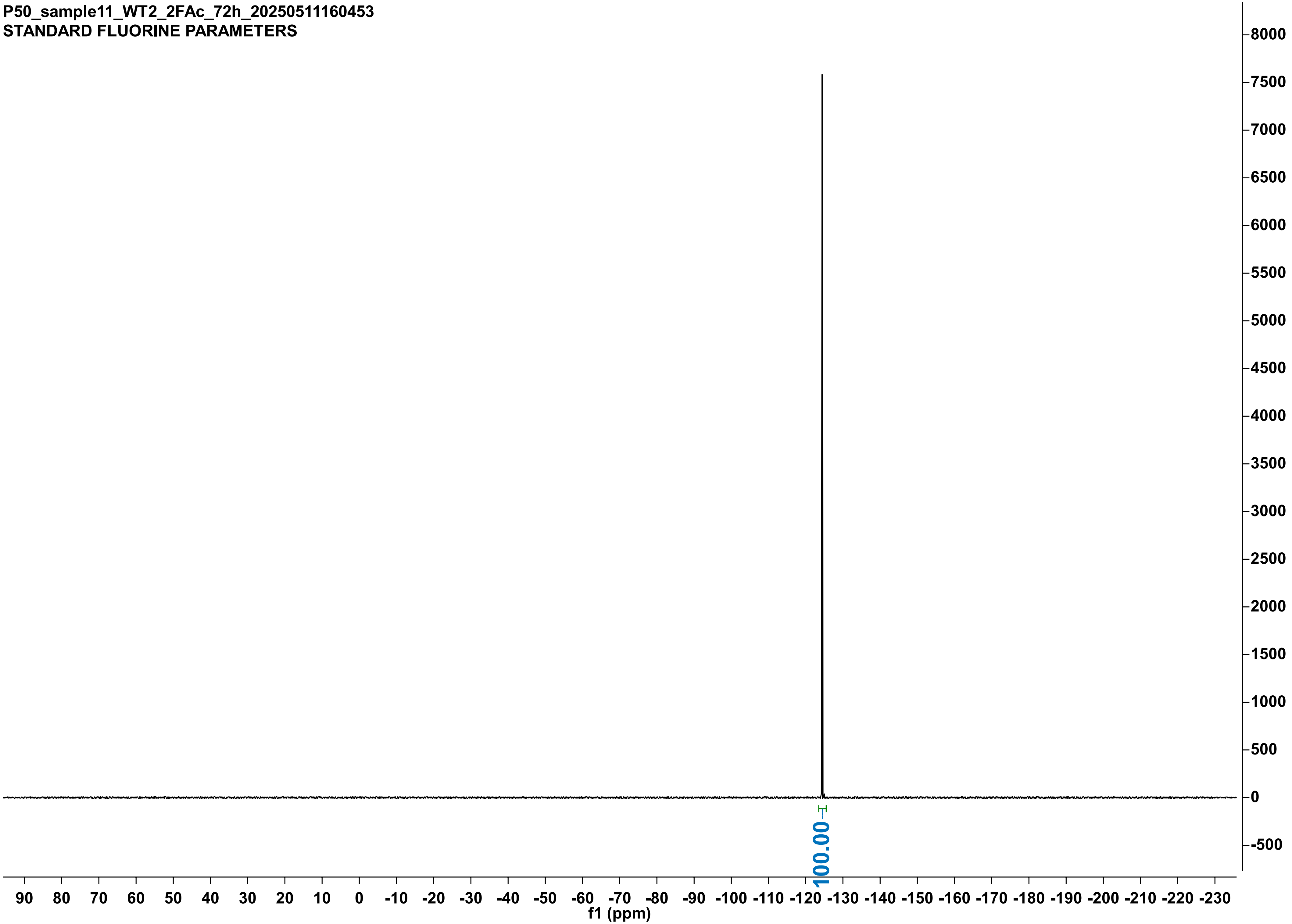

P50_sample12_WT3_2FAc_72h_20250511160503
STANDARD FLUORINE PARAMETERS

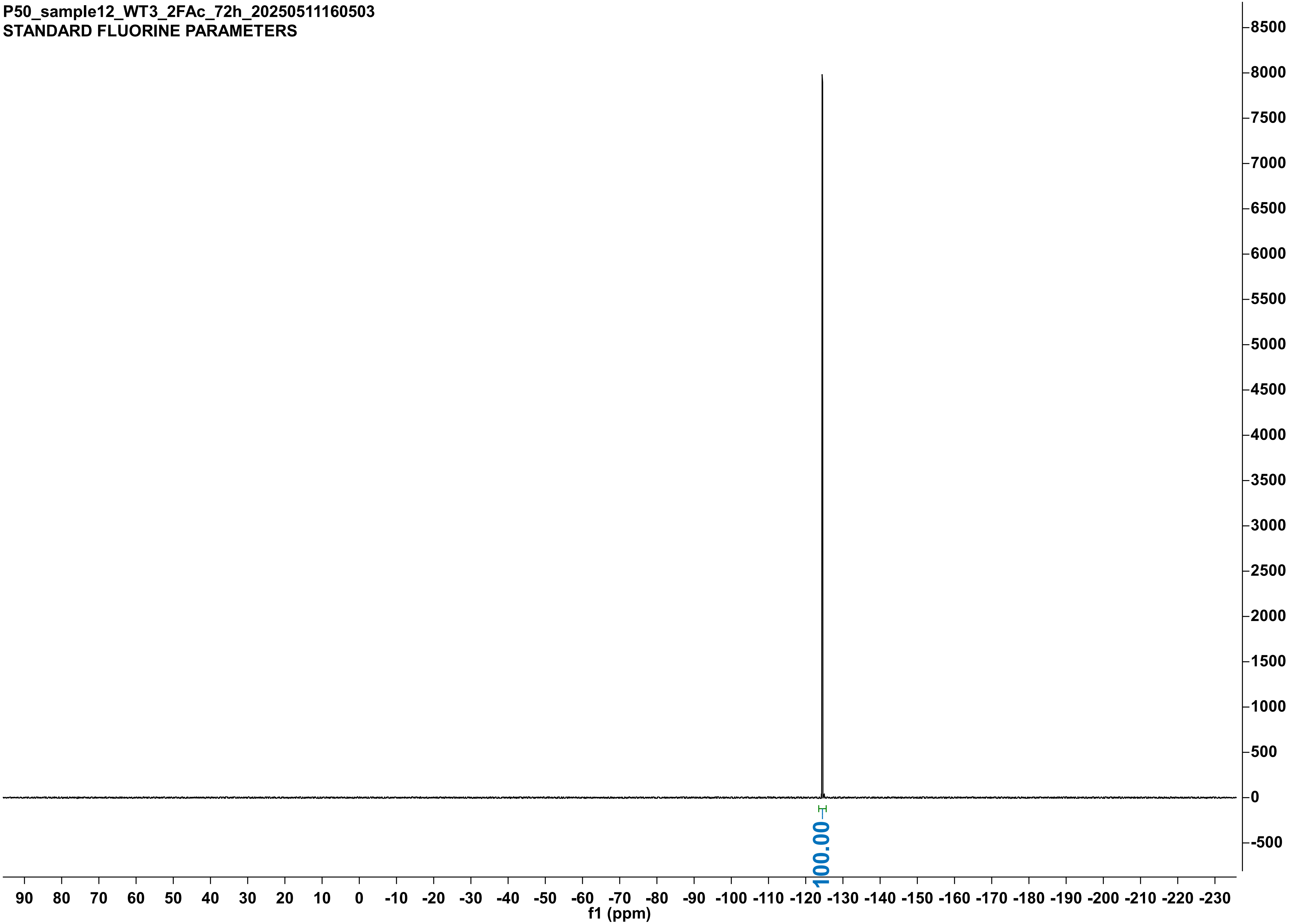

P50_sample7_null1_2FAc_72h_20250511160407
STANDARD FLUORINE PARAMETERS

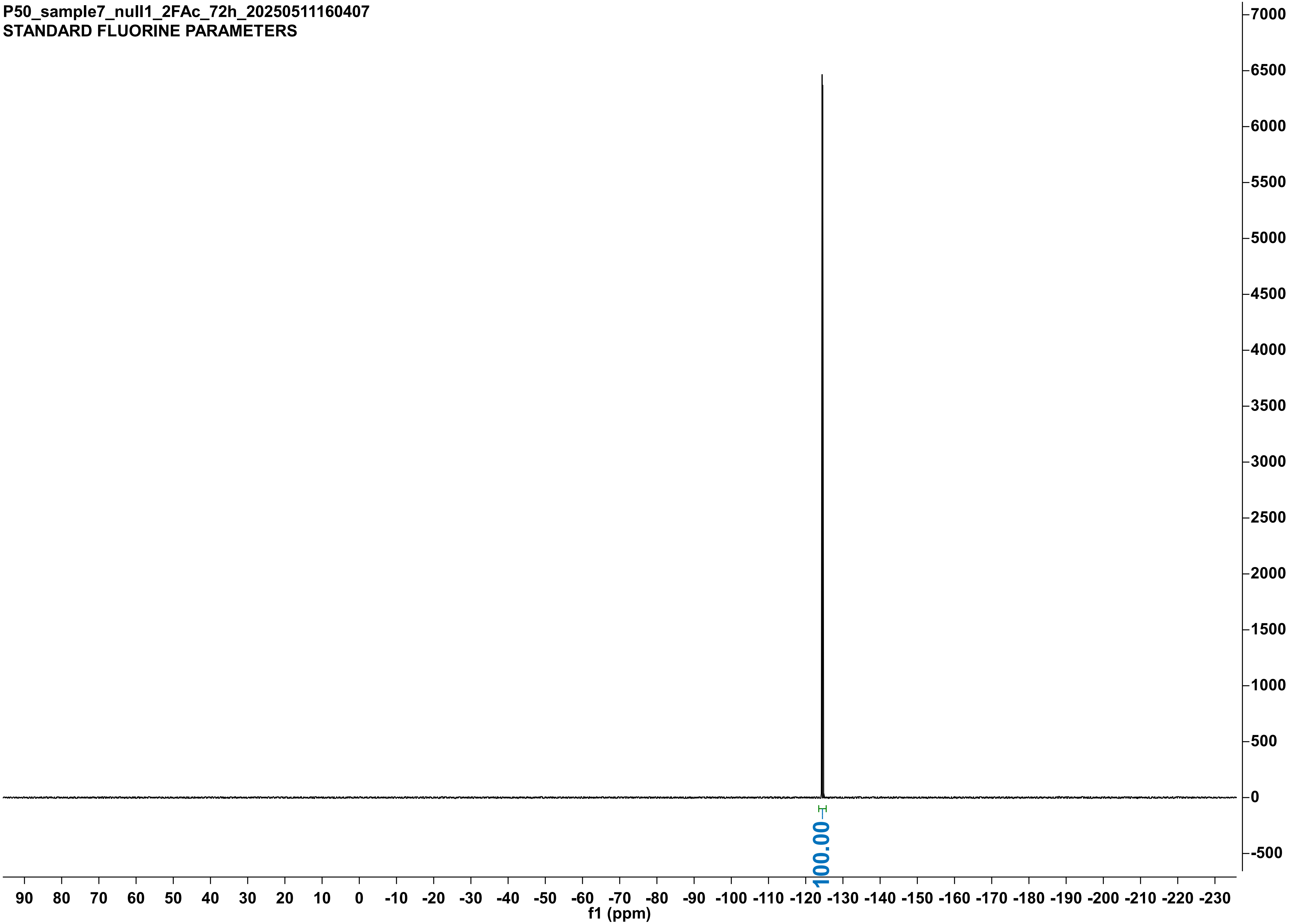

P50_sample8_null2_2FAc_72h_20250511160422
STANDARD FLUORINE PARAMETERS

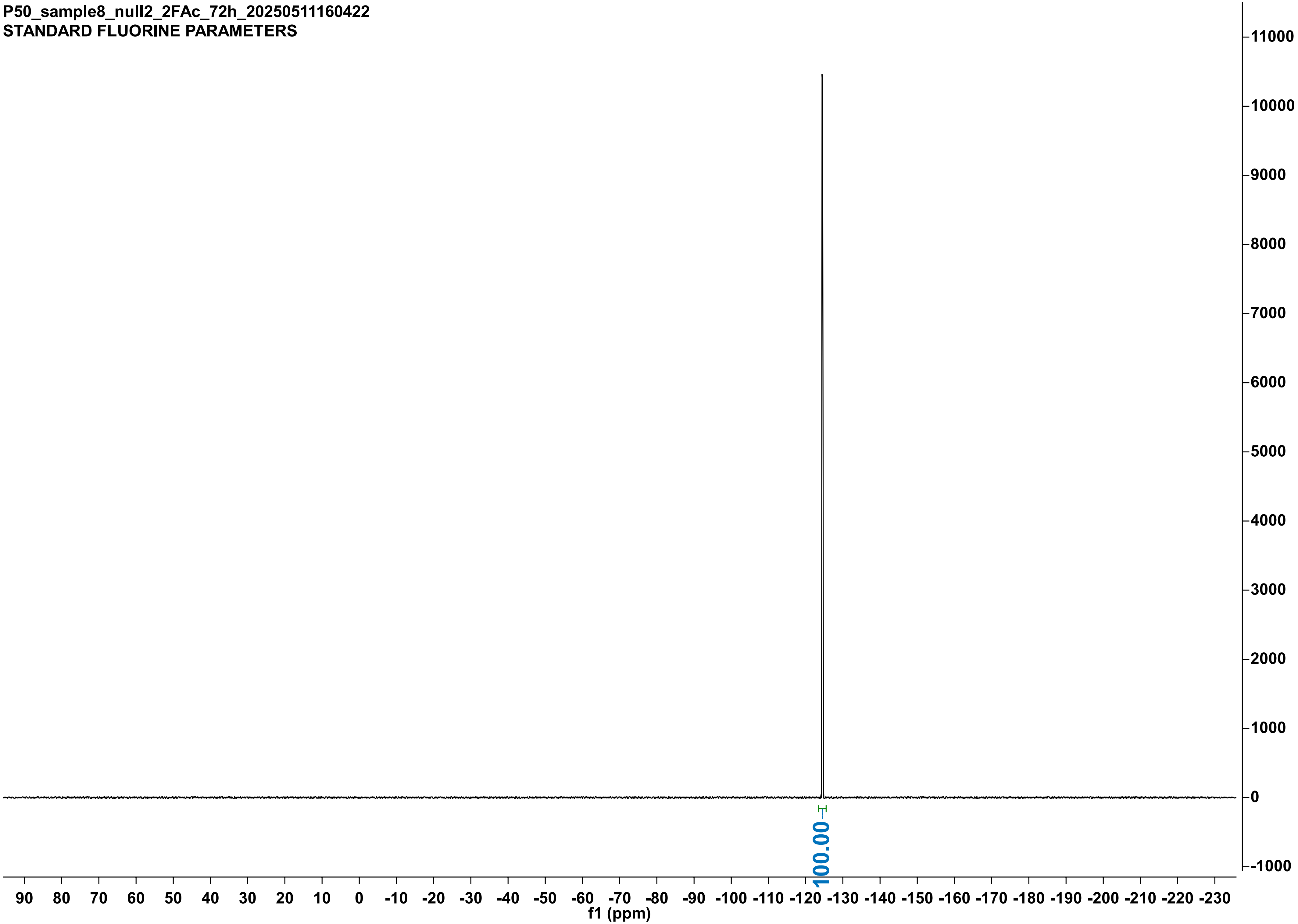

P50_sample9_null3_2FAc_72h_20250511160434
STANDARD FLUORINE PARAMETERS

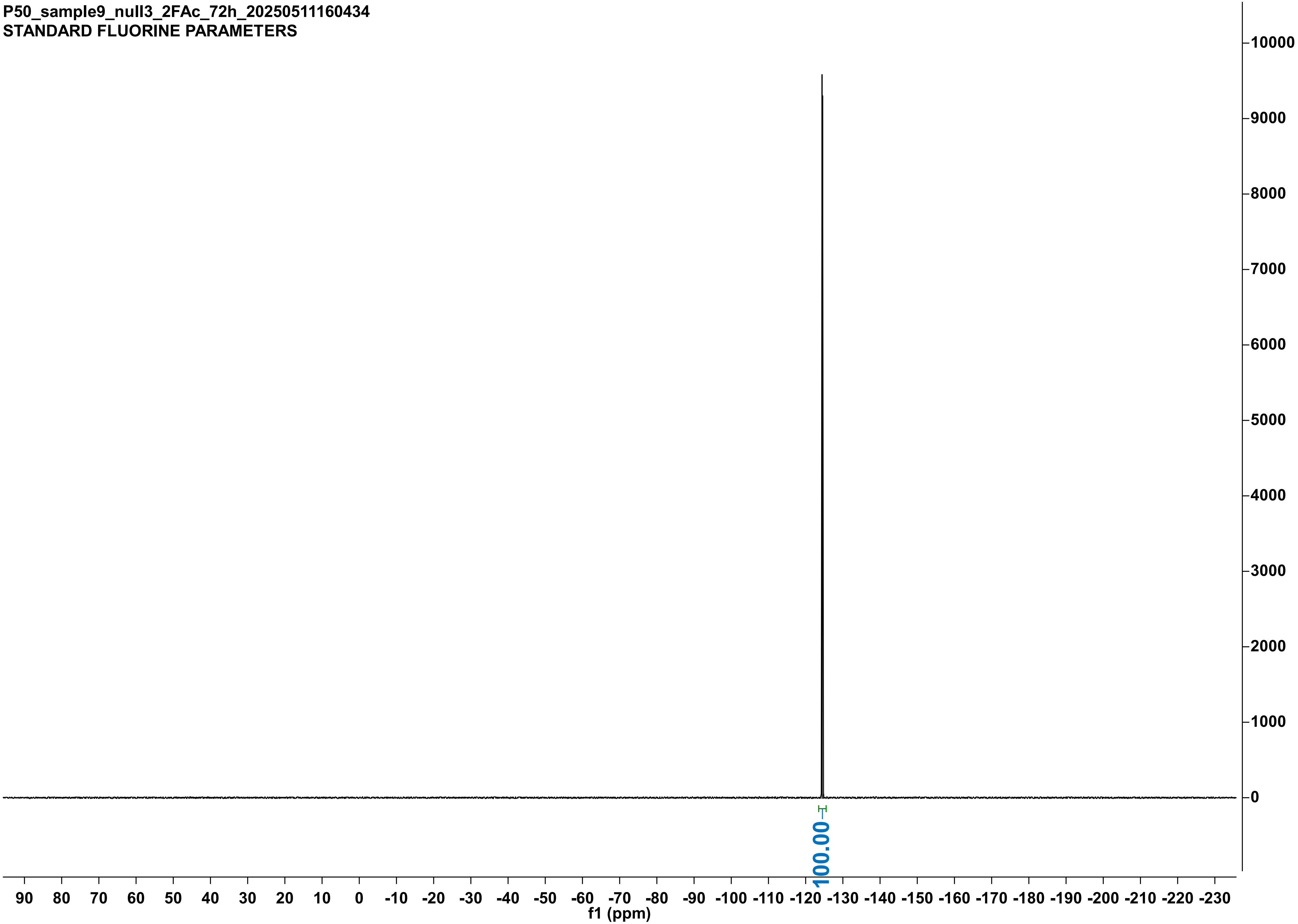

P50_sample16_WT1_2FPA_72h_20250511160616
STANDARD FLUORINE PARAMETERS

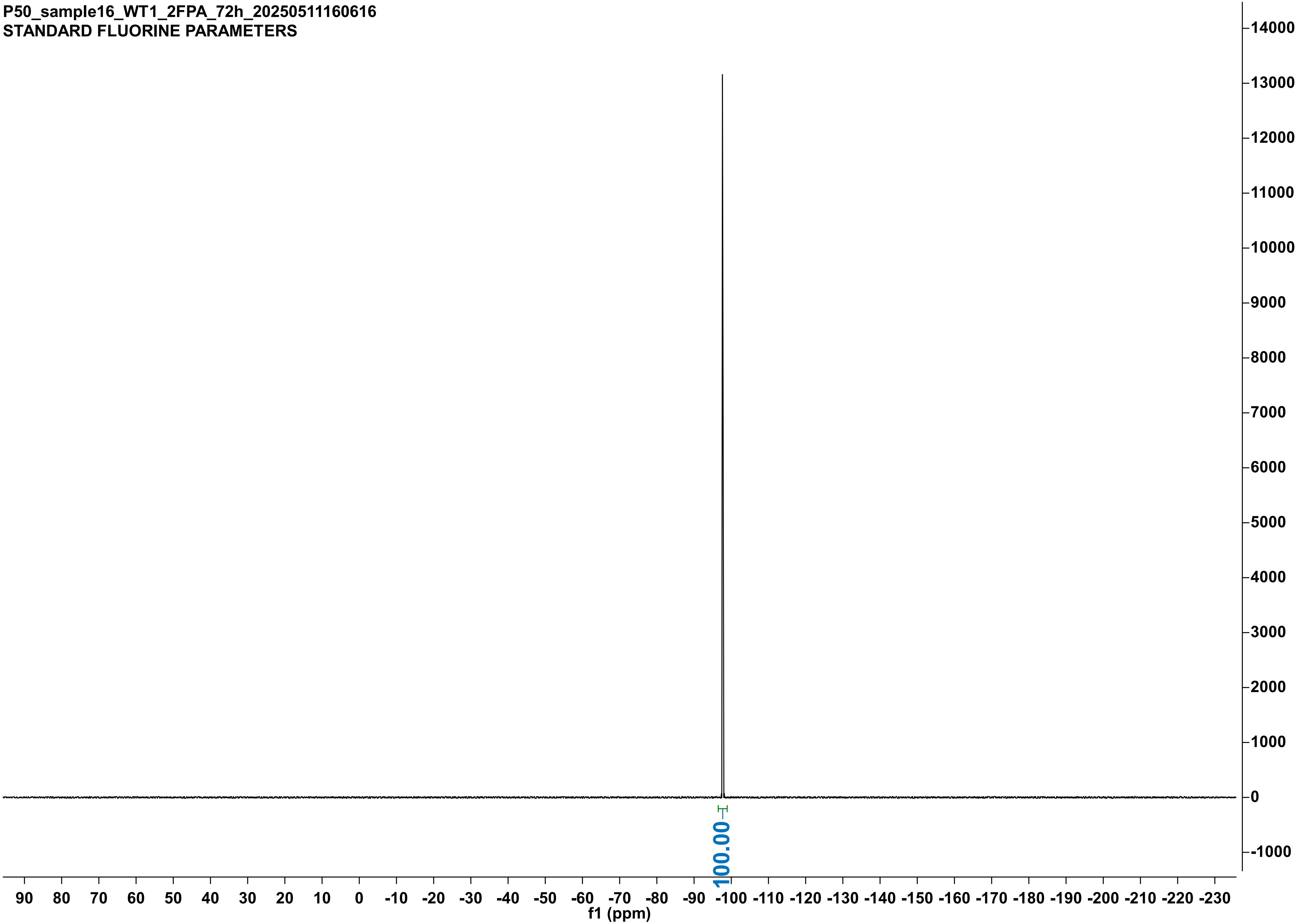

P50_sample17_WT2_2FPA_72h_20250511164442
STANDARD FLUORINE PARAMETERS

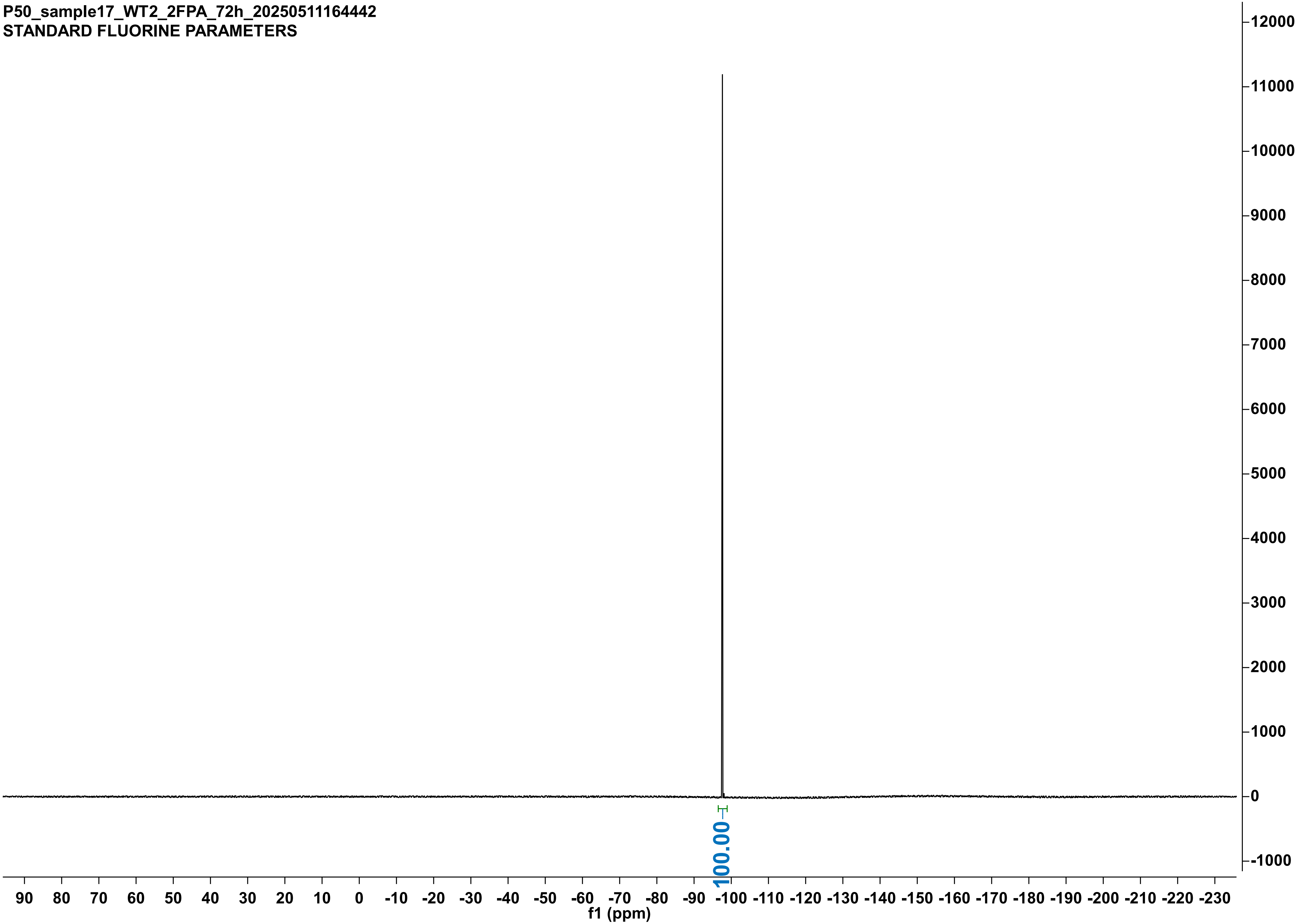

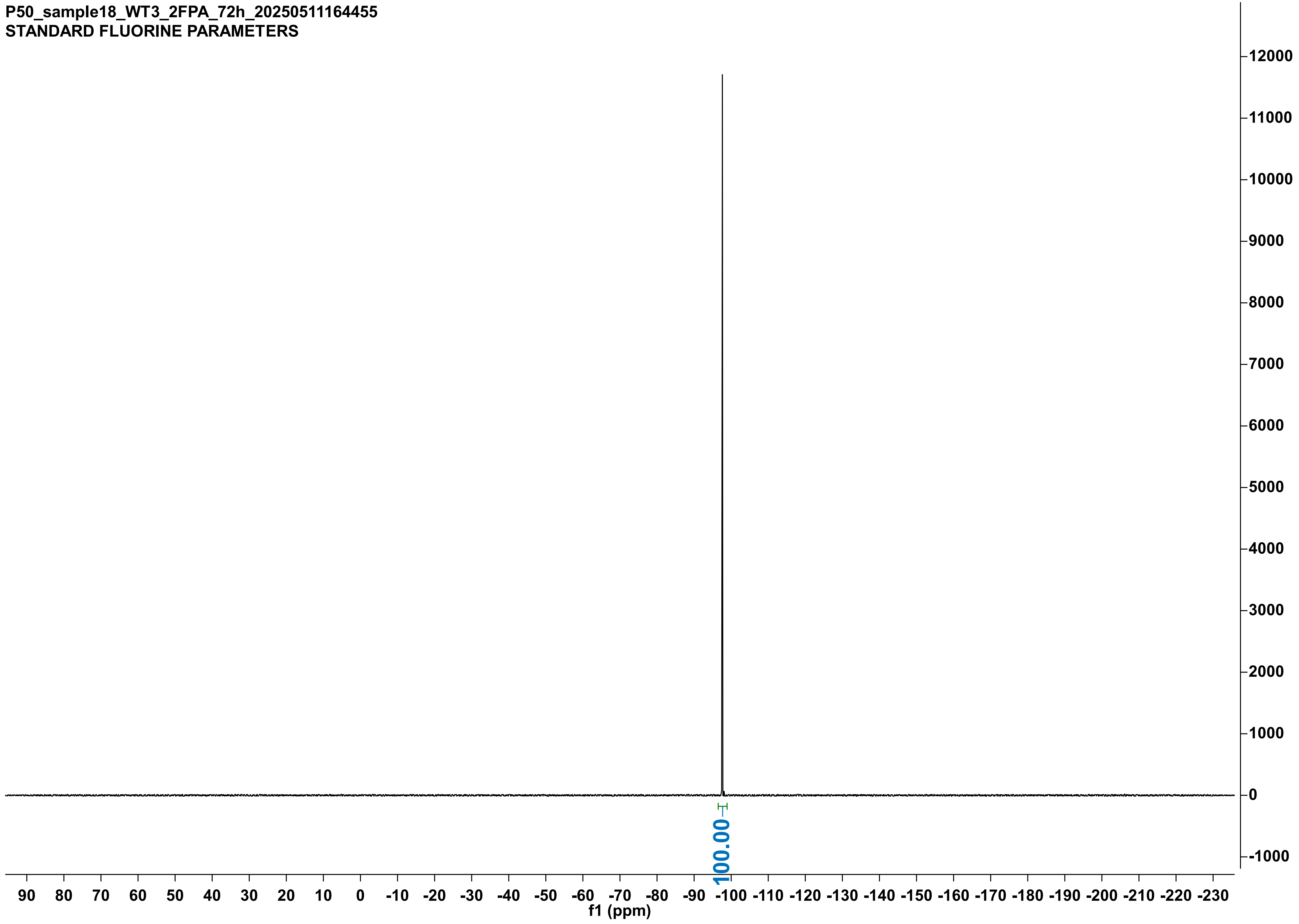

P50_sample18_WT3_2FPA_72h_20250511164455
STANDARD FLUORINE PARAMETERS
100.00
f1 (ppm)

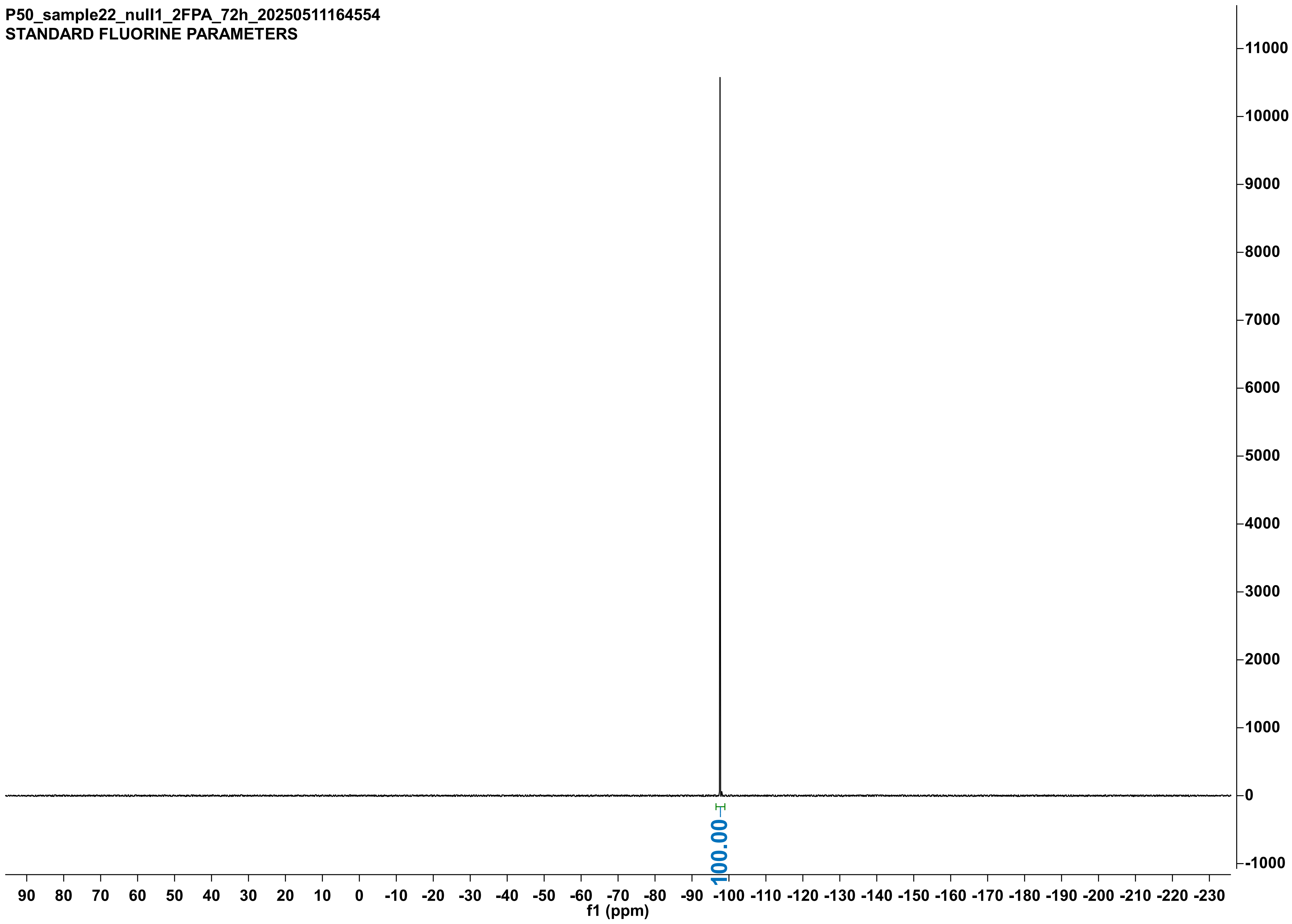
P50_sample22_null1_2FPA_72h_20250511164554
STANDARD FLUORINE PARAMETERS
100.00
f1 (ppm)

P50_sample23_null2_2FPA_72h_20250511164613
STANDARD FLUORINE PARAMETERS

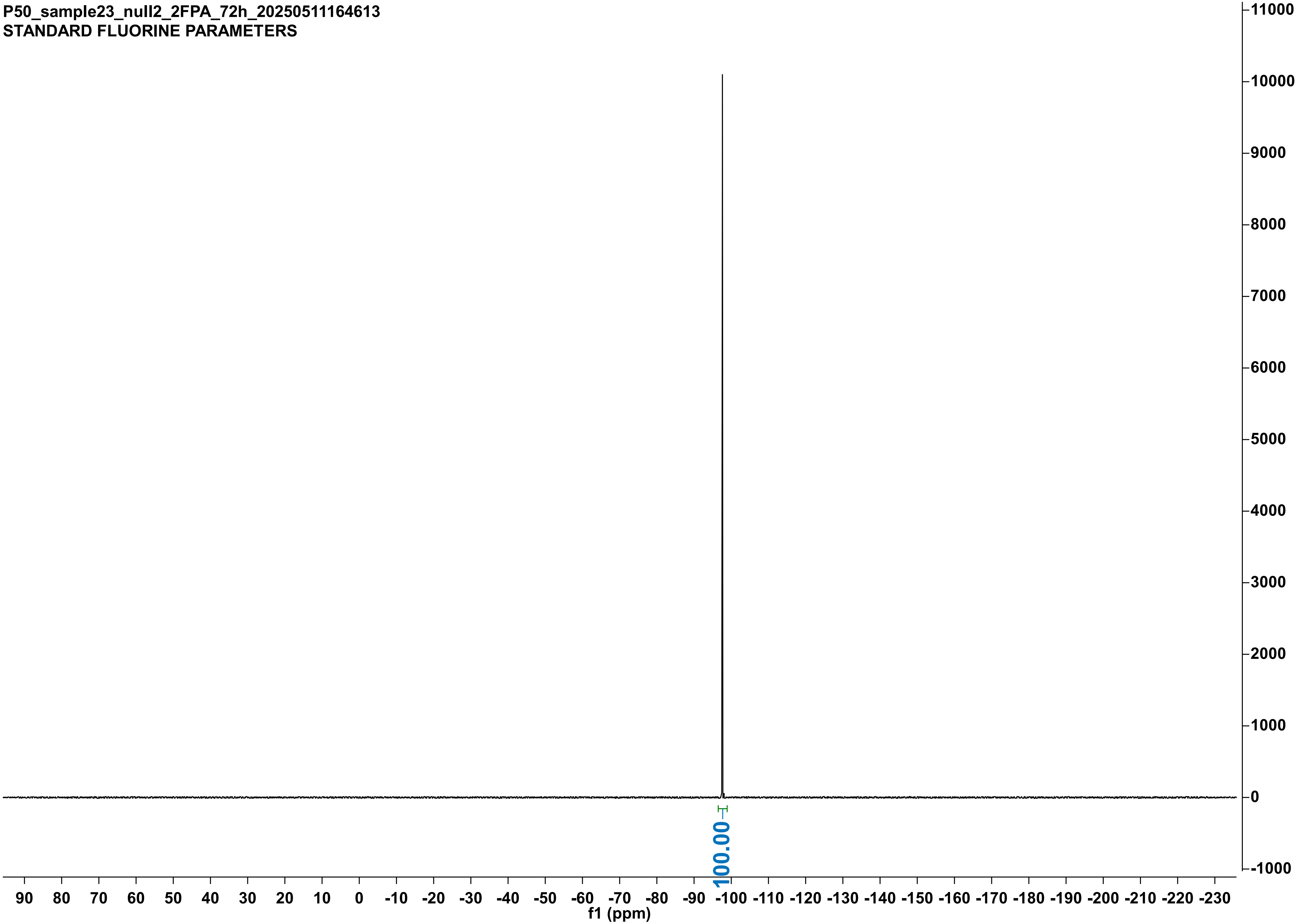

P50_sample24_null3_2FPA_72h_20250511164624
STANDARD FLUORINE PARAMETERS

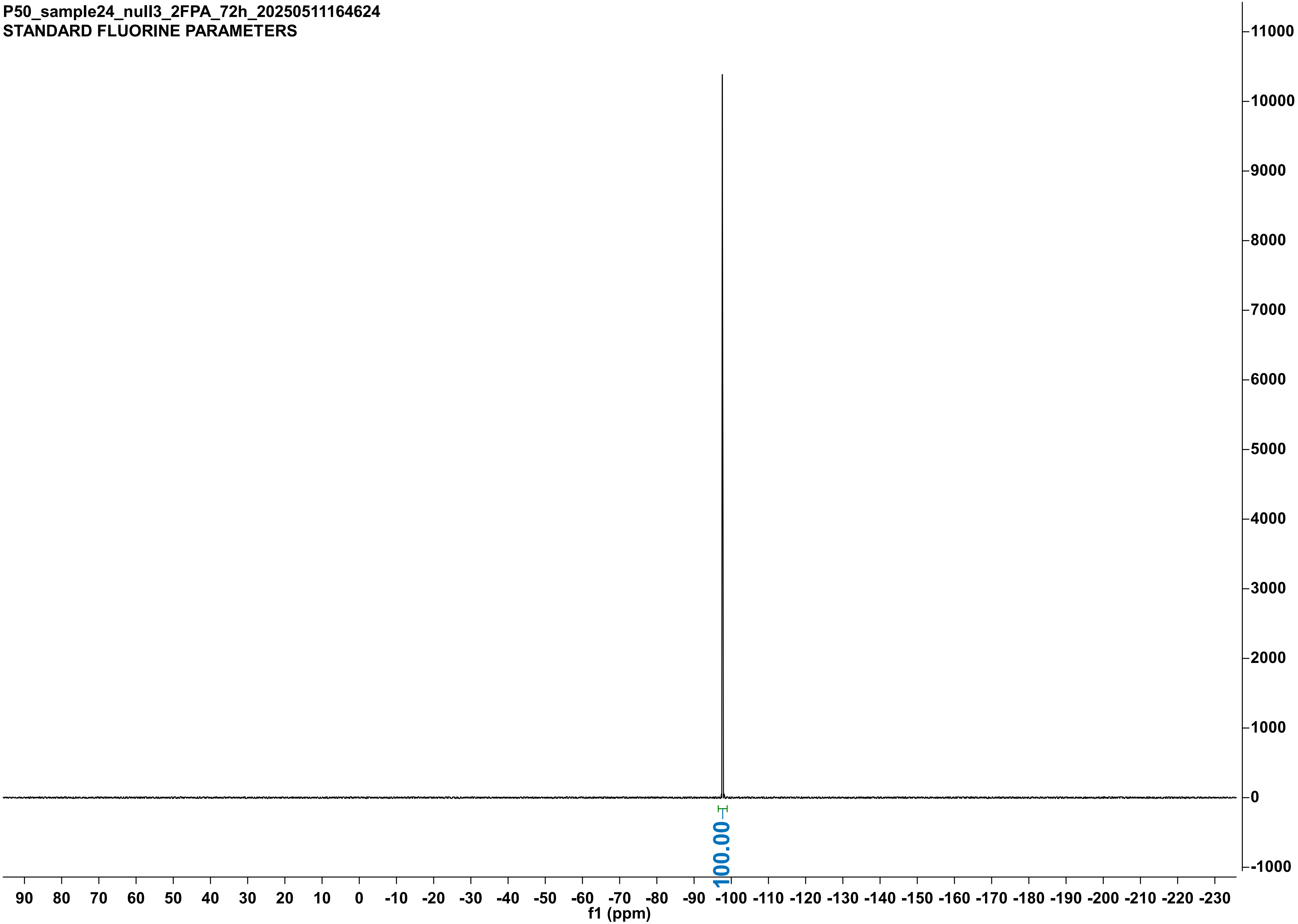

X48-11-4-05uMH272A-2-10mMFAc-20min_20250411112926
STANDARD FLUORINE PARAMETERS

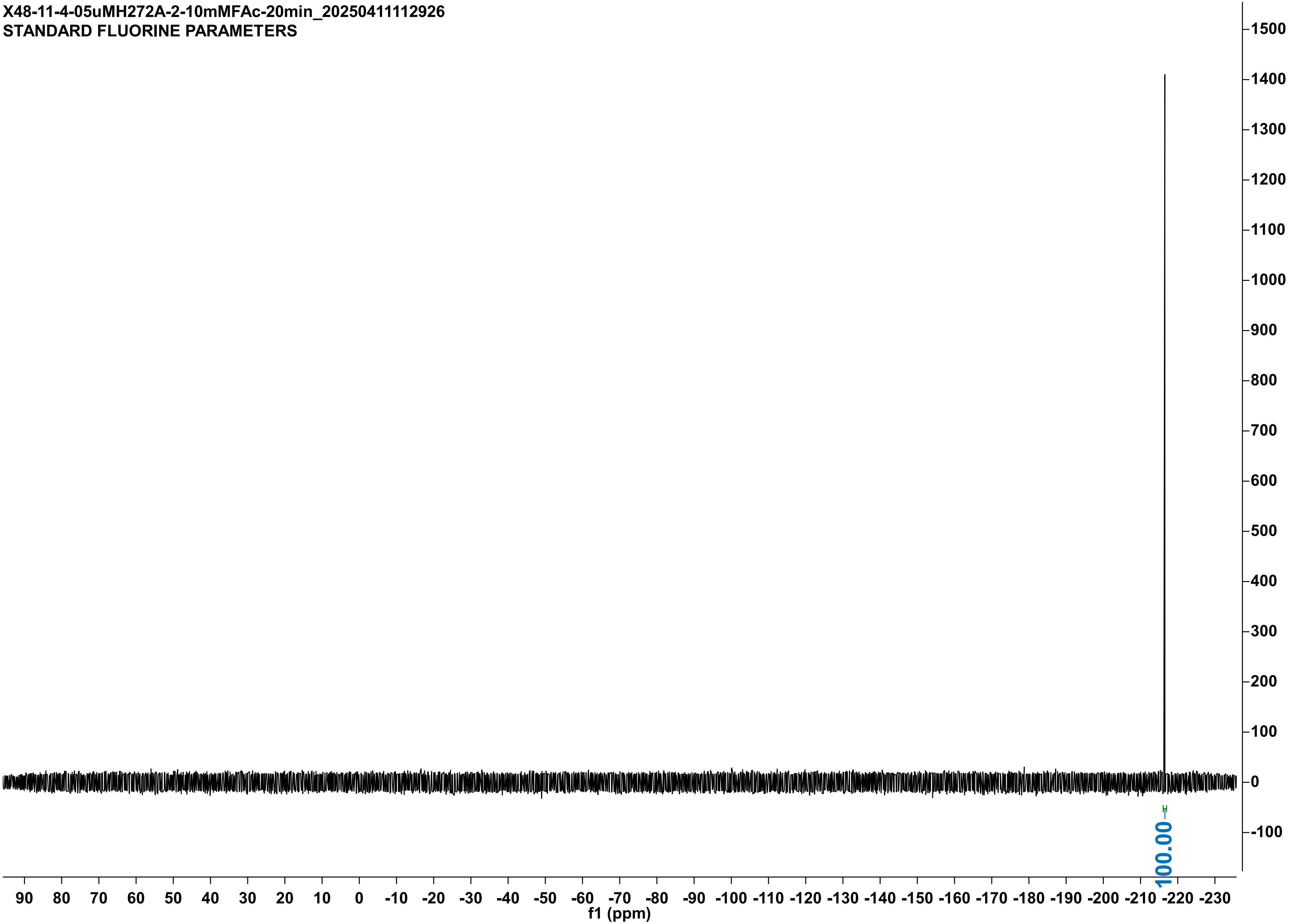

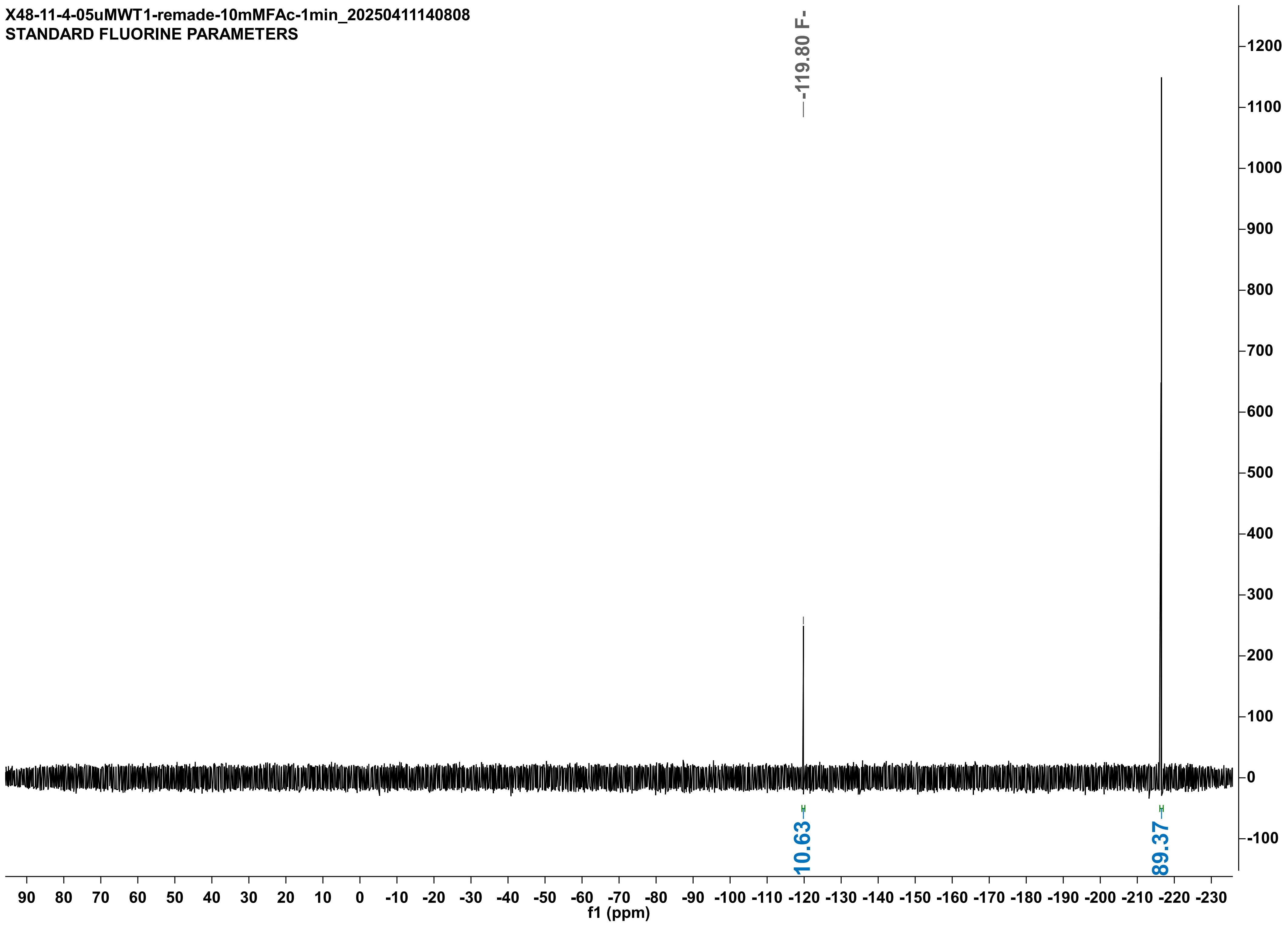

X48-11-4-05uMWT1-remade-10mMFAc-1min_20250411140808
STANDARD FLUORINE PARAMETERS
-119.80 F-
10.63
89.37
f1 (ppm)

X46-05uMWT2-10mMFAc-1min_20250318175327
STANDARD FLUORINE PARAMETERS

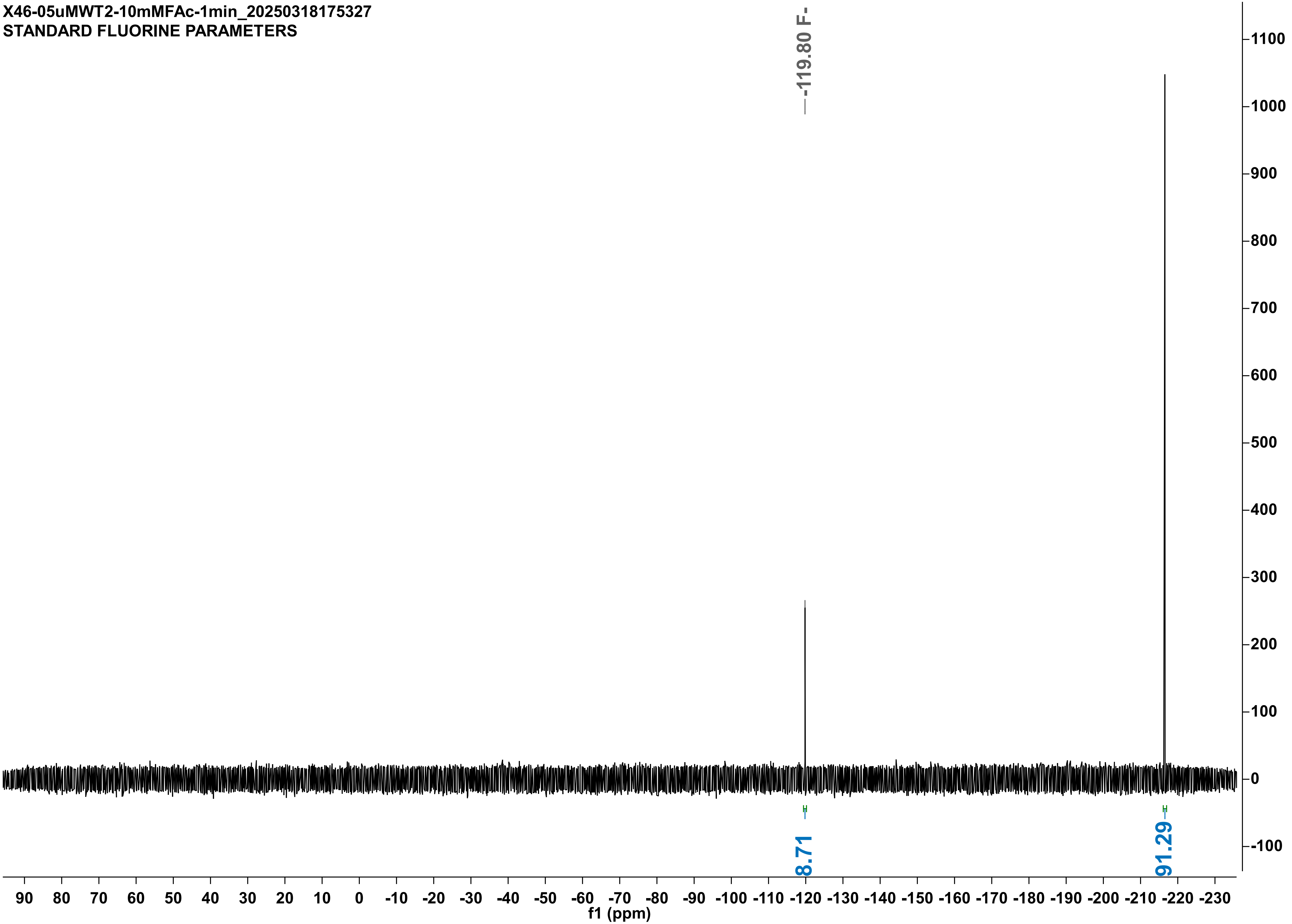

X48-11-4-05uMRPL-10mMFAc-20min_20250411161735
STANDARD FLUORINE PARAMETERS

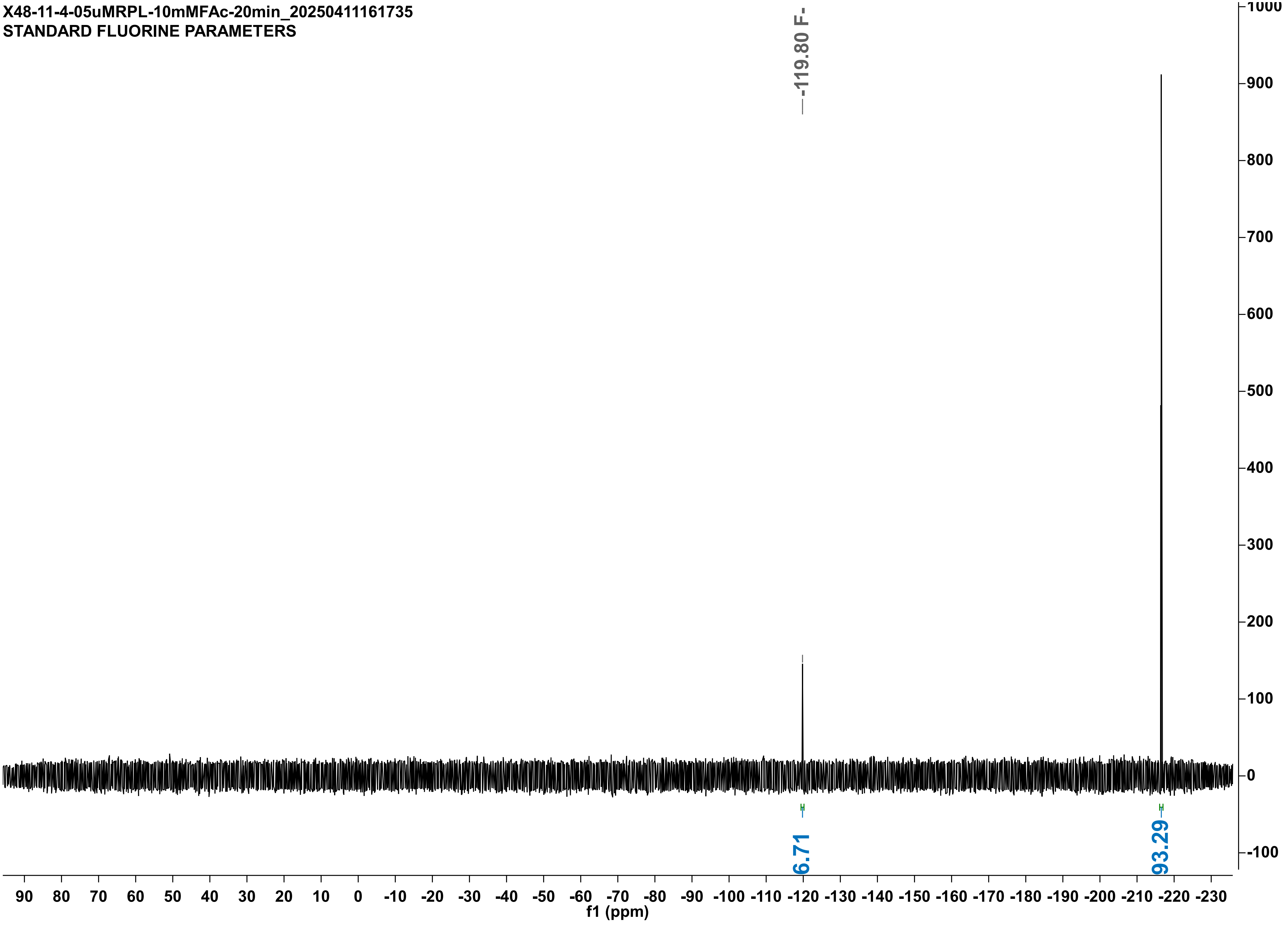

X48-11-4-05uMRPL-DUPLO-10mMFAc-20min_20250411173544
STANDARD FLUORINE PARAMETERS

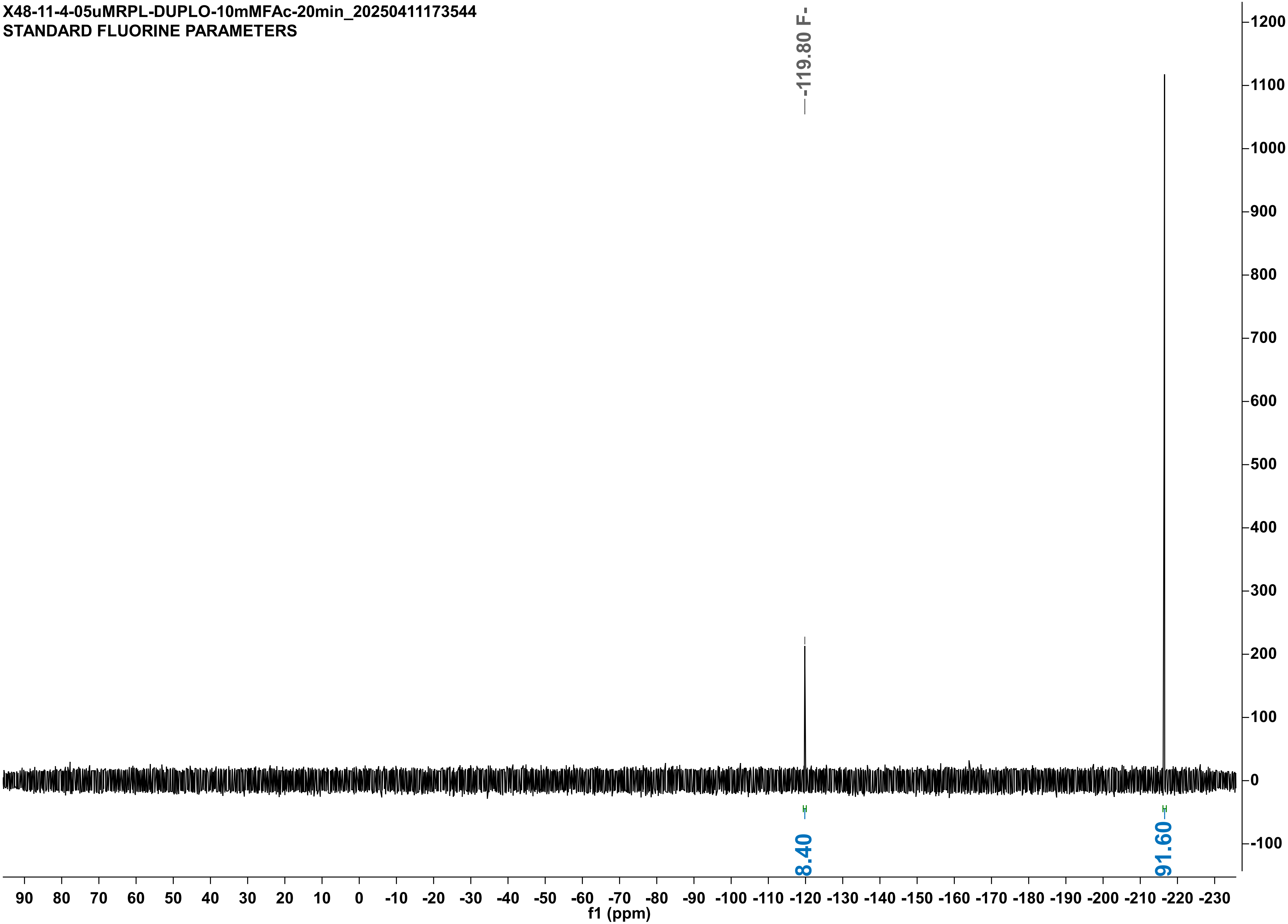

X48-11-4-05uMVW--10mMFAc-5min__20250411140616
STANDARD FLUORINE PARAMETERS

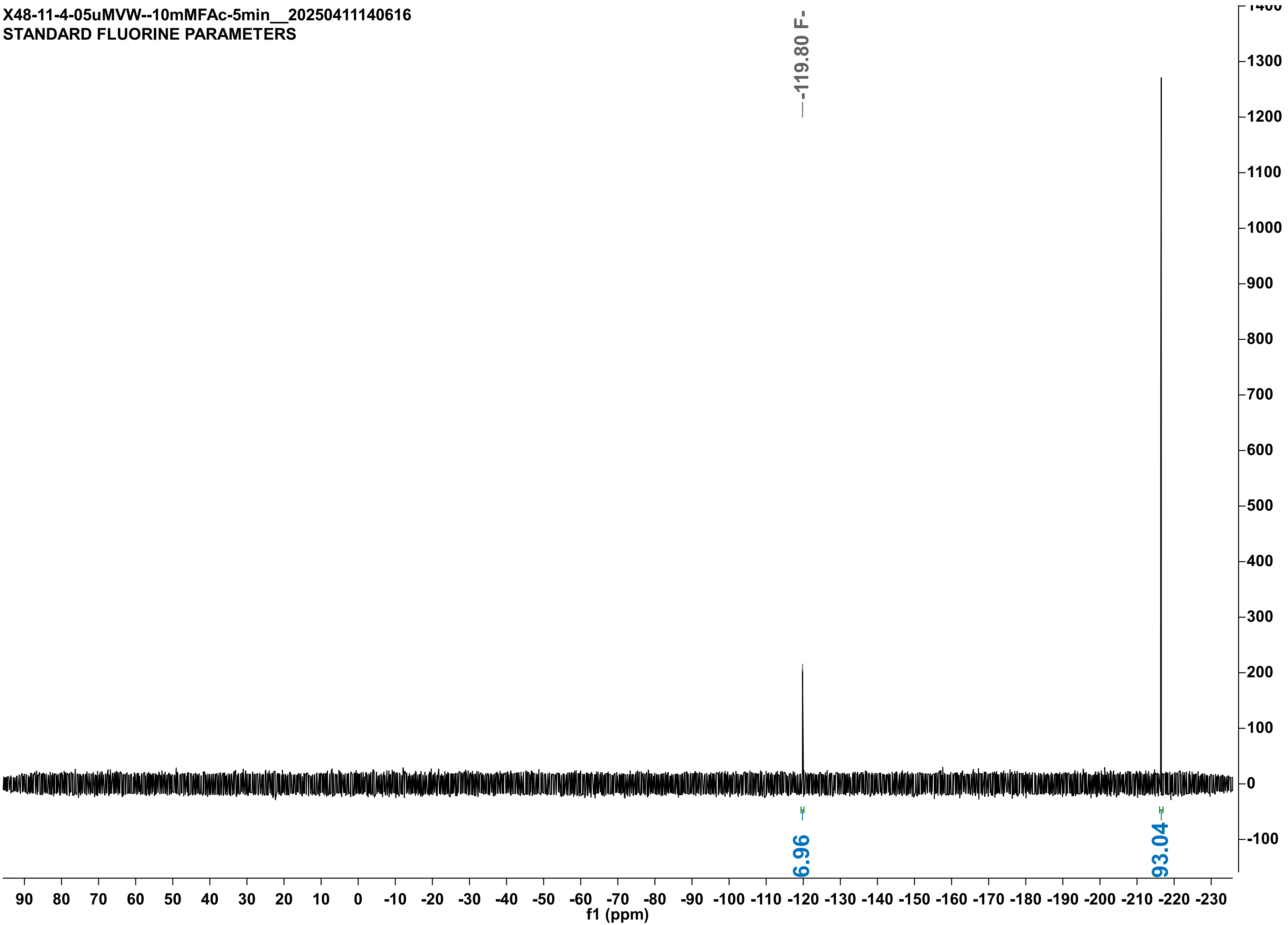

X48-11-4-05uMVW--duplo-10mMFAc-5min_20250411155403
STANDARD FLUORINE PARAMETERS

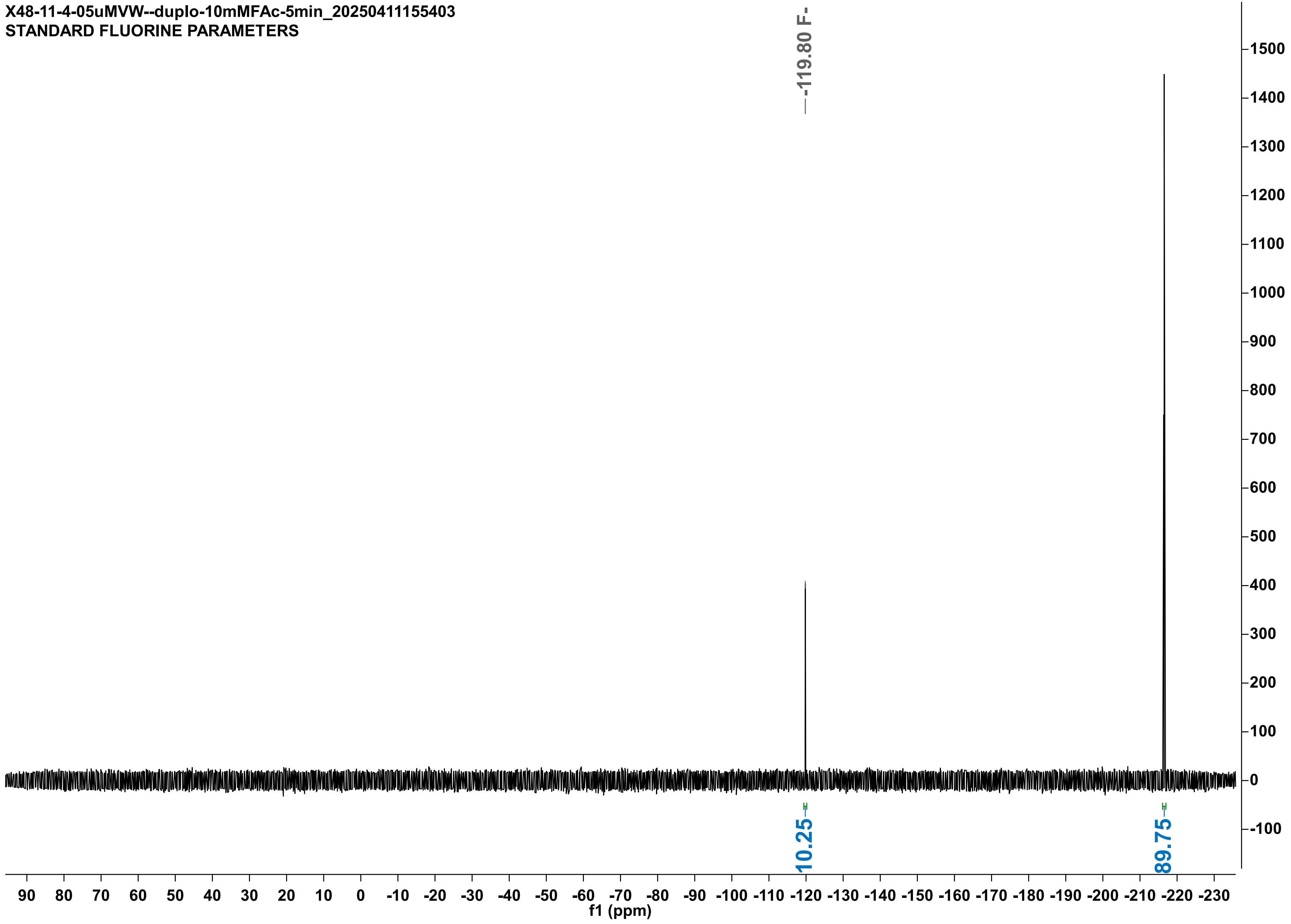

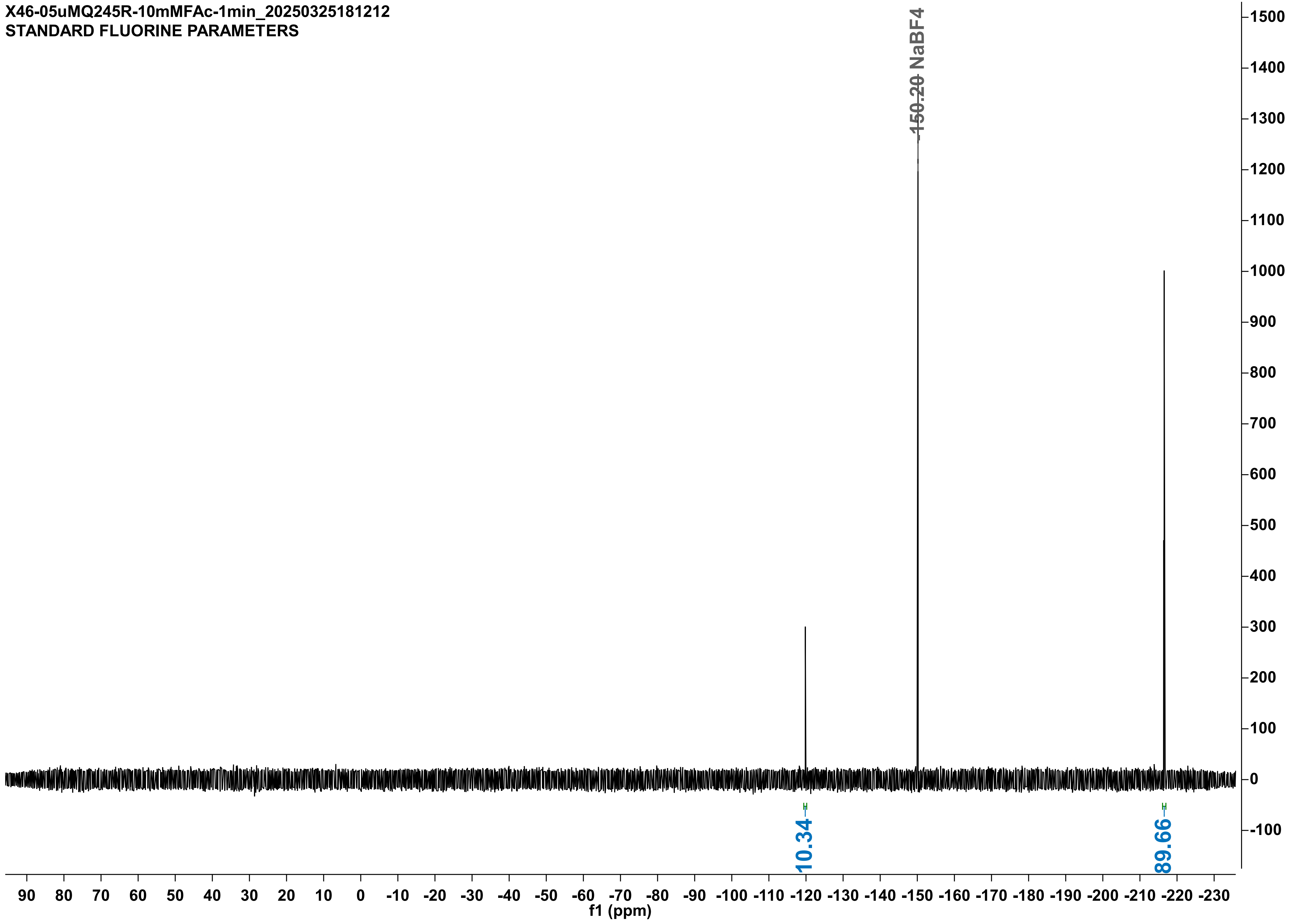
X46-05uMQ245R-10mMFAc-1min_20250325181212
STANDARD FLUORINE PARAMETERS
-150.20 NaBF4
10.34
89.66
f1 (ppm)

X48-16-4-05uMQ245R-10mMFAc-60sec_20250416210642
STANDARD FLUORINE PARAMETERS

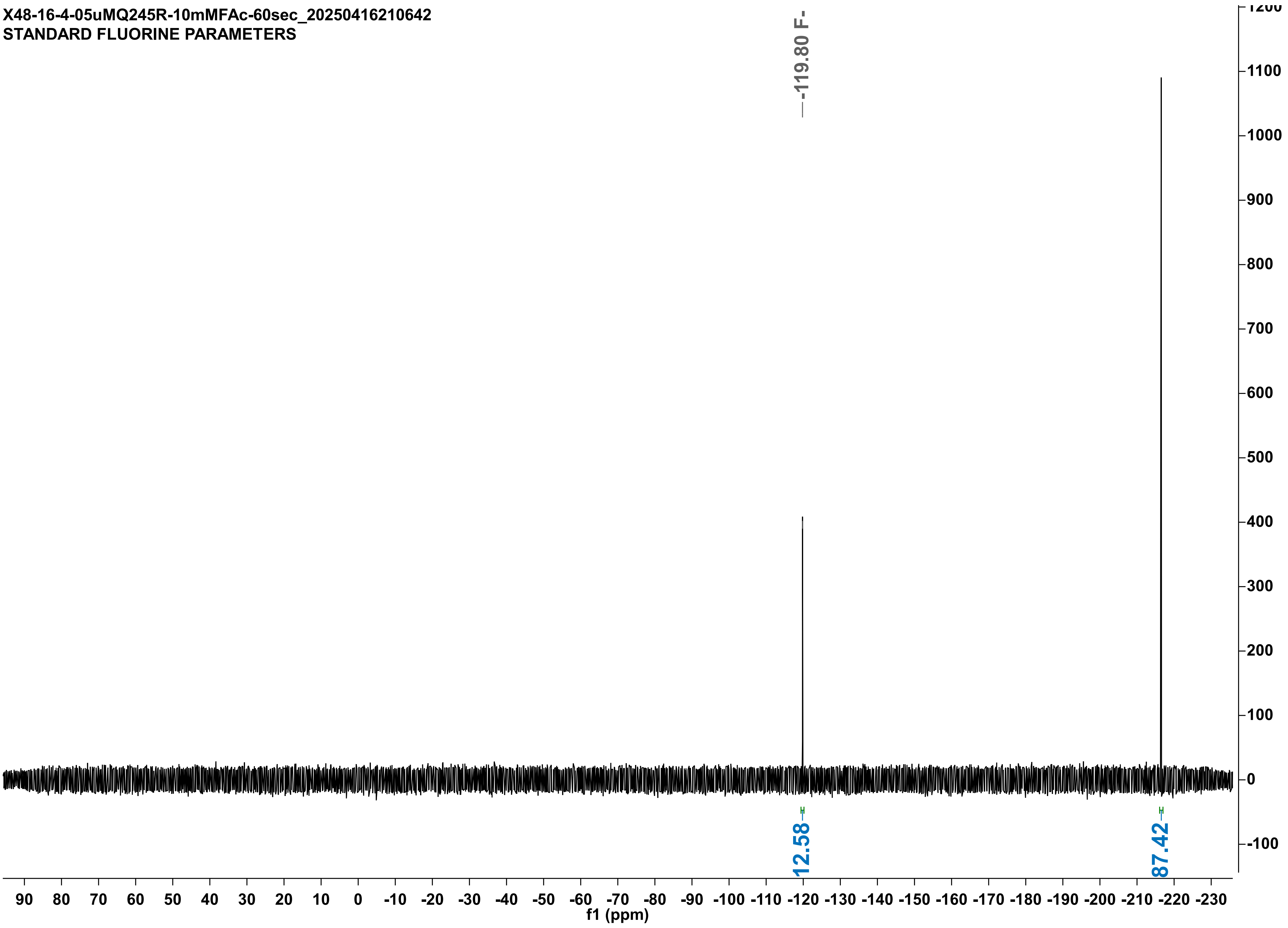

X48-14-4-05uM-ML-10mMFAc-3min_20250414183622
STANDARD FLUORINE PARAMETERS

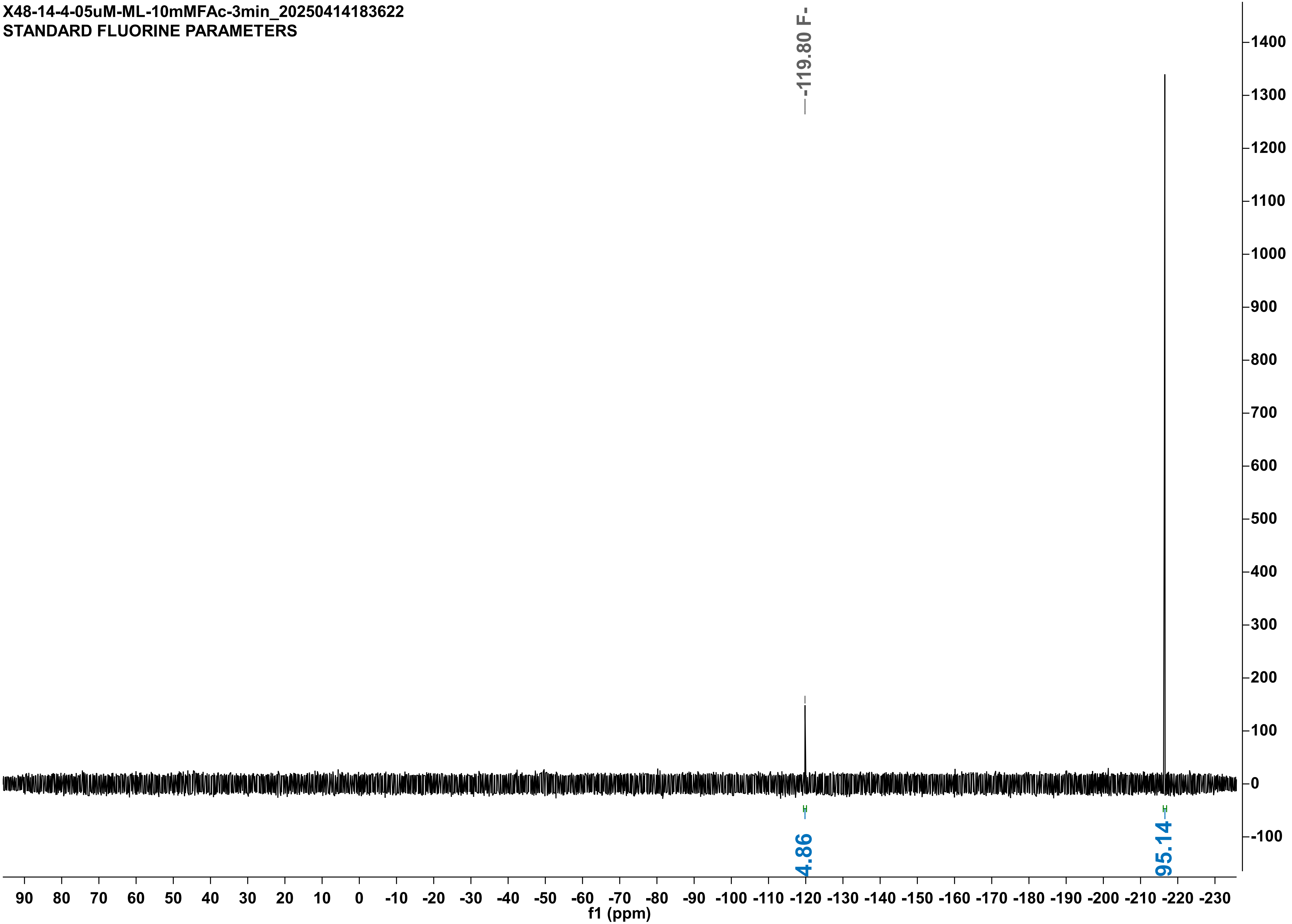

X48-14-4-05uM-ML-DUPLO-10mMFAc-3min_20250414190702
STANDARD FLUORINE PARAMETERS

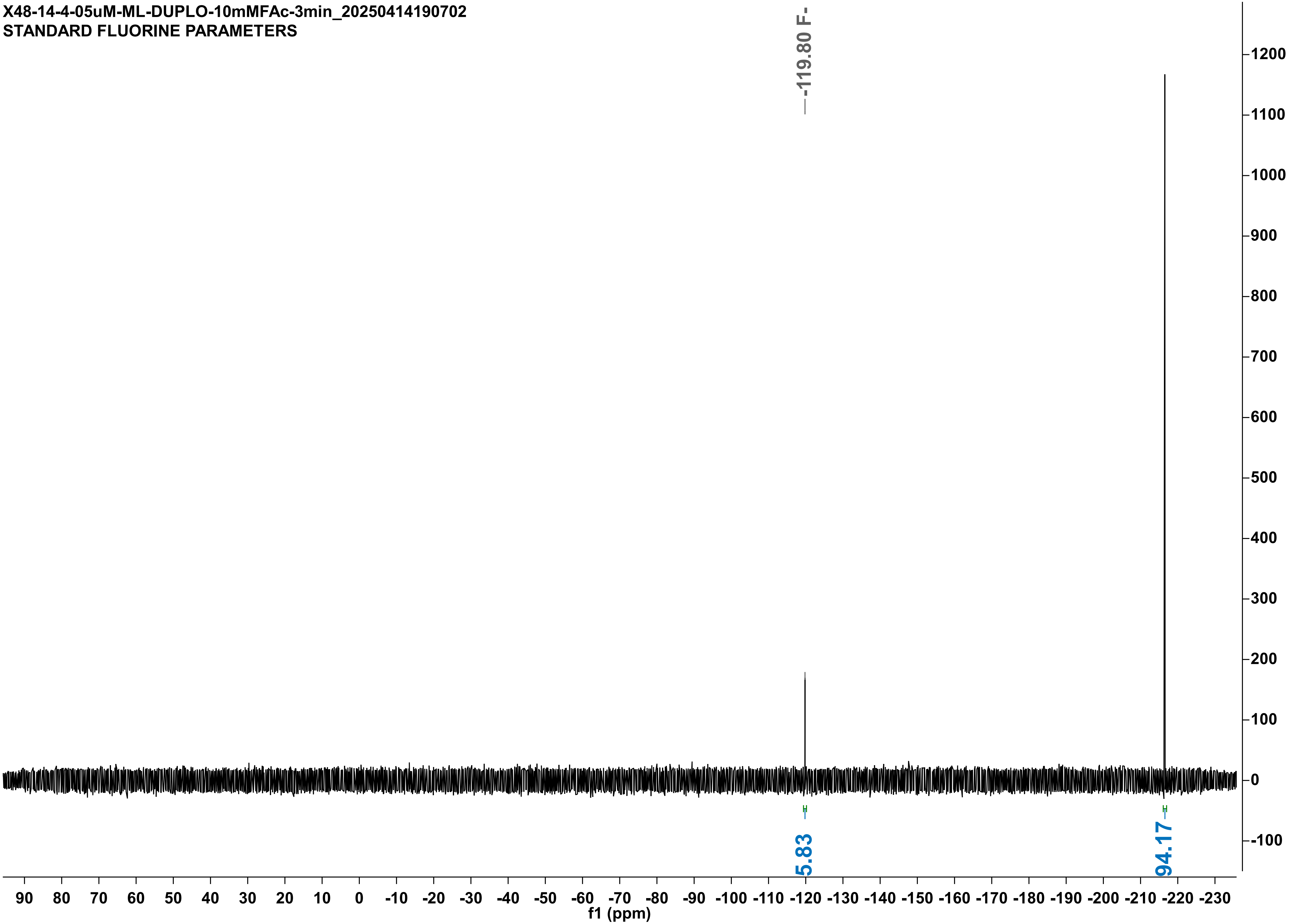

X48-14-4-05uMRHL-10mMFAc-7min_20250414190743
STANDARD FLUORINE PARAMETERS

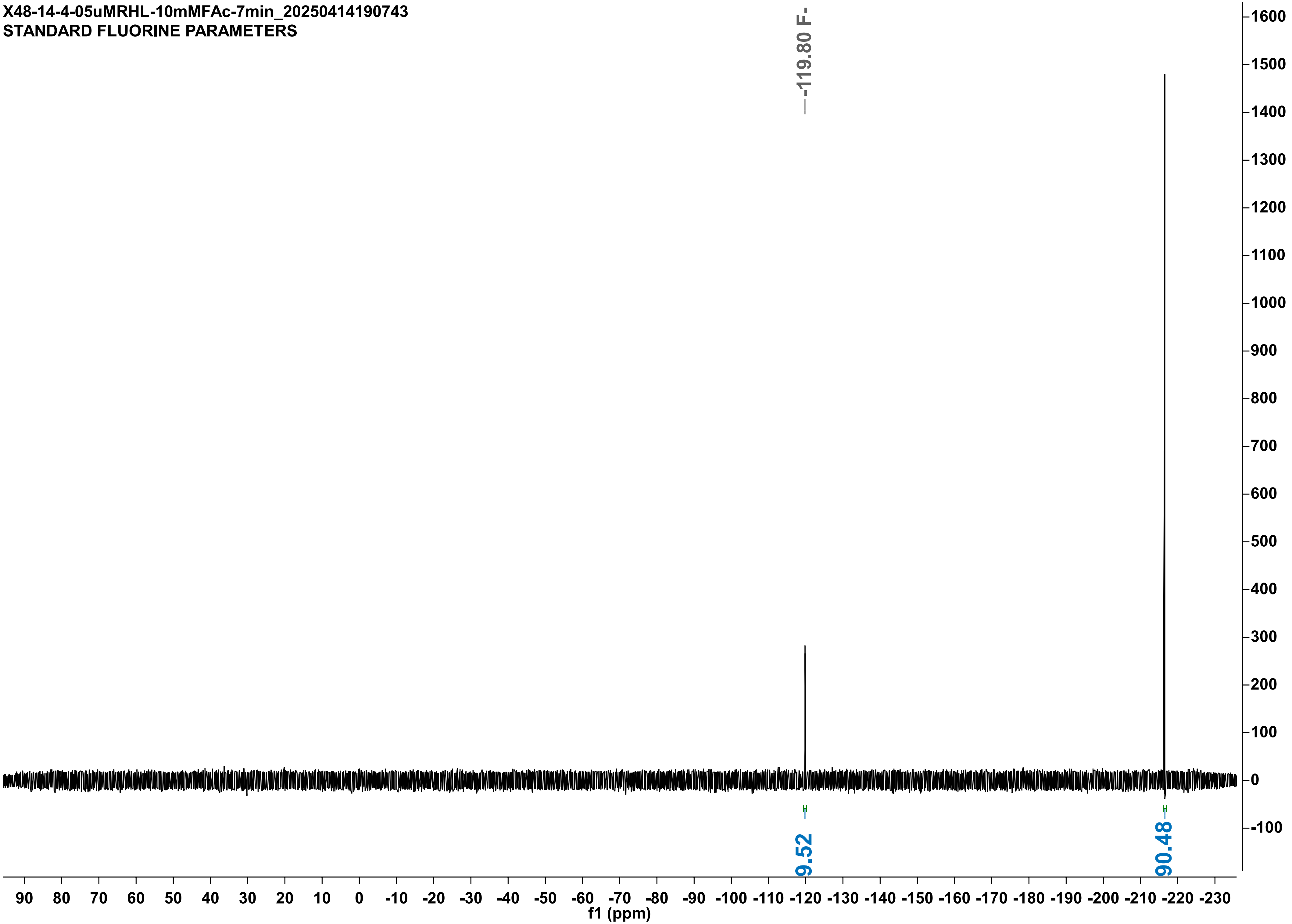

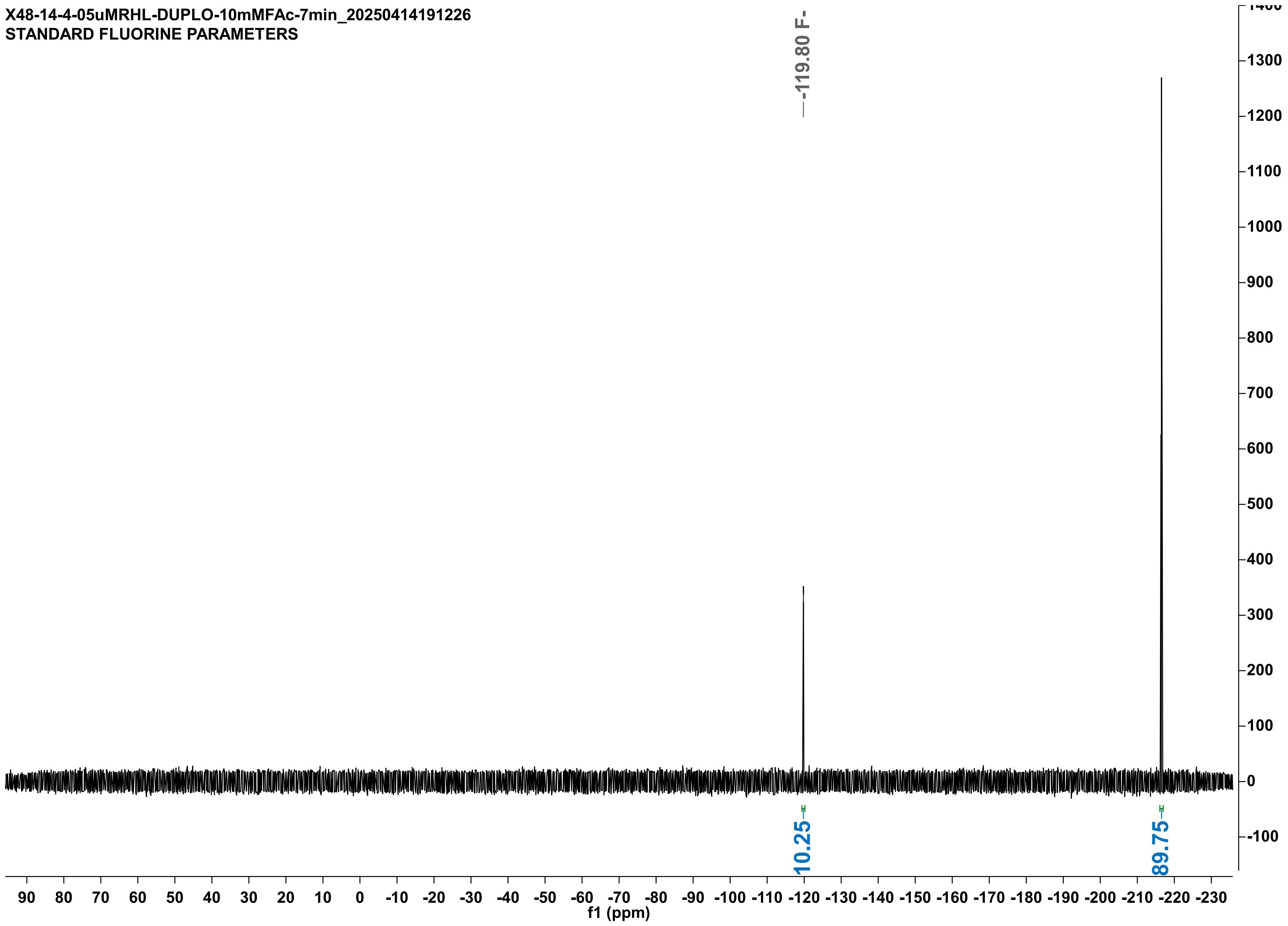
X48-14-4-05uMRHL-DUPLO-10mMFAc-7min_20250414191226
STANDARD FLUORINE PARAMETERS
-119.80 F-
10.25
89.75
f1 (ppm)

X48-11-4-05uMQ245A-10mMFAc-45sec_20250411113955
STANDARD FLUORINE PARAMETERS

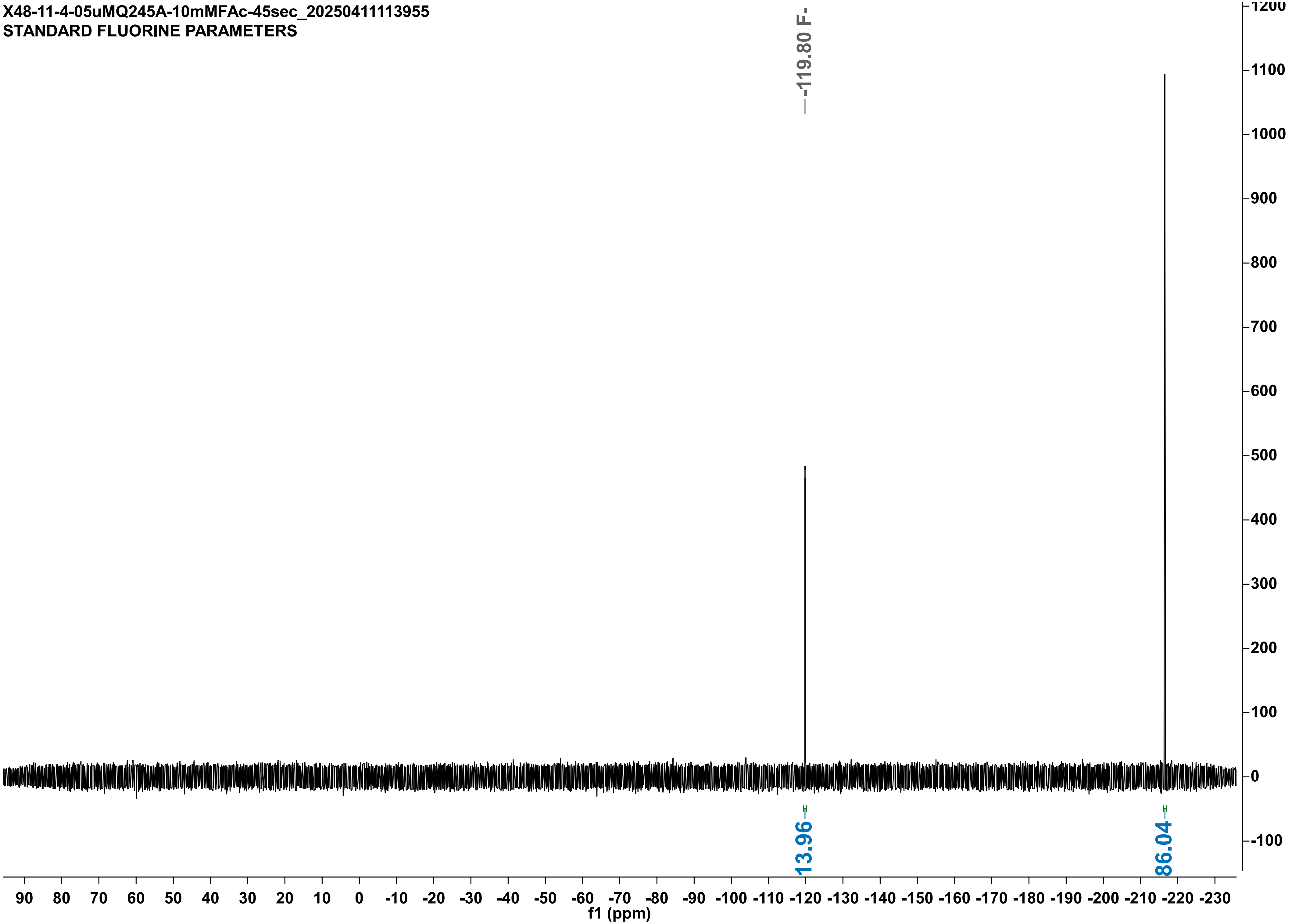

X48-11-4-05uMQ245A-DUPLO-10mMFAc-45sec_20250411132637
STANDARD FLUORINE PARAMETERS

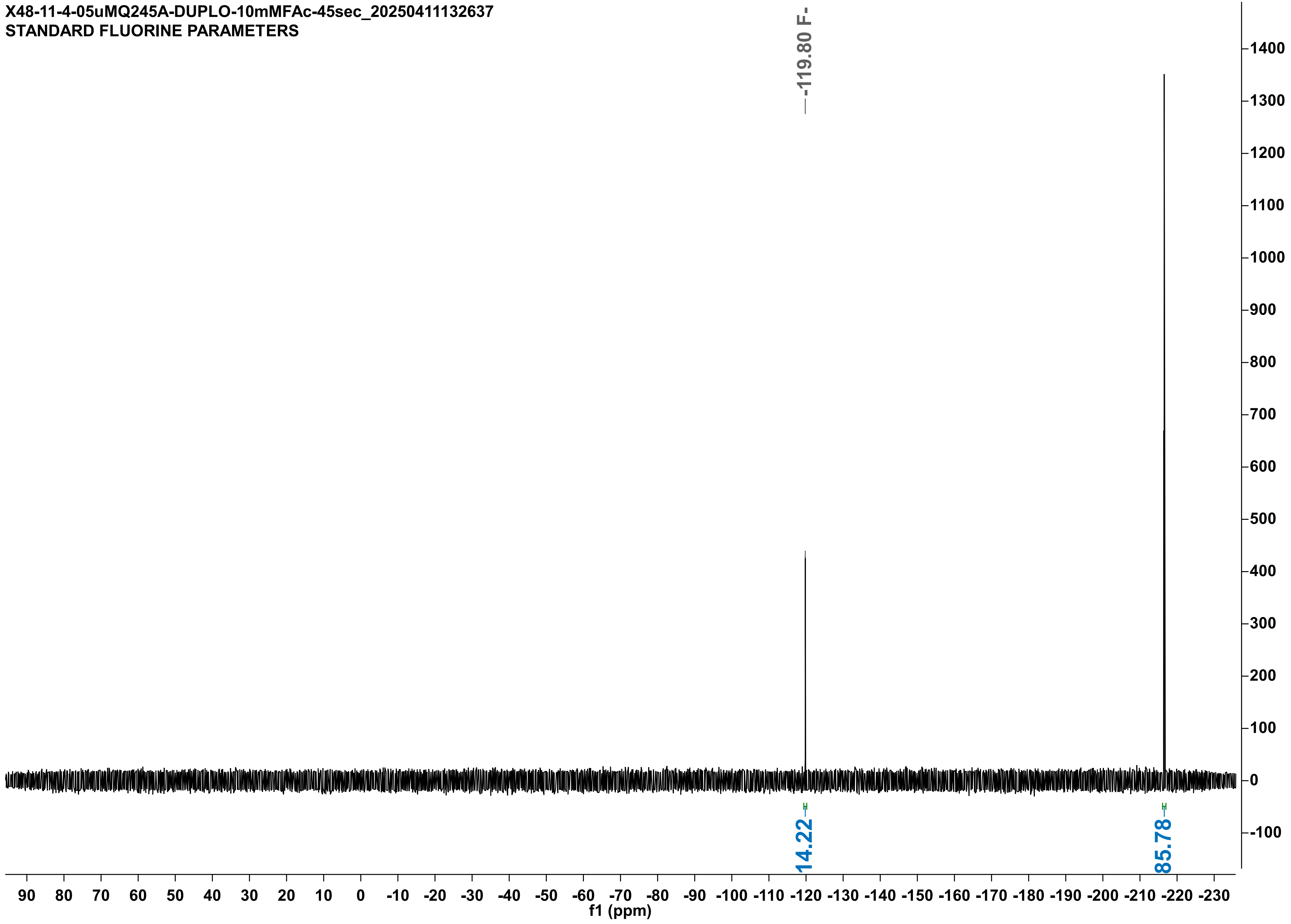

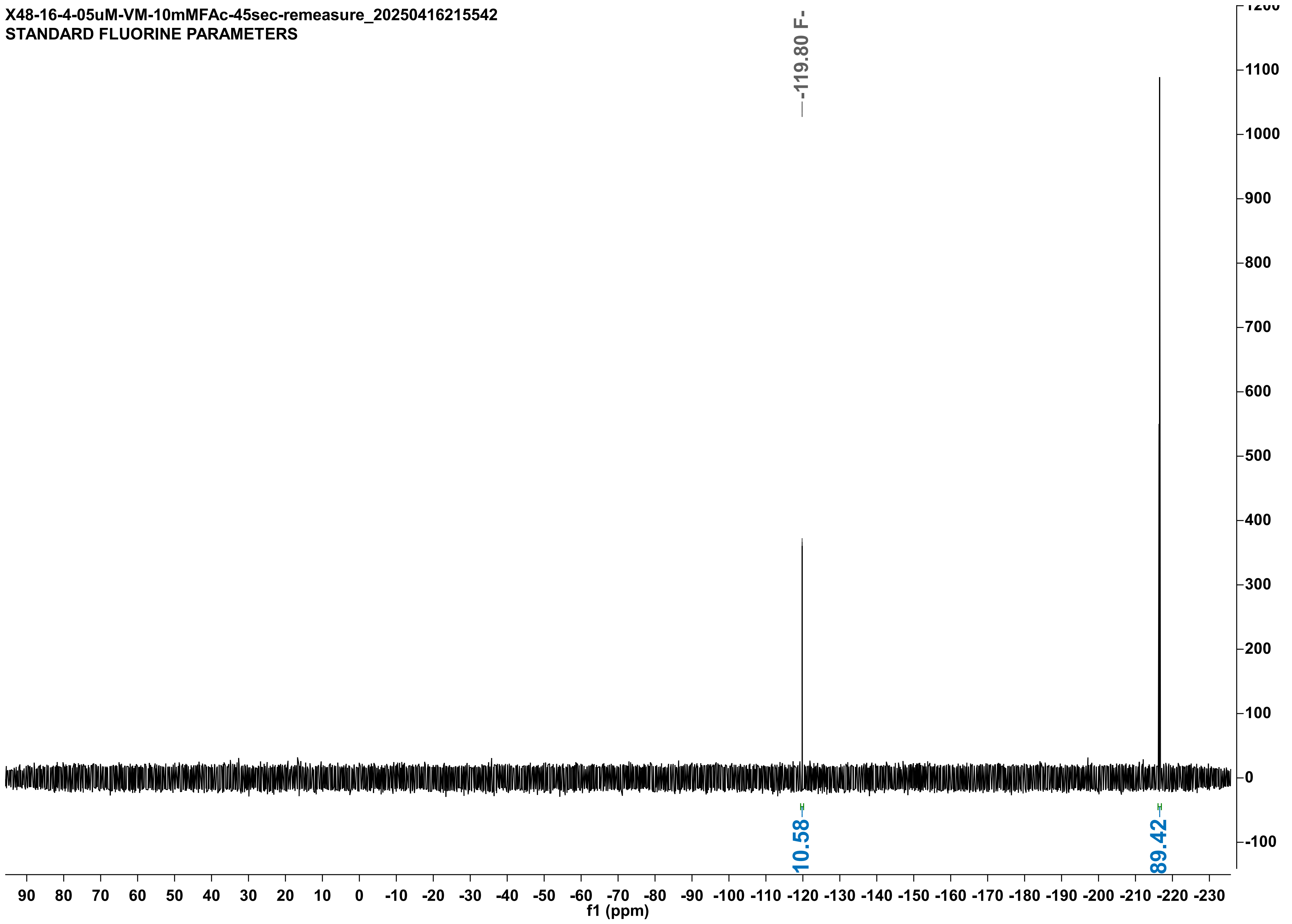

X48-16-4-05uM-VM-10mMFAc-45sec-remeasure_20250416215542
STANDARD FLUORINE PARAMETERS
-119.80 F-
10.58
89.42
f1 (ppm)

X48-16-4-05uM-VM-DUP-10mMFAc-45sec_20250416203831
STANDARD FLUORINE PARAMETERS

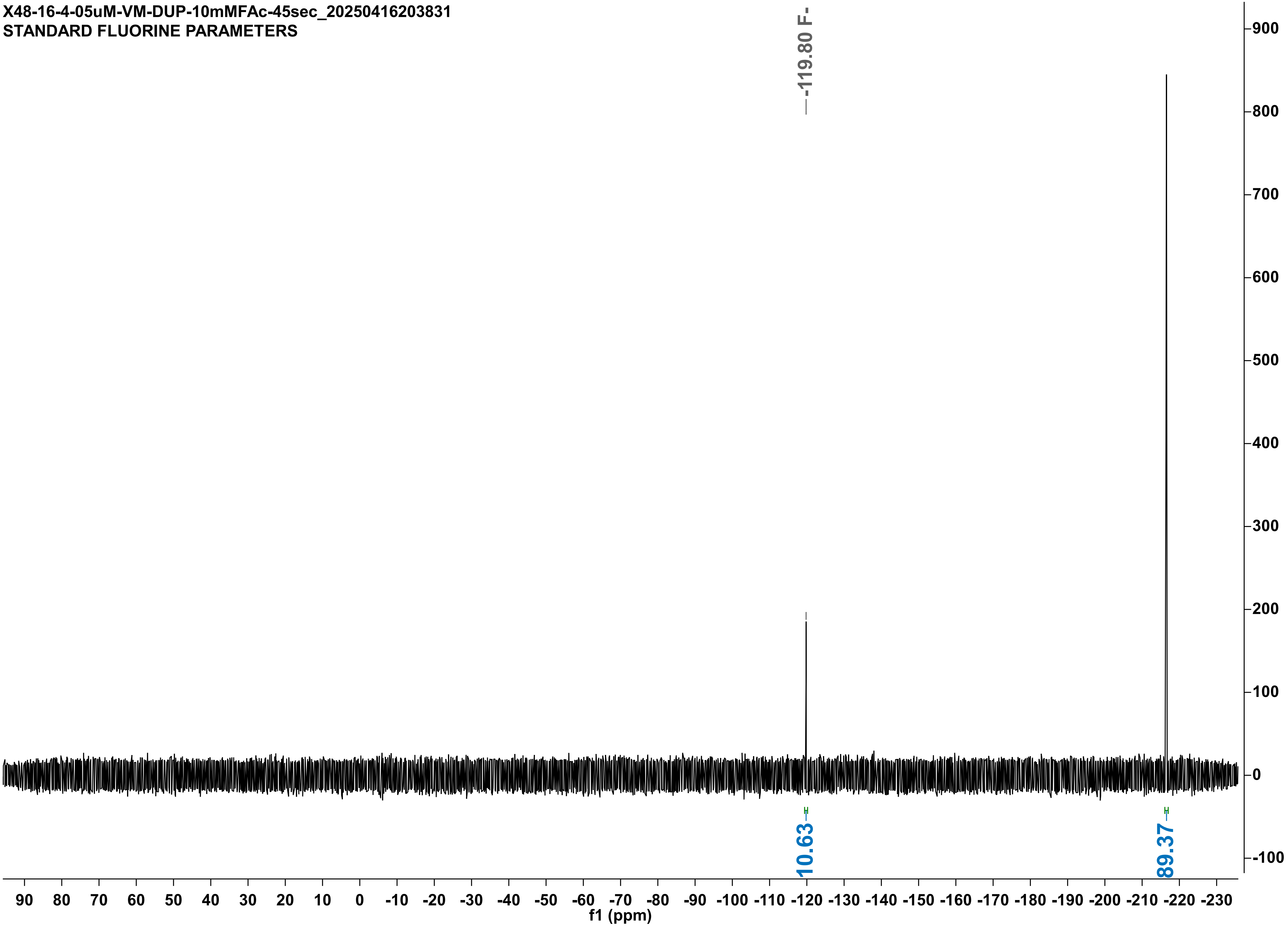

X46-05uMRLM-10mMFAc-1min_20250320160803
STANDARD FLUORINE PARAMETERS

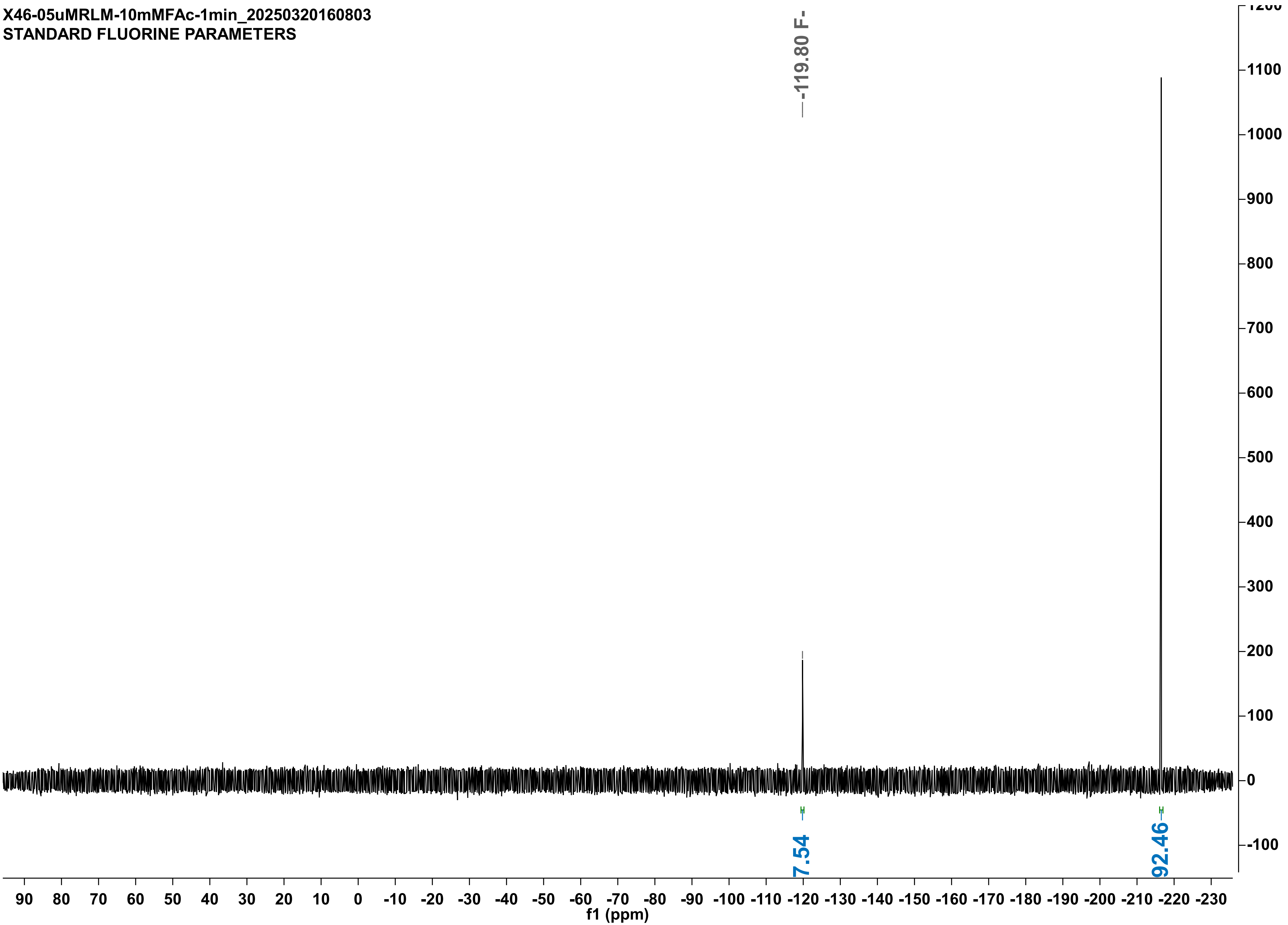

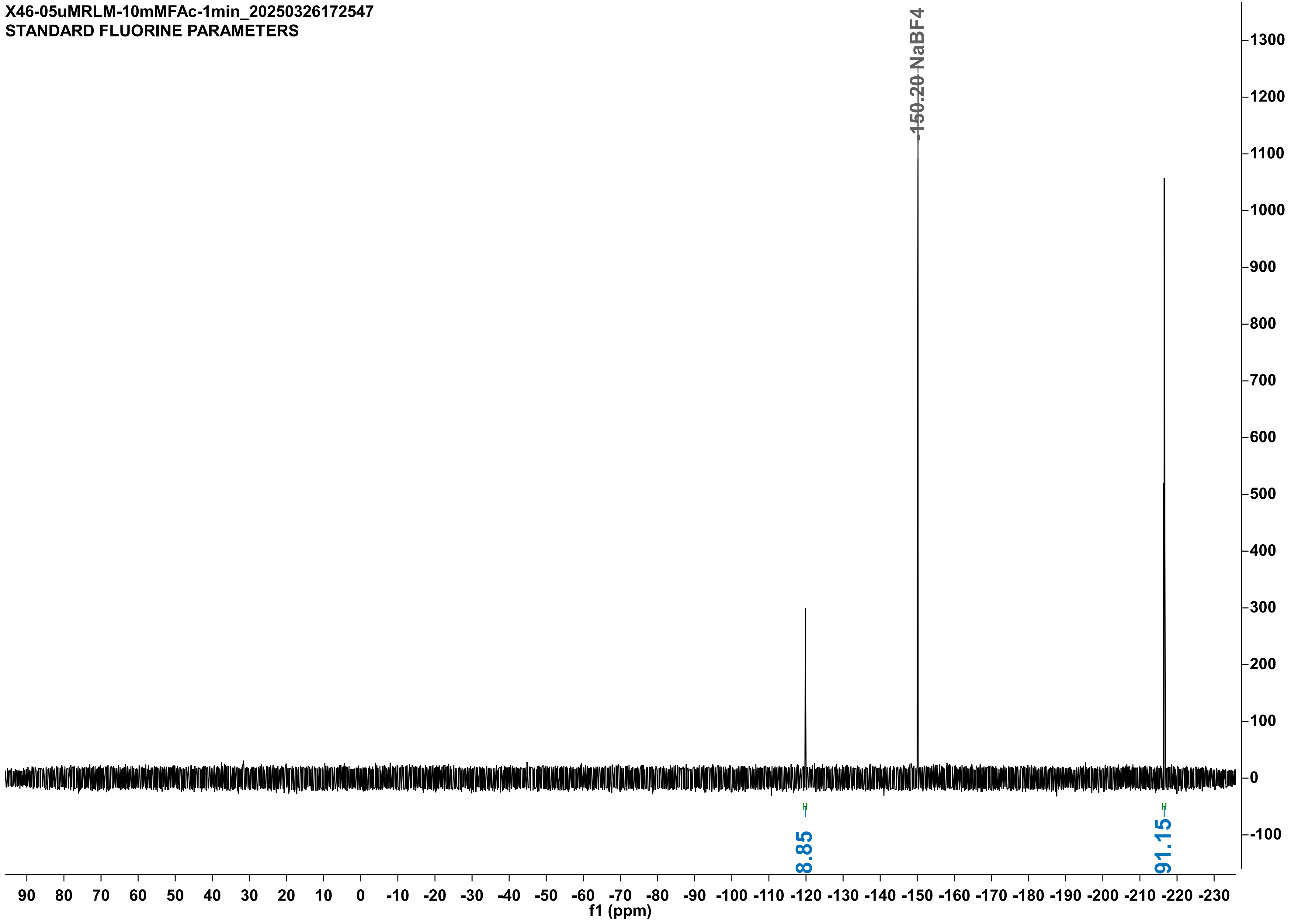
X46-05uMRLM-10mMFAc-1min_20250326172547
STANDARD FLUORINE PARAMETERS
-150.20 NaBF4
8.85
91.15
f1 (ppm)

X48-15-4-05uMQ245G-10mMFAc-1min_20250415191946
STANDARD FLUORINE PARAMETERS

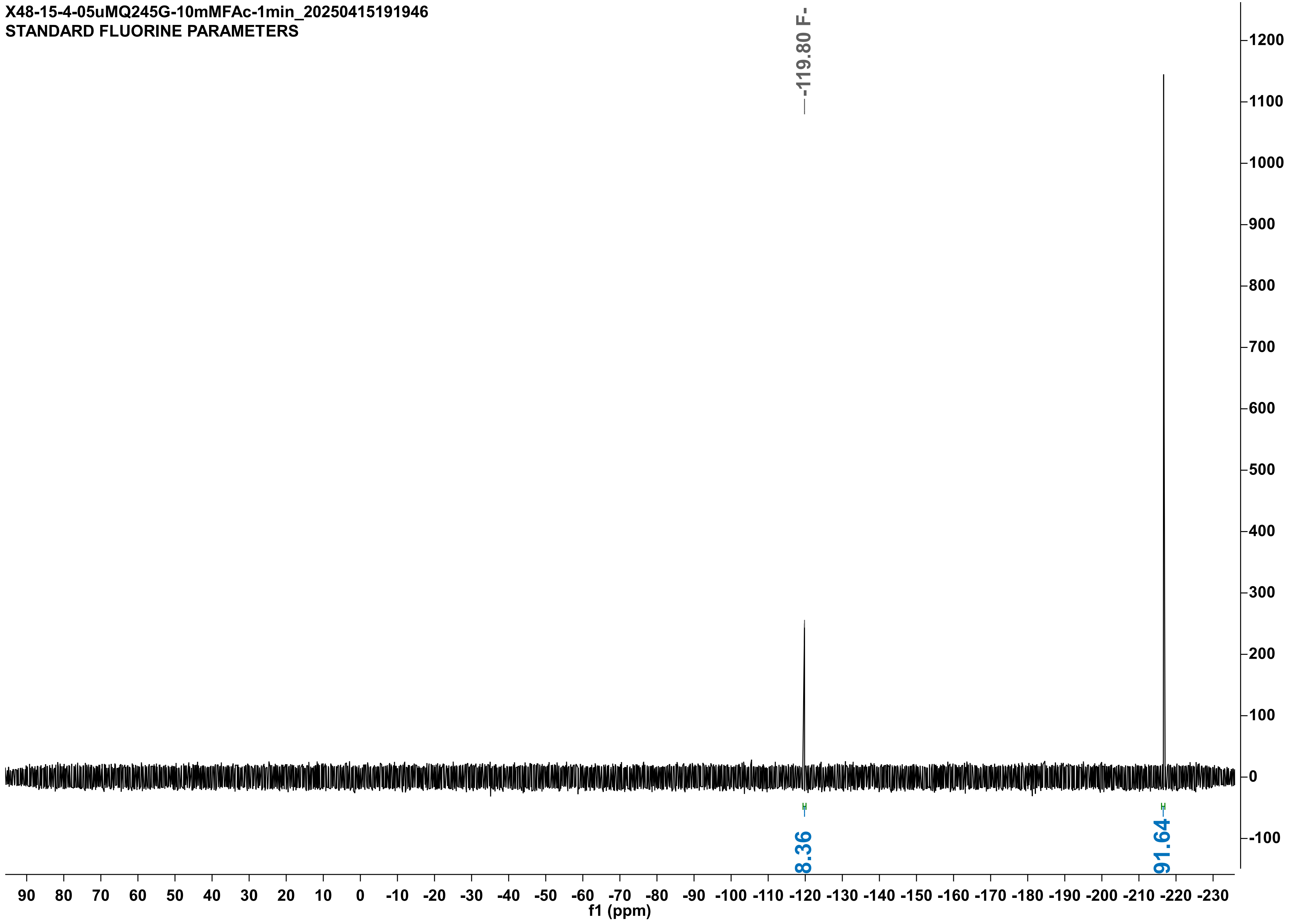

X48-15-4-05uMQ245G-DUP-10mMFAc-1min_20250415192016
STANDARD FLUORINE PARAMETERS

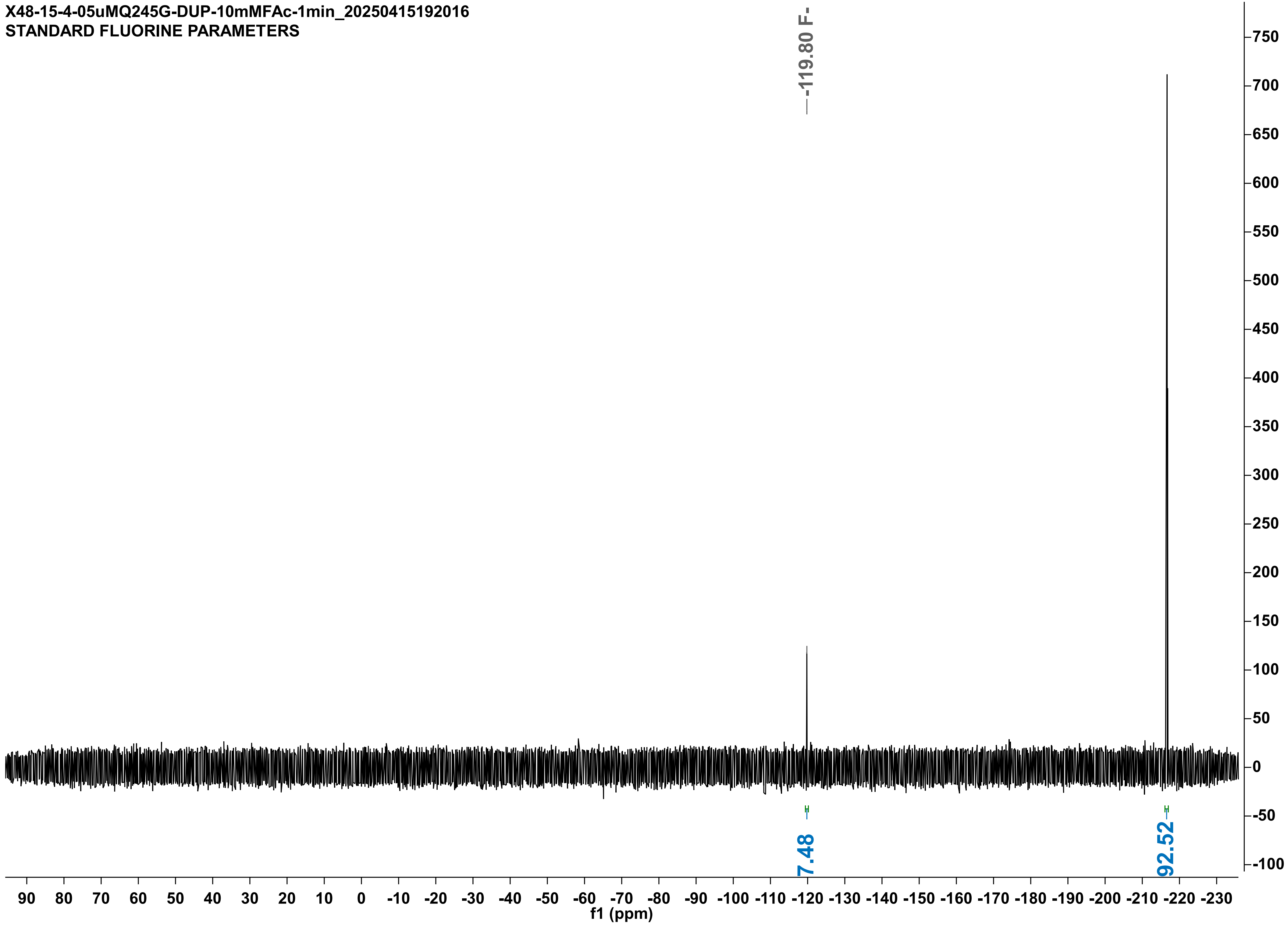

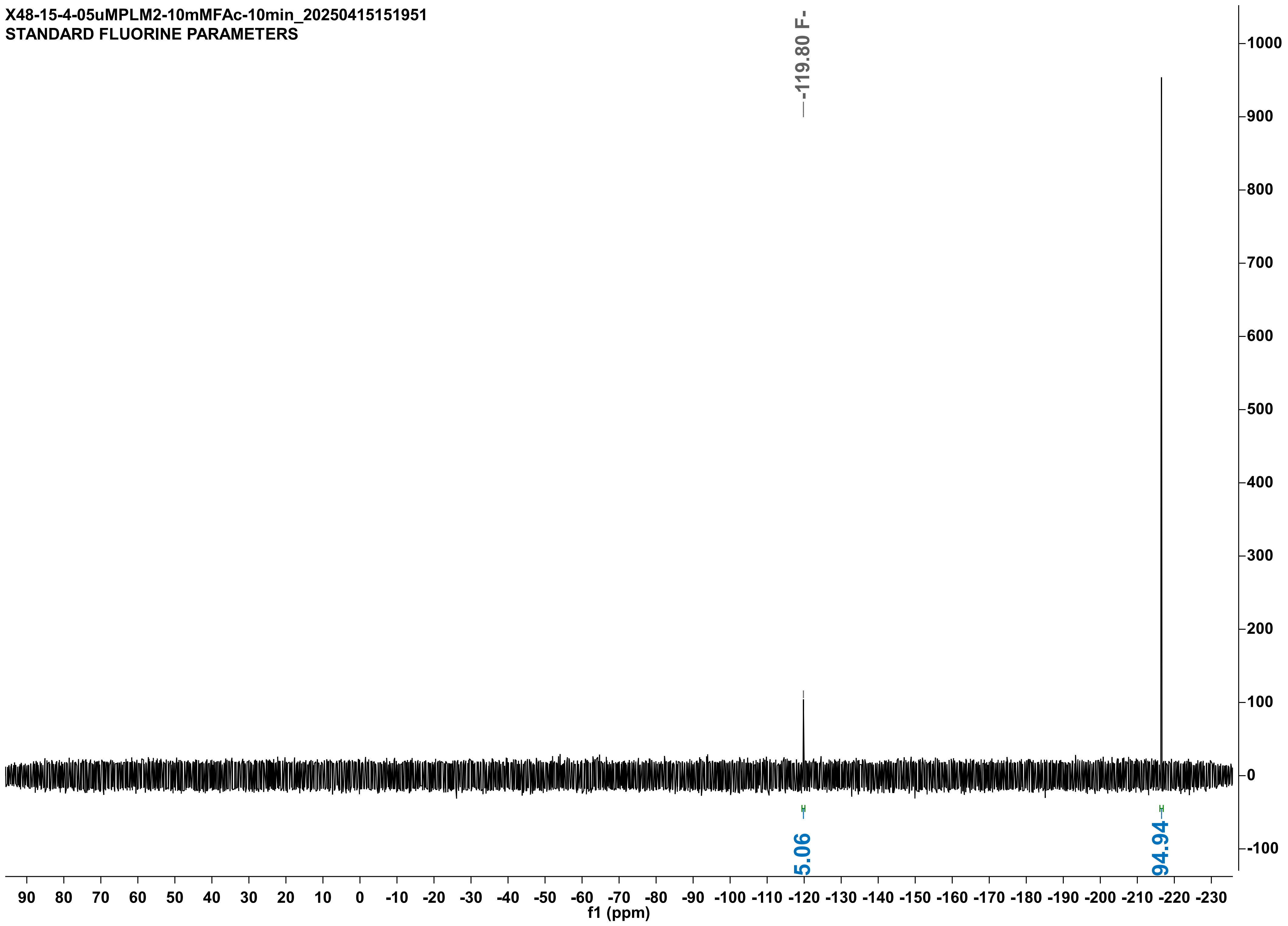
X48-15-4-05uMPLM2-10mMFAc-10min_20250415151951
STANDARD FLUORINE PARAMETERS
-119.80 F-
5.06
94.94
f1 (ppm)

X48-16-4-05uMPLM2-DUP-10mMFAc-10min_20250416210748
STANDARD FLUORINE PARAMETERS

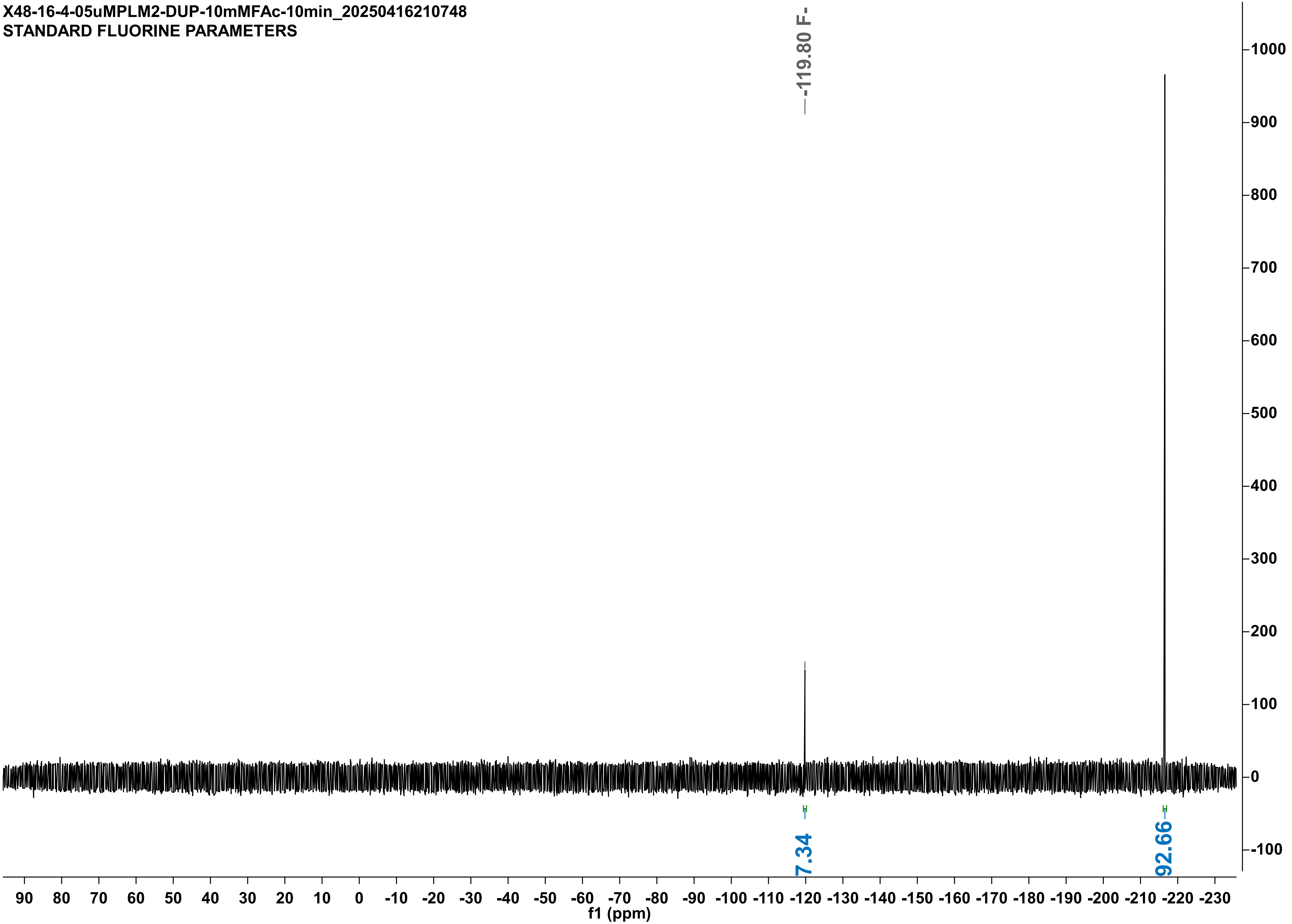

X48-15-4-05uMLW--10mMFAc-2min_20250415162833
STANDARD FLUORINE PARAMETERS

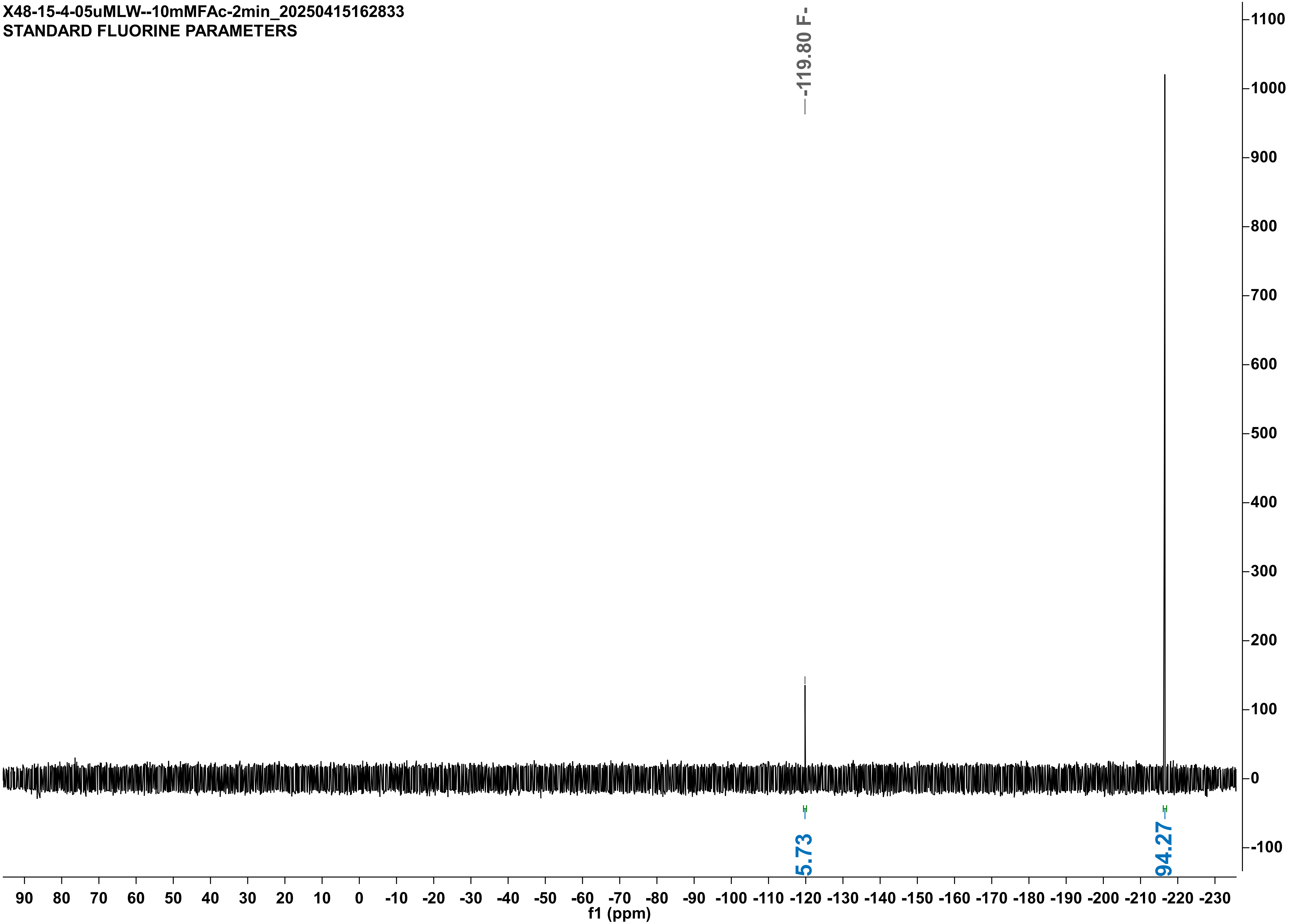

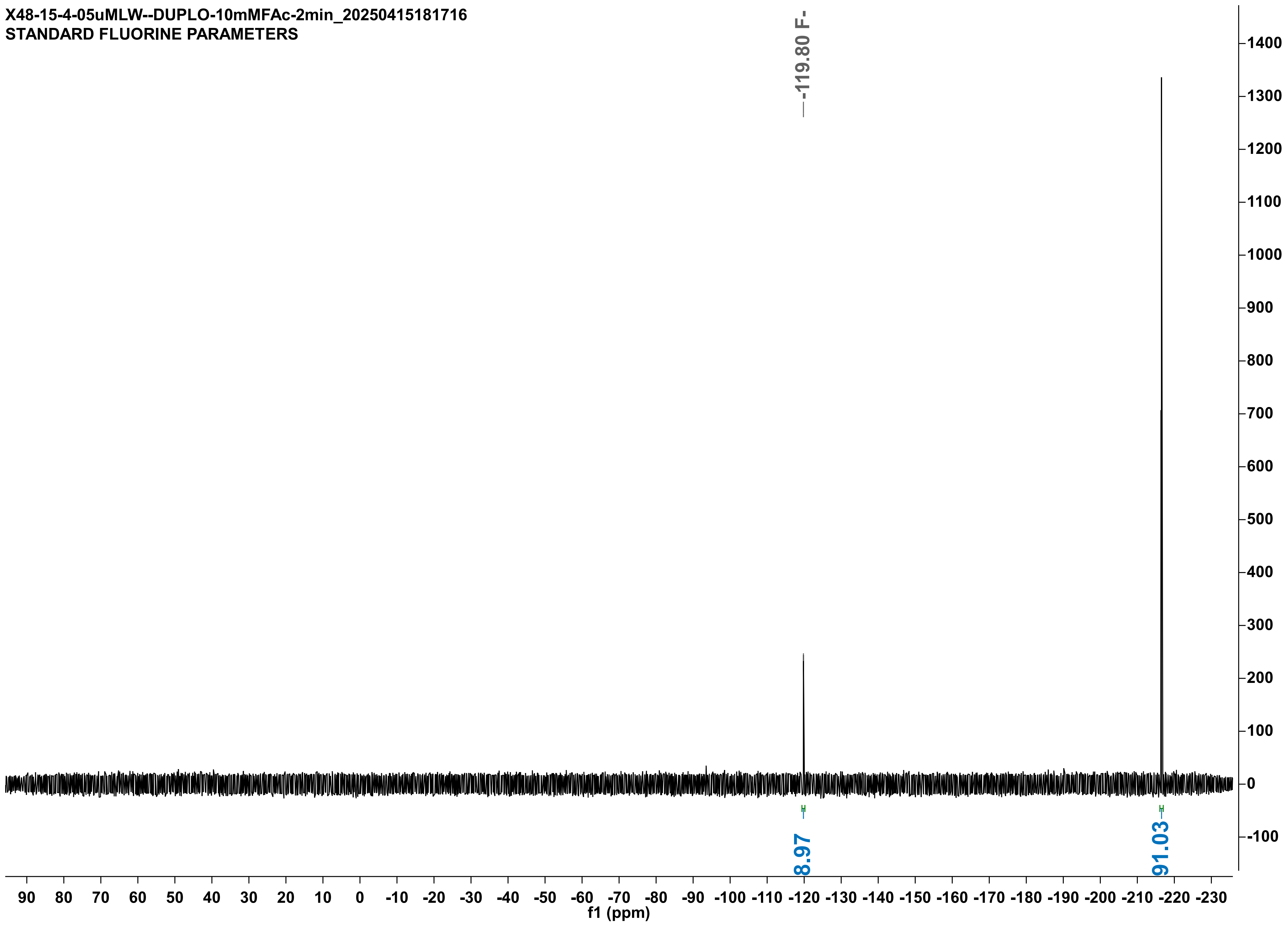

X48-15-4-05uMLW--DUPLO-10mMFAc-2min_20250415181716
STANDARD FLUORINE PARAMETERS
-119.80 F-
8.97
91.03
f1 (ppm)

X48-16-4-05uMLVM-10mMFAc-45sec_20250416203054
STANDARD FLUORINE PARAMETERS

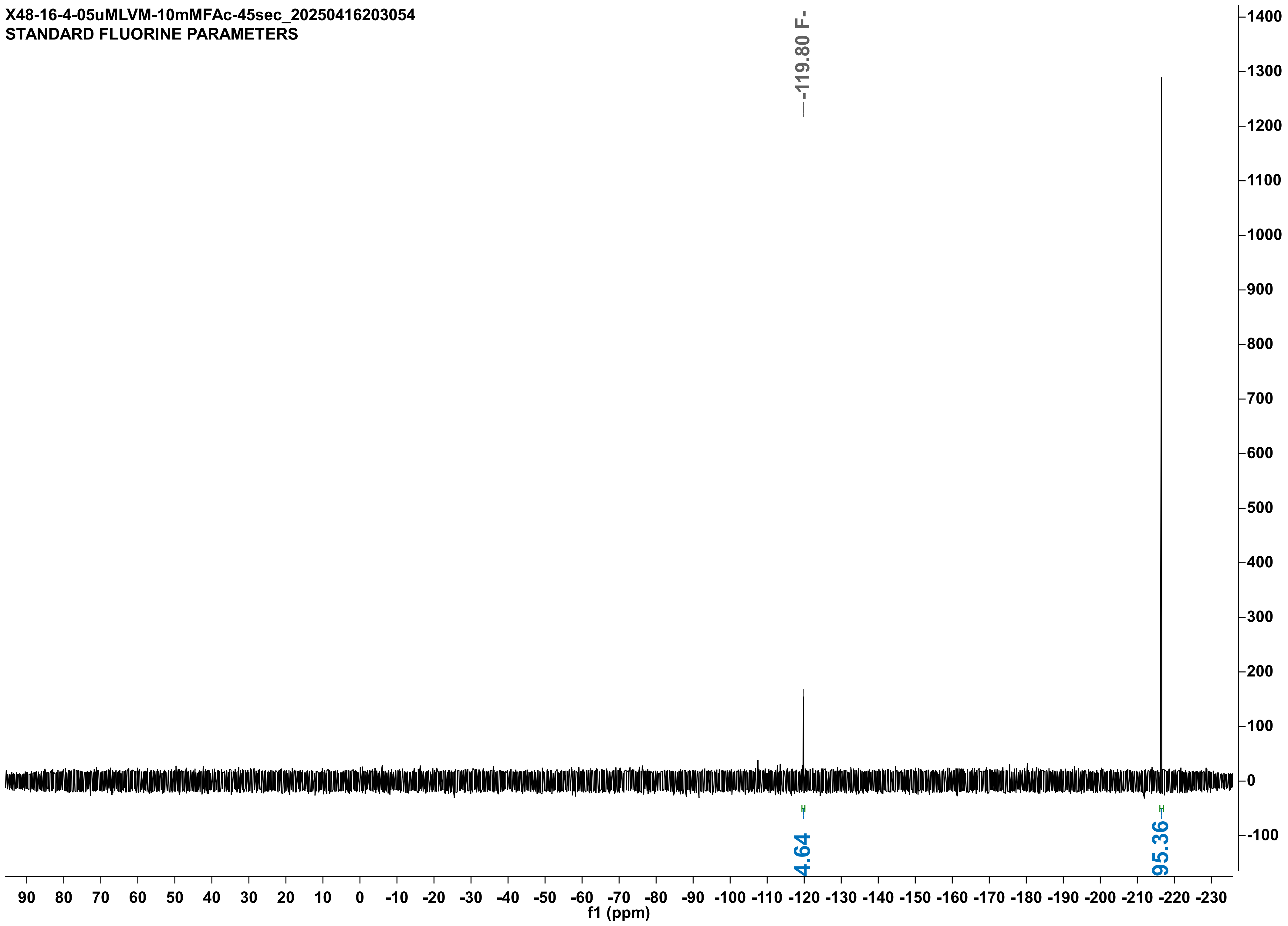

X48-16-4-05uMLVM-DUP-10mMFAc-45sec_20250416203114
STANDARD FLUORINE PARAMETERS

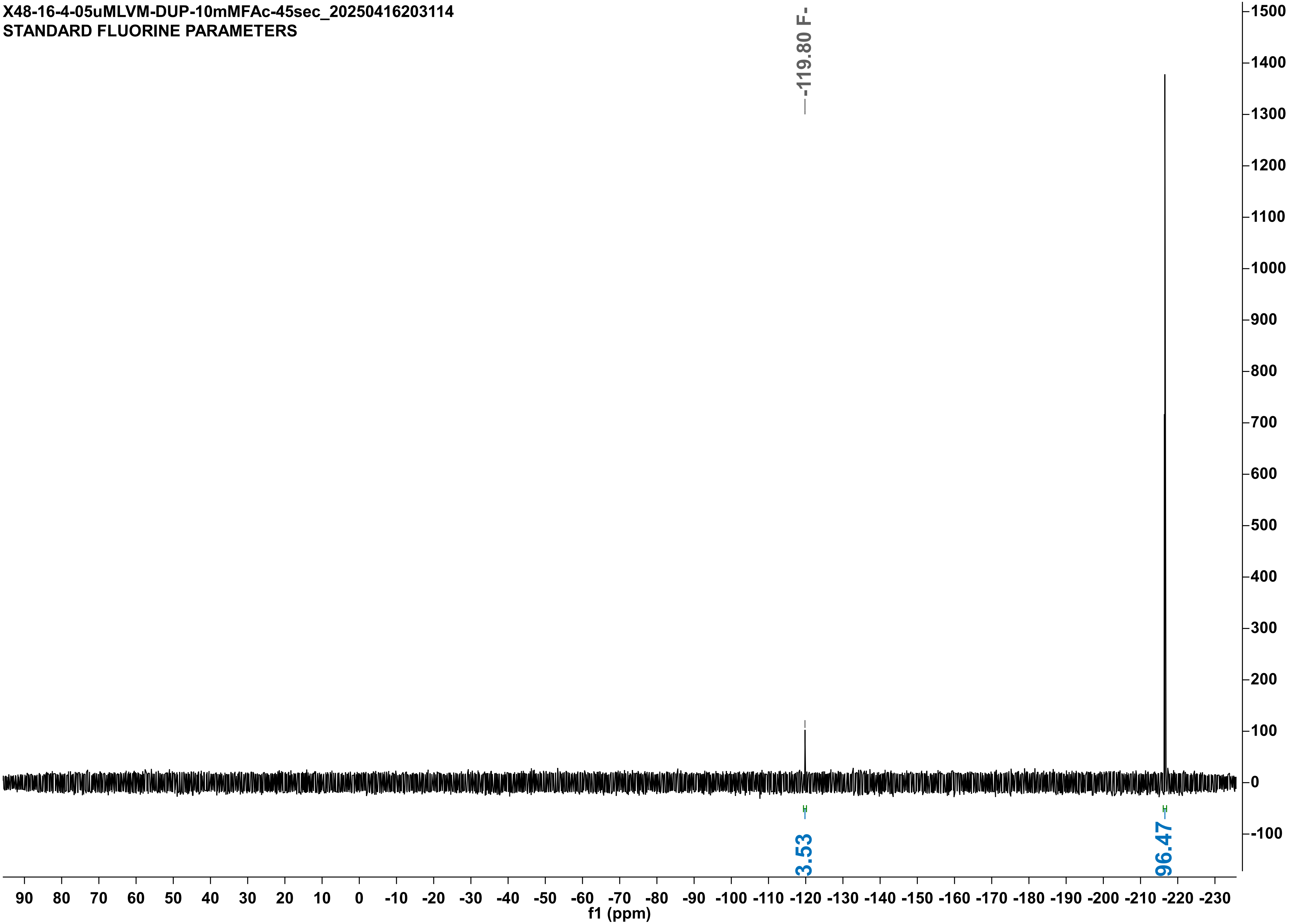

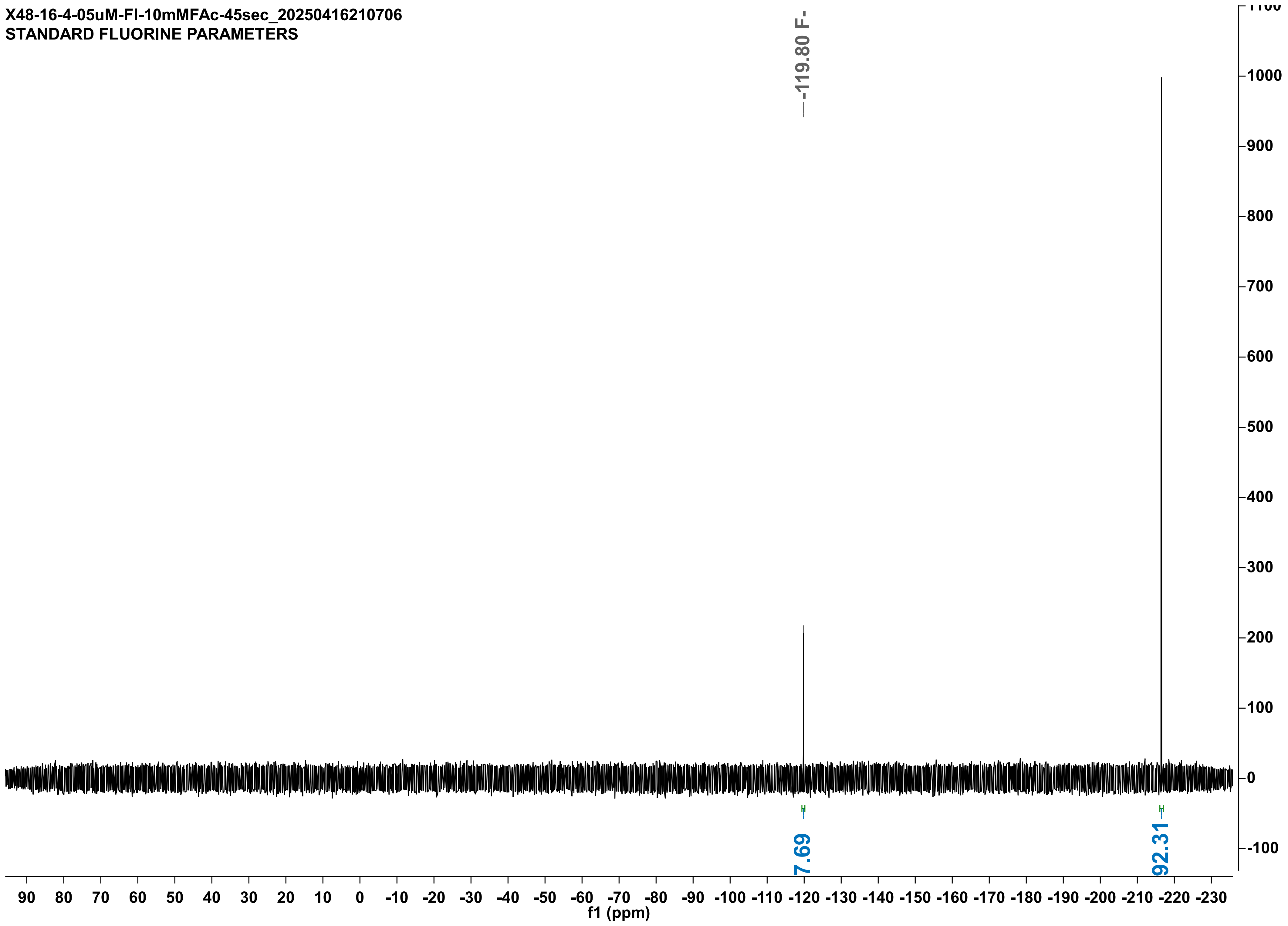

X48-16-4-05uM-FI-10mMFAc-45sec_20250416210706
STANDARD FLUORINE PARAMETERS
-119.80 F-
7.69
92.31
f1 (ppm)

X48-16-4-05uM-FI-DUPLO-10mMFAc-45sec_20250416210716
STANDARD FLUORINE PARAMETERS

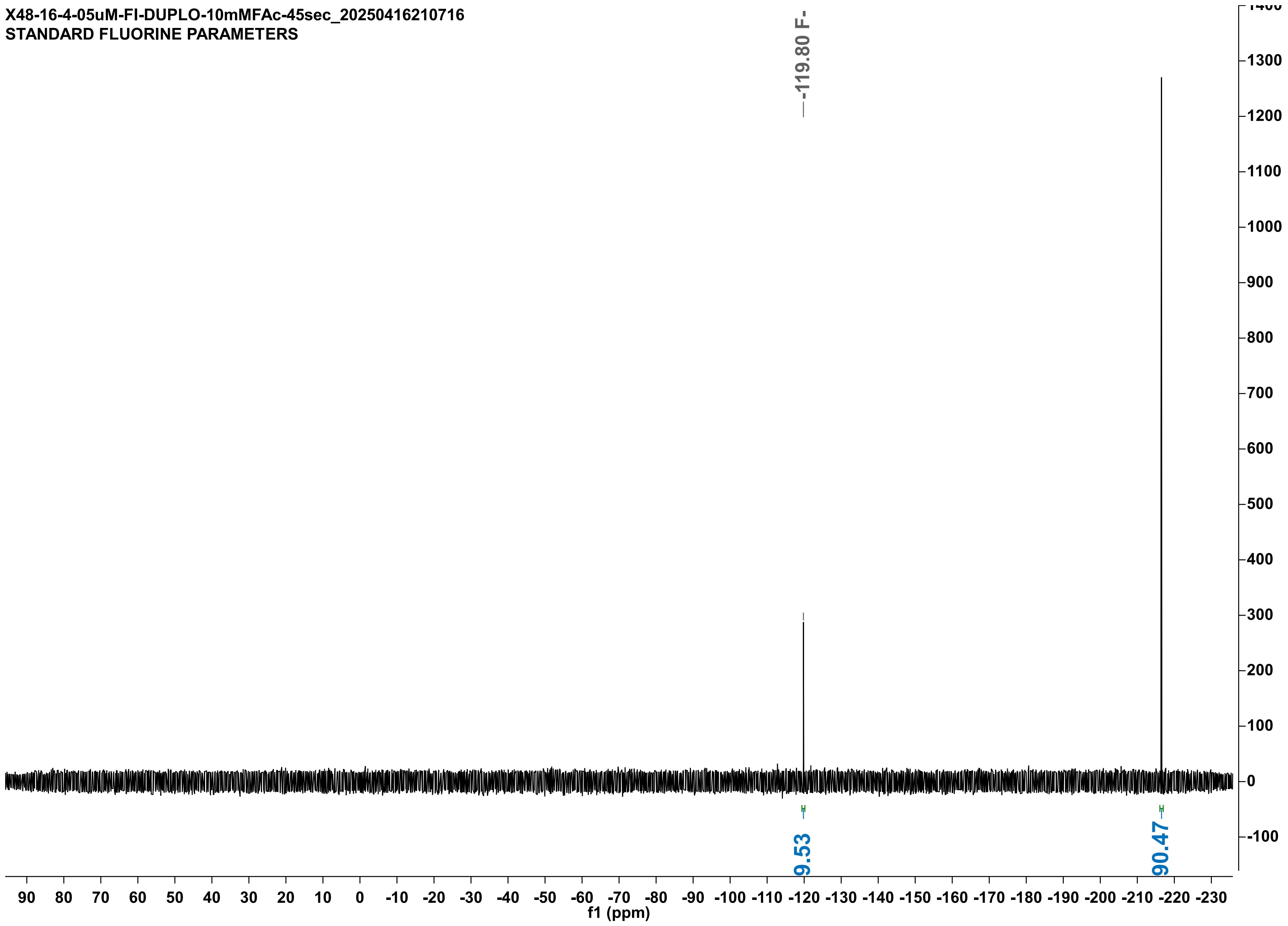

X46-9-4-NOENZ-10mMFPA-20min-noIS_20250409154323
STANDARD FLUORINE PARAMETERS

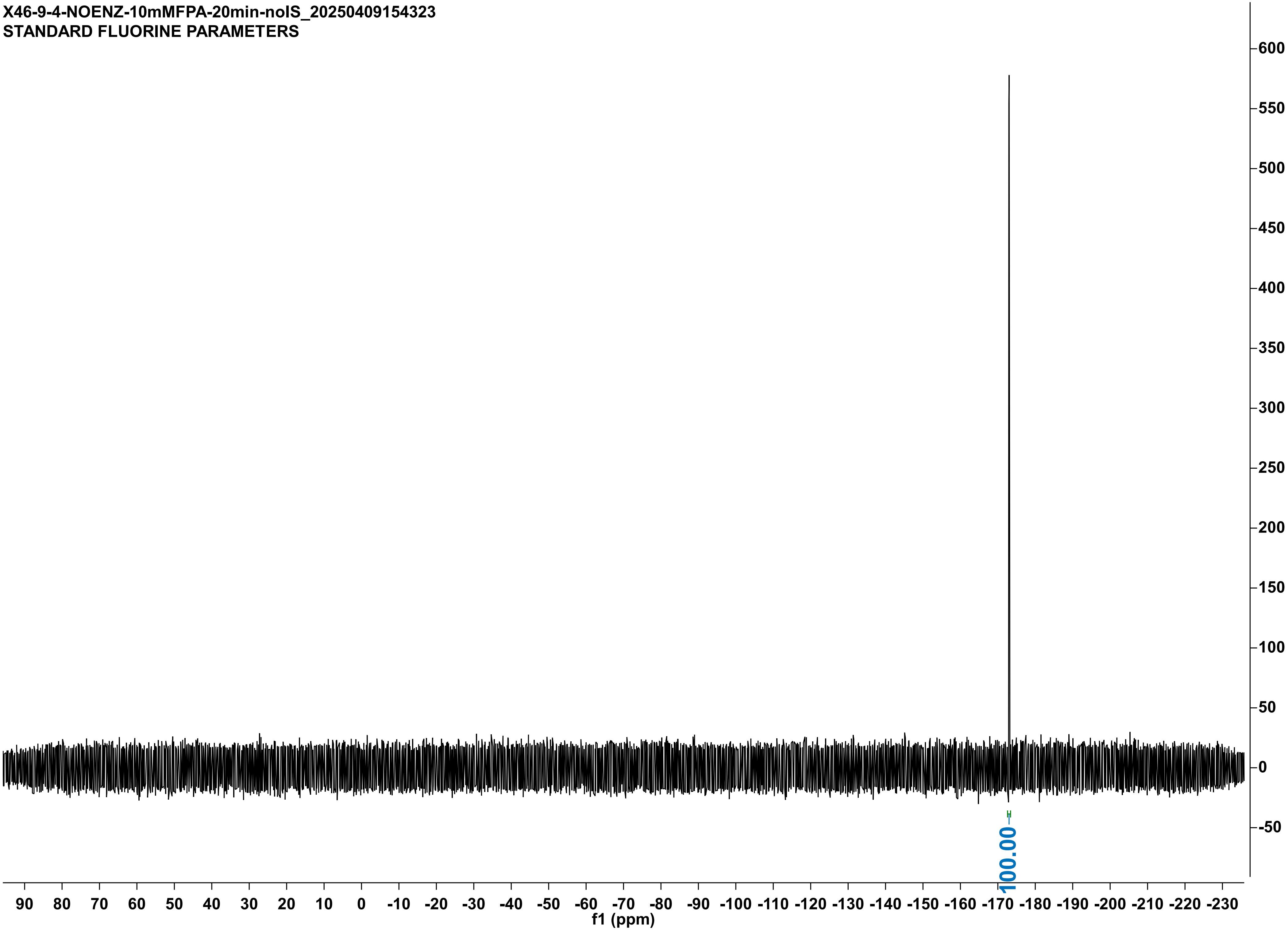

X46-9-4-05uMH272A-2-10mMFPA-20min-noIS_20250409154342
STANDARD FLUORINE PARAMETERS

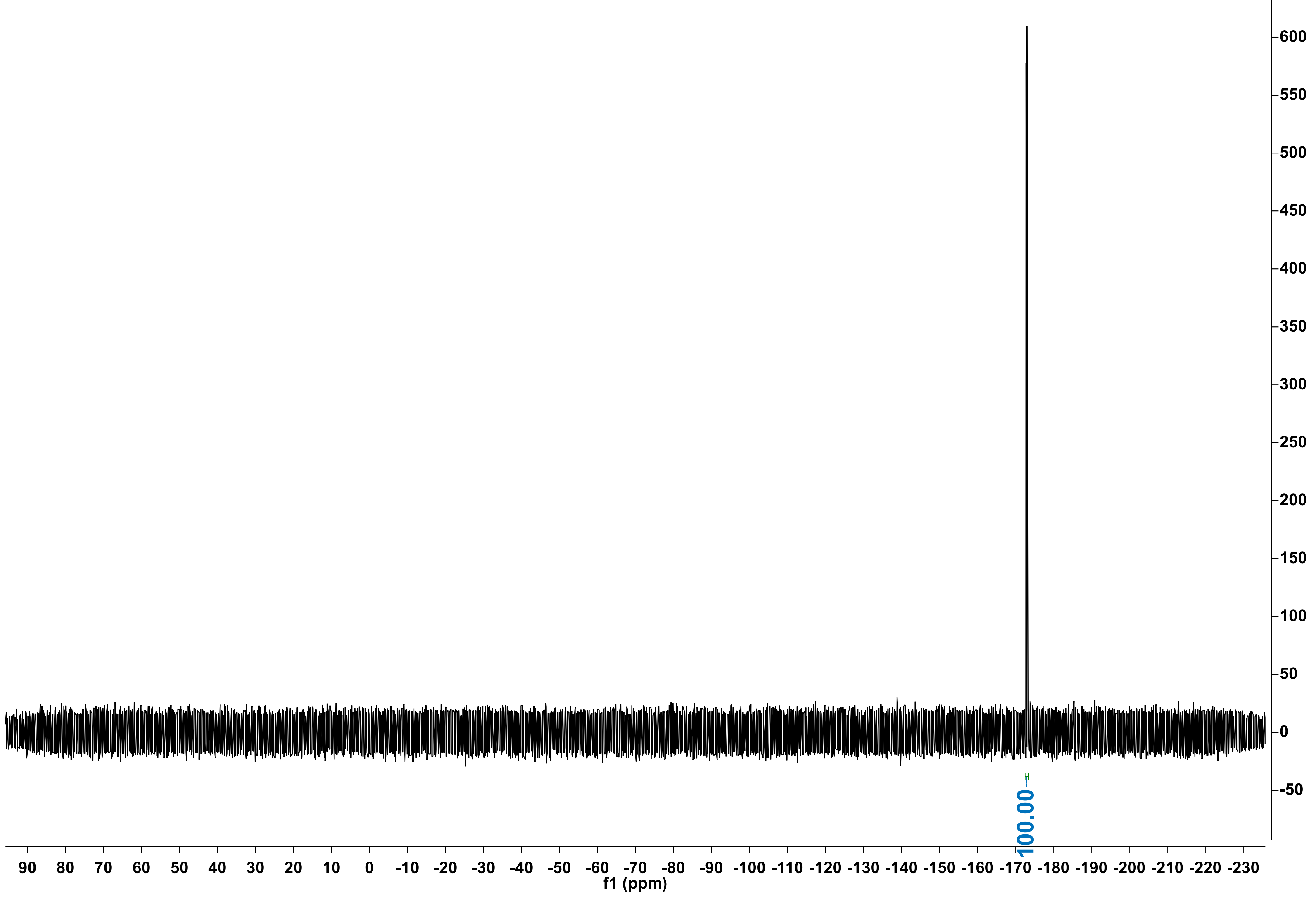

X48-11-4-05uMWT1-10mMFPA-10min_20250411170910
STANDARD FLUORINE PARAMETERS

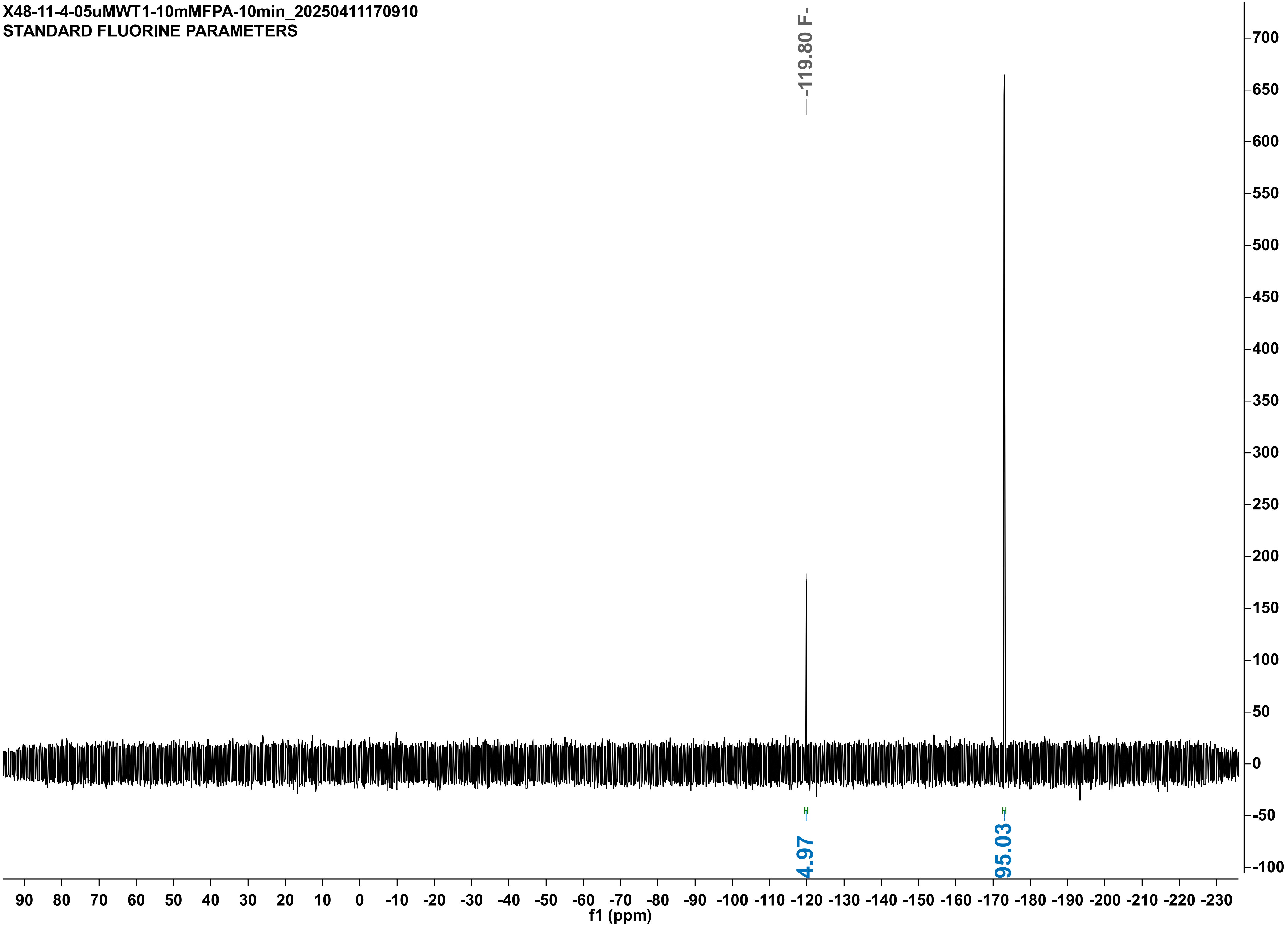

X48-11-4-05uMWT2-10mMFPA-10min_20250411170920
STANDARD FLUORINE PARAMETERS

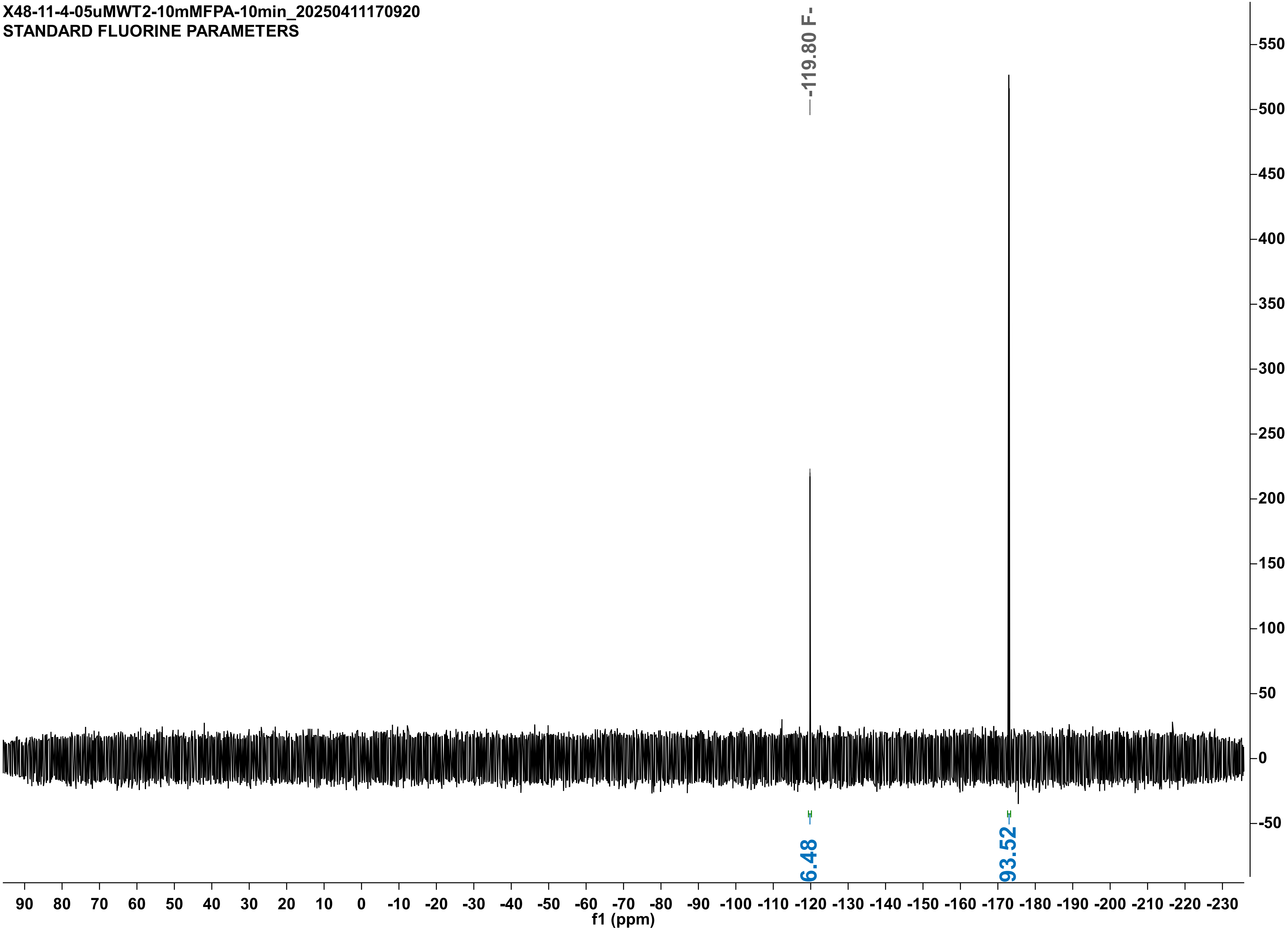

X48-15-4-5uMPLM2-10mMFPA-40min_20250415144111
STANDARD FLUORINE PARAMETERS

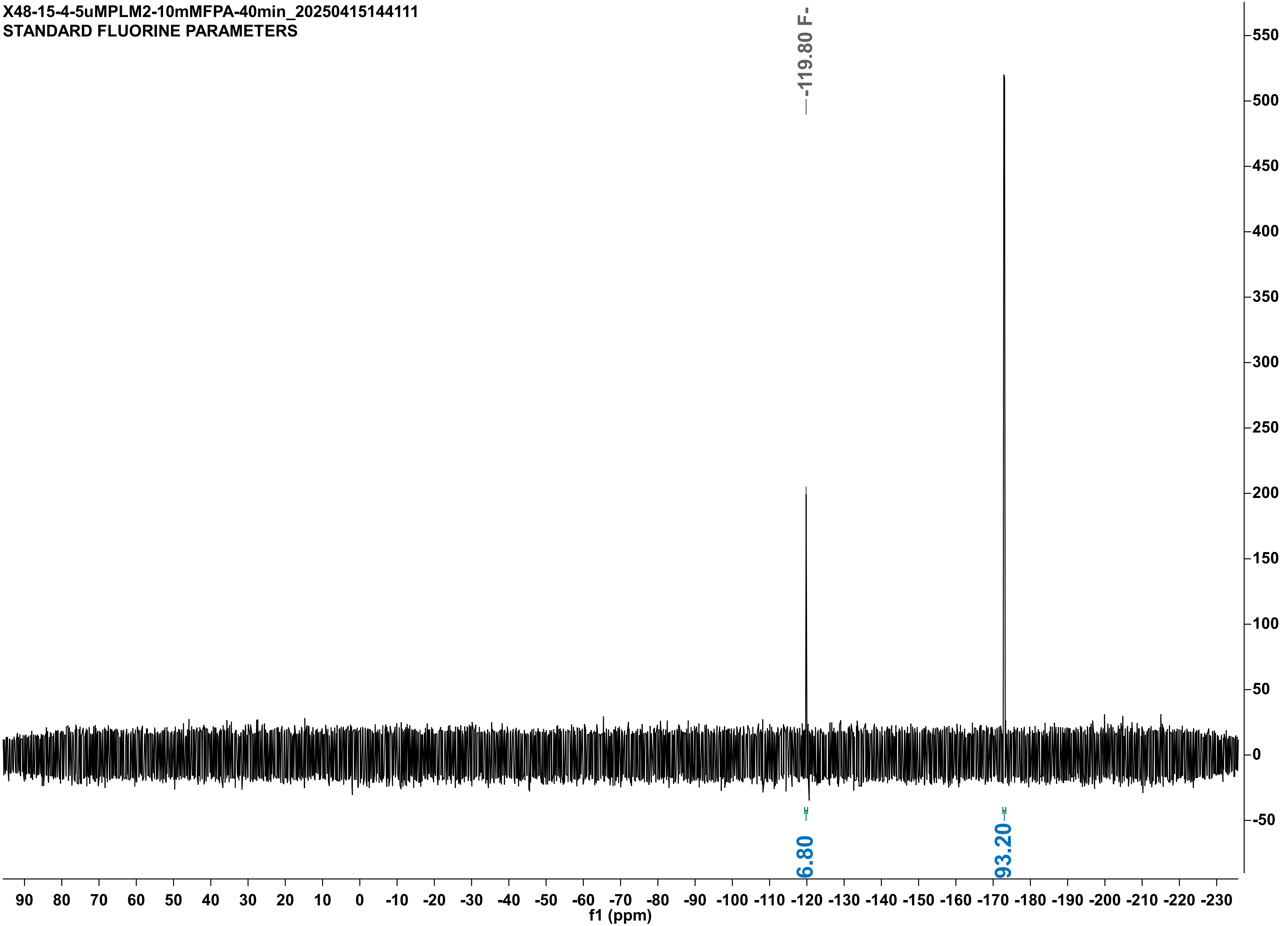

**X48-15-4-5uMPLM2-duplo-10mMFPA-40min_20250415154332**
**STANDARD FLUORINE PARAMETERS**

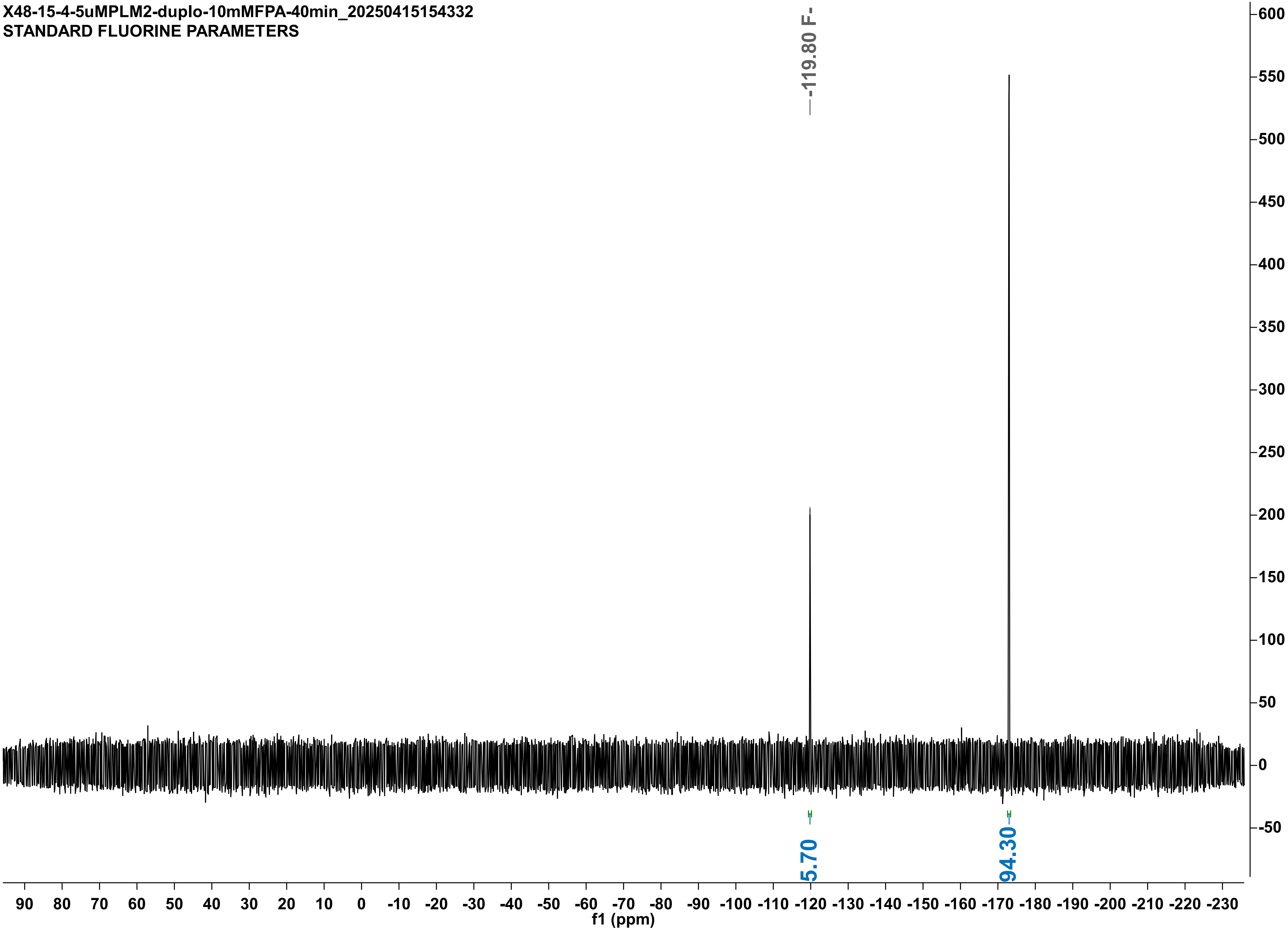

X48-15-4-5uMRPL-10mMFPA-2h30min_20250415151629
STANDARD FLUORINE PARAMETERS

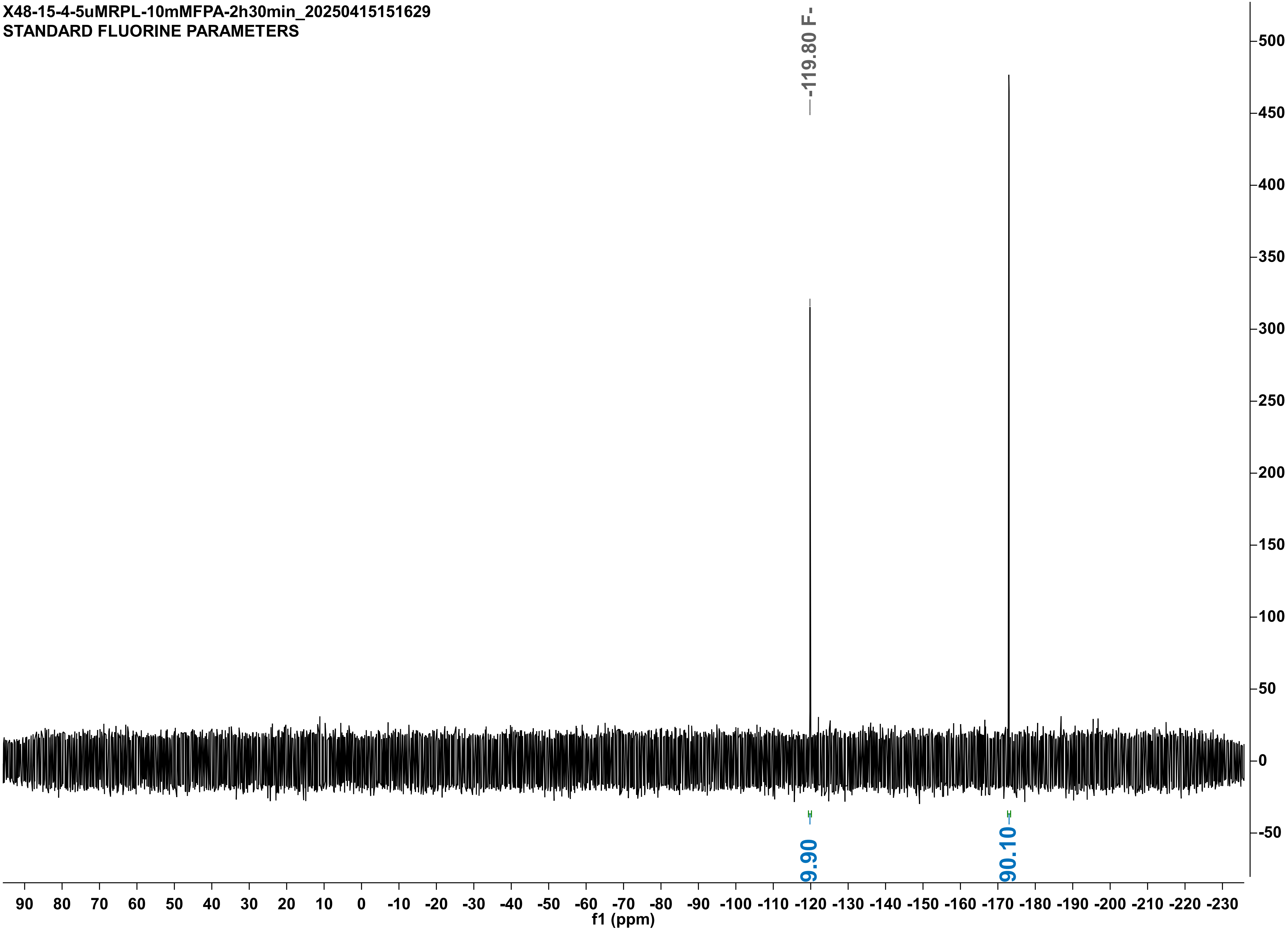

X48-15-4-5uMRPL-DUPLO-10mMFPA-2h30min_20250415174344
STANDARD FLUORINE PARAMETERS

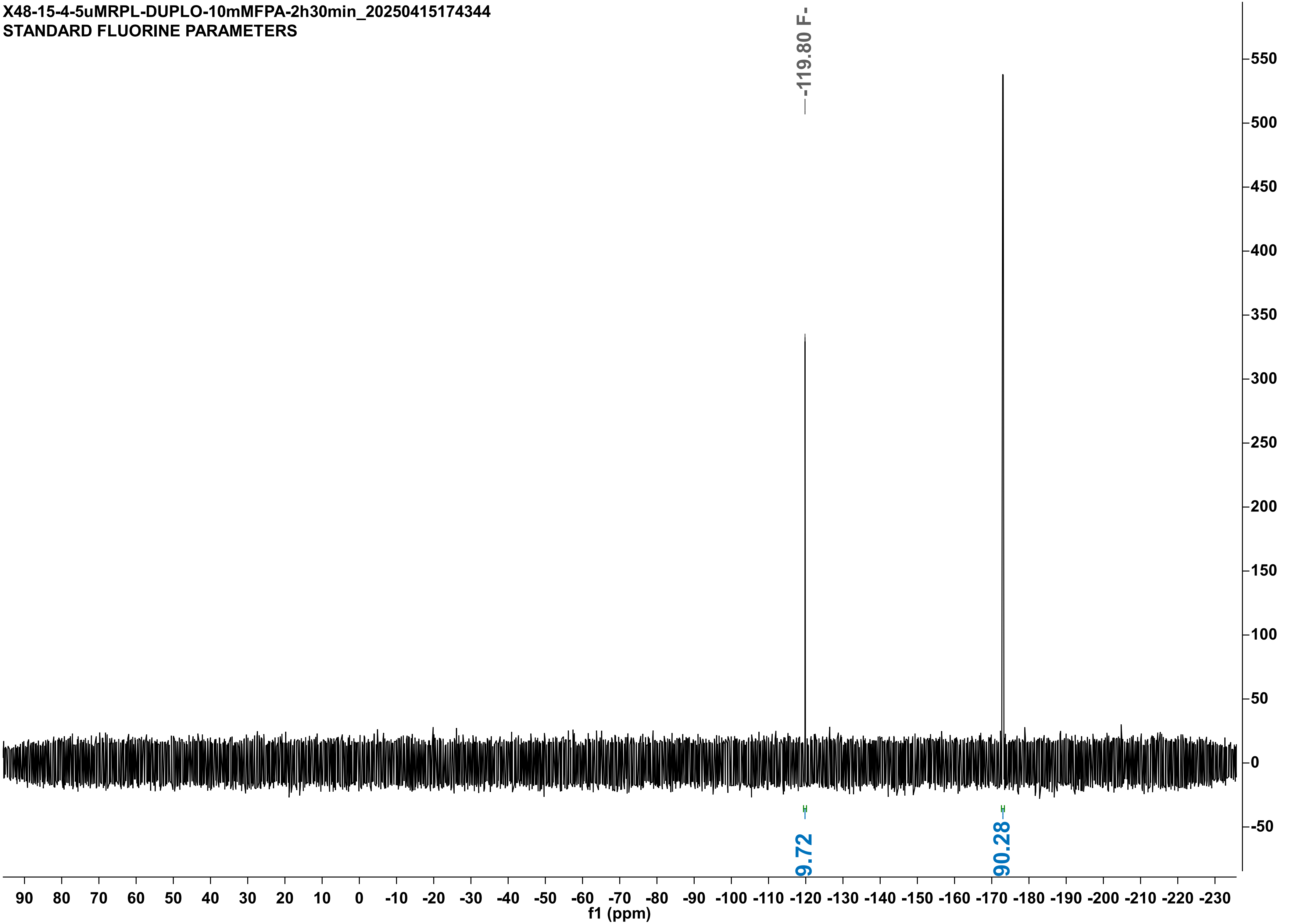

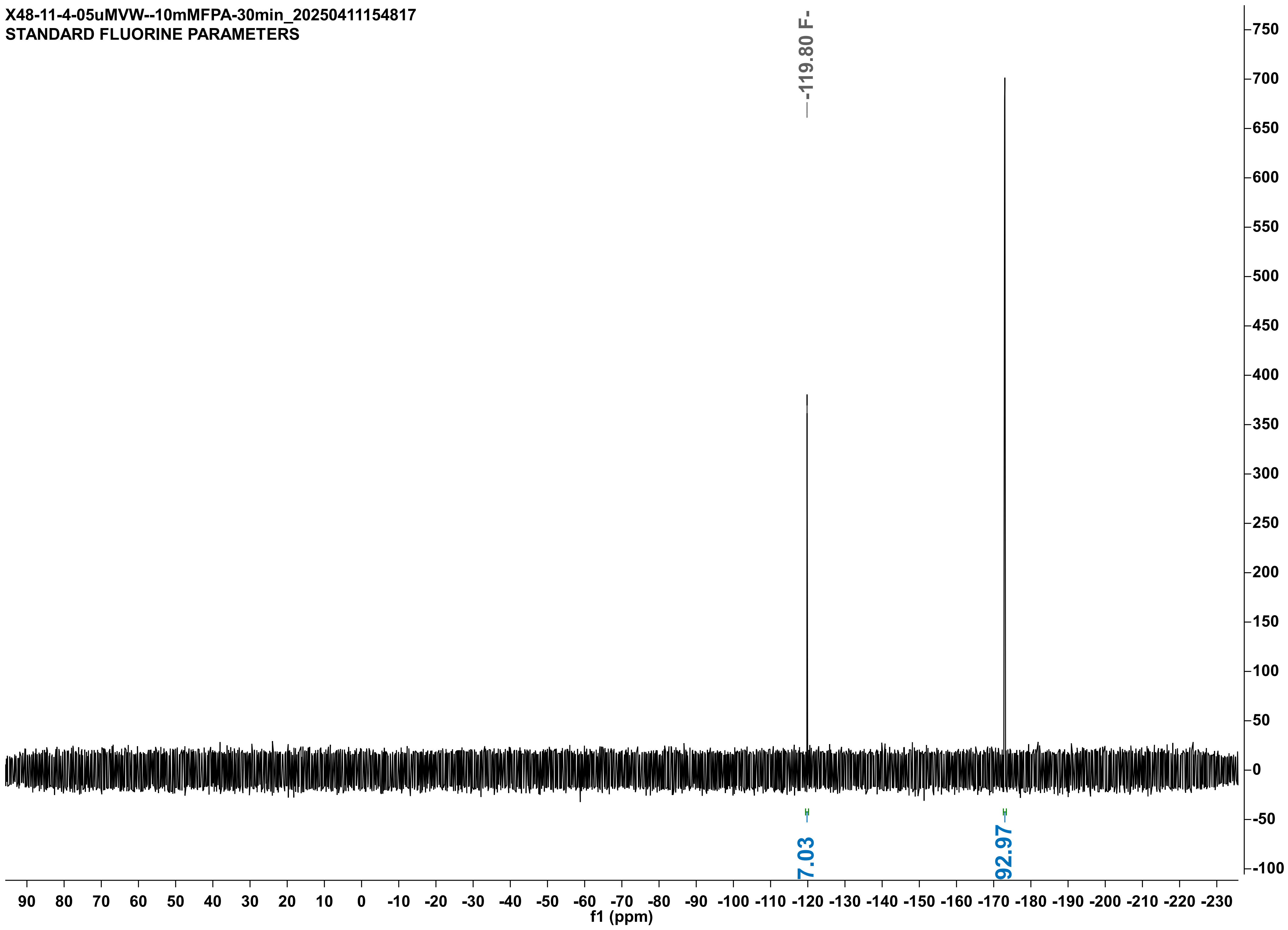

X48-11-4-05uMVW--10mMFPA-30min_20250411154817
STANDARD FLUORINE PARAMETERS
-119.80 F-
7.03
92.97
f1 (ppm)

X48-11-4-05uMVW--DUPLO-10mMFPA-30min_20250411170727
STANDARD FLUORINE PARAMETERS

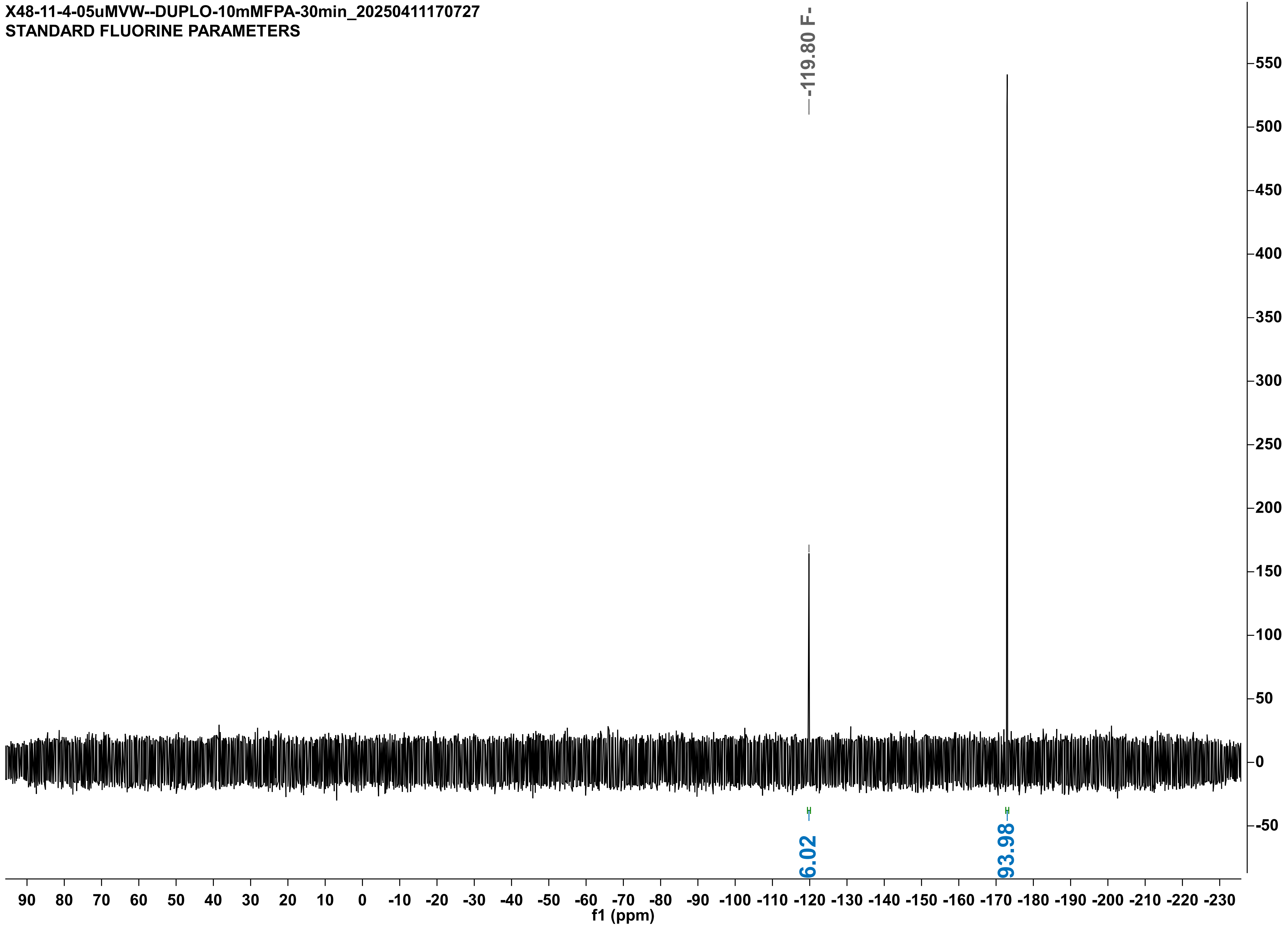

X48-16-4-05uM-VM-DUP-10mMFPA-15min_20250416184333
STANDARD FLUORINE PARAMETERS

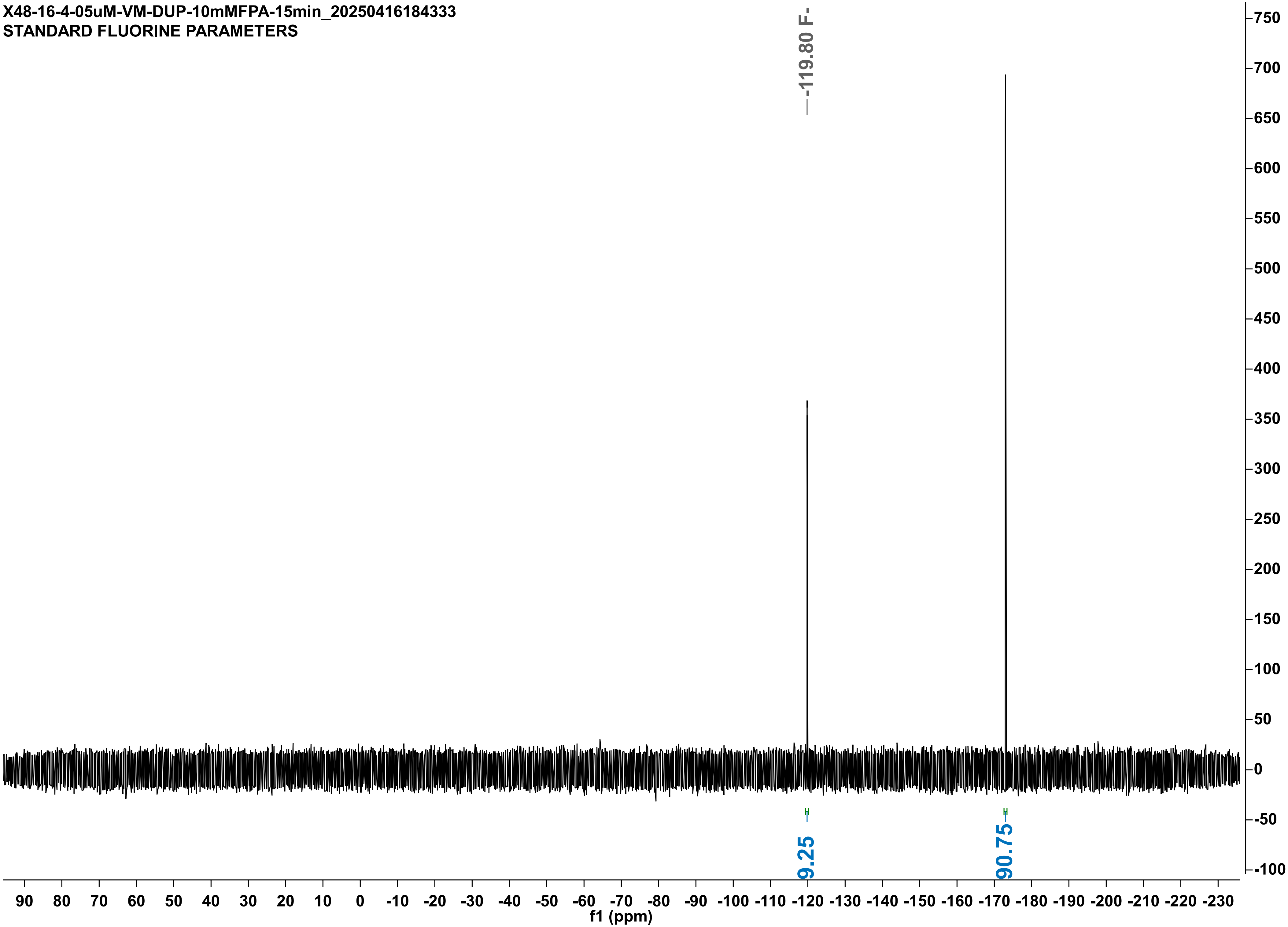

X48-16-4-05uM-VM-10mMFPA-15min_20250416184332
STANDARD FLUORINE PARAMETERS

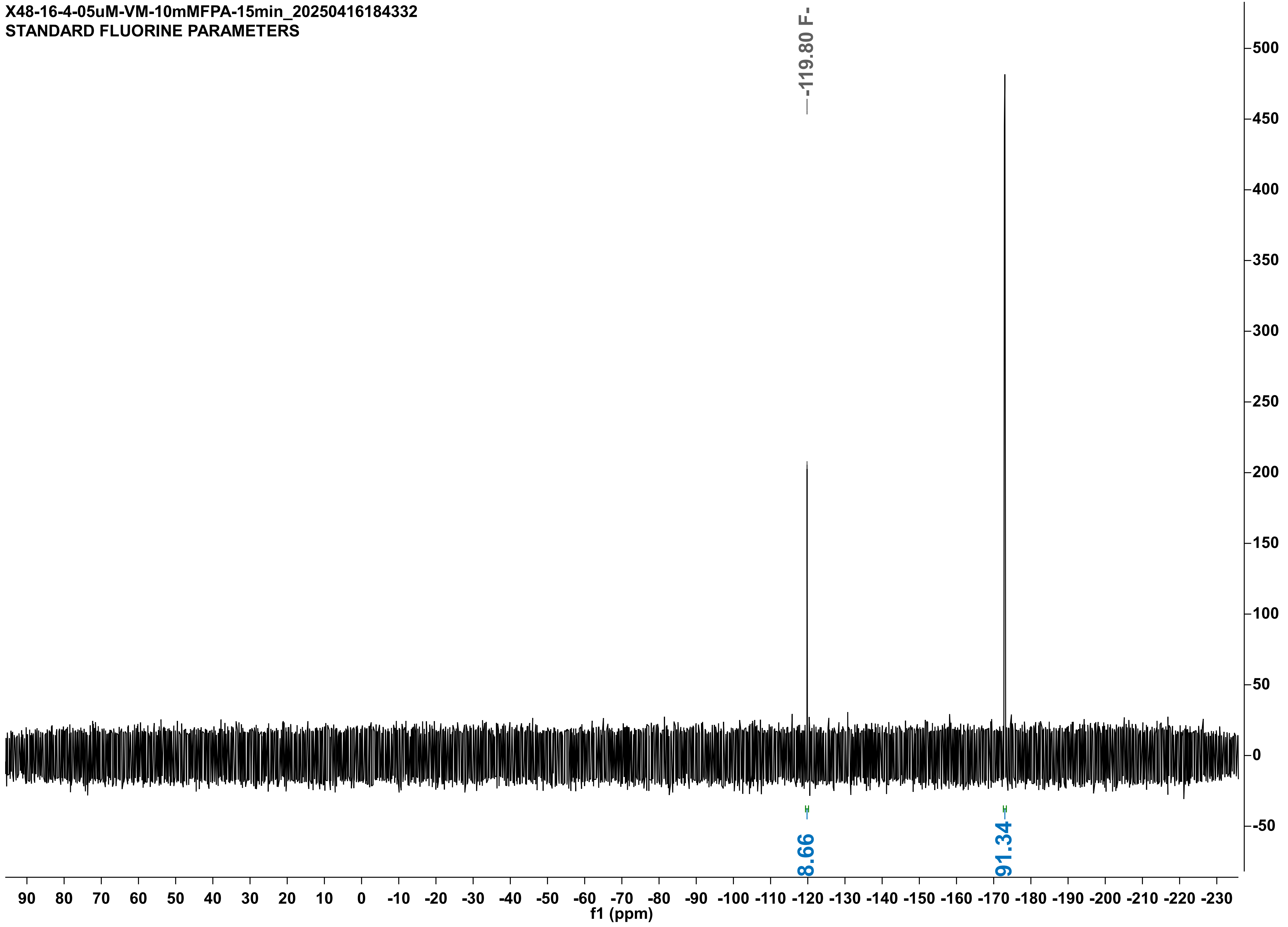

X48-15-4-05uMLW--10mMFPA-30min_20250415181748
STANDARD FLUORINE PARAMETERS

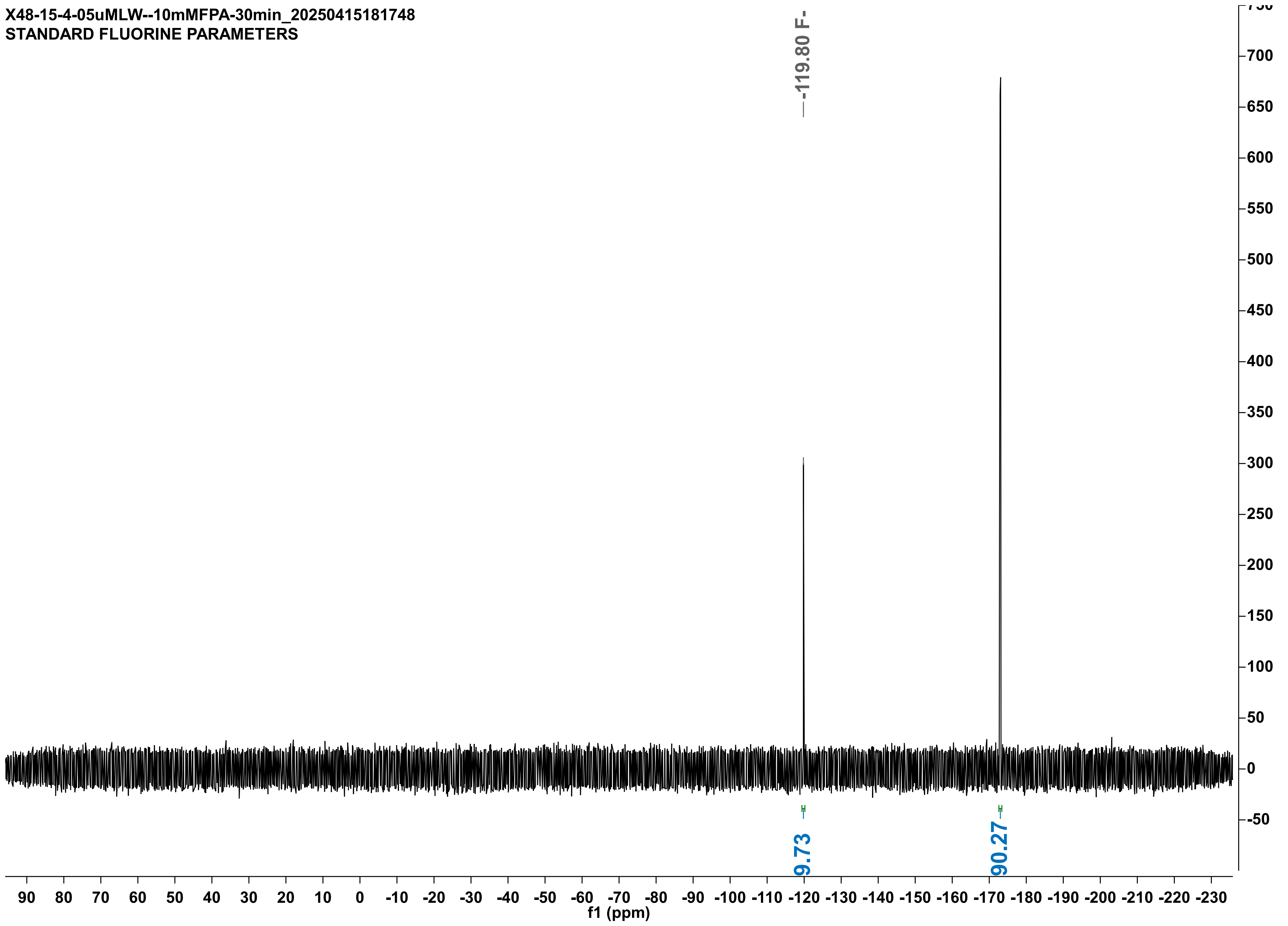

X48-16-4-05uMLW--10mMFPA-30min_20250416202647
STANDARD FLUORINE PARAMETERS

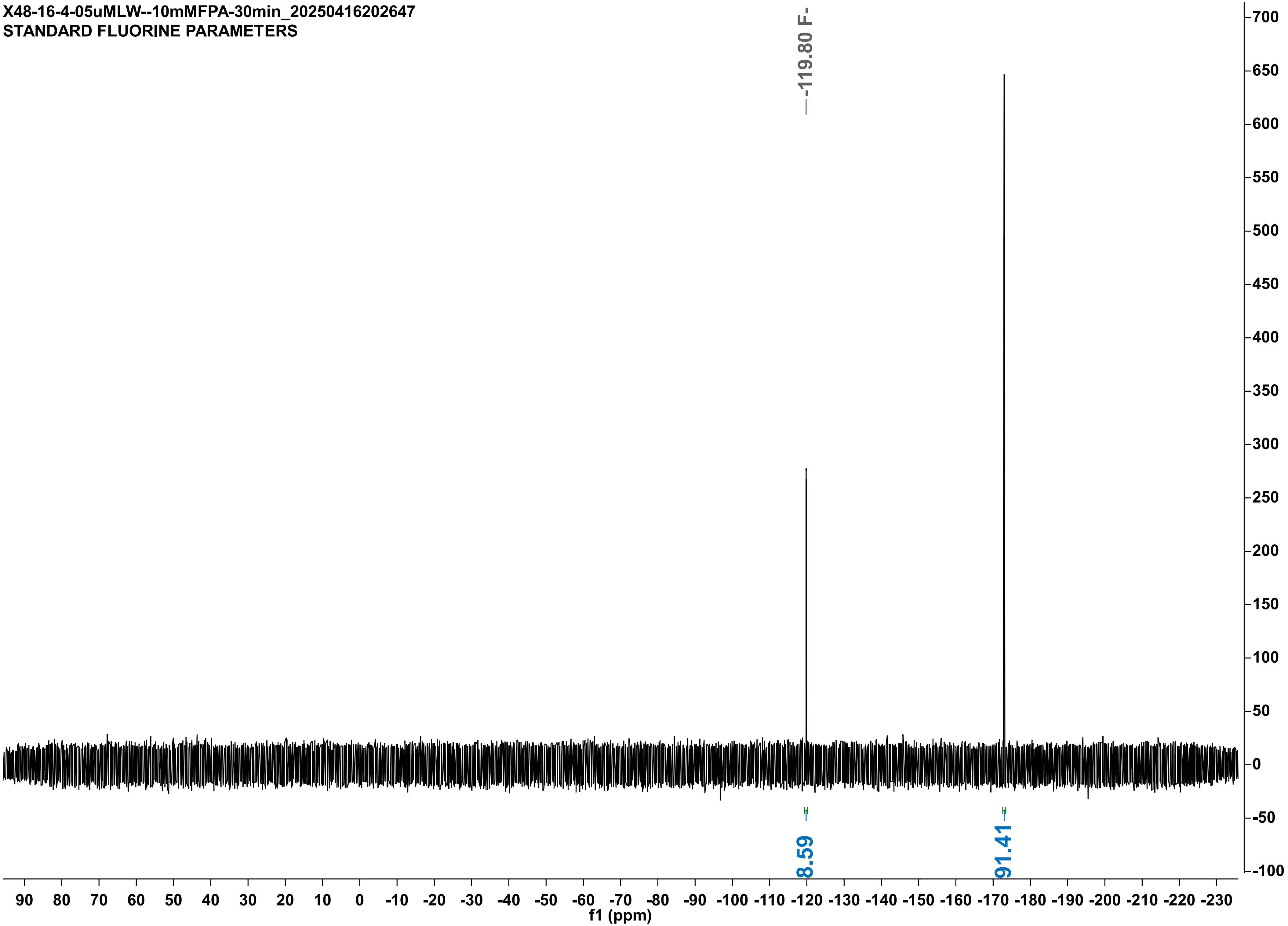

X48-16-4-05uMRLM-10mMFPA-25min_20250416203025
STANDARD FLUORINE PARAMETERS

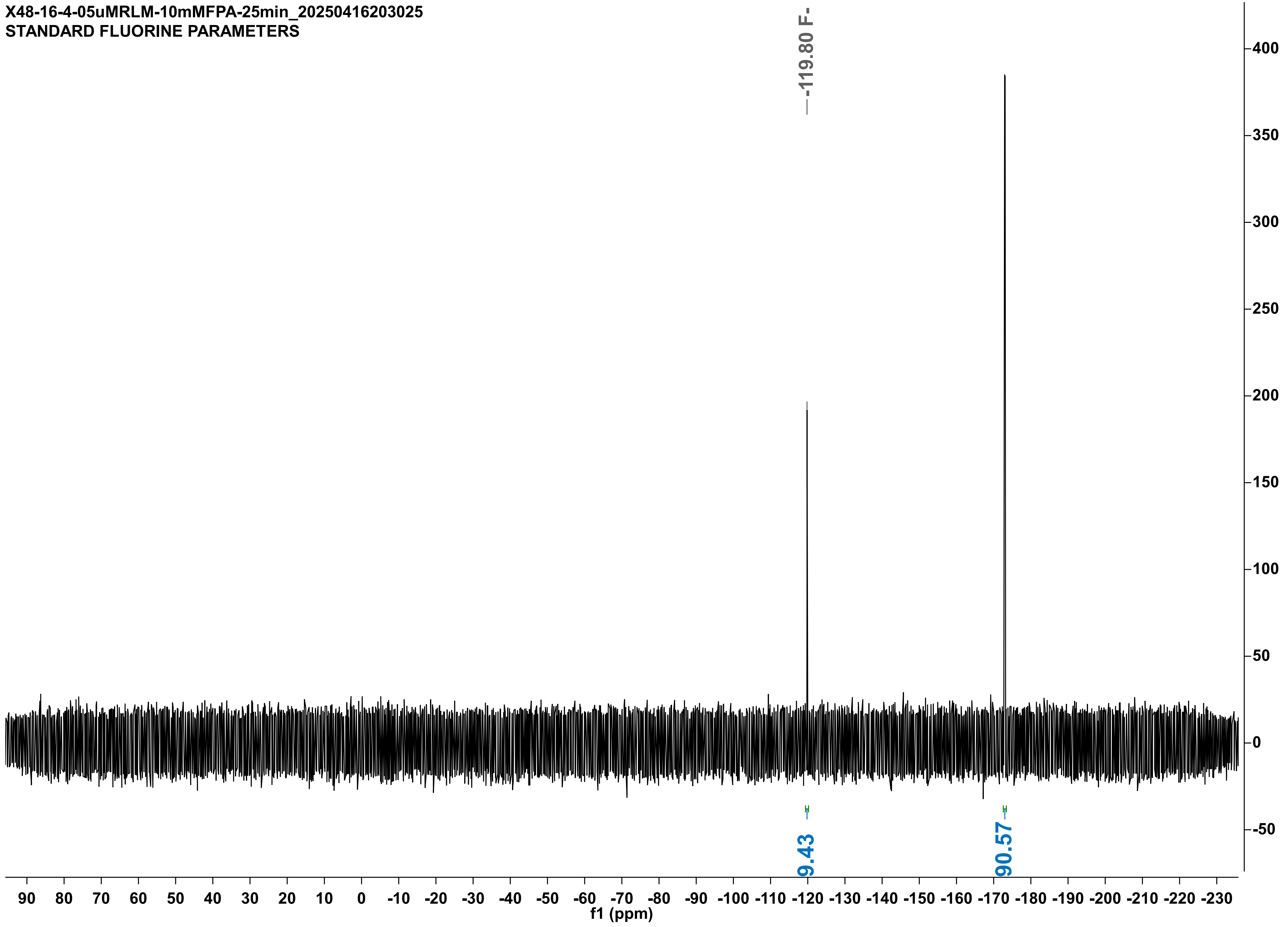

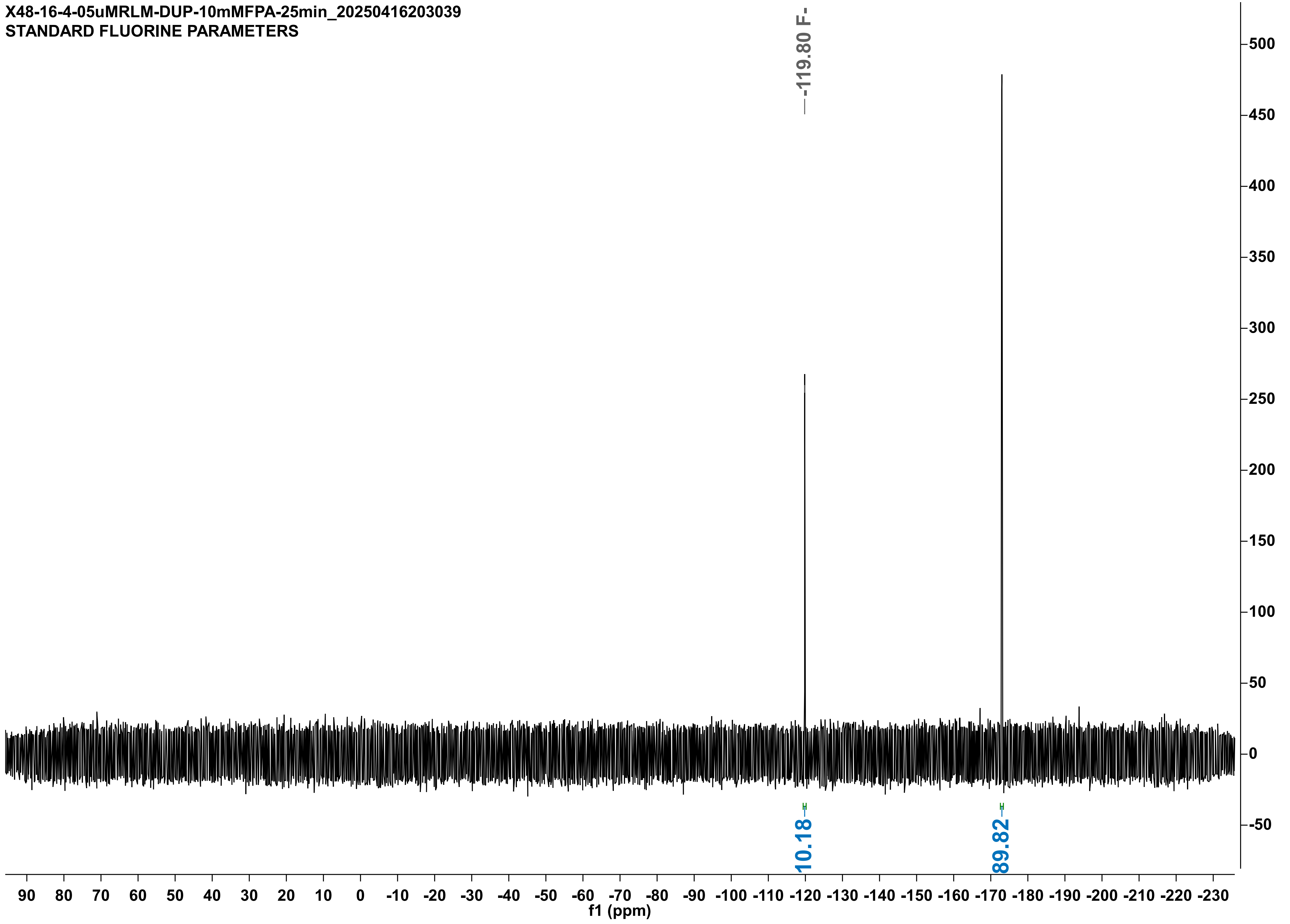

X48-16-4-05uMRLM-DUP-10mMFPA-25min_20250416203039
STANDARD FLUORINE PARAMETERS
-119.80 F-
10.18
89.82
f1 (ppm)

X48-15-4-5uMRHL-DUPLO-10mMFPA-90min_20250415174520
STANDARD FLUORINE PARAMETERS

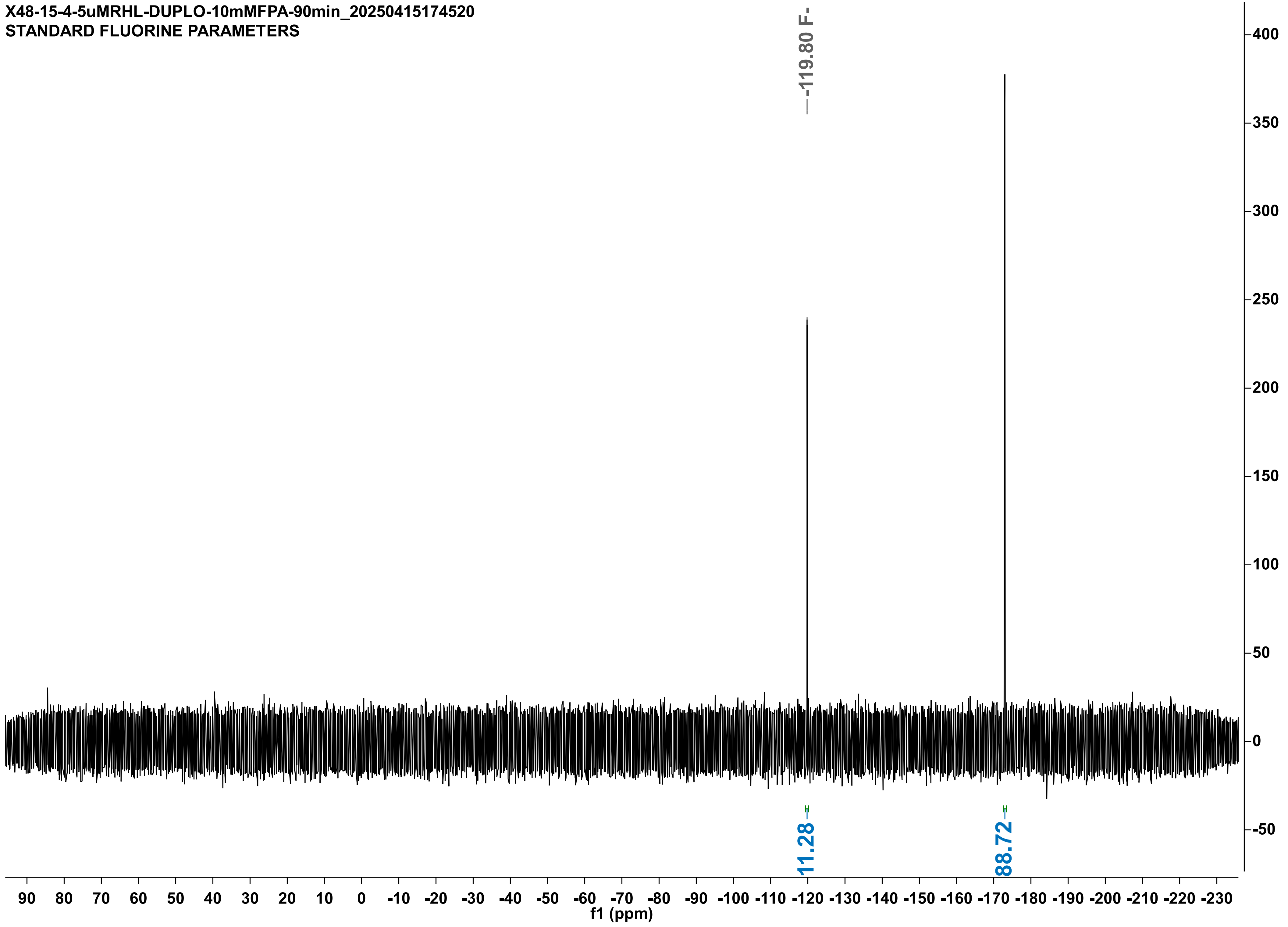

X48-15-4-5uMRHL-10mMFPA-90min_20250415140510
STANDARD FLUORINE PARAMETERS

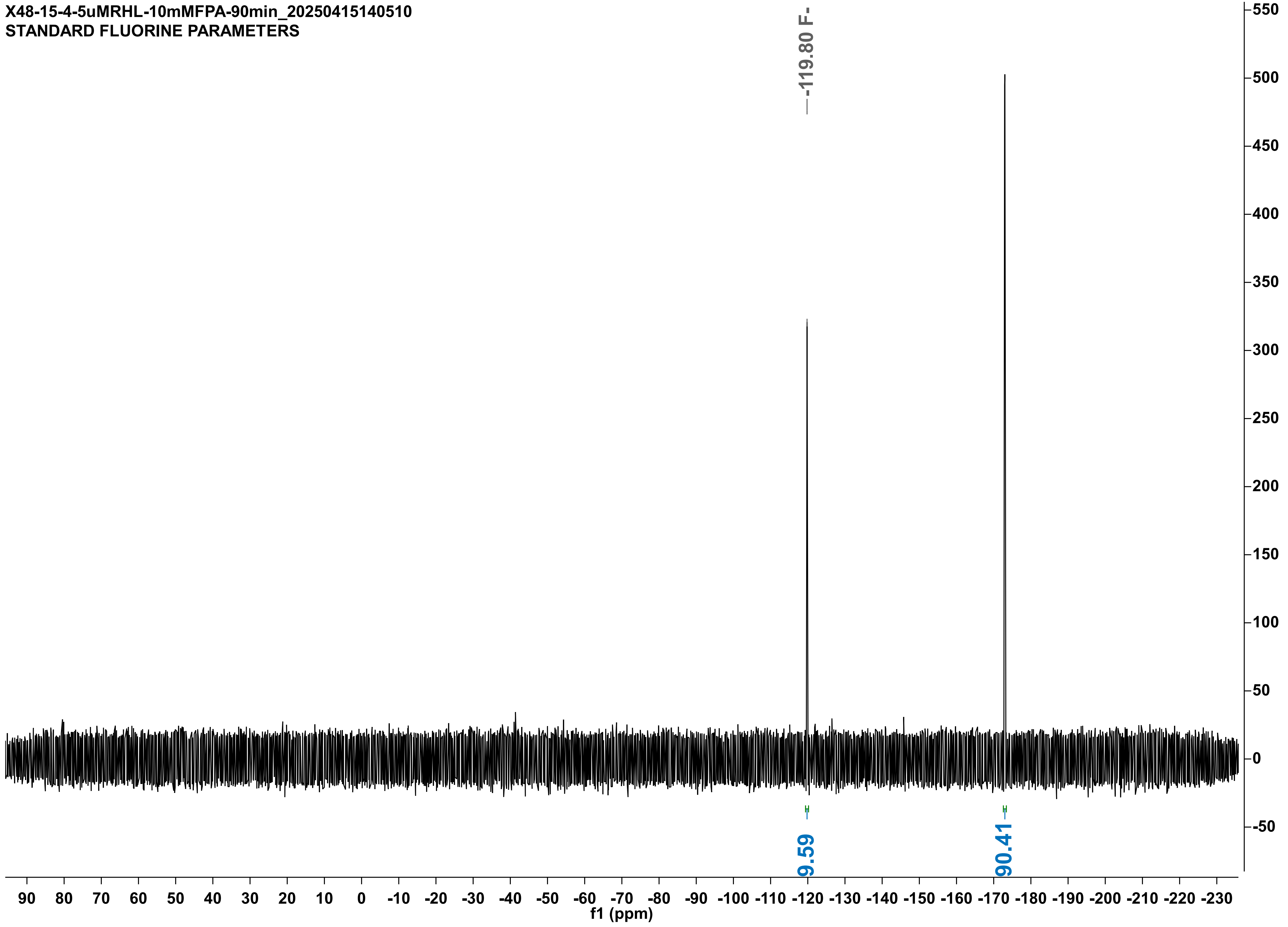

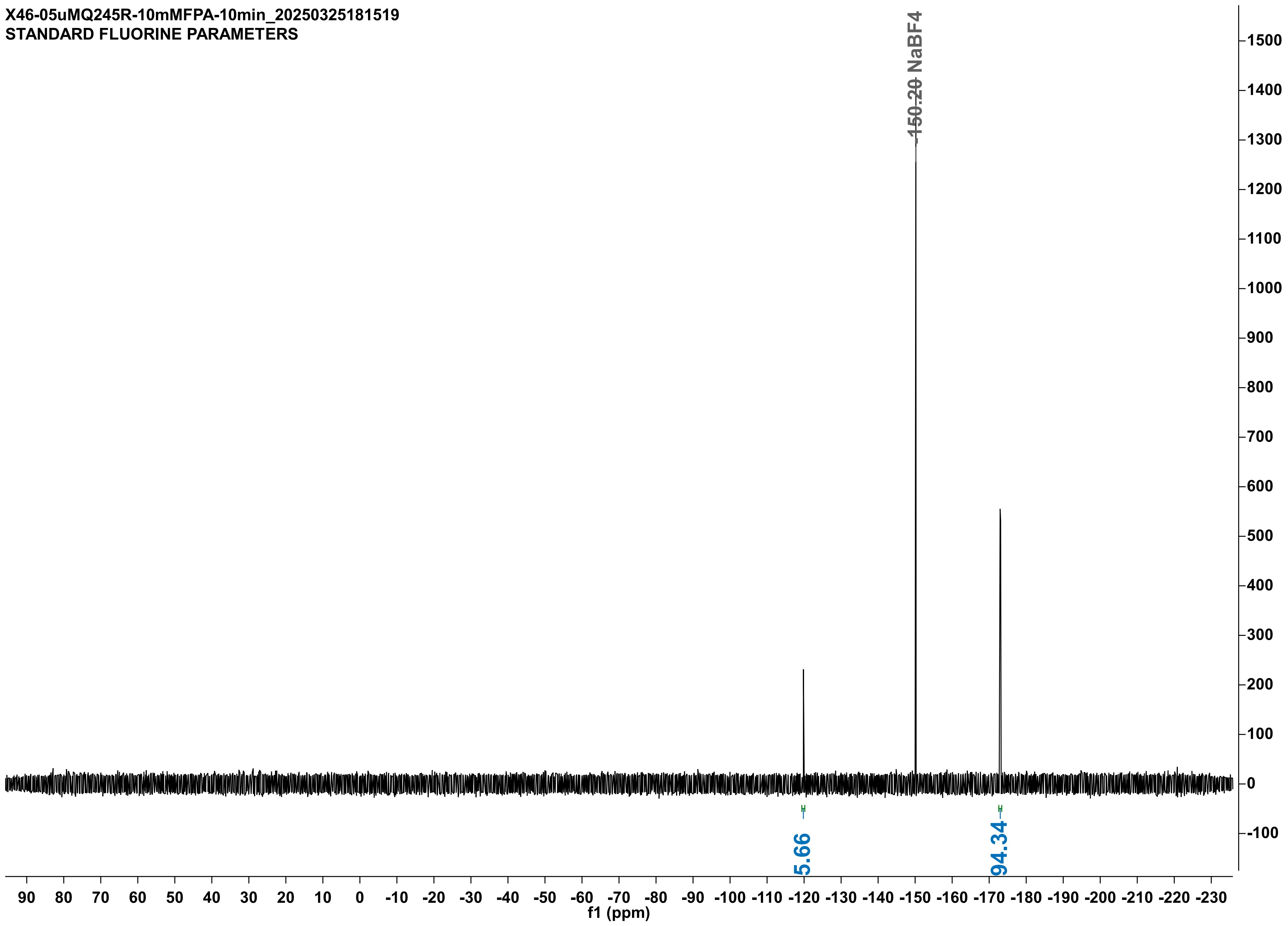
X46-05uMQ245R-10mMFPA-10min_20250325181519
STANDARD FLUORINE PARAMETERS
-150.20 NaBF4
5.66
94.34
f1 (ppm)

X46-9-4-05uMQ245R-DUPLO-10mMFPA-10min-noIS_20250409152234
STANDARD FLUORINE PARAMETERS

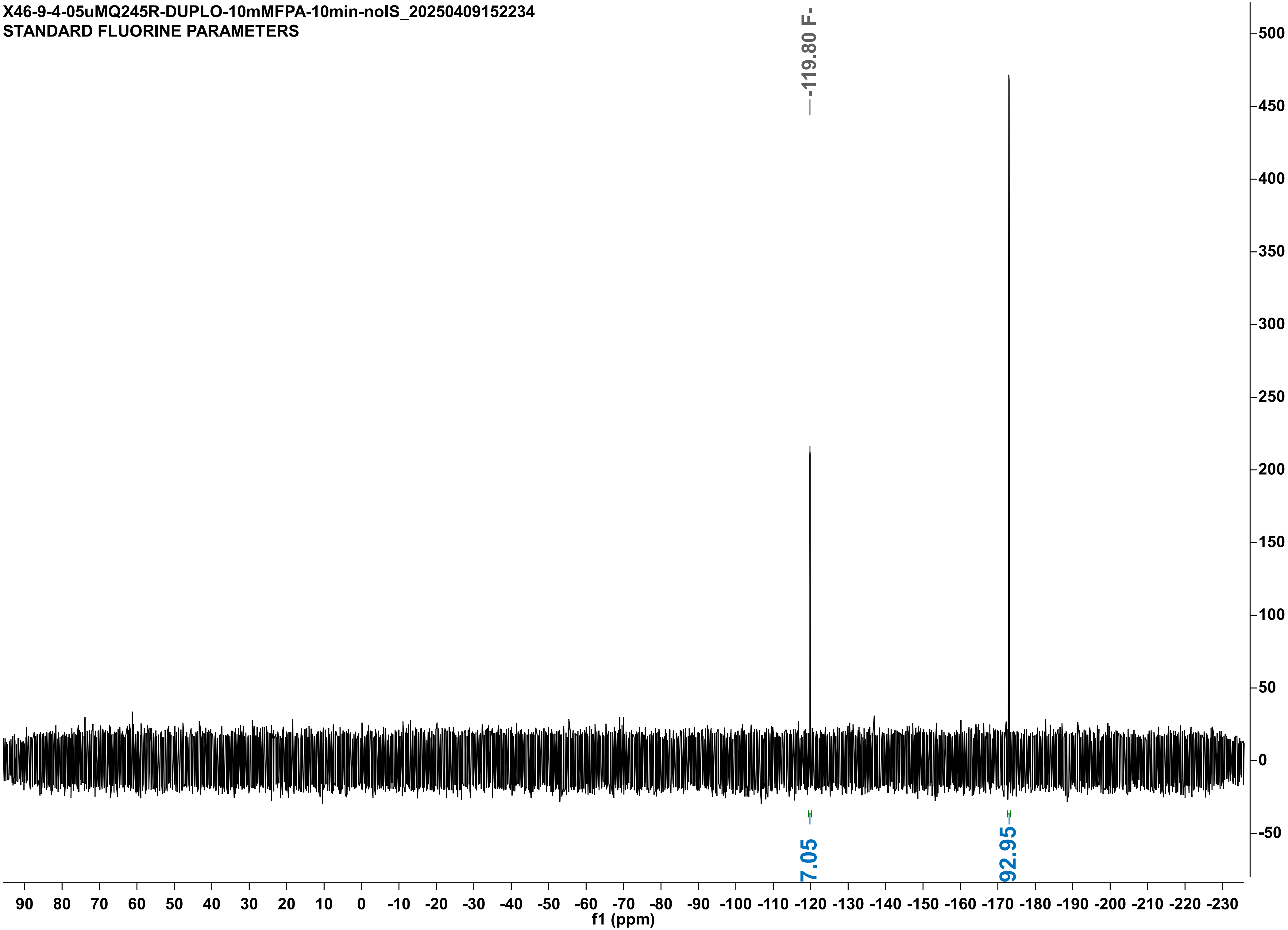

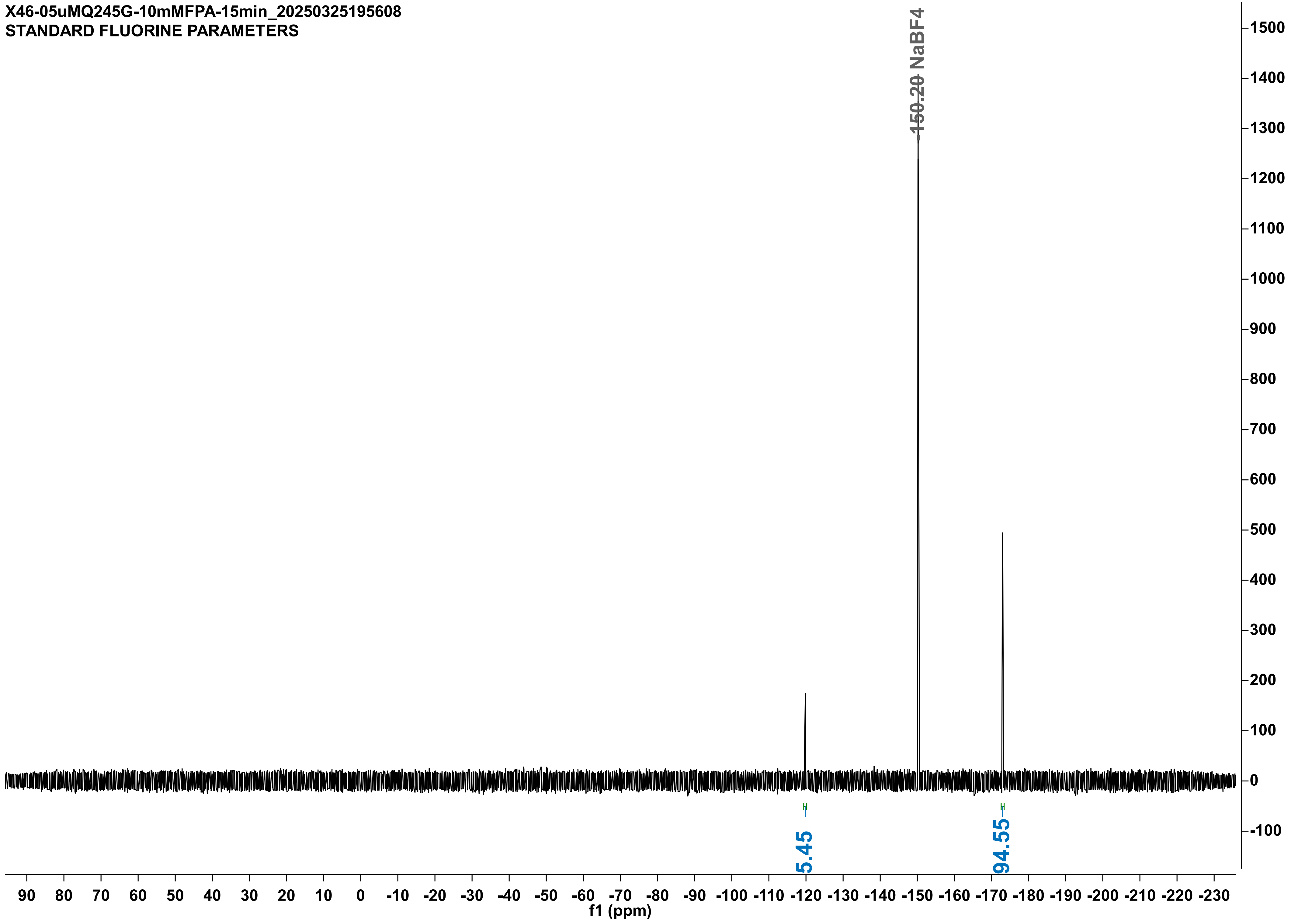
X46-05uMQ245G-10mMFPA-15min_20250325195608
STANDARD FLUORINE PARAMETERS
-150.20 NaBF4
5.45
94.55
f1 (ppm)

X46-9-4-05uMQ245G-DUPLO-10mMFPA-15min-noIS_20250409154236
STANDARD FLUORINE PARAMETERS

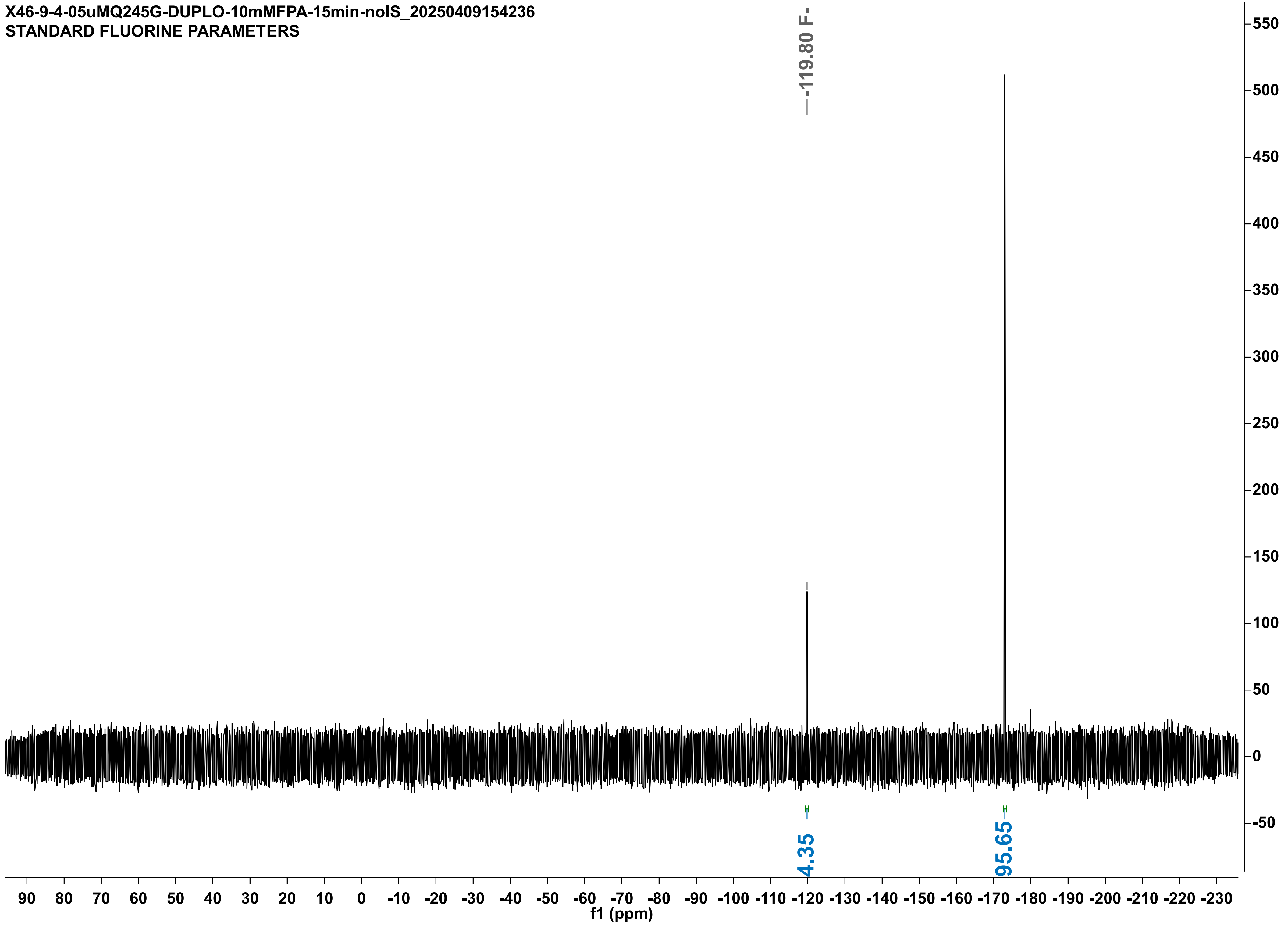

X48-11-4-05uMQ245A-10mMFPA-10min_20250411132706
STANDARD FLUORINE PARAMETERS

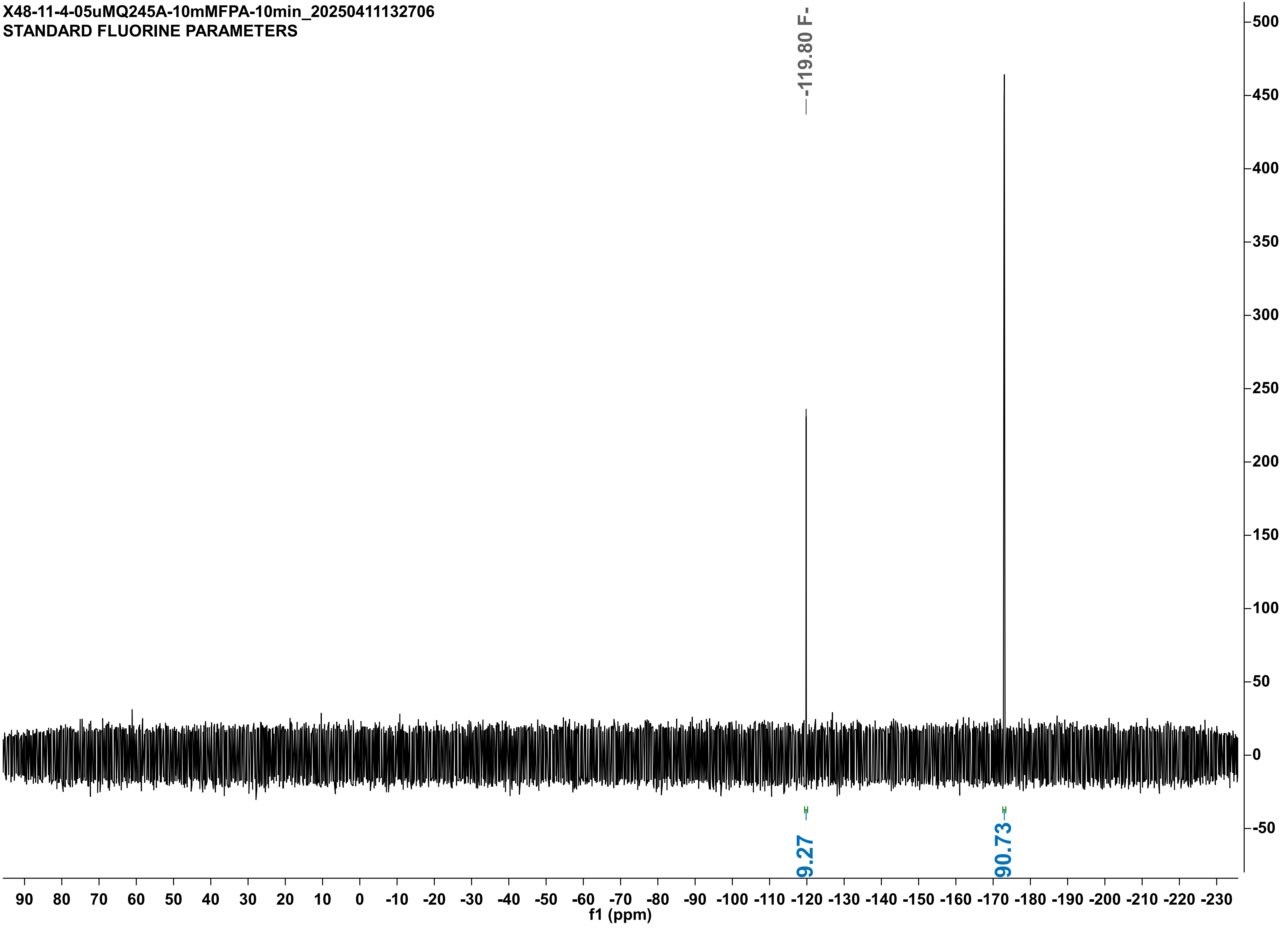

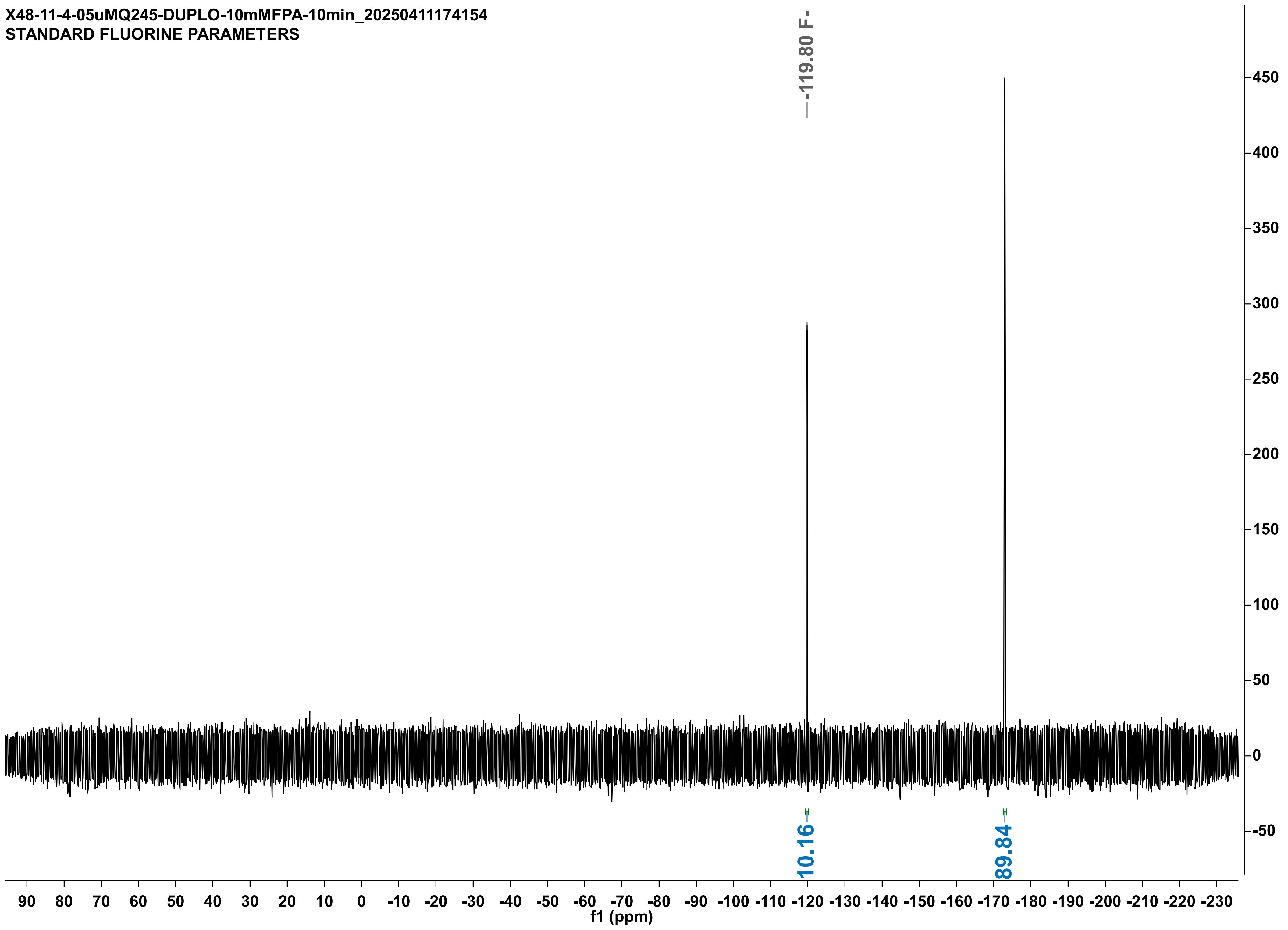

X48-11-4-05uMQ245-DUPLO-10mMFPA-10min_20250411174154
STANDARD FLUORINE PARAMETERS
-119.80 F-
10.16
89.84
450
400
350
300
250
200
150
100
50
0
-50
90 80 70 60 50 40 30 20 10 0 -10 -20 -30 -40 -50 -60 -70 -80 -90 -100 -110 -120 -130 -140 -150 -160 -170 -180 -190 -200 -210 -220 -230
f1 (ppm)

X48-15-4-5uM-ML-duplo-10mMFPA-40min_20250415155244
STANDARD FLUORINE PARAMETERS

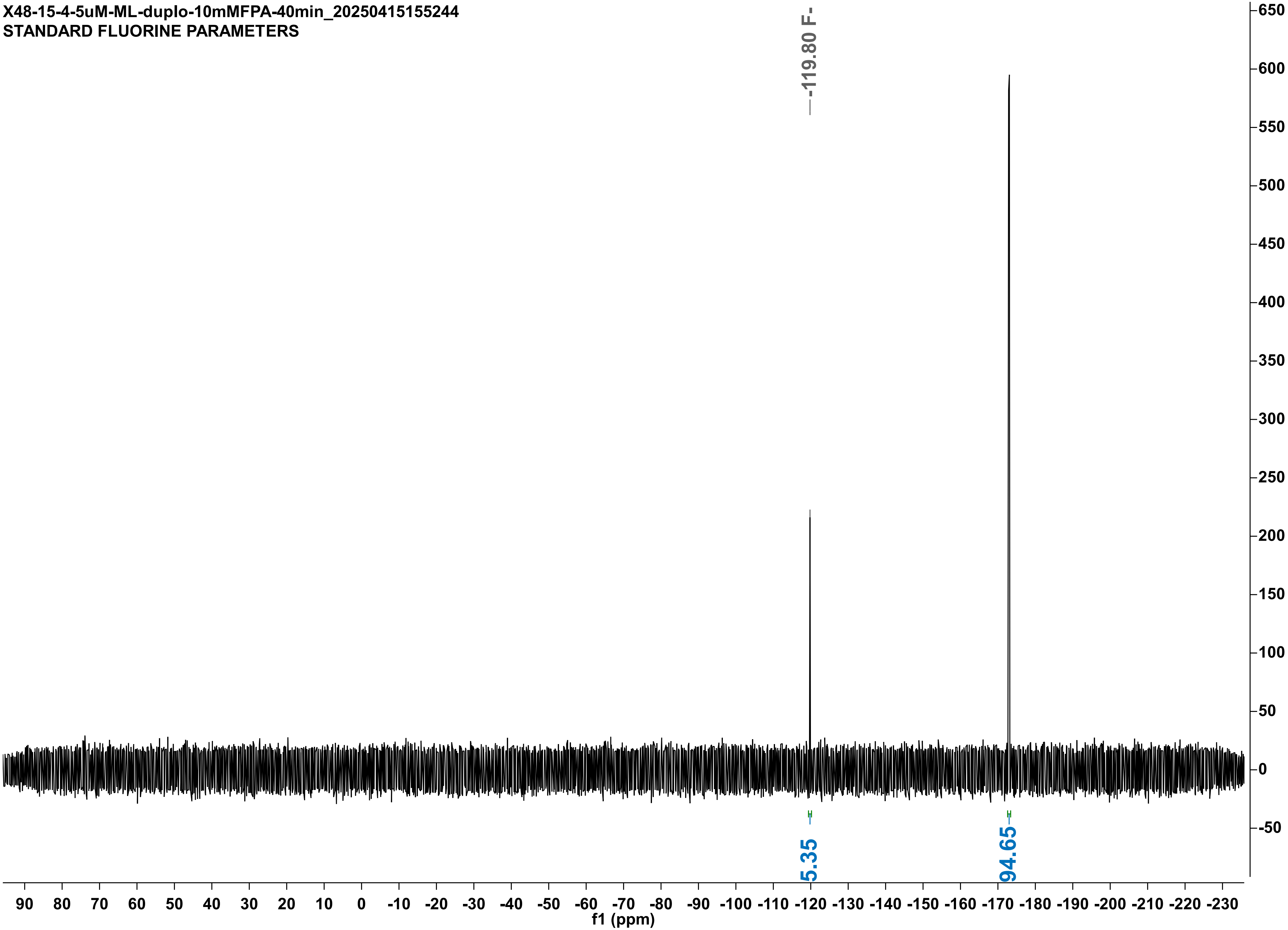

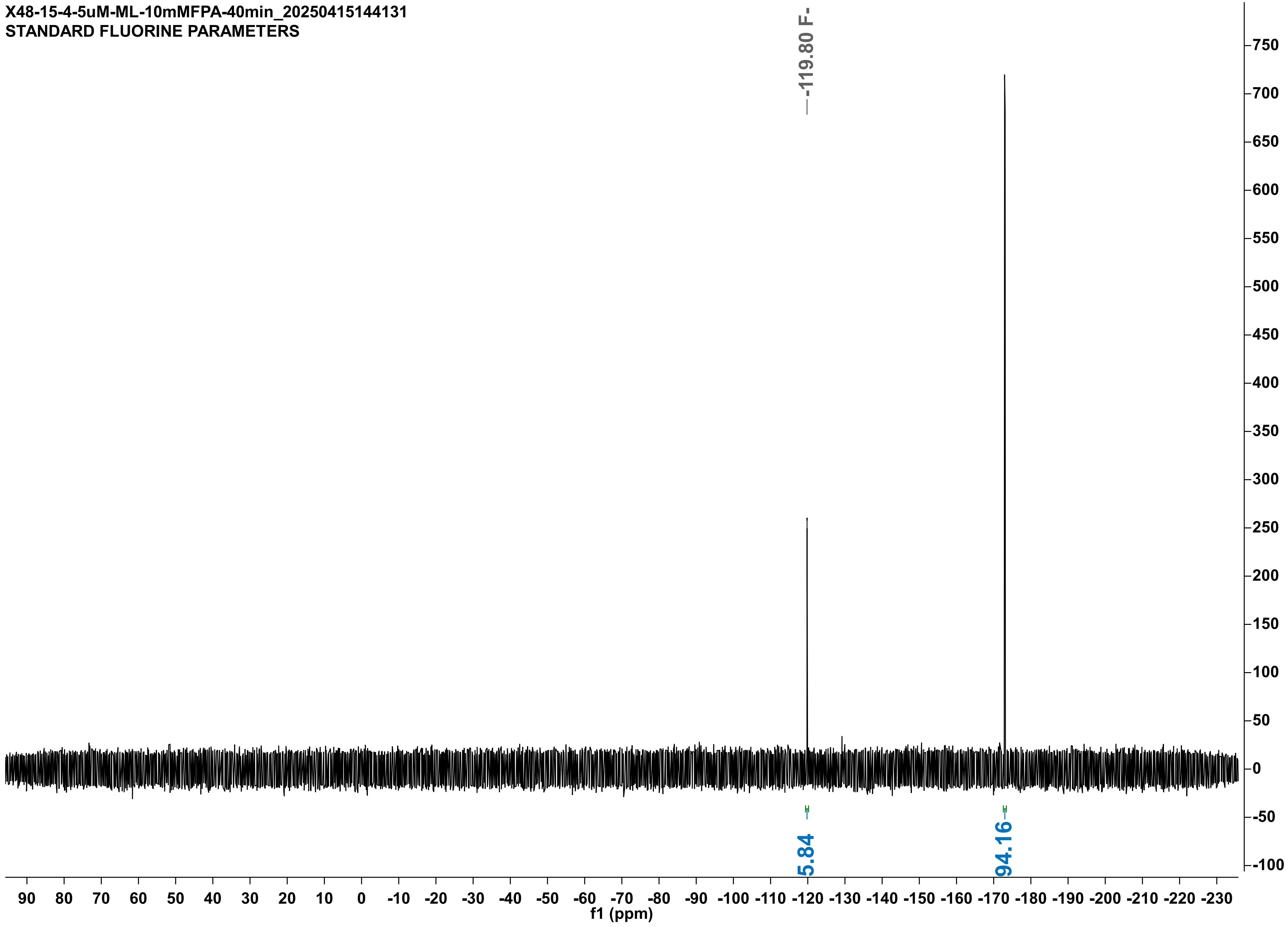

X48-15-4-5uM-ML-10mMFPA-40min_20250415144131
STANDARD FLUORINE PARAMETERS
-119.80 F-
5.84
94.16
f1 (ppm)

X48-16-4-05uMLVM-DUP-10mMFPA-15min_20250416184330
STANDARD FLUORINE PARAMETERS

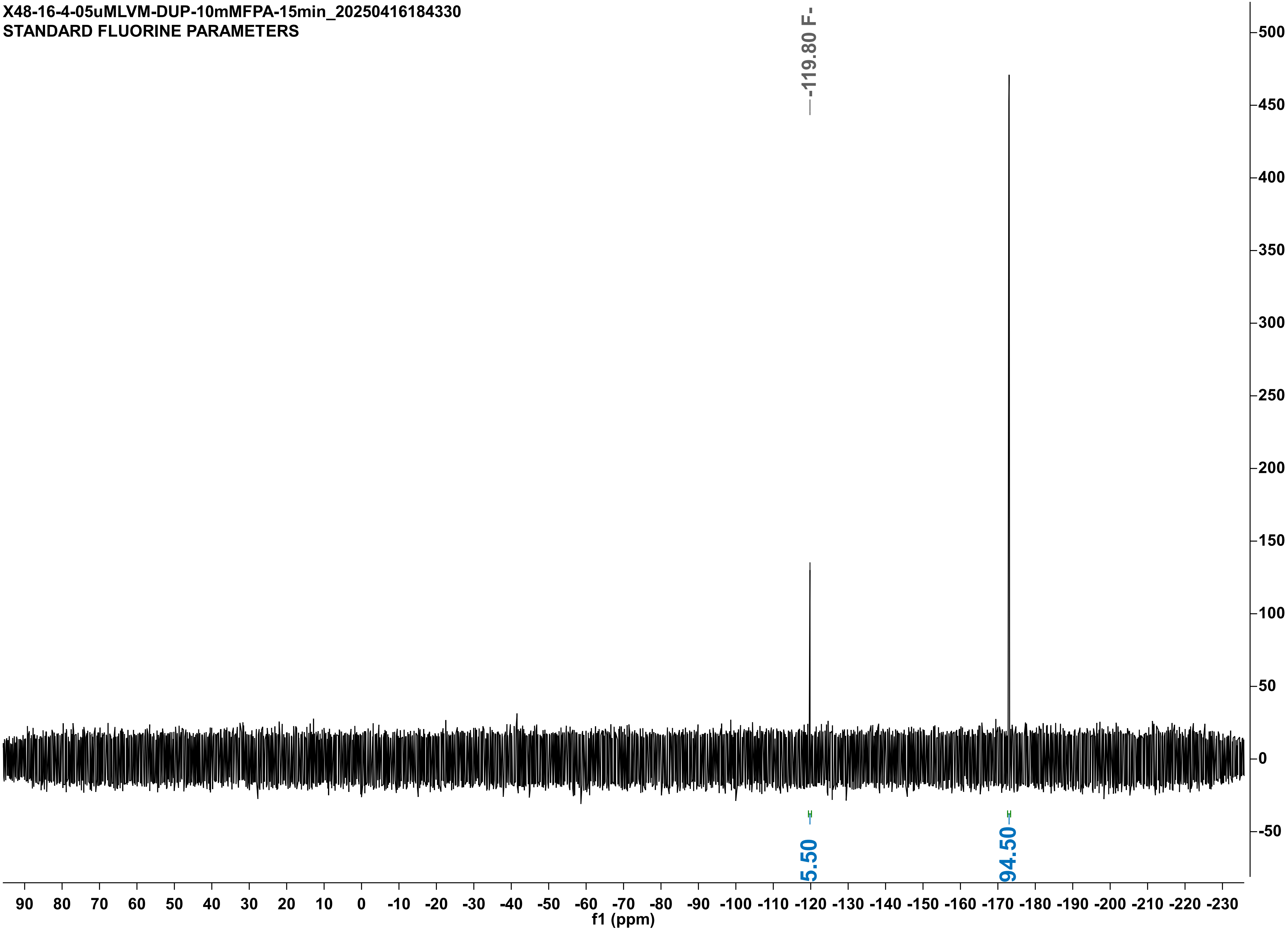

X48-16-4-05uMLVM-10mMFPA-15min_20250416184329
STANDARD FLUORINE PARAMETERS

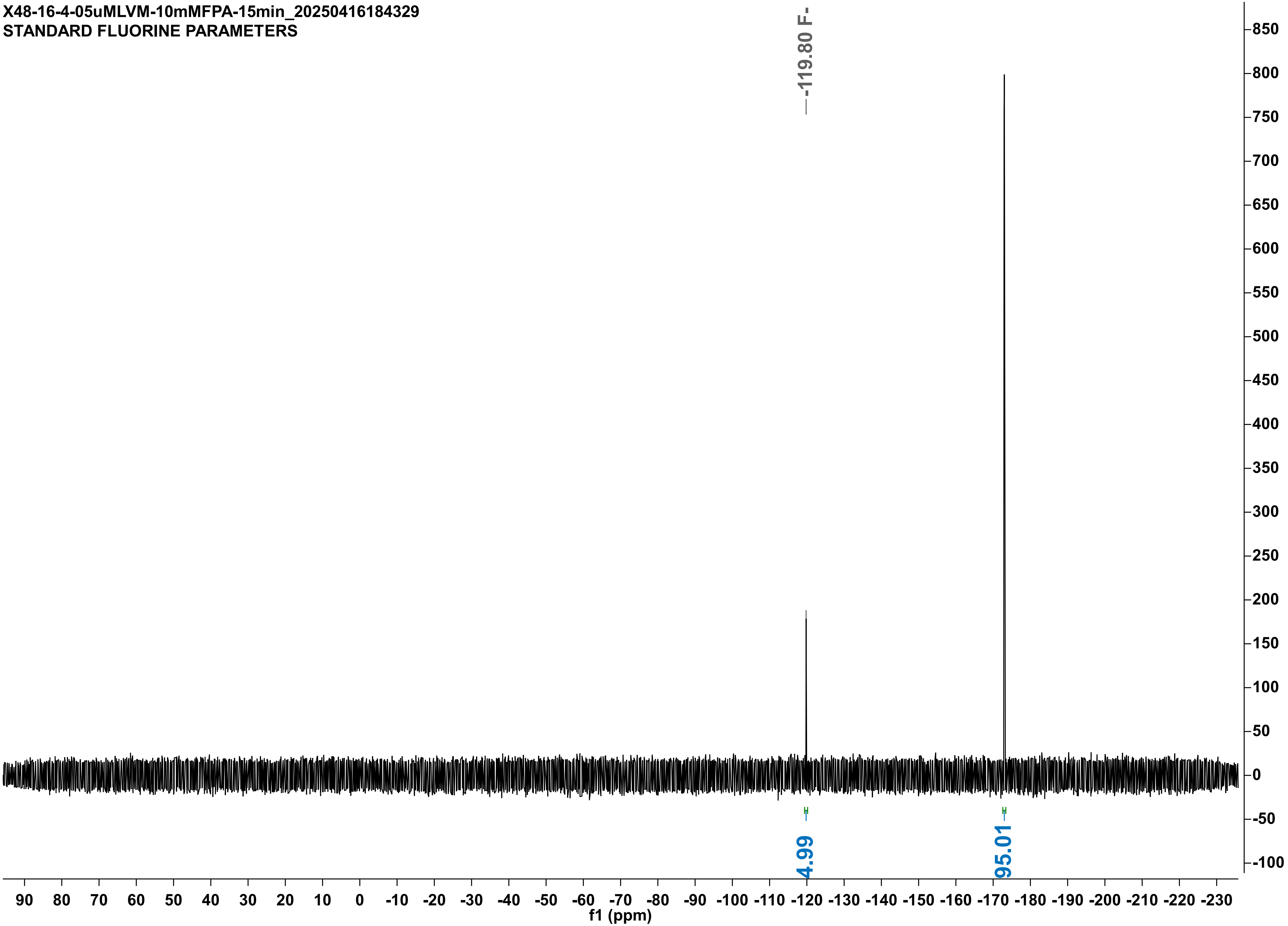

X46-9-4-05uM-FI-DUPLO-10mMFPA-15min-noIS_20250409154310
STANDARD FLUORINE PARAMETERS

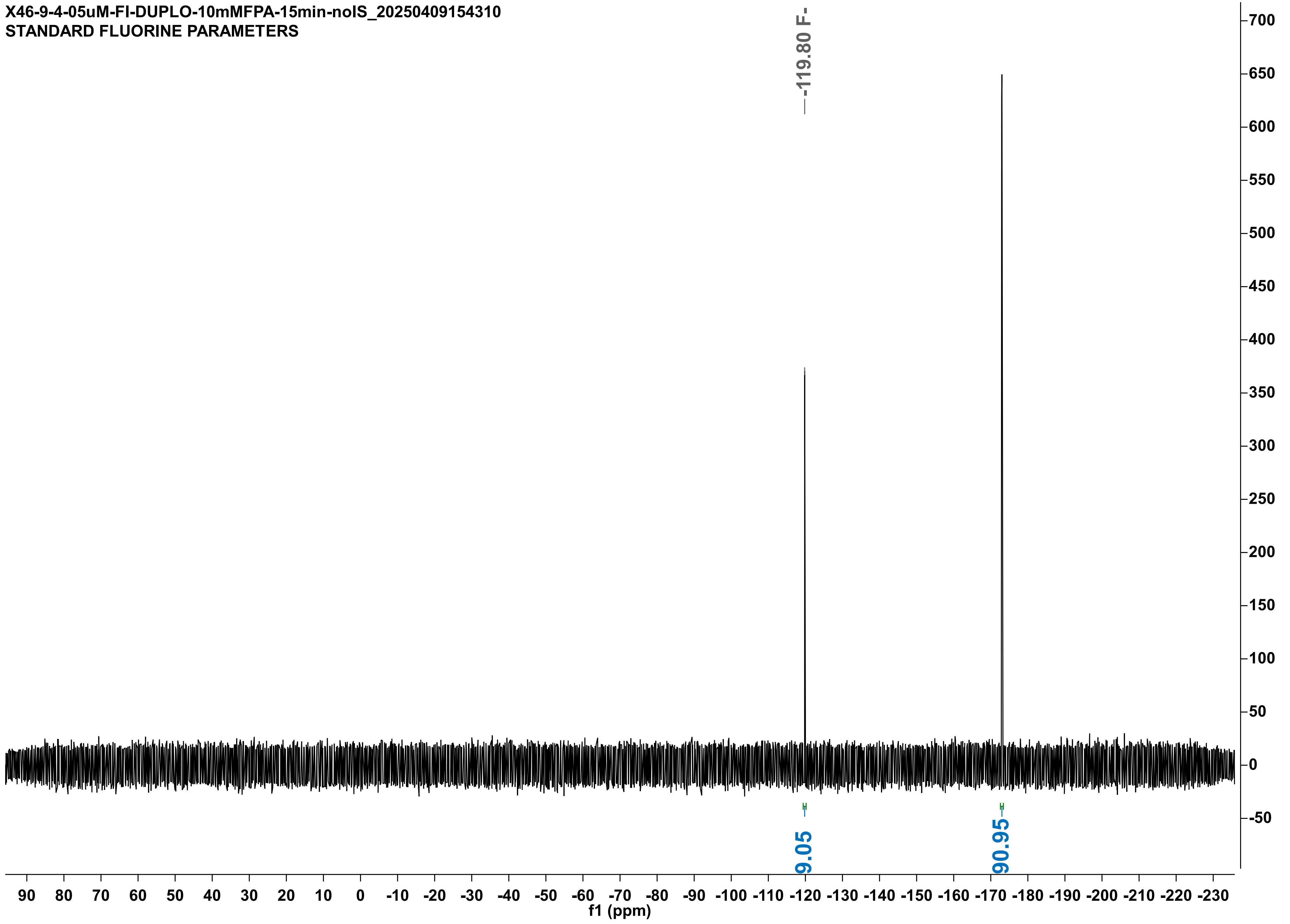

X48-16-4-05uM-FI-10mMFPA-15min_20250416184331
STANDARD FLUORINE PARAMETERS

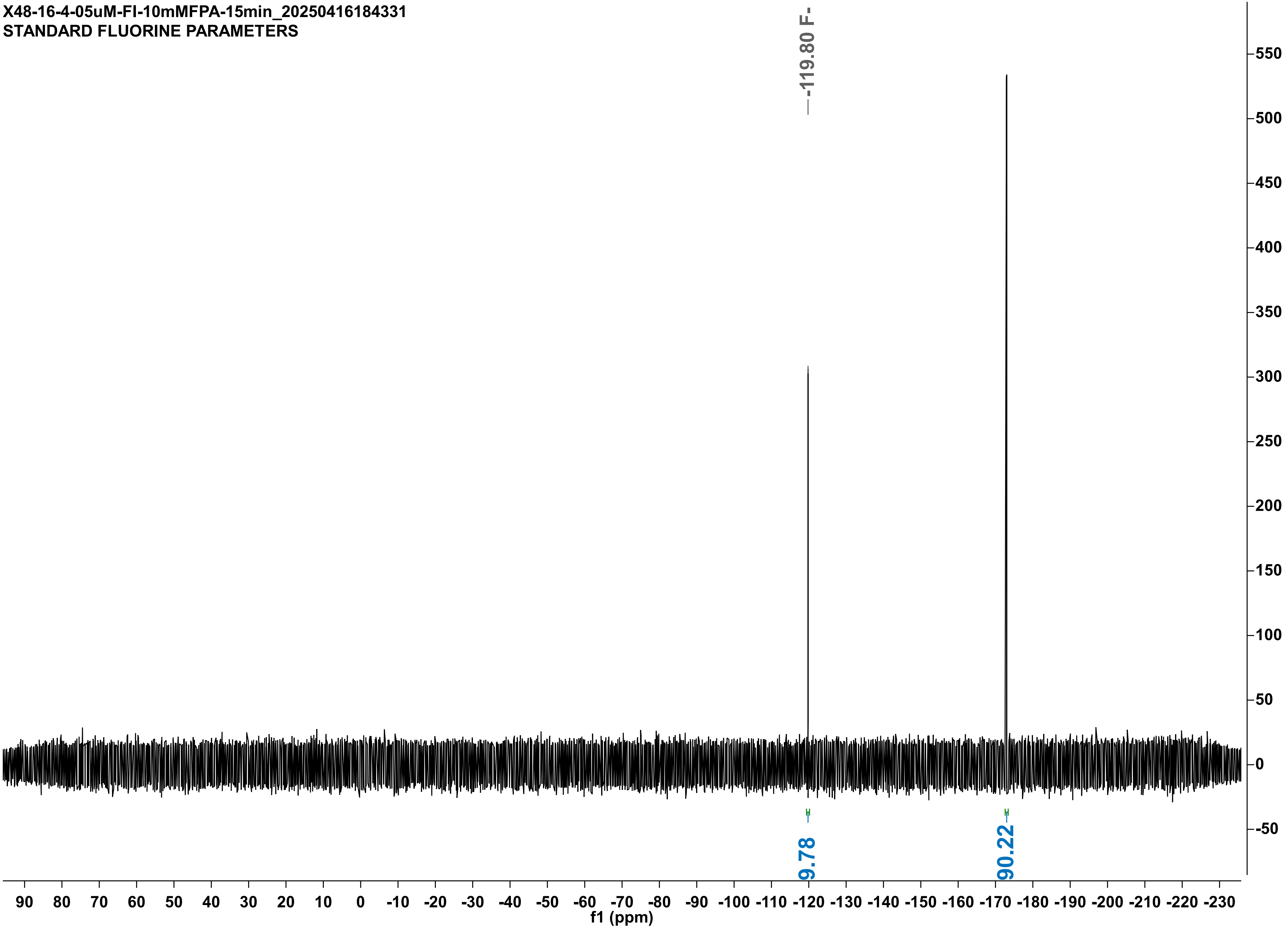

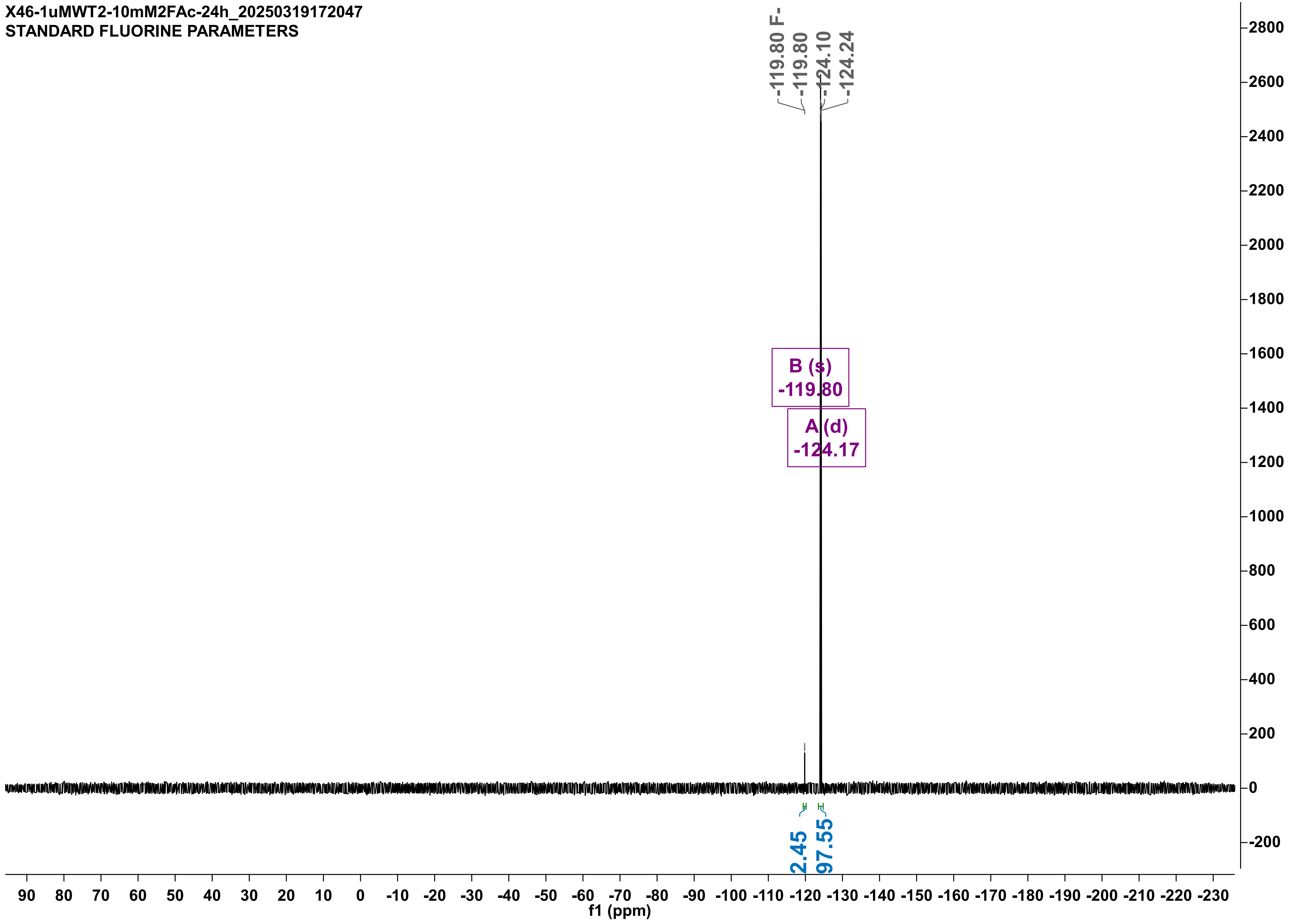
X46-1uMWT2-10mM2FAc-24h_20250319172047
STANDARD FLUORINE PARAMETERS
-119.80 F-
-119.80
-124.10
-124.24
B (s)
-119.80
A (d)
-124.17
2.45
97.55
f1 (ppm)

X46-2-1uMWT1-10mM2FAc-24h_20250328144605
STANDARD FLUORINE PARAMETERS

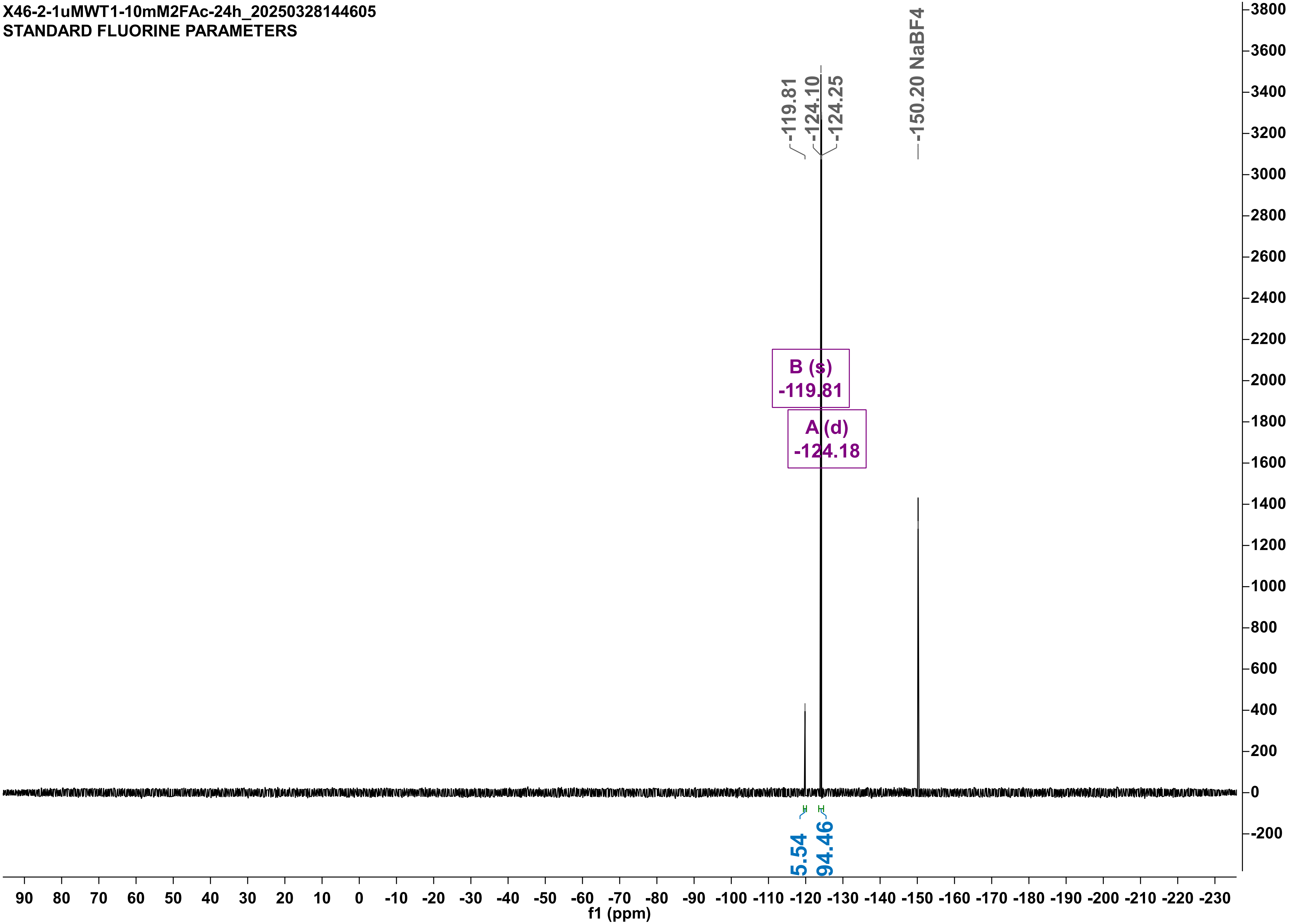

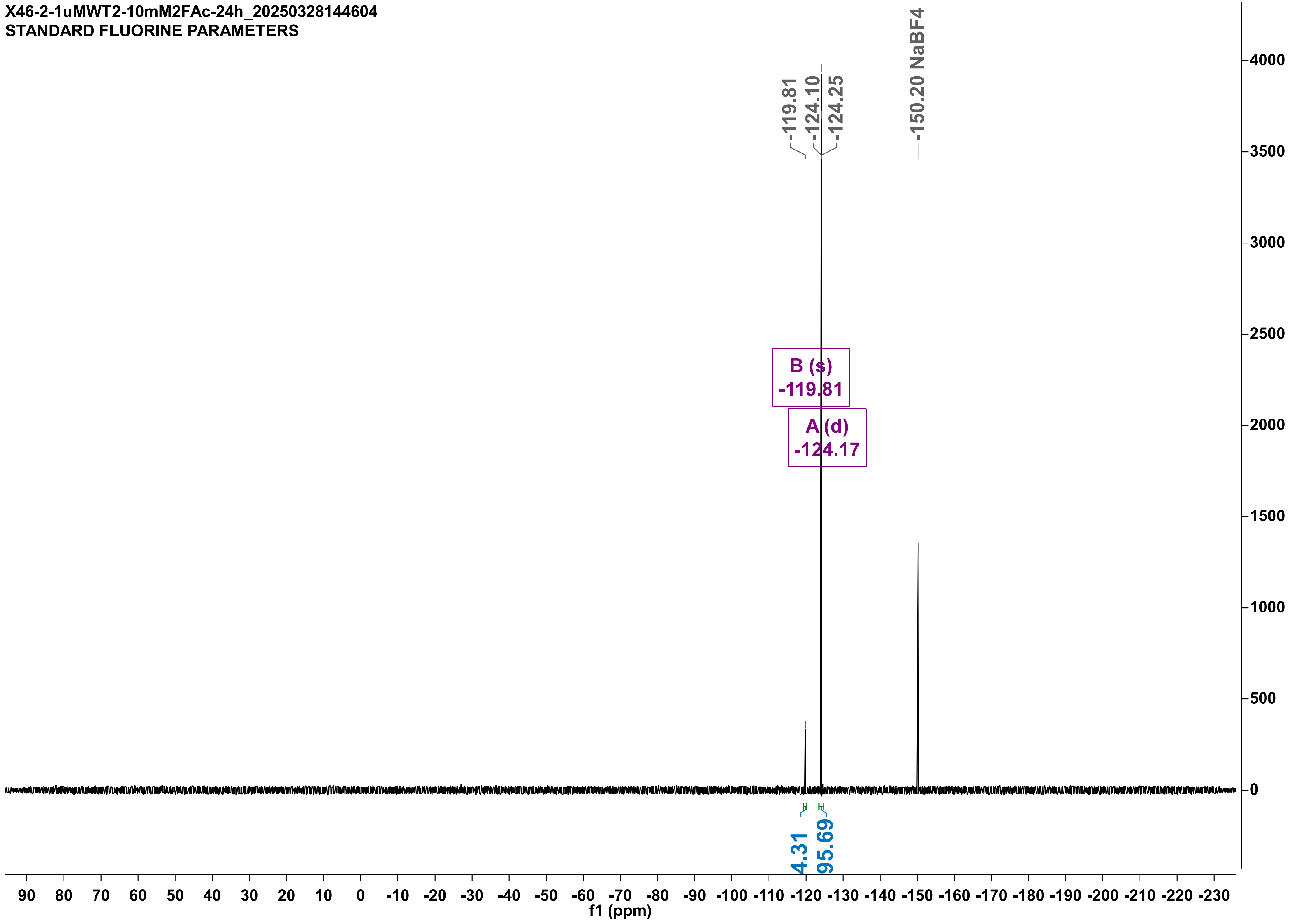
X46-2-1uMWT2-10mM2FAc-24h_20250328144604
STANDARD FLUORINE PARAMETERS
-119.81
-124.10
-124.25
-150.20 NaBF4
B (s)
-119.81
A (d)
-124.17
4.31
95.69
f1 (ppm)

X46-1uMRPL-10mM2FAc-24h_20250319172412
STANDARD FLUORINE PARAMETERS

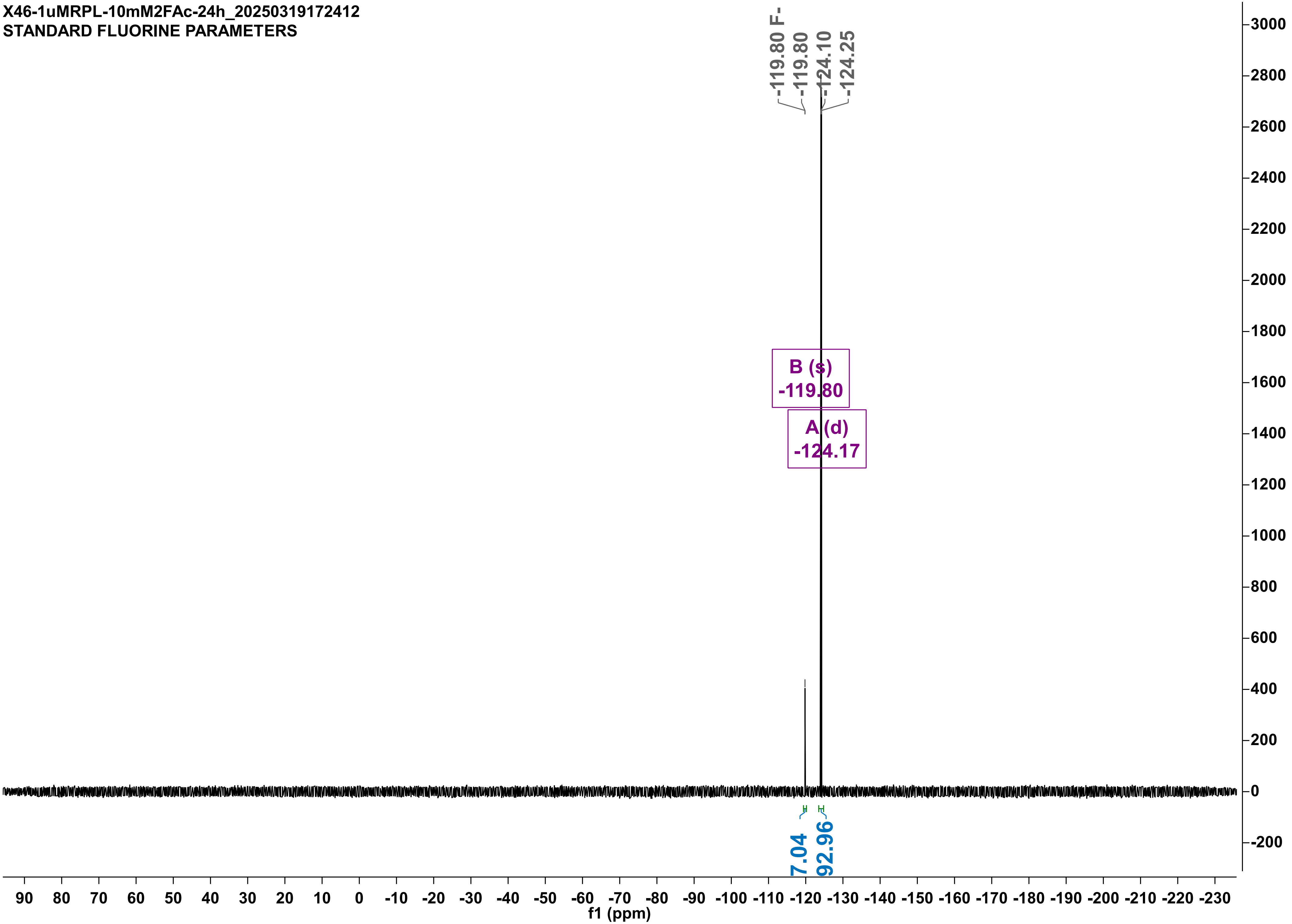

X46-2-1uMRPL-duplo-10mM2FAc-24H-noIS_20250328184254
STANDARD FLUORINE PARAMETERS

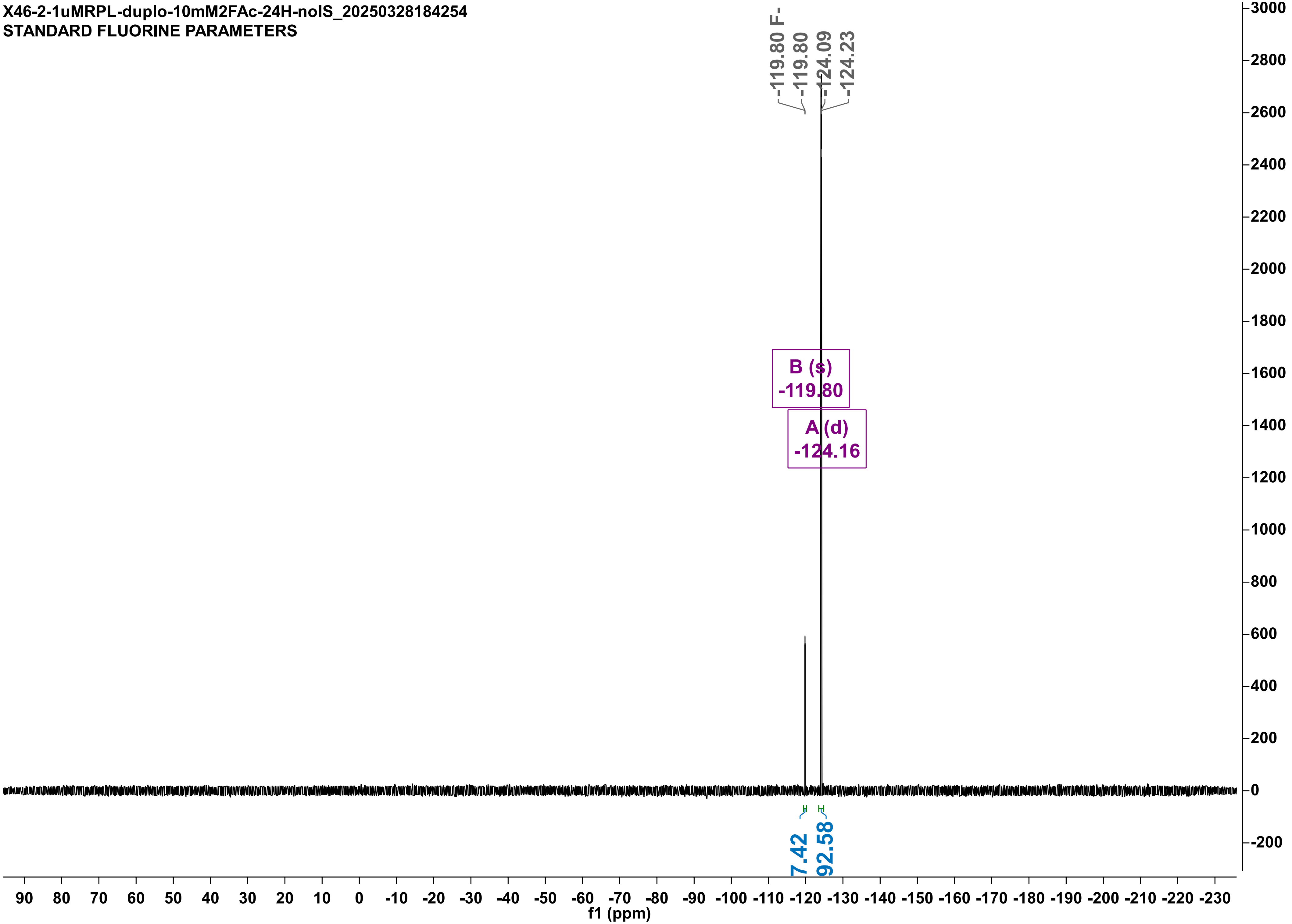

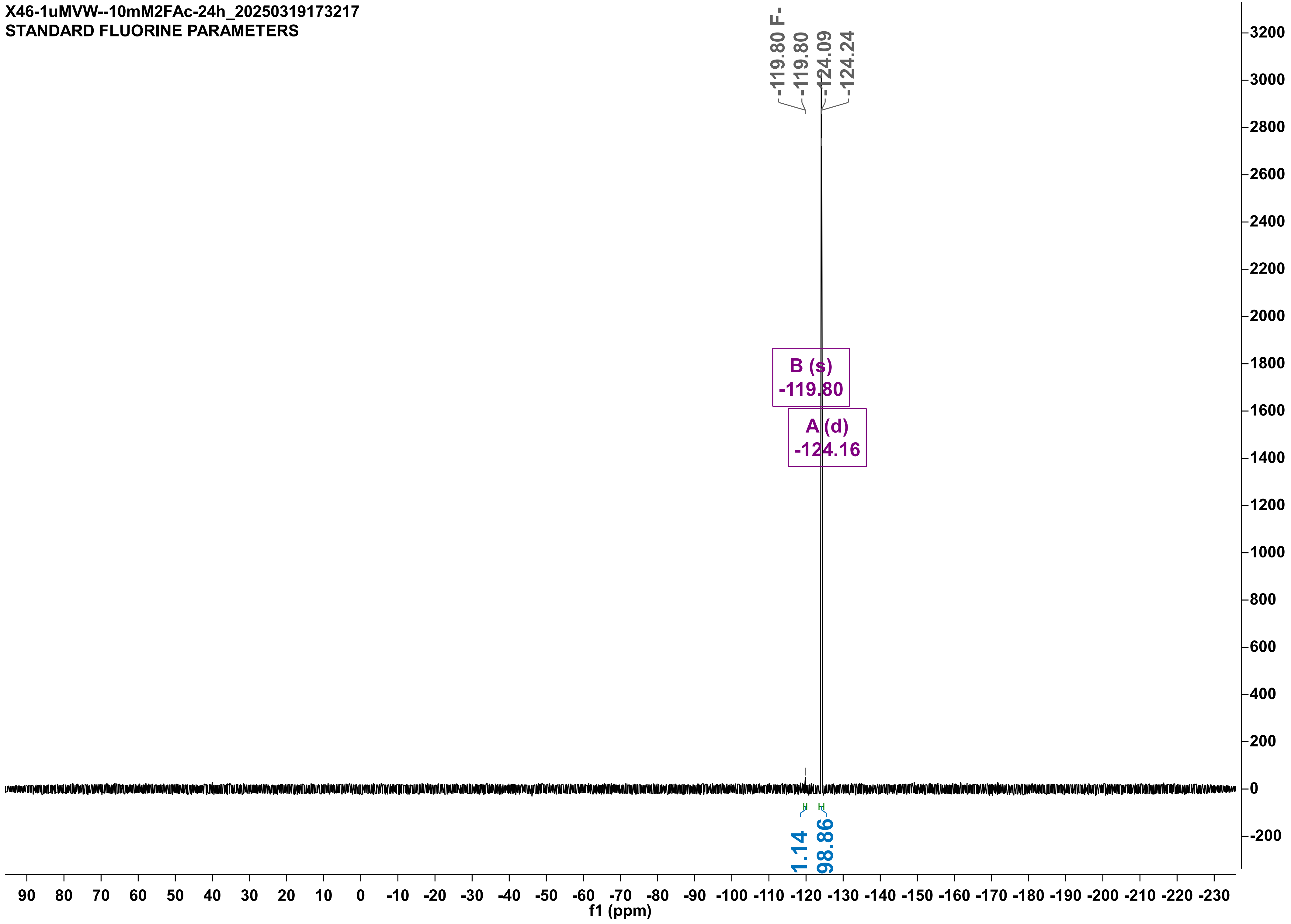

X46-1uMVW--10mM2FAc-24h_20250319173217
STANDARD FLUORINE PARAMETERS
-119.80 F-
-119.80
-124.09
-124.24
B (s)
-119.80
A (d)
-124.16
1.14
98.86
3200
3000
2800
2600
2400
2200
2000
1800
1600
1400
1200
1000
800
600
400
200
0
-200
90 80 70 60 50 40 30 20 10 0 -10 -20 -30 -40 -50 -60 -70 -80 -90 -100 -110 -120 -130 -140 -150 -160 -170 -180 -190 -200 -210 -220 -230
f1 (ppm)

X46-9-4-1uMVW--remeasure-10mM2FAc-24h-noIS_20250410134628
STANDARD FLUORINE PARAMETERS

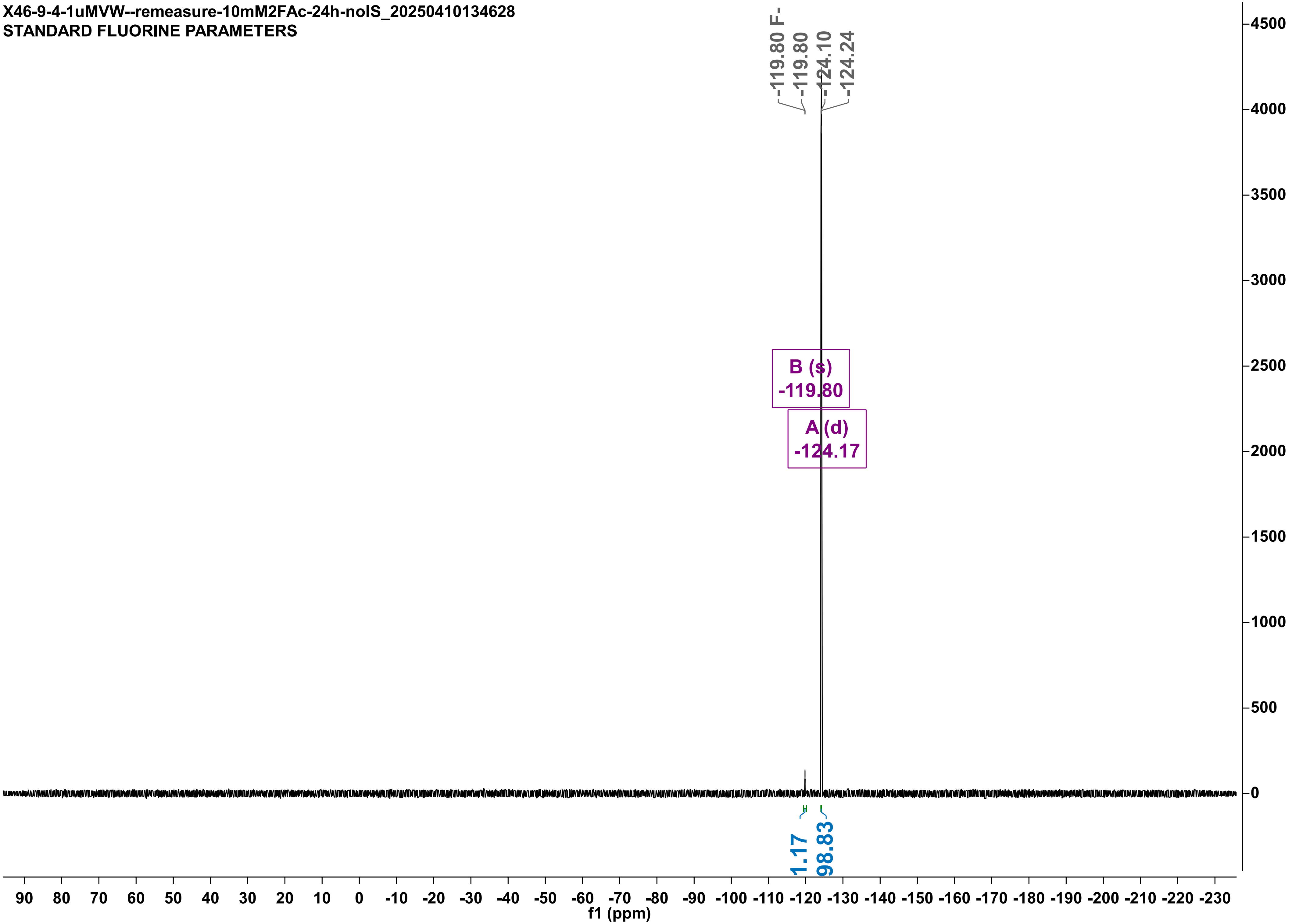

X46-1uMLW--10mM2FAc-24h_20250319173817
STANDARD FLUORINE PARAMETERS

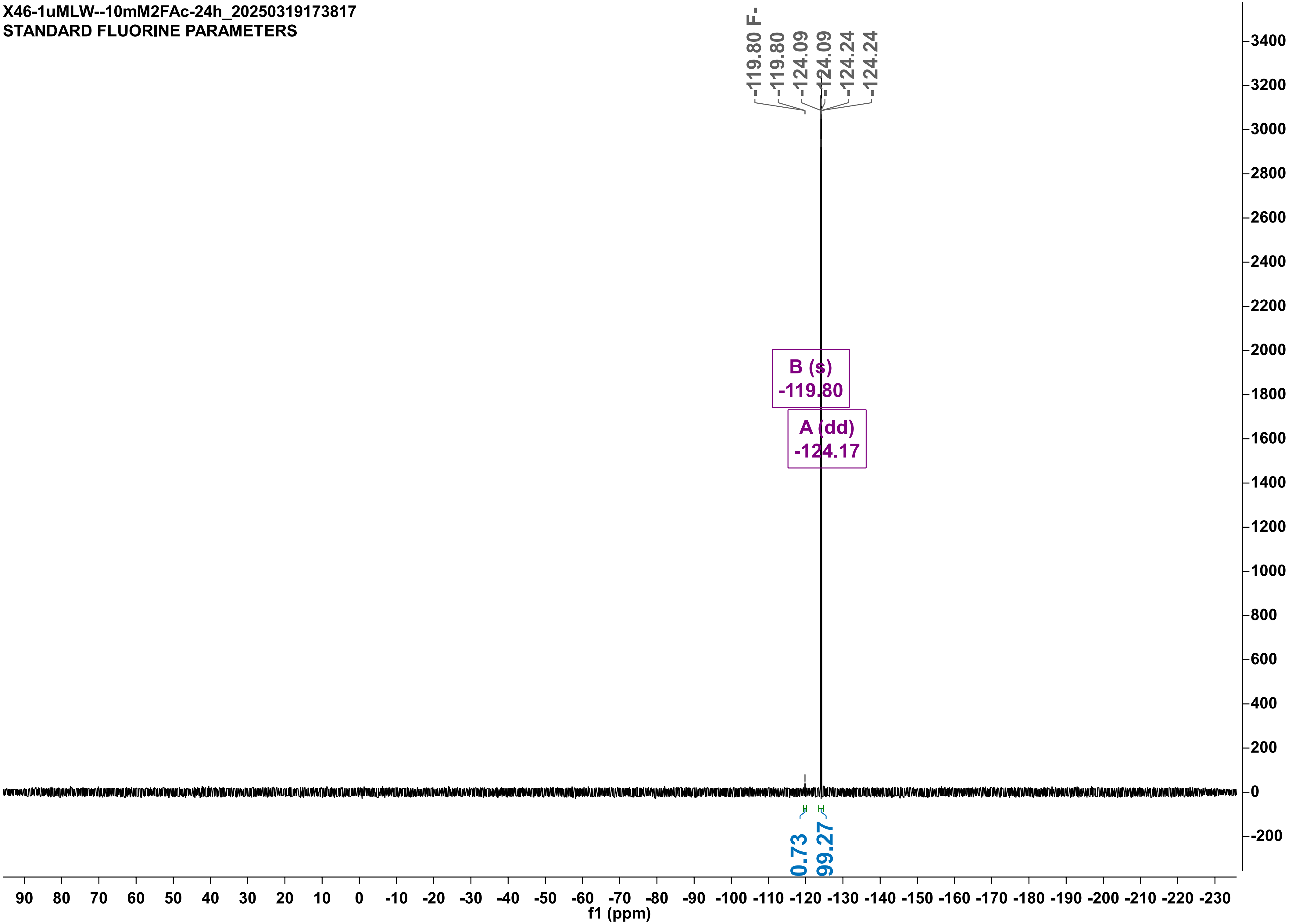

X46-9-4-1uMLW--10mM2FAc-24h-noIS_20250410124351
STANDARD FLUORINE PARAMETERS

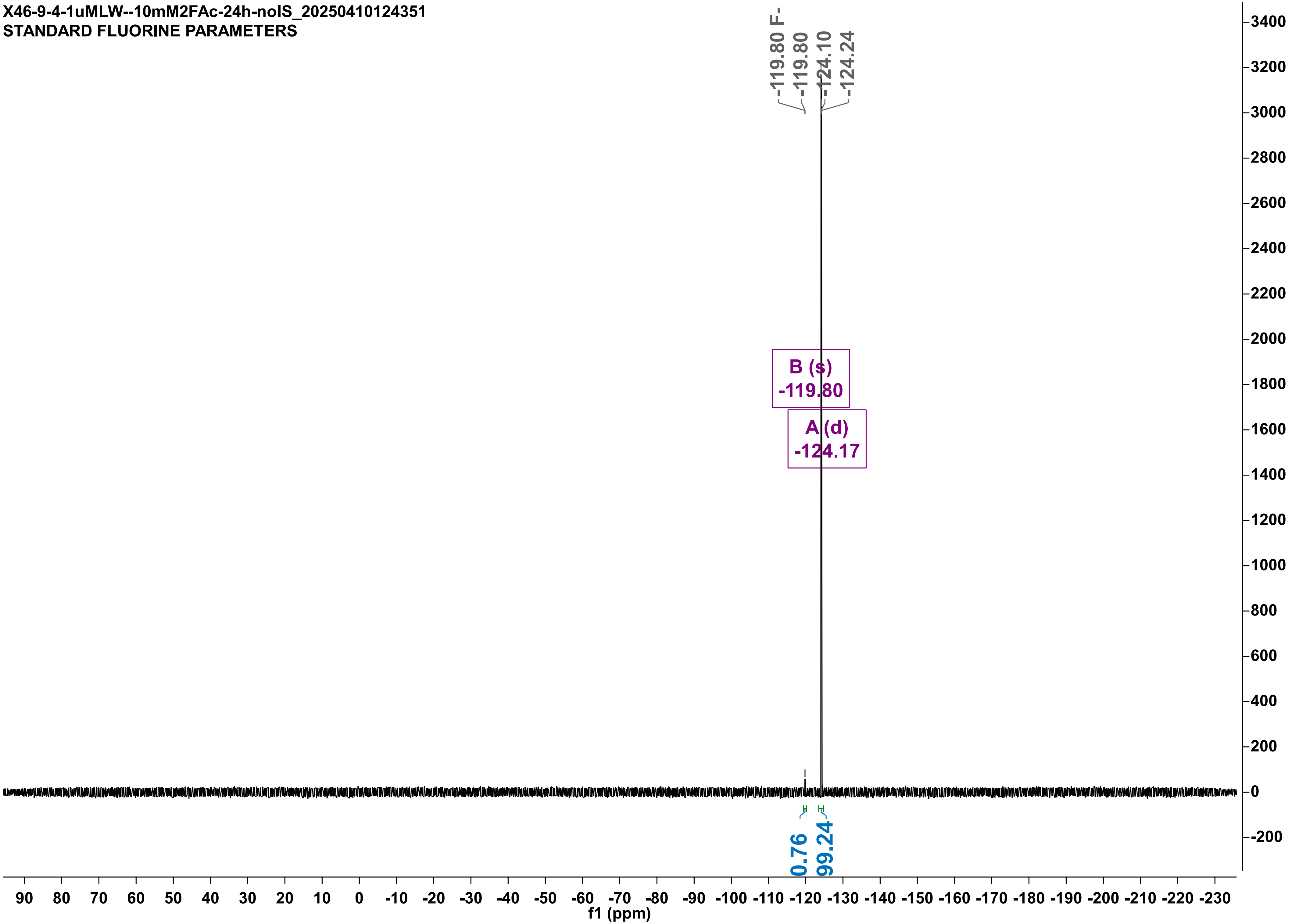

X46-1uMRLM-10mM2FAc-24h_20250321155455
STANDARD FLUORINE PARAMETERS

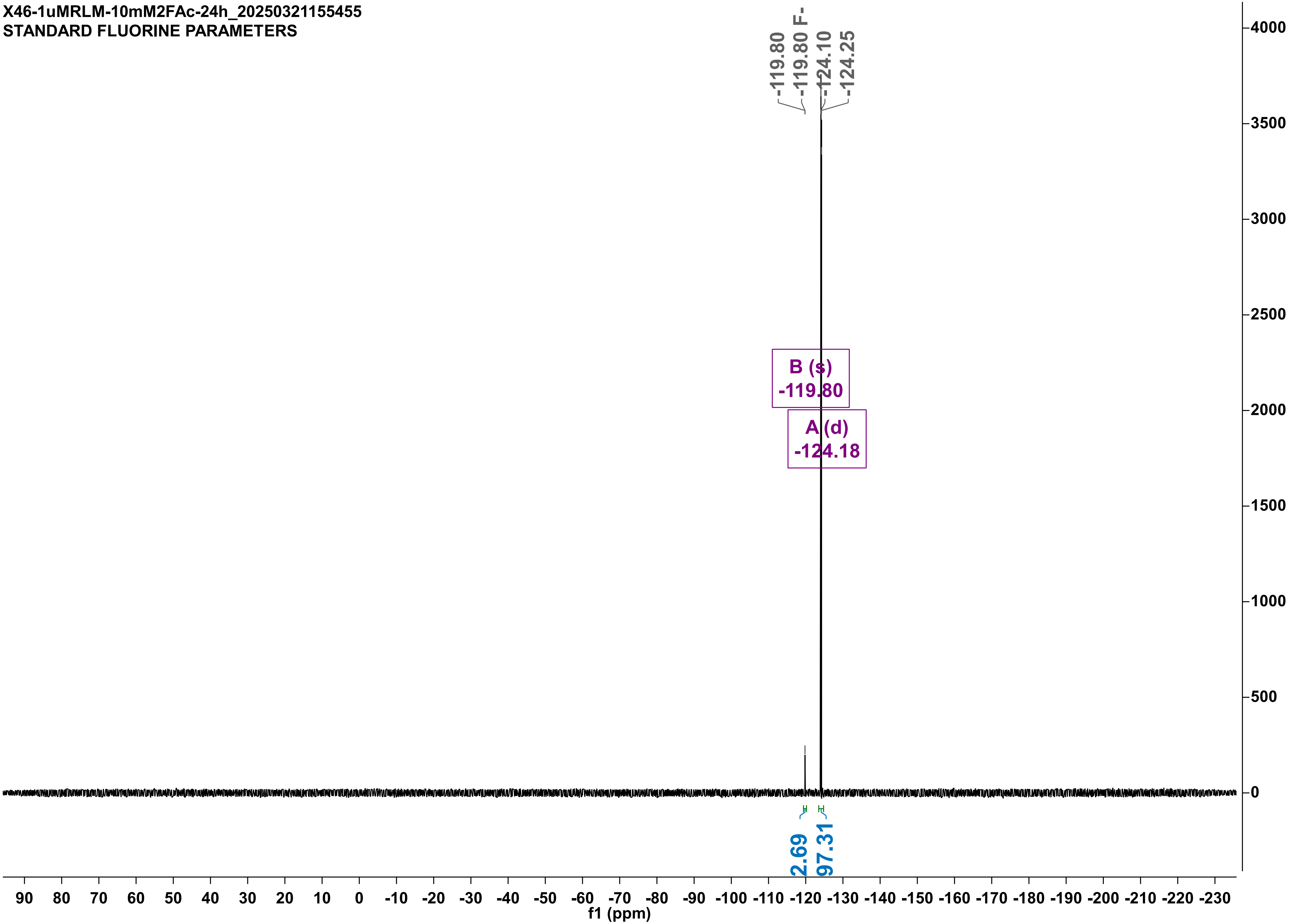

X46-1uMRLM-10mM2FAc-24H_20250327174911
STANDARD FLUORINE PARAMETERS

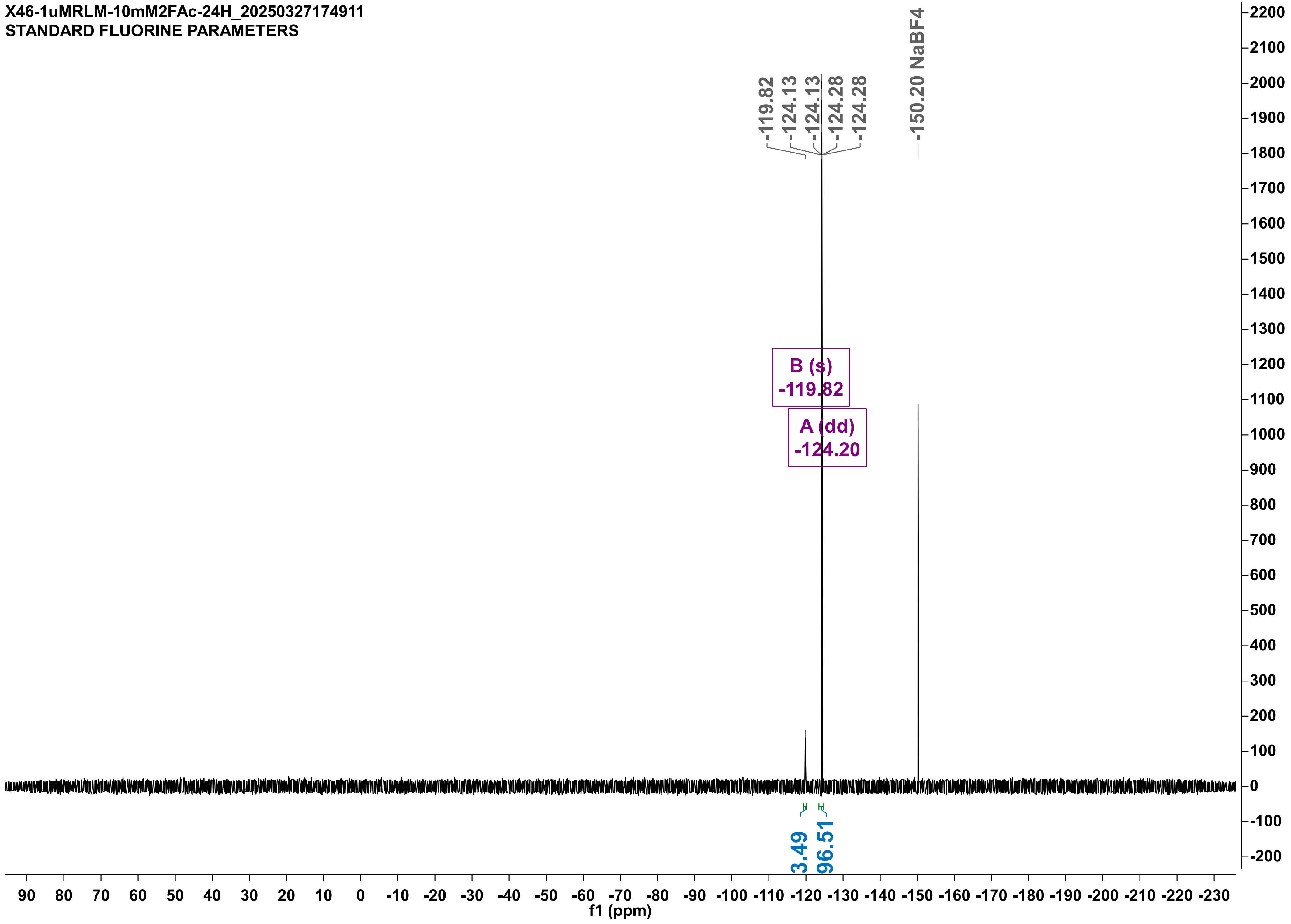

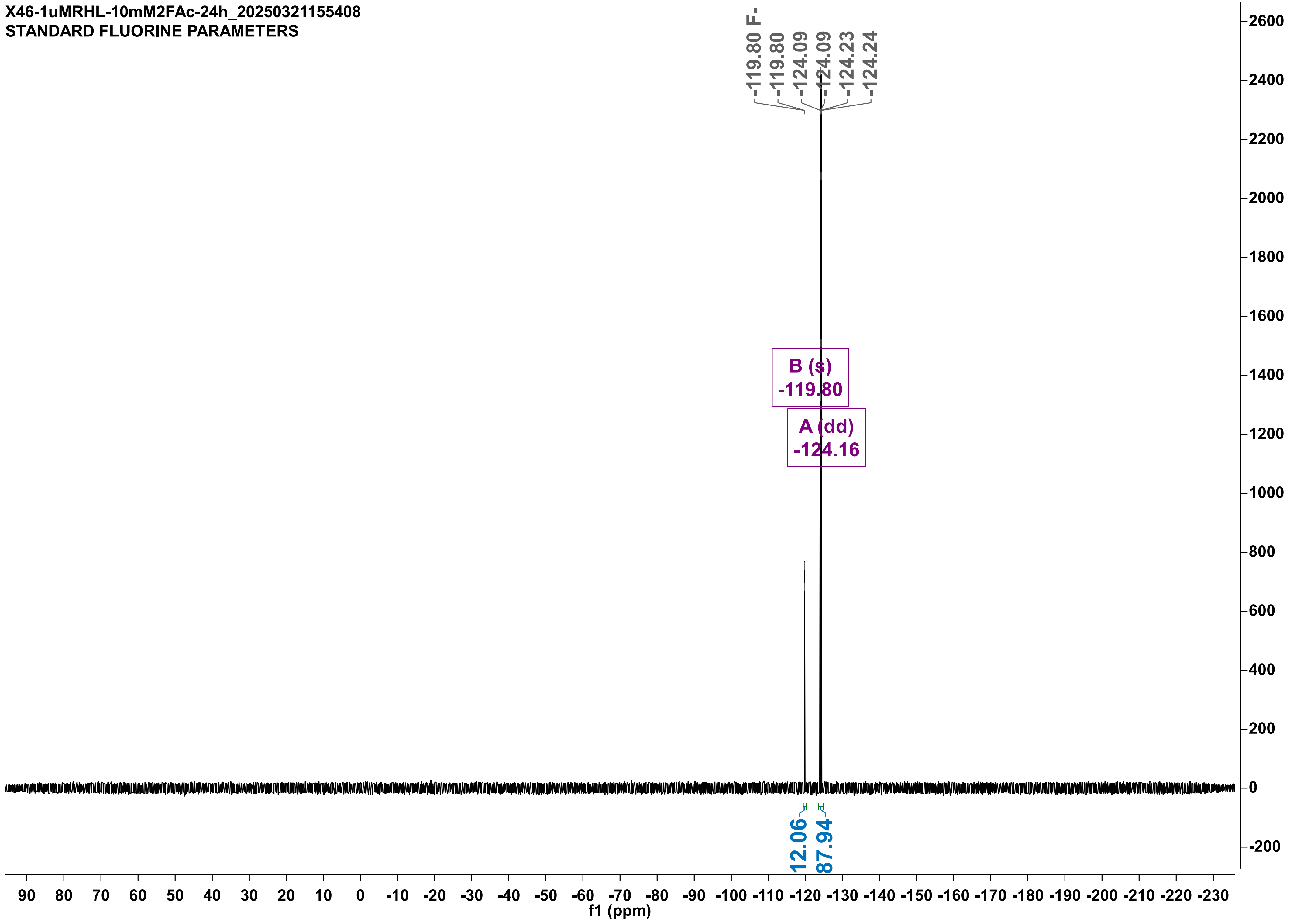

X46-1uMRHL-10mM2FAc-24h_20250321155408
STANDARD FLUORINE PARAMETERS
-119.80 F-
-119.80
-124.09
-124.09
-124.23
-124.24
B (s)
-119.80
A (dd)
-124.16
12.06
87.94
f1 (ppm)

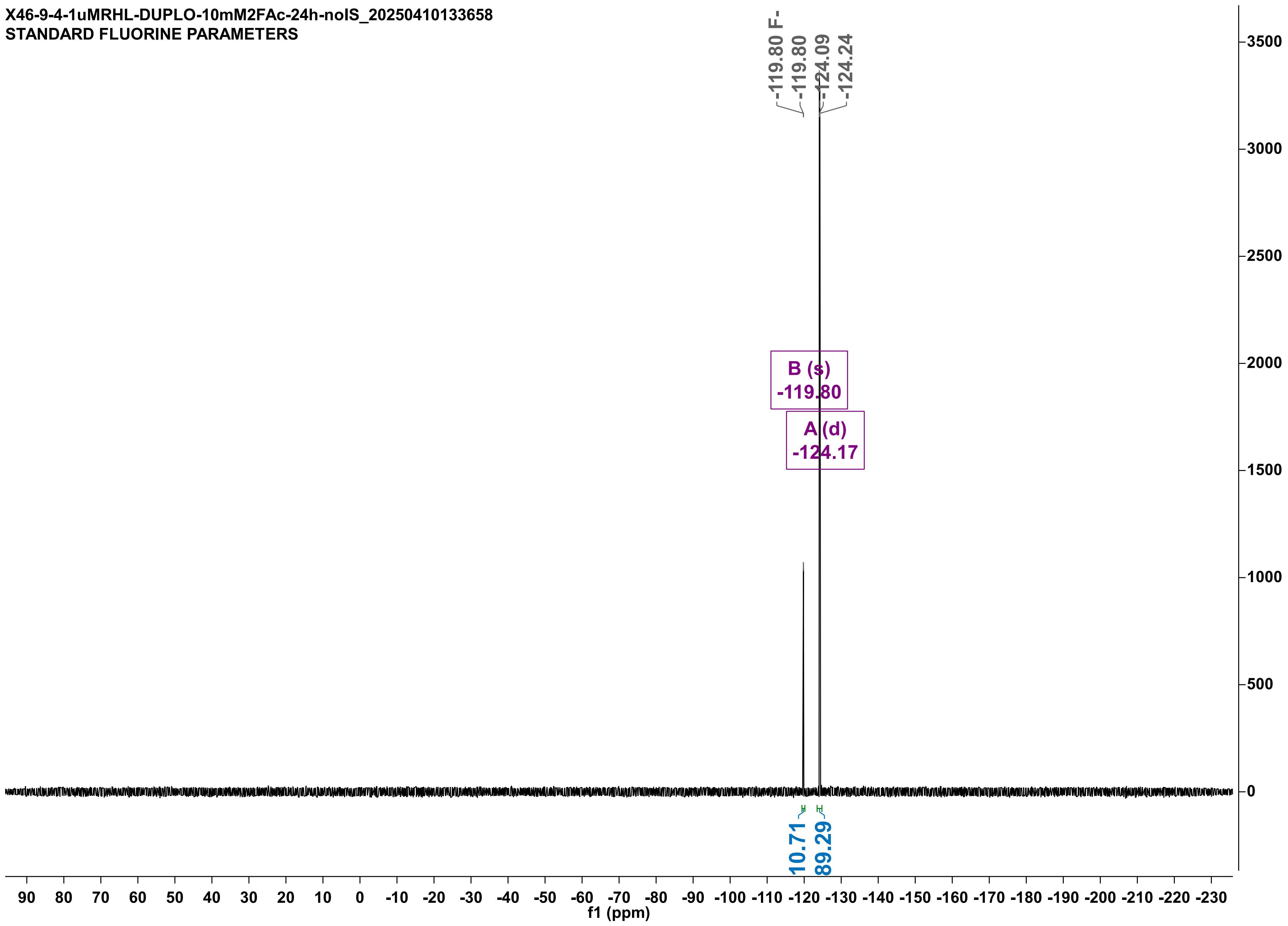

X46-9-4-1uMRHL-DUPLO-10mM2FAc-24h-noIS_20250410133658
STANDARD FLUORINE PARAMETERS
-119.80 F-
-119.80
-124.09
-124.24
B (s)
-119.80
A (d)
-124.17
10.71
89.29
f1 (ppm)

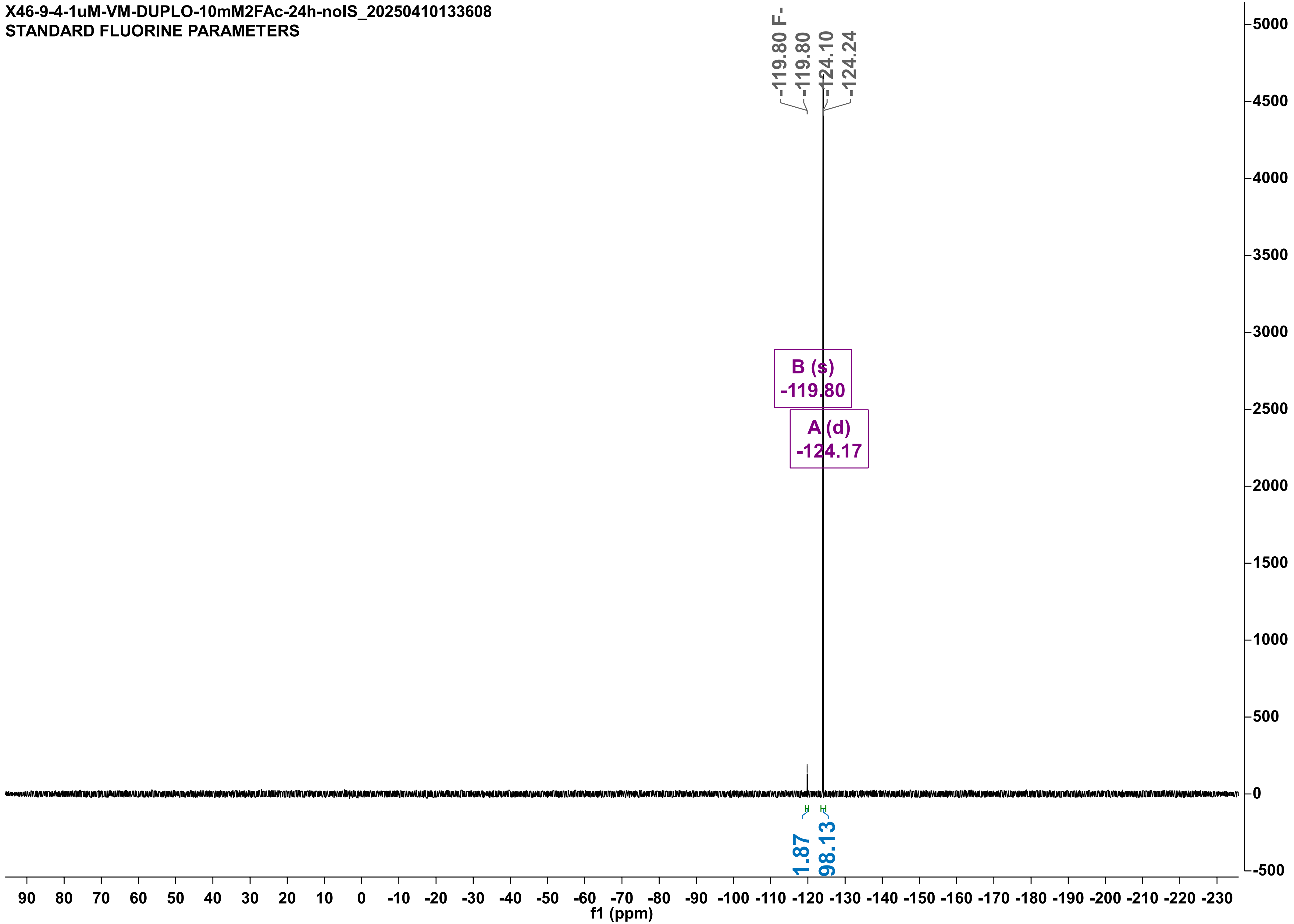

X46-9-4-1uM-VM-DUPLO-10mM2FAc-24h-noIS_20250410133608
STANDARD FLUORINE PARAMETERS
-119.80 F-
-119.80
-124.10
-124.24
B (s)
-119.80
A (d)
-124.17
1.87
98.13
f1 (ppm)

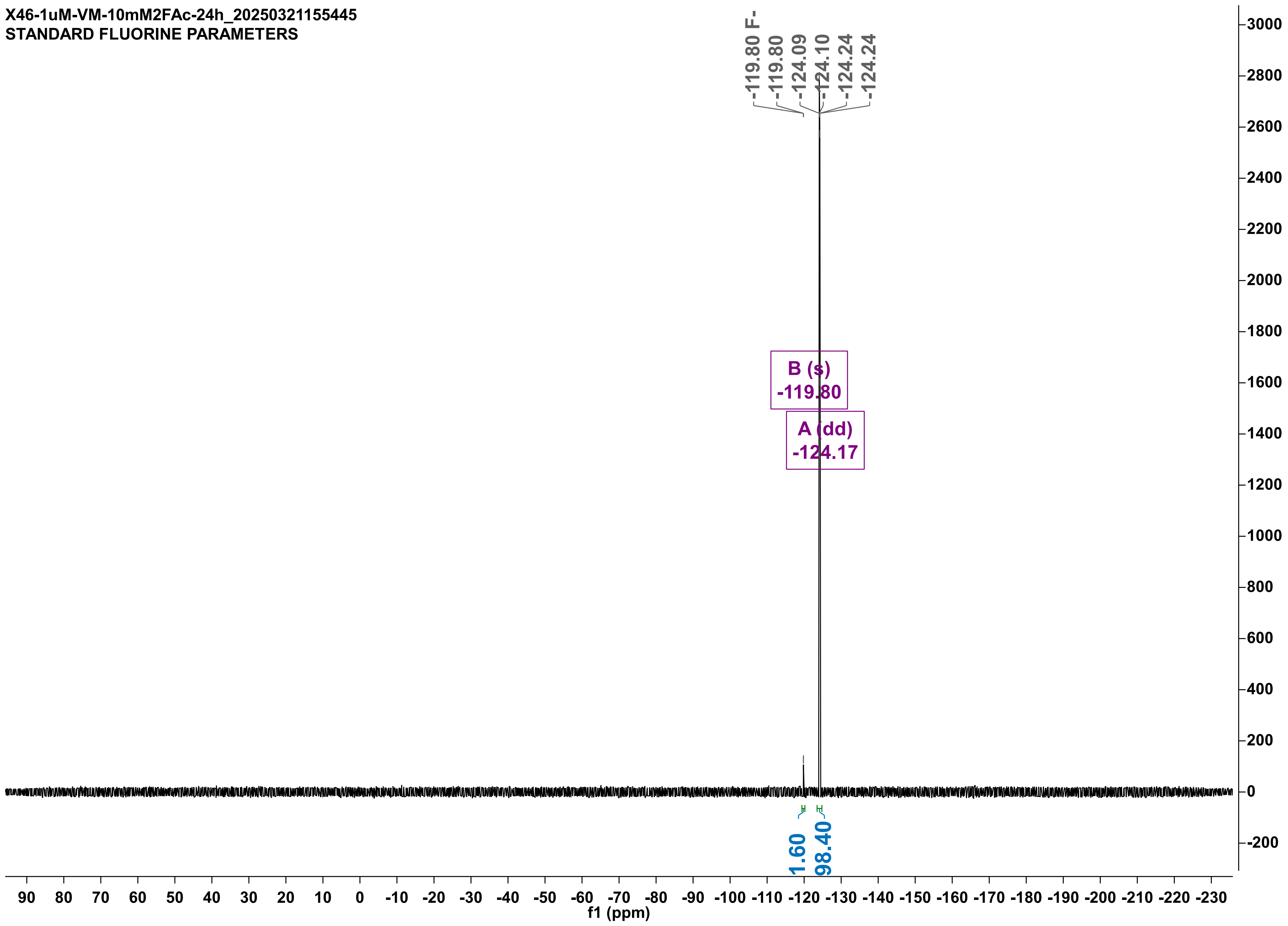
X46-1uM-VM-10mM2FAc-24h_20250321155445
STANDARD FLUORINE PARAMETERS
-119.80 F-
-119.80
-124.09
-124.10
-124.24
-124.24
B (s)
-119.80
A (dd)
-124.17
1.60
98.40
f1 (ppm)

X46-1uMPLM2-10mM2FAc-24h_20250327182014
STANDARD FLUORINE PARAMETERS

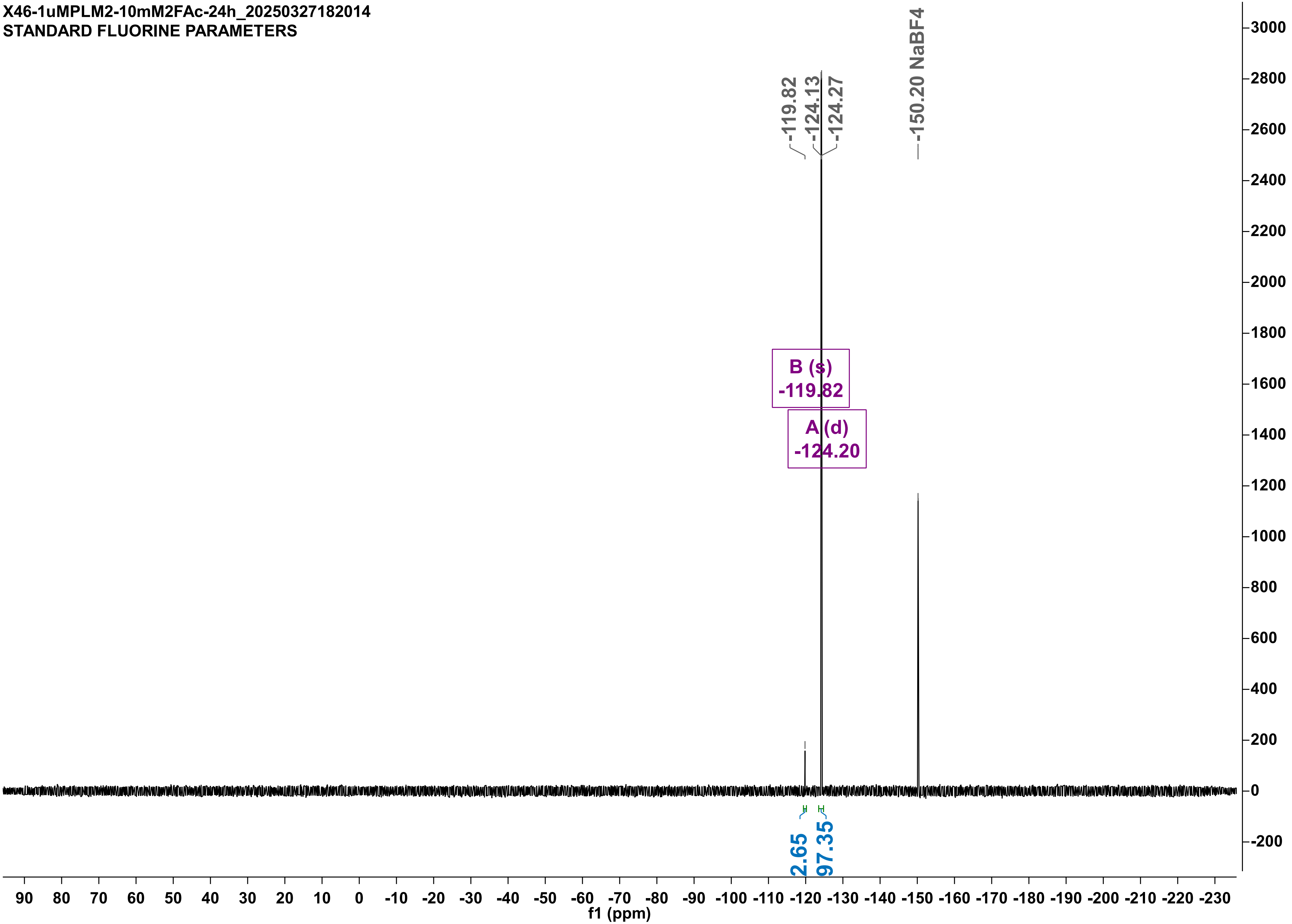

X46-2-1uMPLM2-duplo-10mM2FAc-24H-noIS_20250328184251
STANDARD FLUORINE PARAMETERS

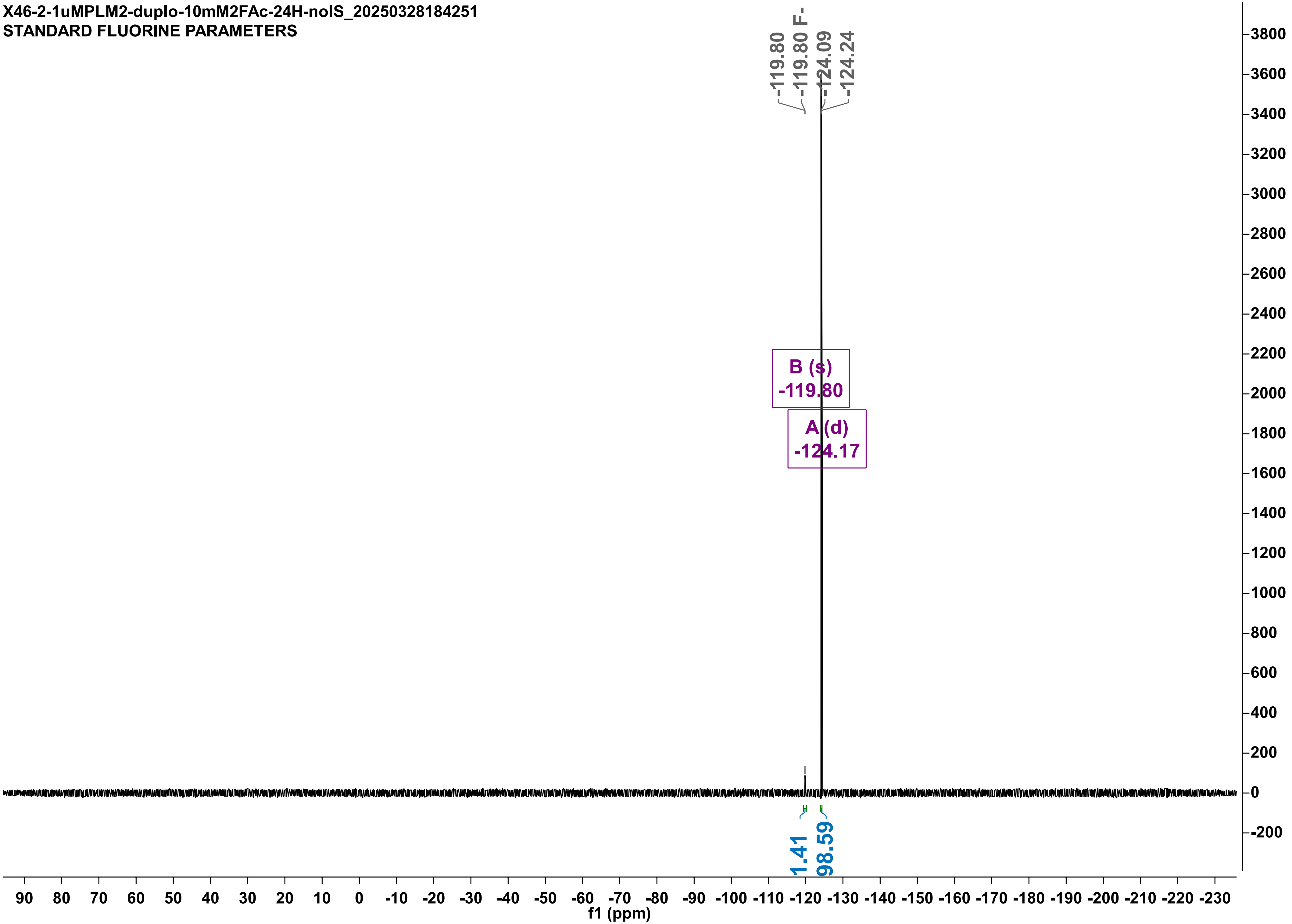

X48-15-4-1uMPLM2-rep-10mM2FAc-24h_20250415175946
STANDARD FLUORINE PARAMETERS

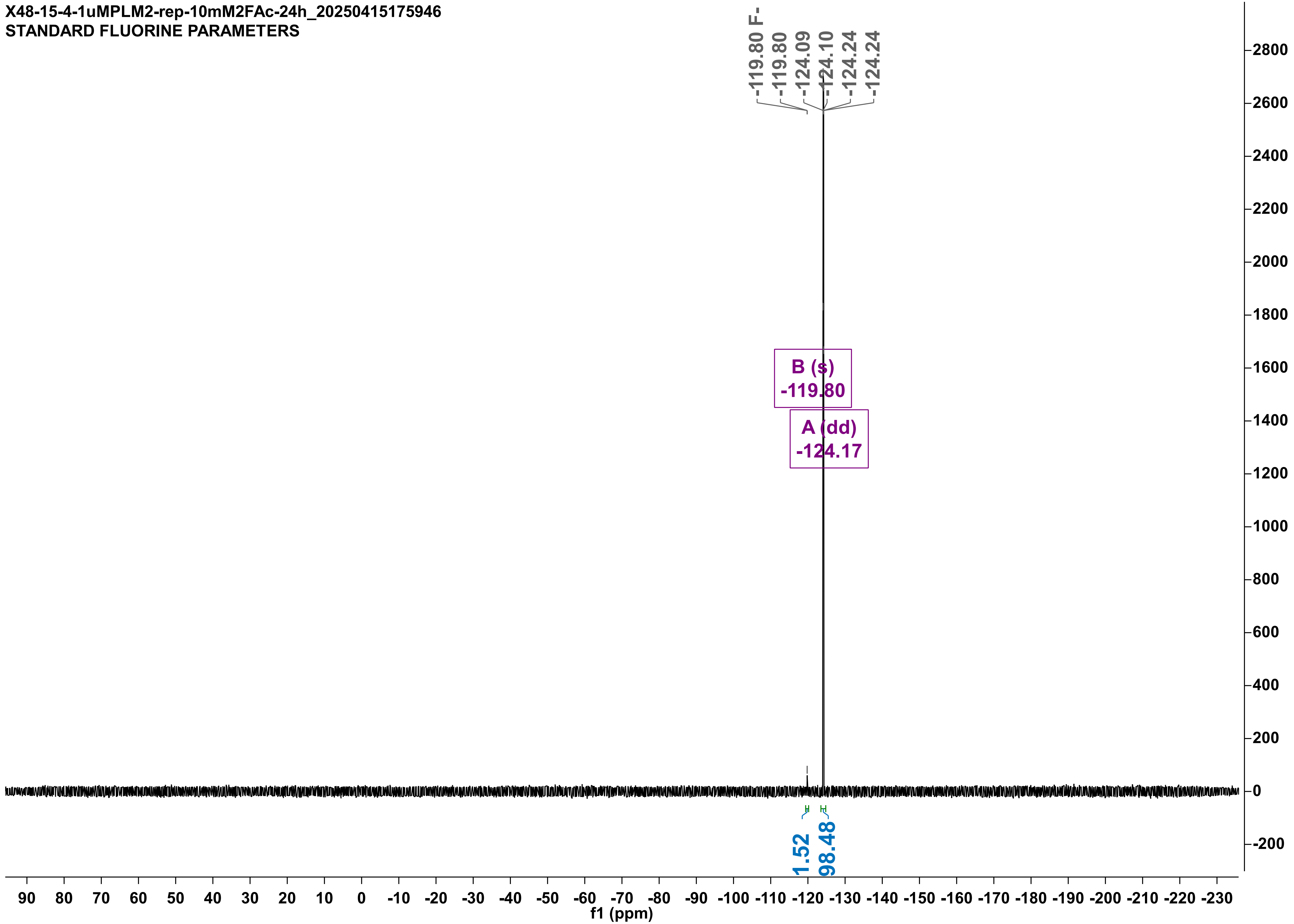

X46-1uM-ML-10mM2FAc-24H_20250327180540
STANDARD FLUORINE PARAMETERS

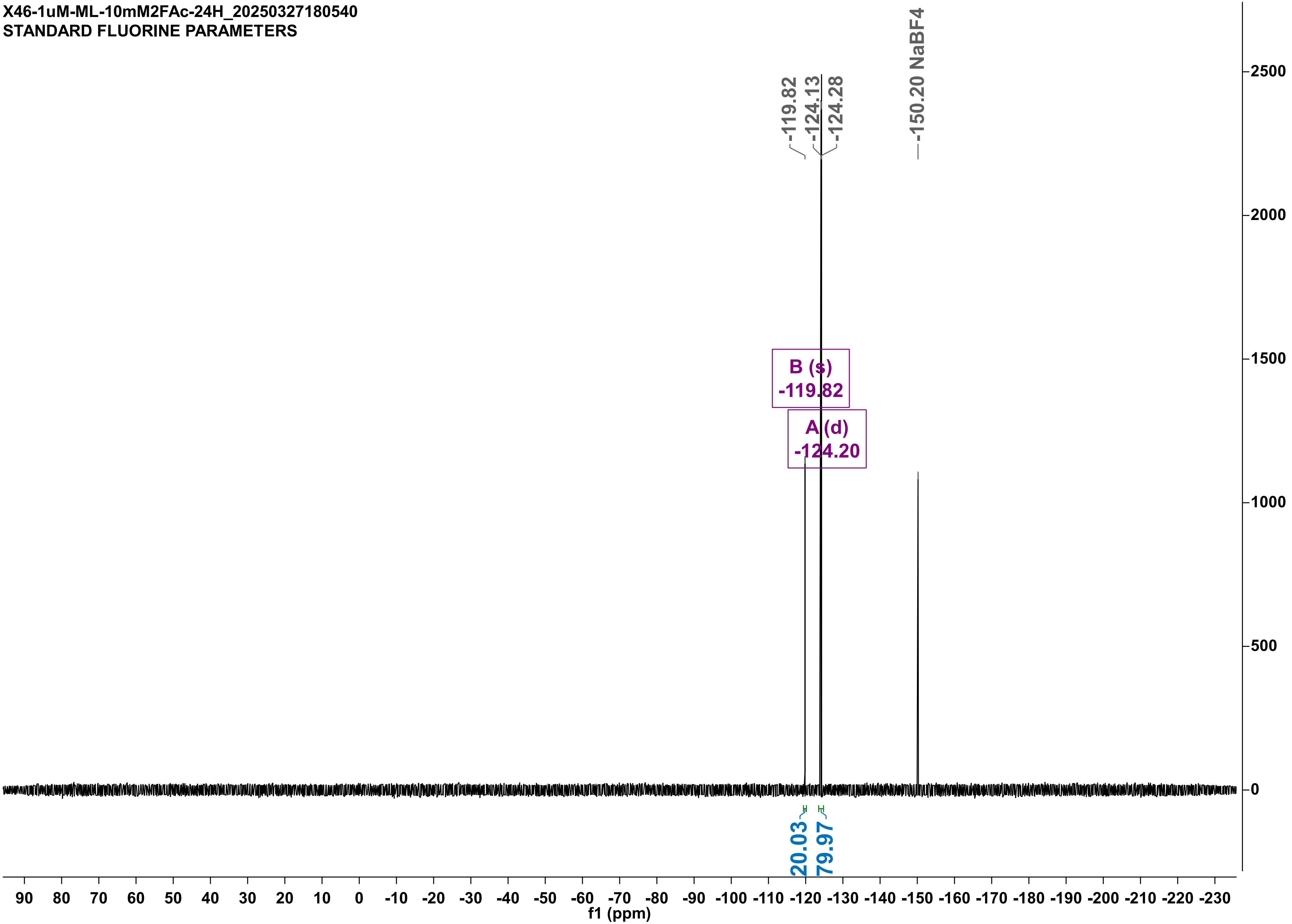

X46-2-1uM-ML-duplo-10mM2FAc-24H-noIS_20250328172252
STANDARD FLUORINE PARAMETERS

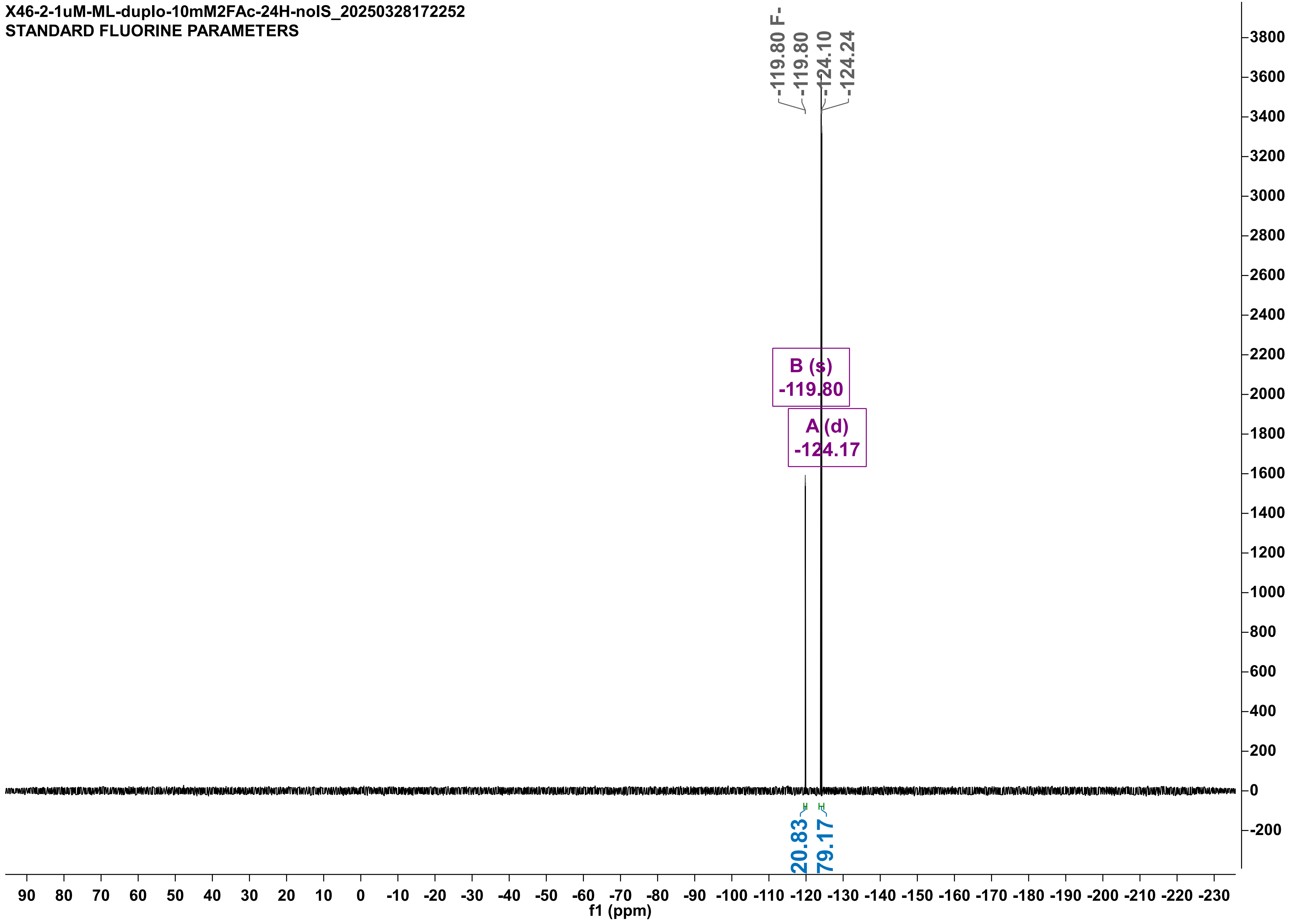

X48-15-4-1uM-ML-rep-10mM2FAc-24h_20250415180014
STANDARD FLUORINE PARAMETERS

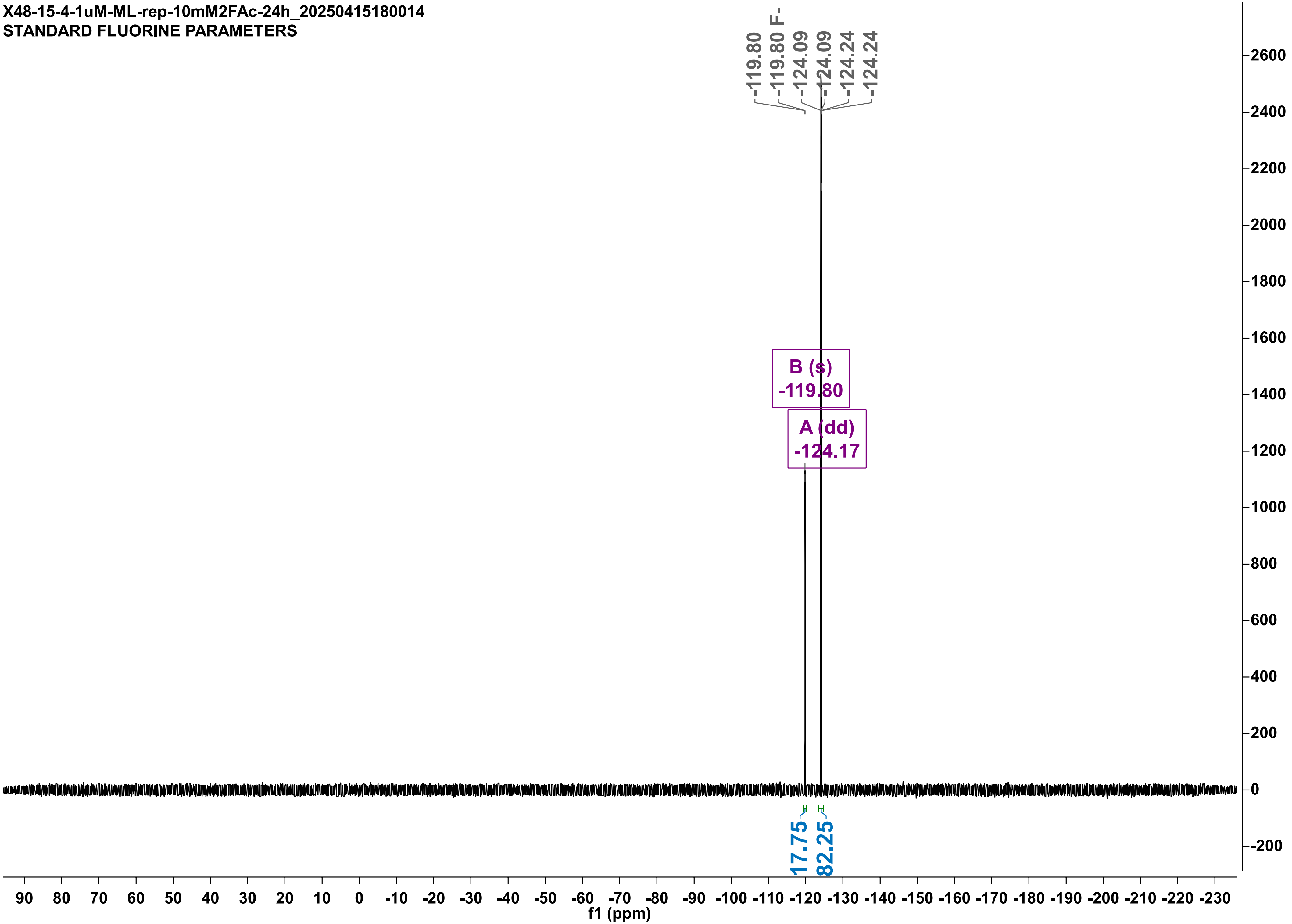

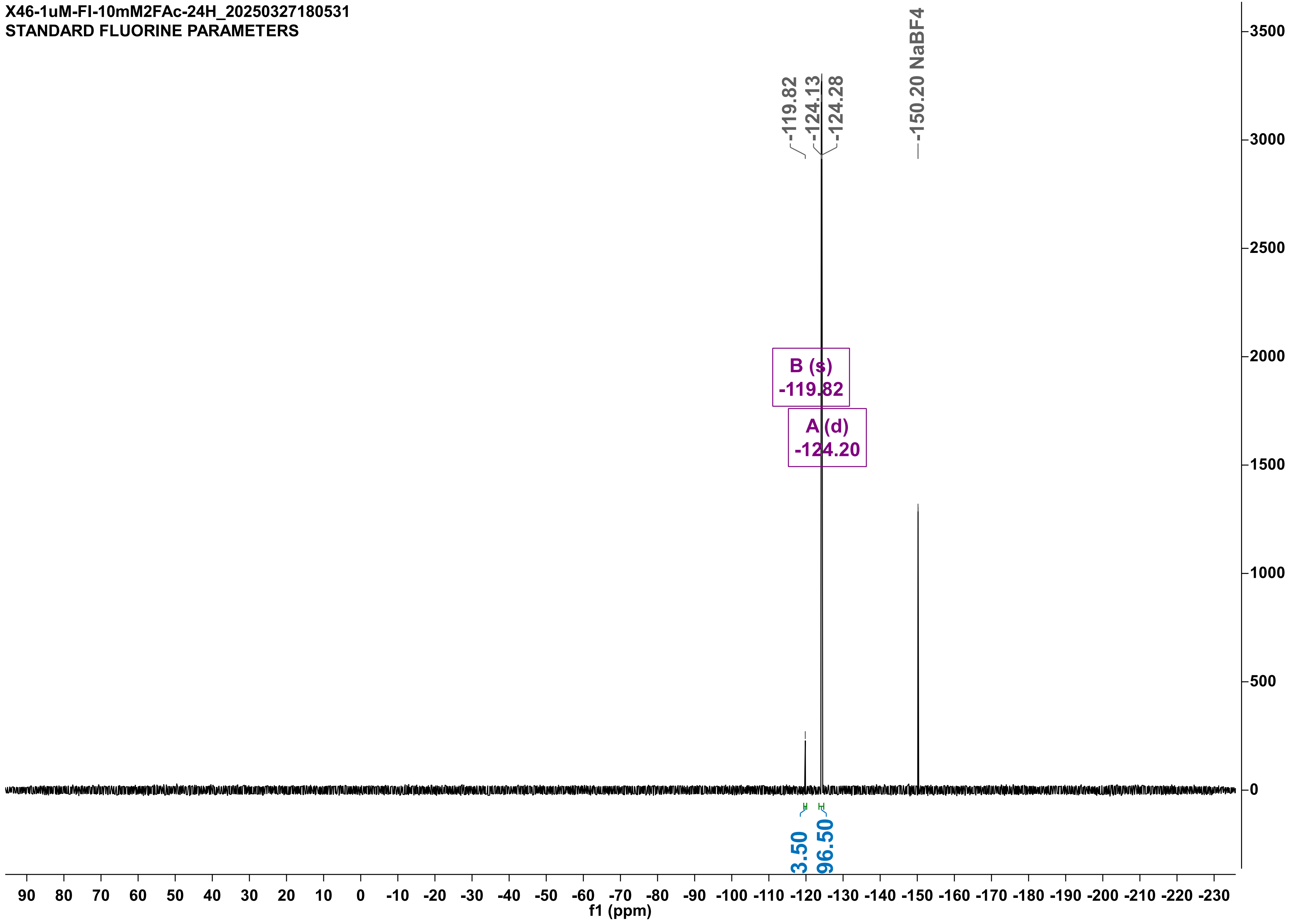
X46-1uM-FI-10mM2FAc-24H_20250327180531
STANDARD FLUORINE PARAMETERS
-119.82
-124.13
-124.28
-150.20 NaBF4
B (s)
-119.82
A (d)
-124.20
3.50
96.50
f1 (ppm)

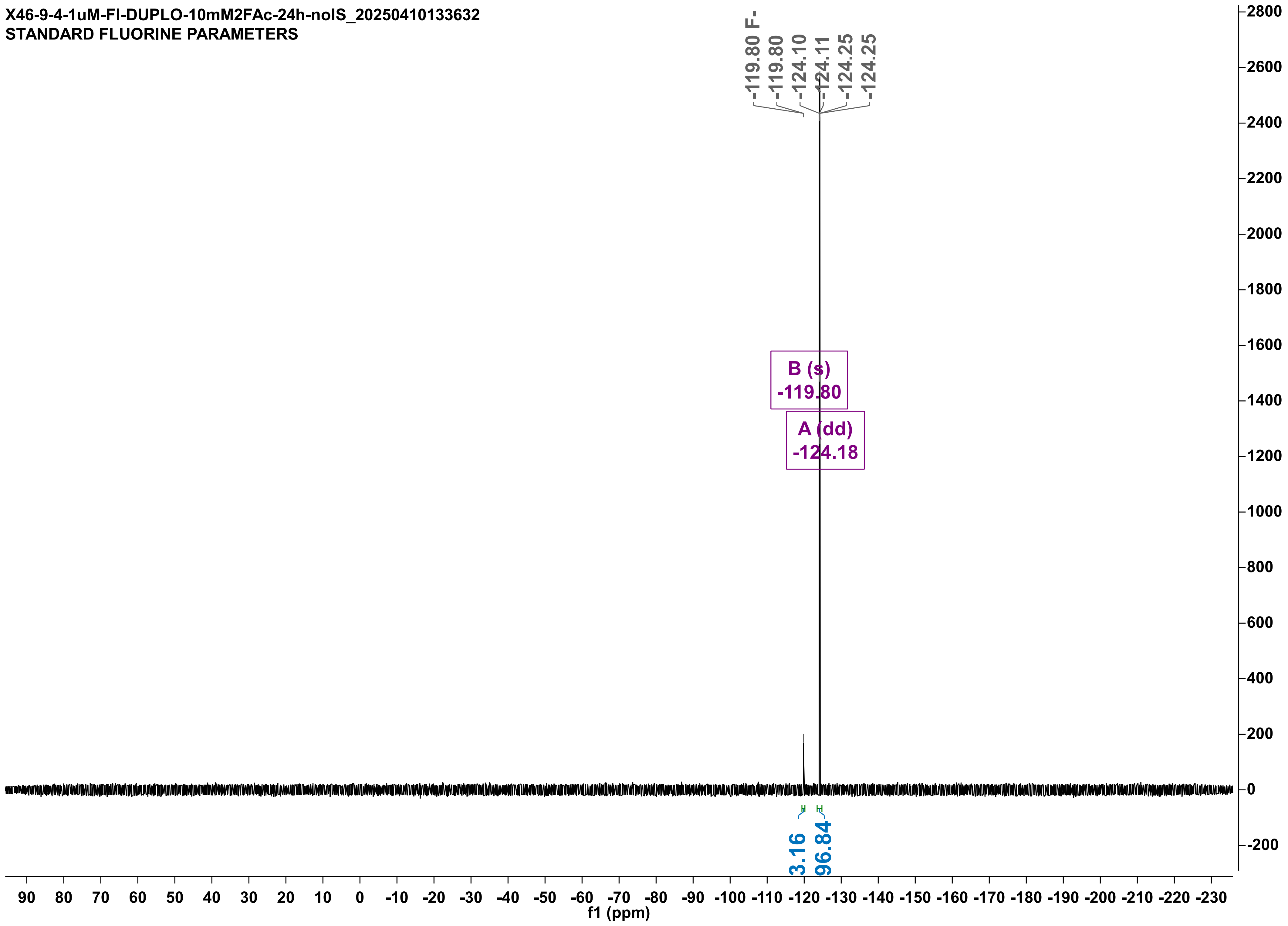
X46-9-4-1uM-FI-DUPLO-10mM2FAc-24h-noIS_20250410133632
STANDARD FLUORINE PARAMETERS
-119.80 F-
-119.80
-124.10
-124.11
-124.25
-124.25
B (s)
-119.80
A (dd)
-124.18
3.16
96.84
f1 (ppm)

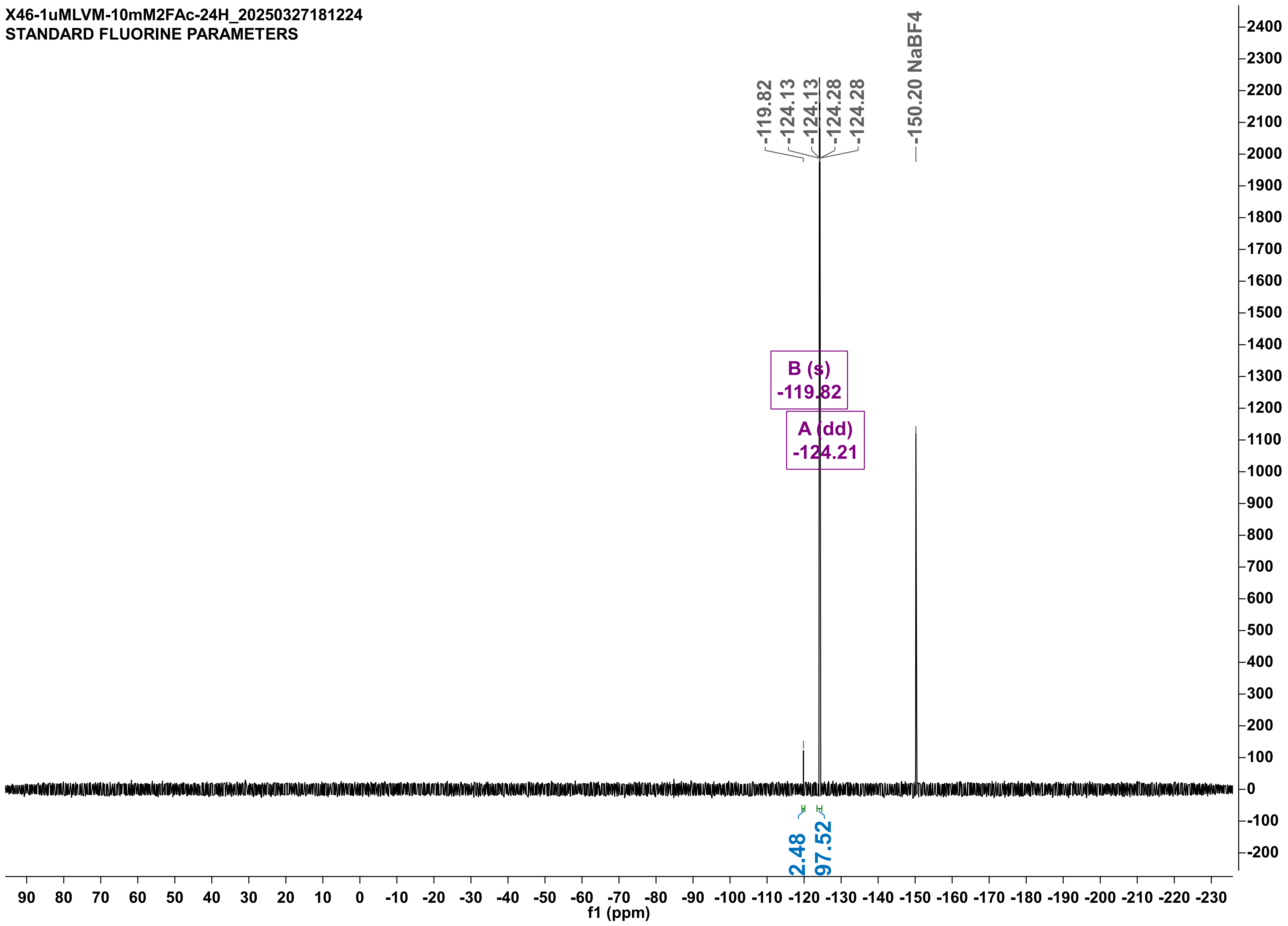
X46-1uMLVM-10mM2FAc-24H_20250327181224
STANDARD FLUORINE PARAMETERS
-119.82
-124.13
-124.13
-124.28
-124.28
-150.20 NaBF4
B (s)
-119.82
A (dd)
-124.21
2.48
97.52
f1 (ppm)

X46-2-1uMLVM-duplo-10mM2FAc-24H-noIS_20250328184256
STANDARD FLUORINE PARAMETERS

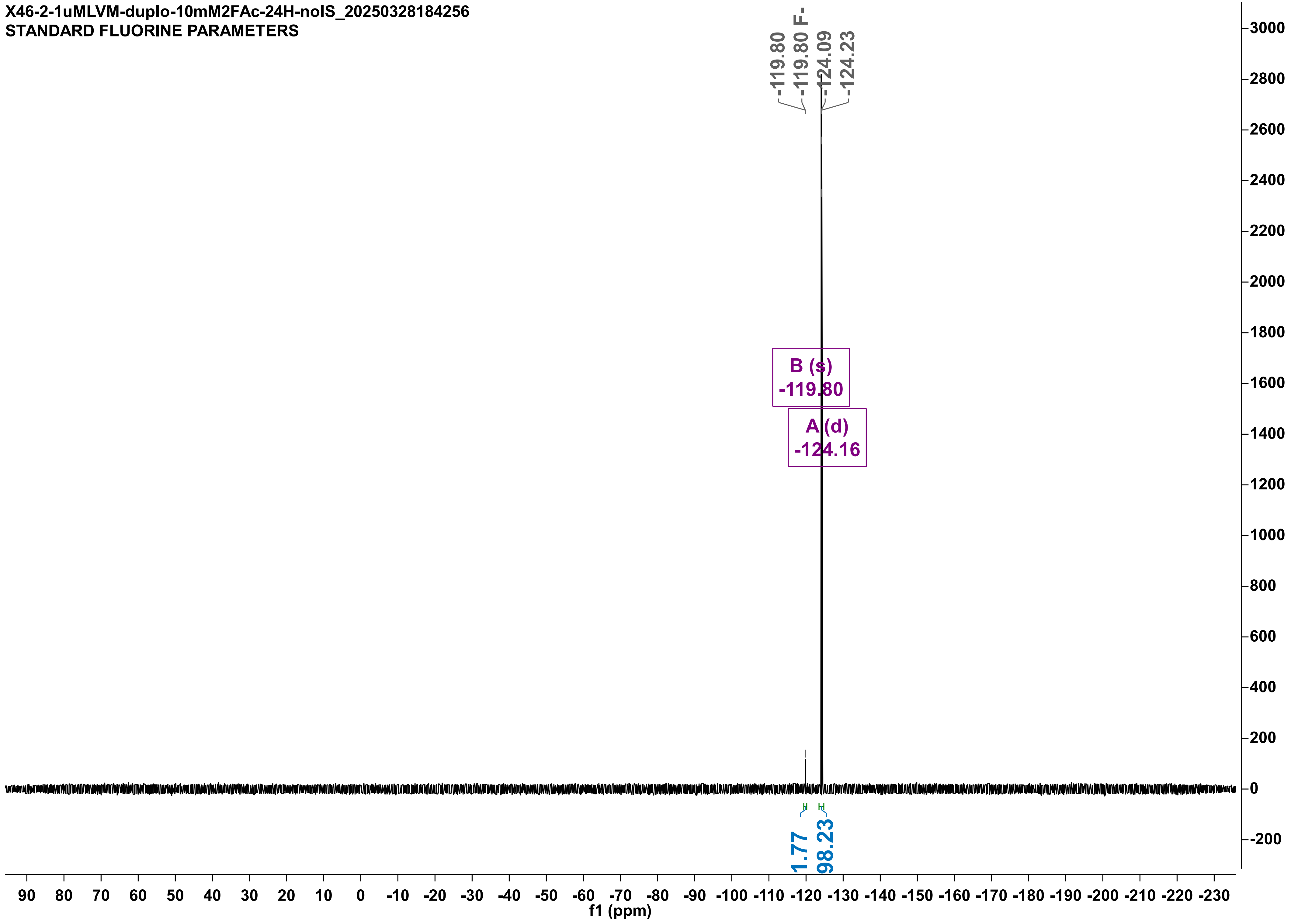

X46-1uMQ245A-10mMF2Ac-24H_20250326185253
STANDARD FLUORINE PARAMETERS

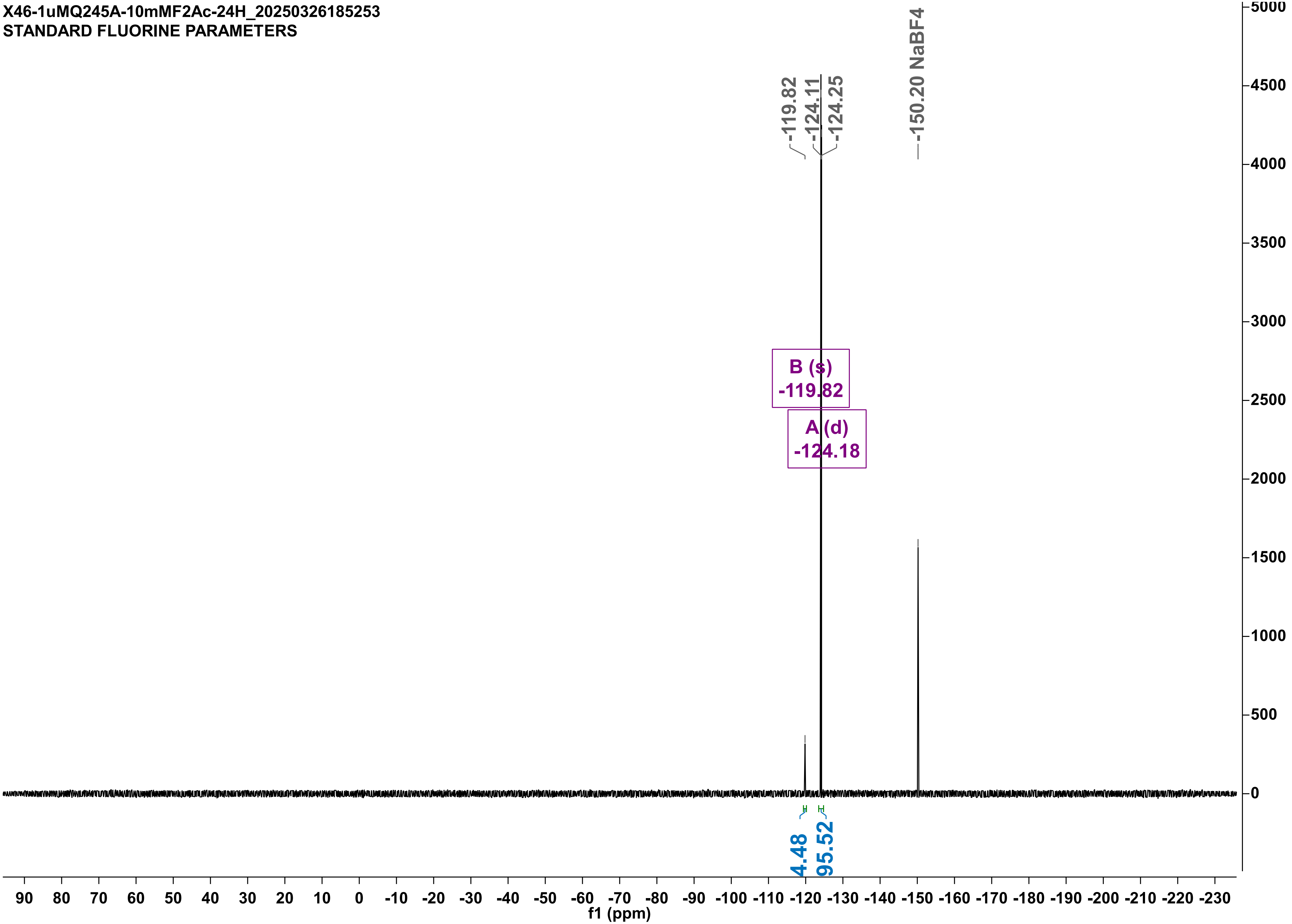

X46-9-4-1uMQ245A-DUPLO-10mM2FAc-24h-noIS_20250410133640
STANDARD FLUORINE PARAMETERS

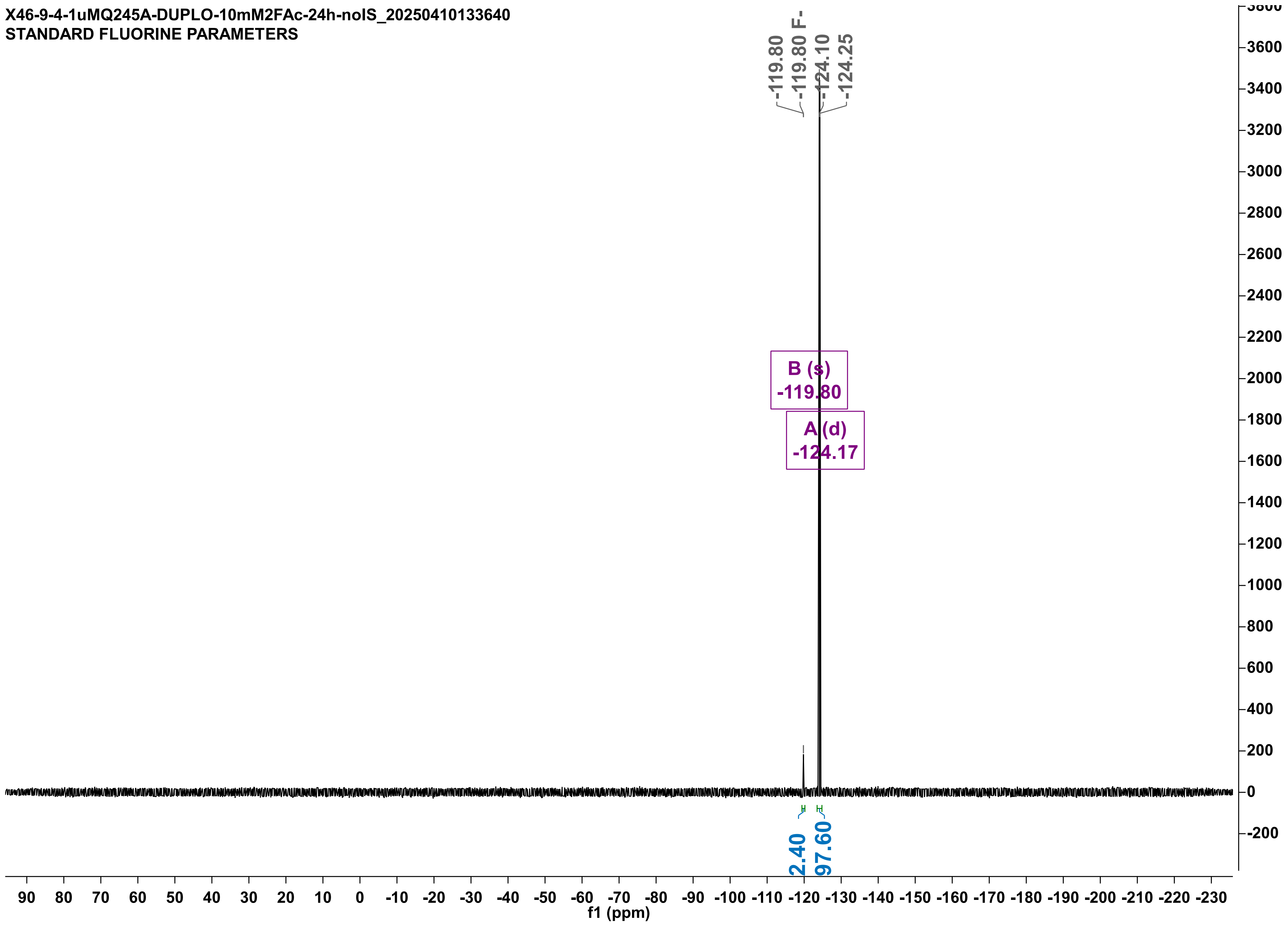

X46-1uMQ245G-10mMF2Ac-24H_20250326191725
STANDARD FLUORINE PARAMETERS

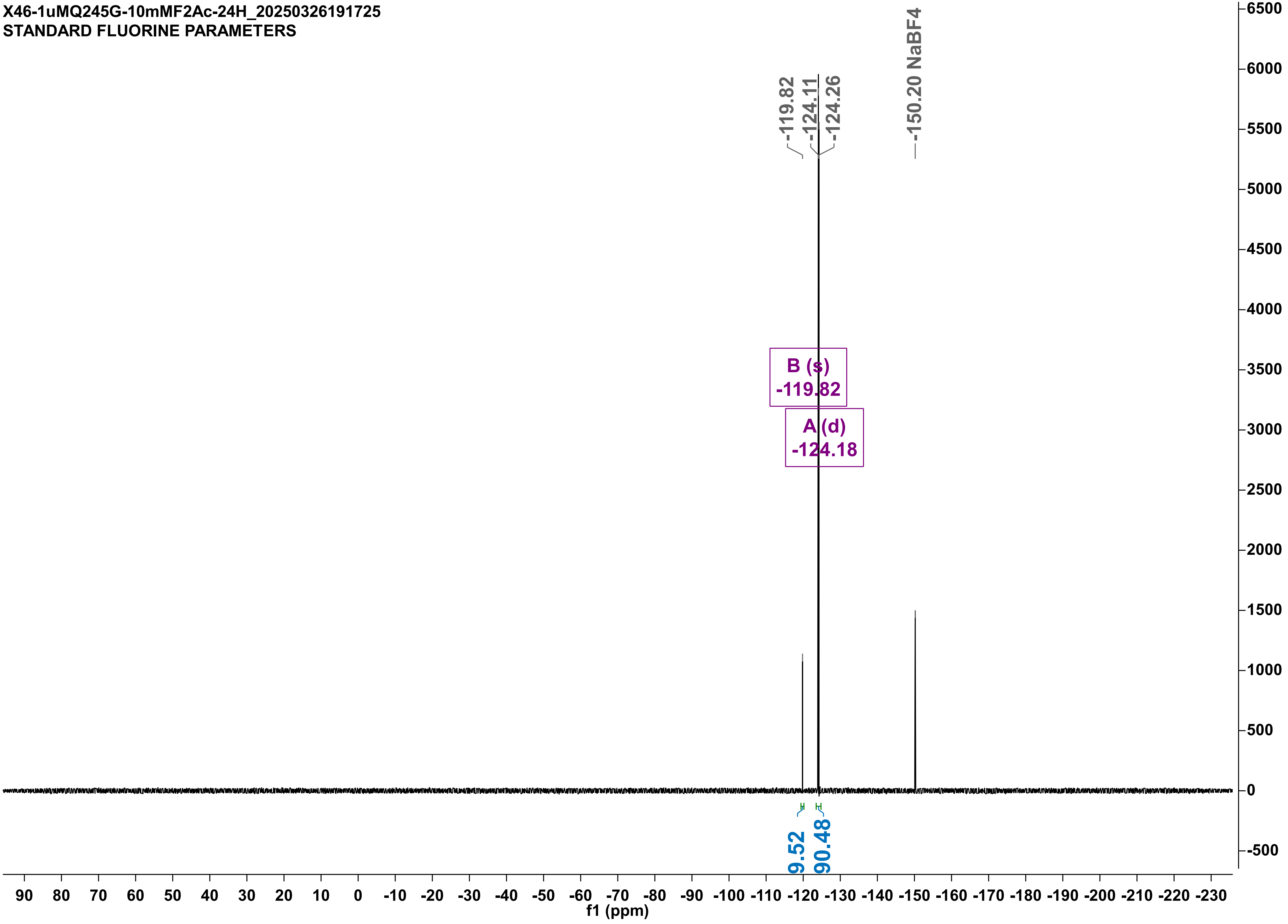

X46-9-4-1uMQ245G-DUPLO-10mM2FAc-24h-noIS_20250410133648
STANDARD FLUORINE PARAMETERS

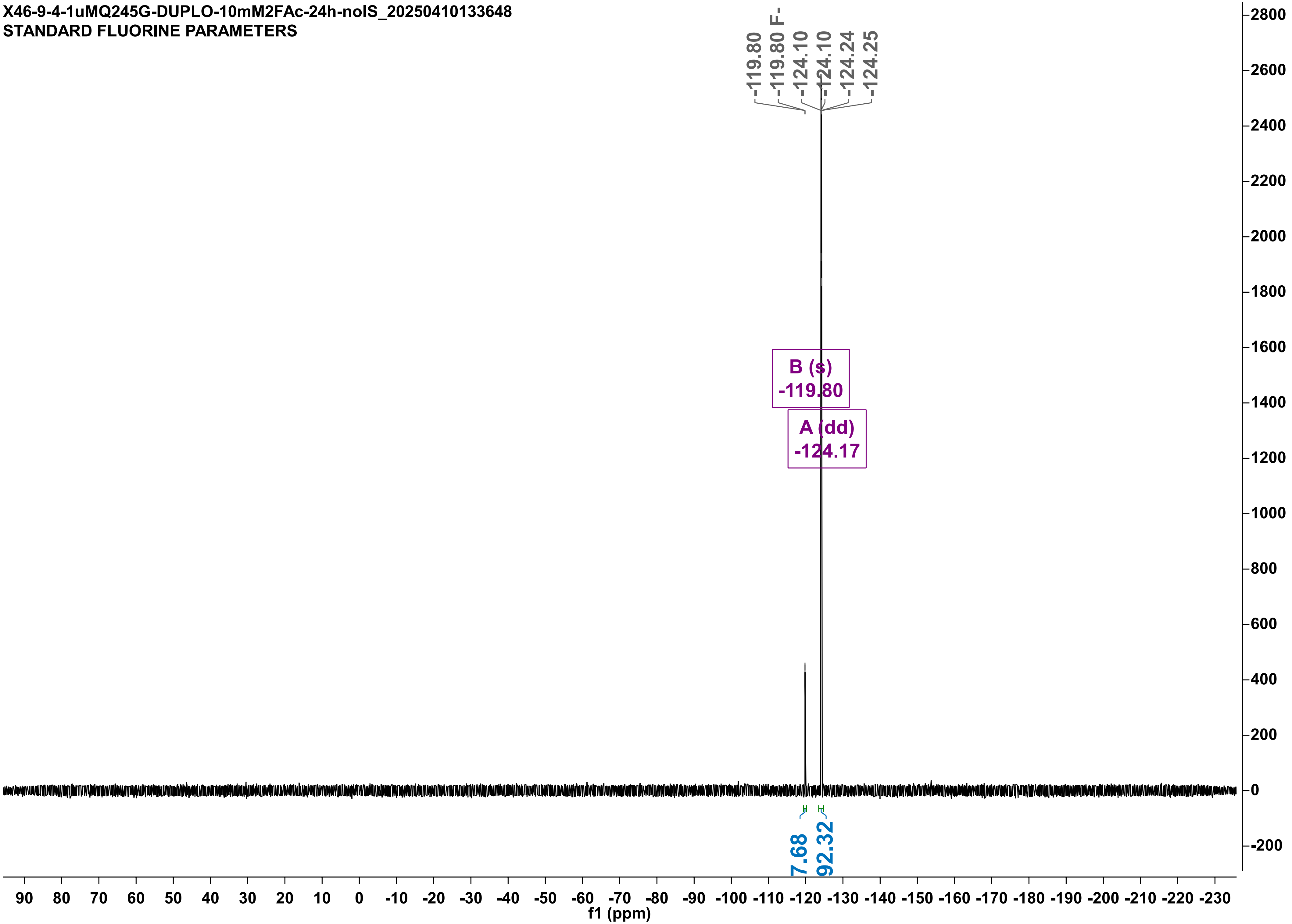

X46-1uMQ245R-10mMF2Ac-24H_20250326191736
STANDARD FLUORINE PARAMETERS

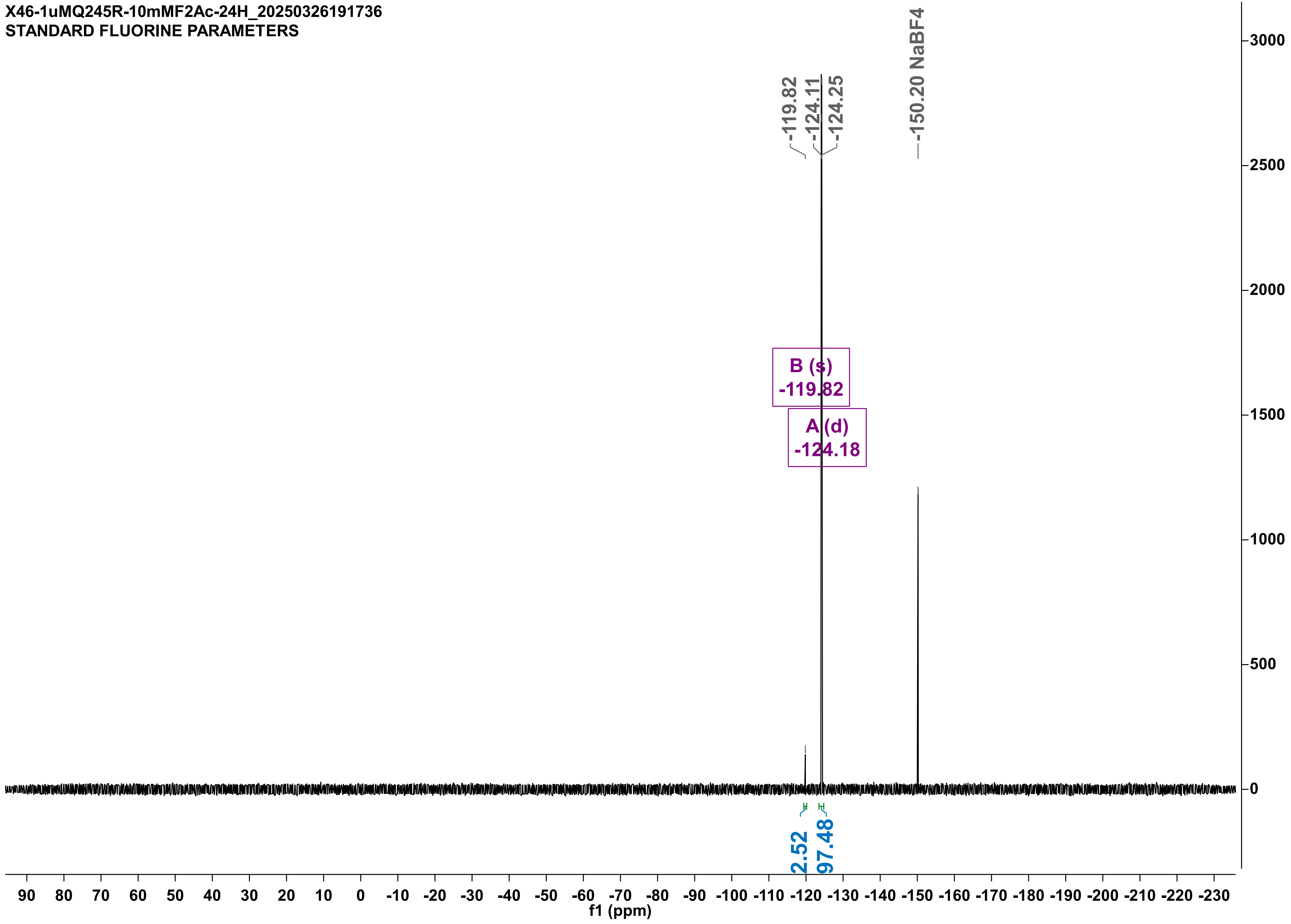

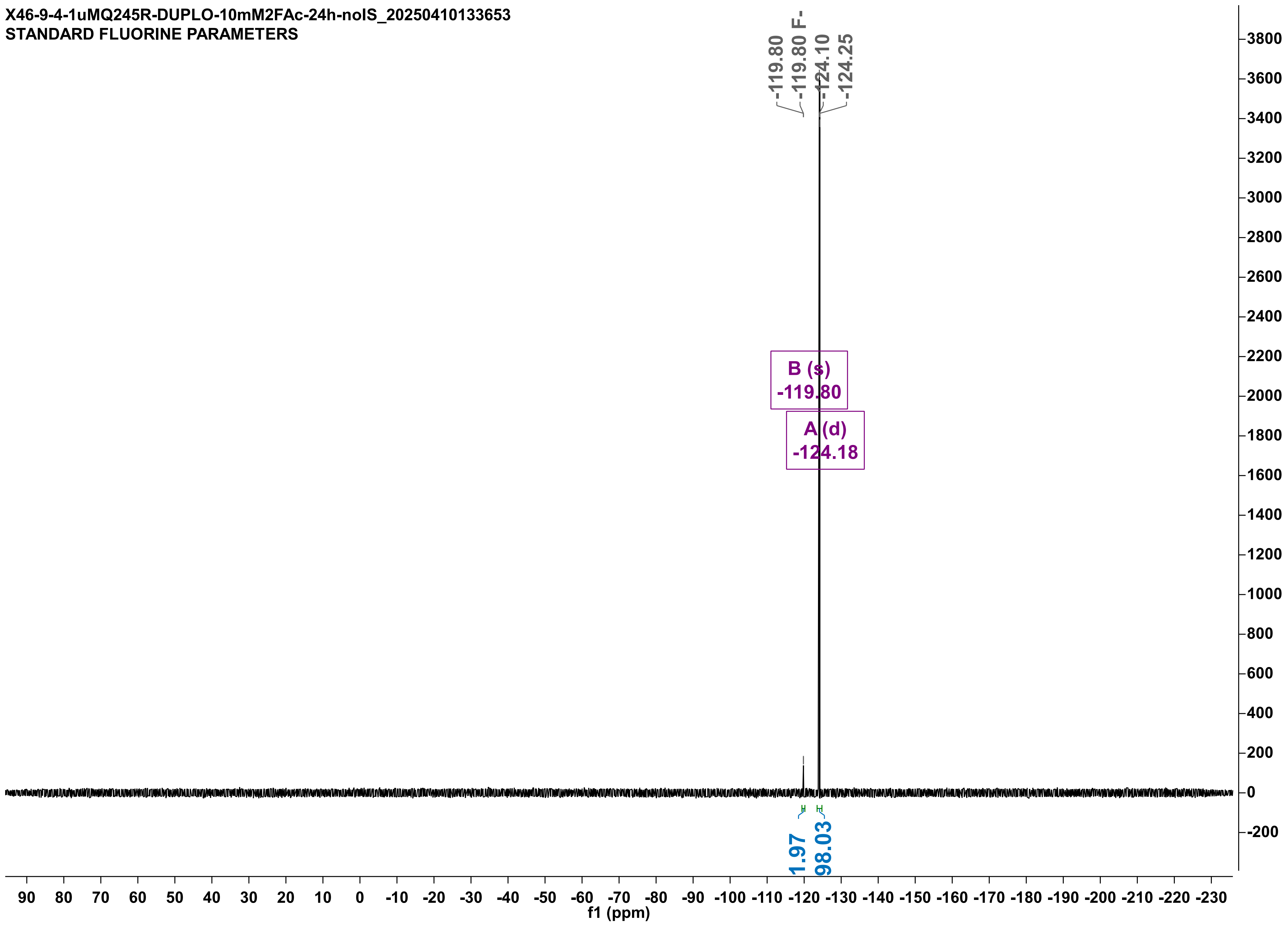
X46-9-4-1uMQ245R-DUPLO-10mM2FAc-24h-noIS_20250410133653
STANDARD FLUORINE PARAMETERS
-119.80
-119.80 F-
-124.10
-124.25
B (s)
-119.80
A (d)
-124.18
1.97
98.03
f1 (ppm)

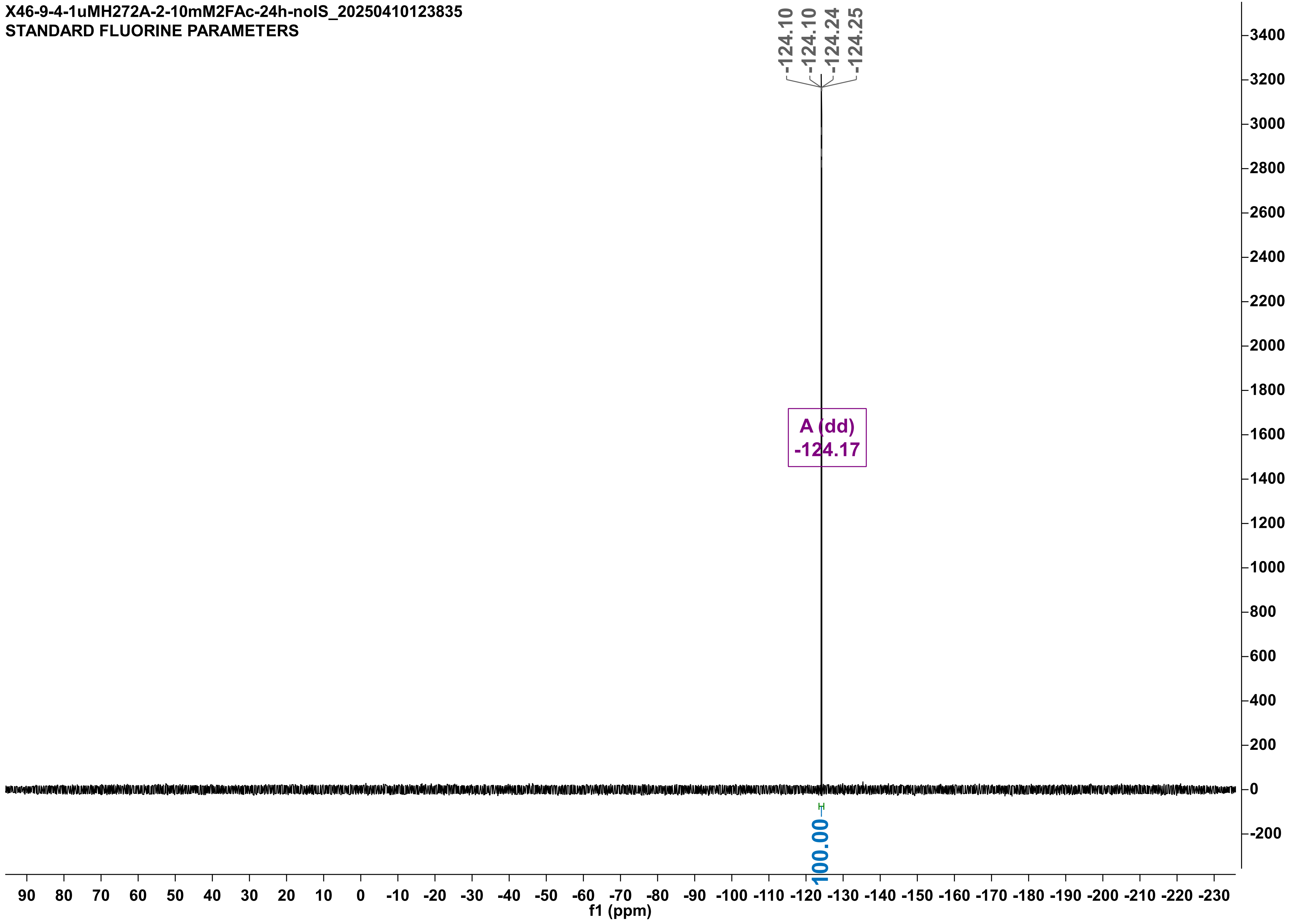

X46-9-4-1uMH272A-2-10mM2FAc-24h-noIS_20250410123835
STANDARD FLUORINE PARAMETERS
-124.10
-124.10
-124.24
-124.25
A (dd)
-124.17
100.00
f1 (ppm)

X46-9-4-noenz-10mM2FAc-24h-noIS_20250410123746
STANDARD FLUORINE PARAMETERS

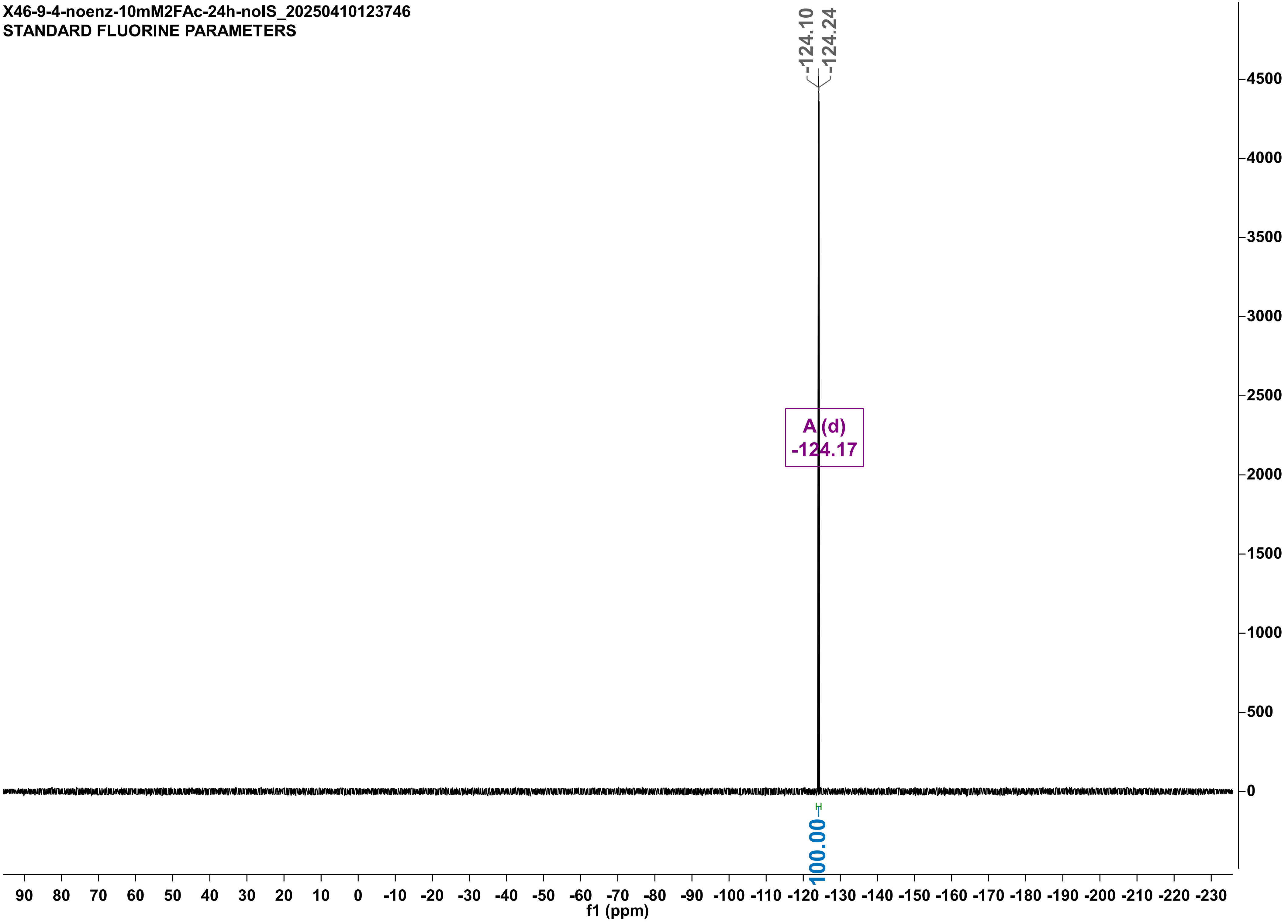

X46-1uMH272A-duplo-10mM2FAc-24h_20250327181949
STANDARD FLUORINE PARAMETERS

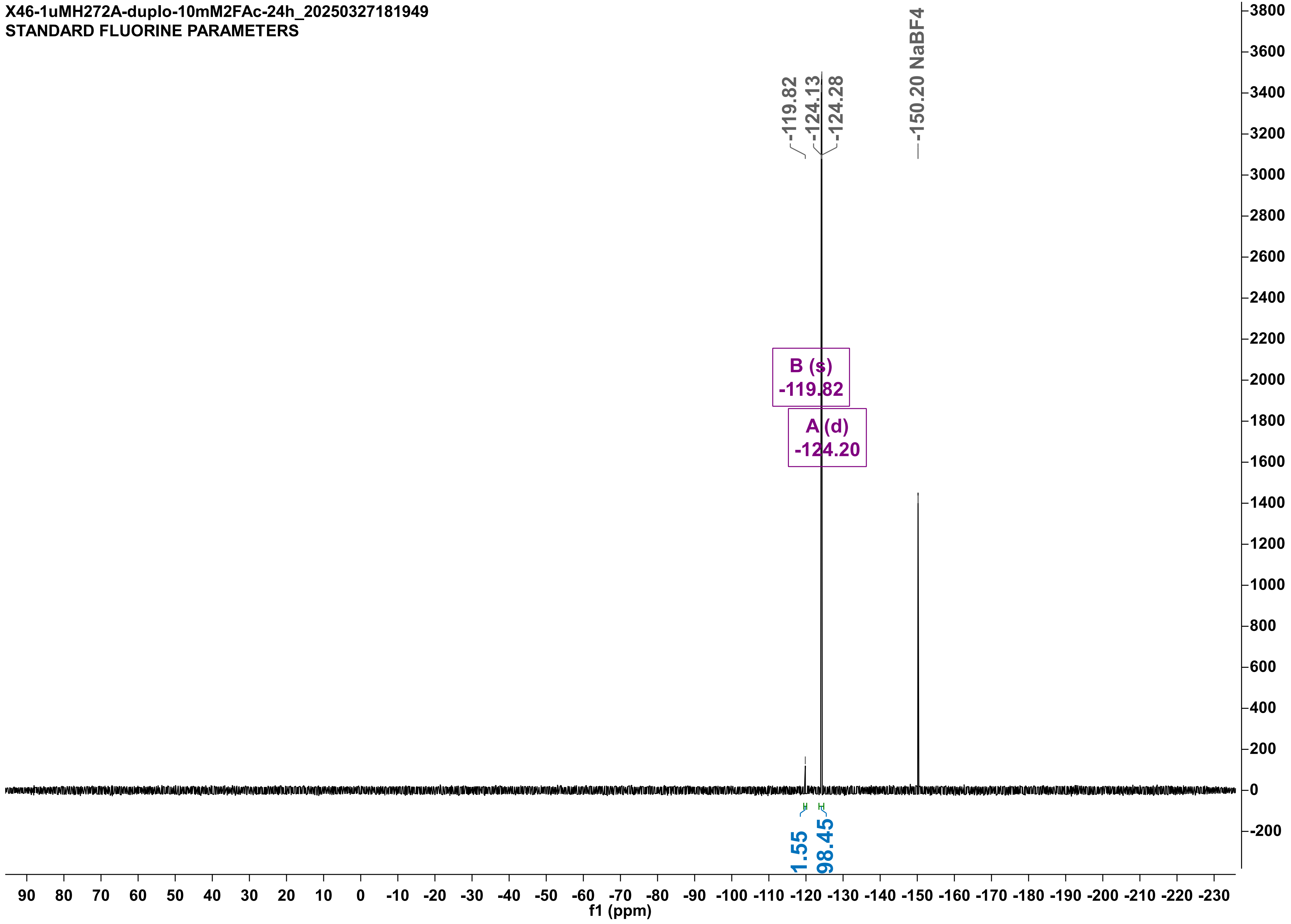

X49-17-4-NOENZ-10mM2FPA-time24h_20250417143022
STANDARD FLUORINE PARAMETERS

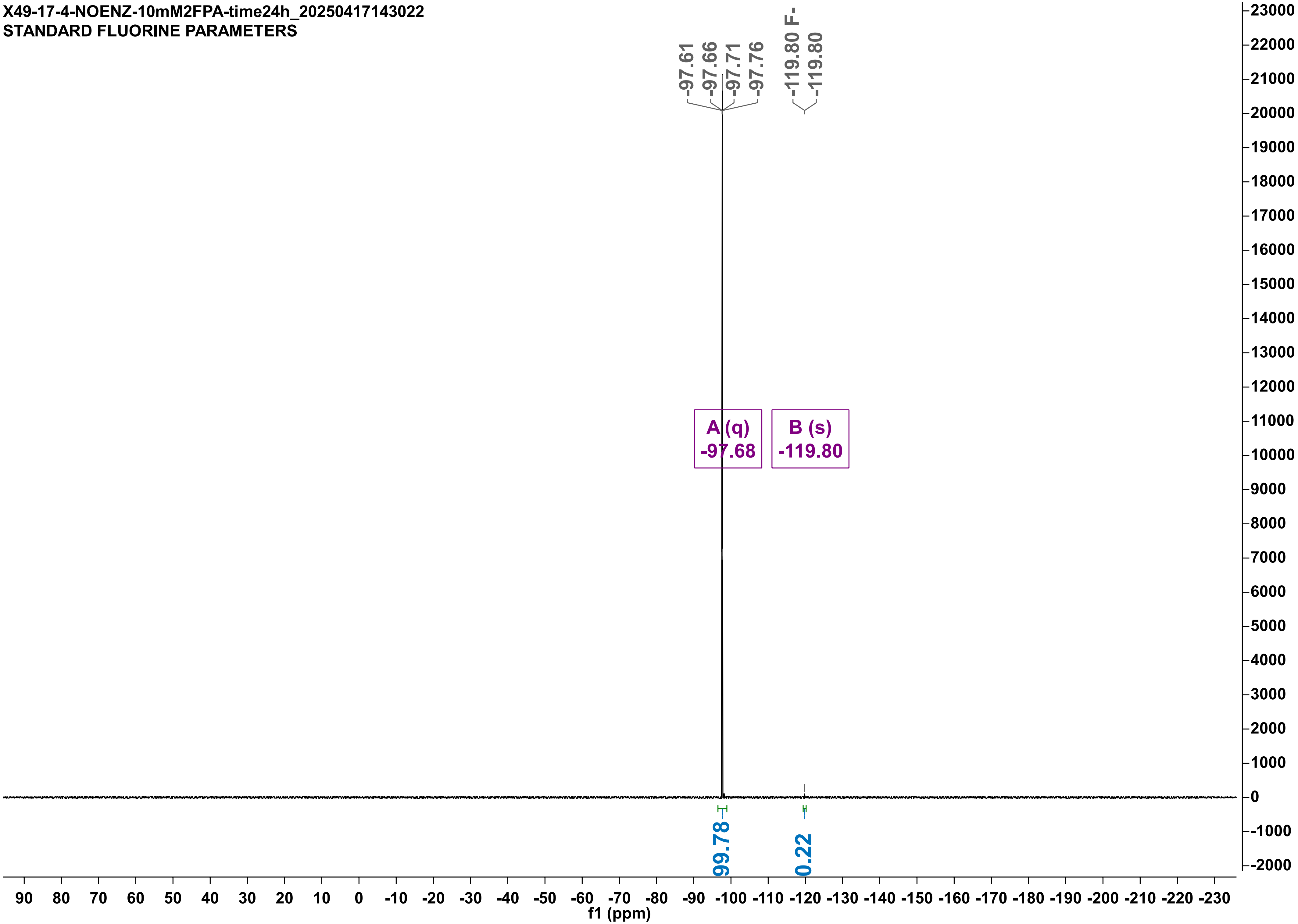

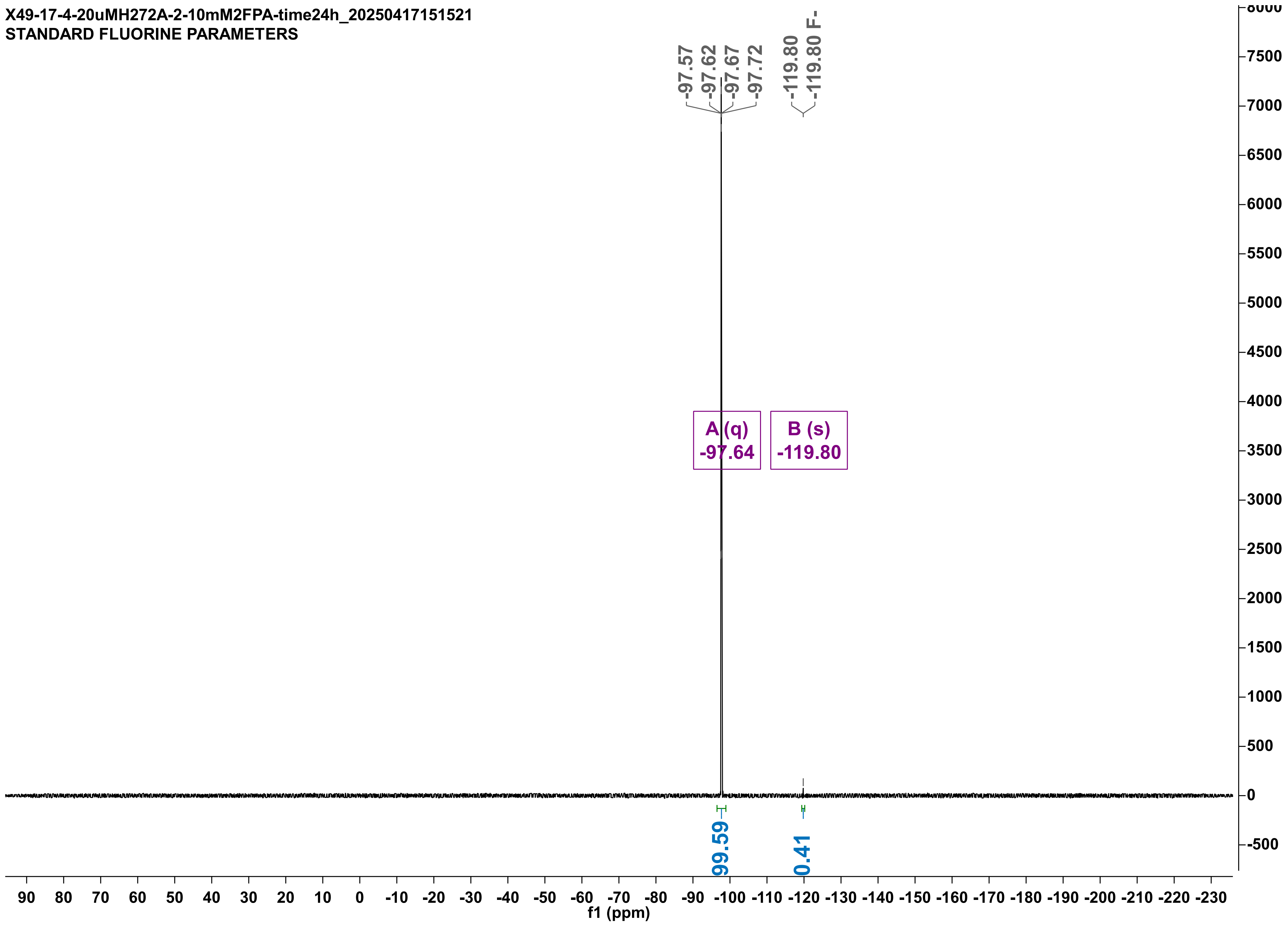
X49-17-4-20uMH272A-2-10mM2FPA-time24h_20250417151521
STANDARD FLUORINE PARAMETERS
-97.57
-97.62
-97.67
-97.72
-119.80
-119.80 F-
A (q)
-97.64
B (s)
-119.80
99.59
0.41
f1 (ppm)

X49-17-4-20uMWT1-10mM2FPA-time24h_20250417151701
STANDARD FLUORINE PARAMETERS

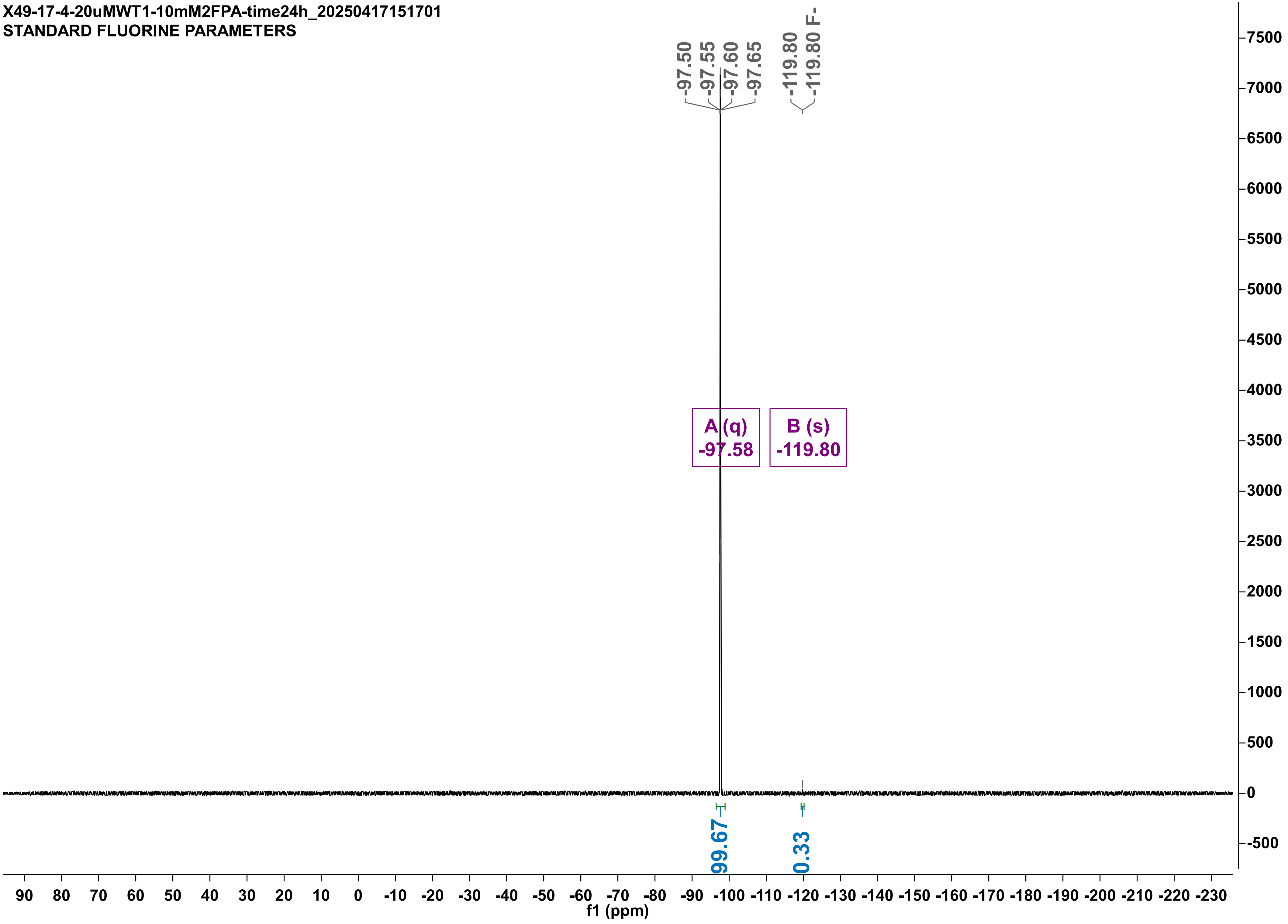

X49-17-4-20uMWT2-10mM2FPA-time24h_20250417161746
STANDARD FLUORINE PARAMETERS

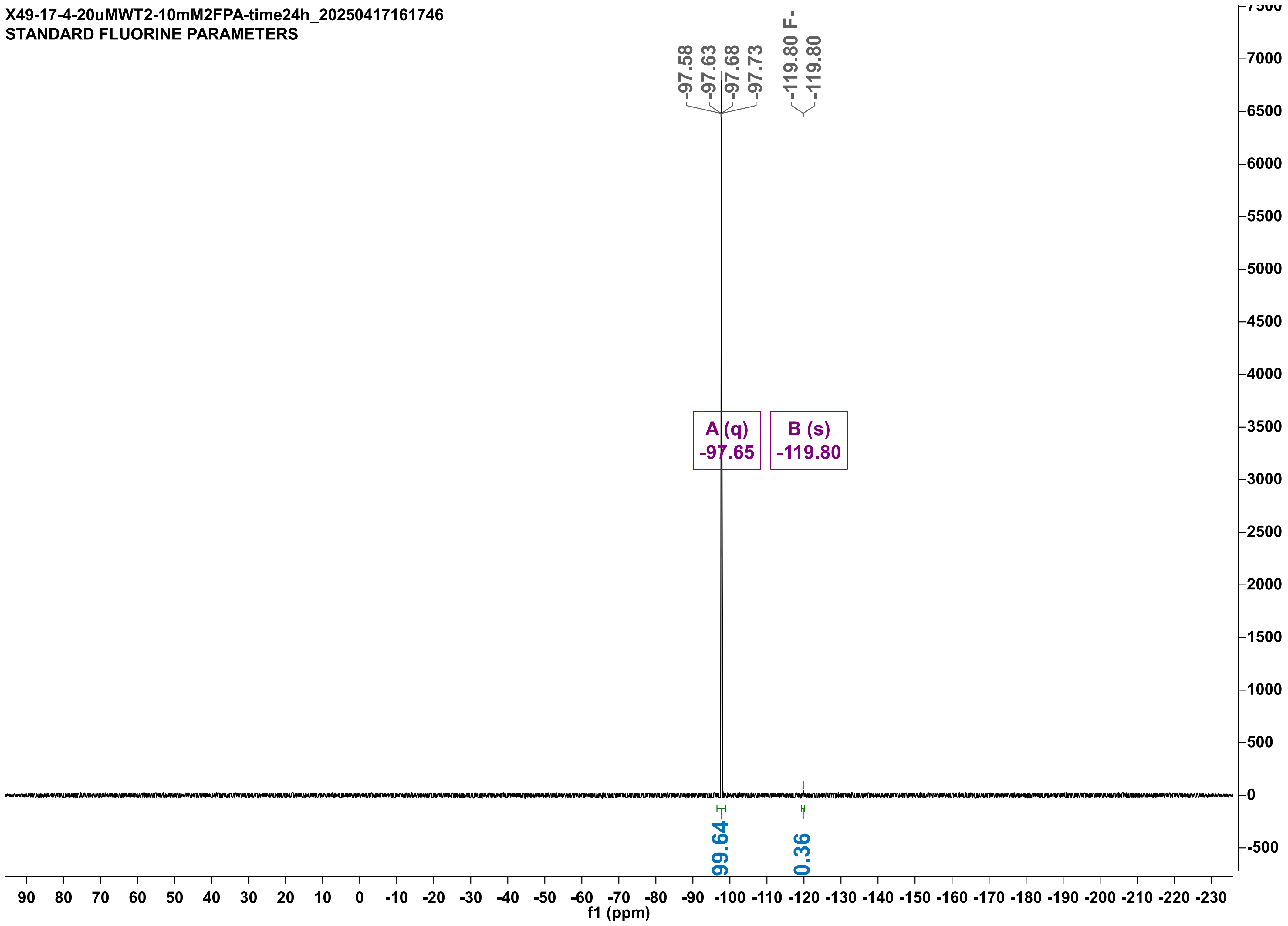

X46-20uMVW--10mMF2PA-24h_20250319165040
STANDARD FLUORINE PARAMETERS

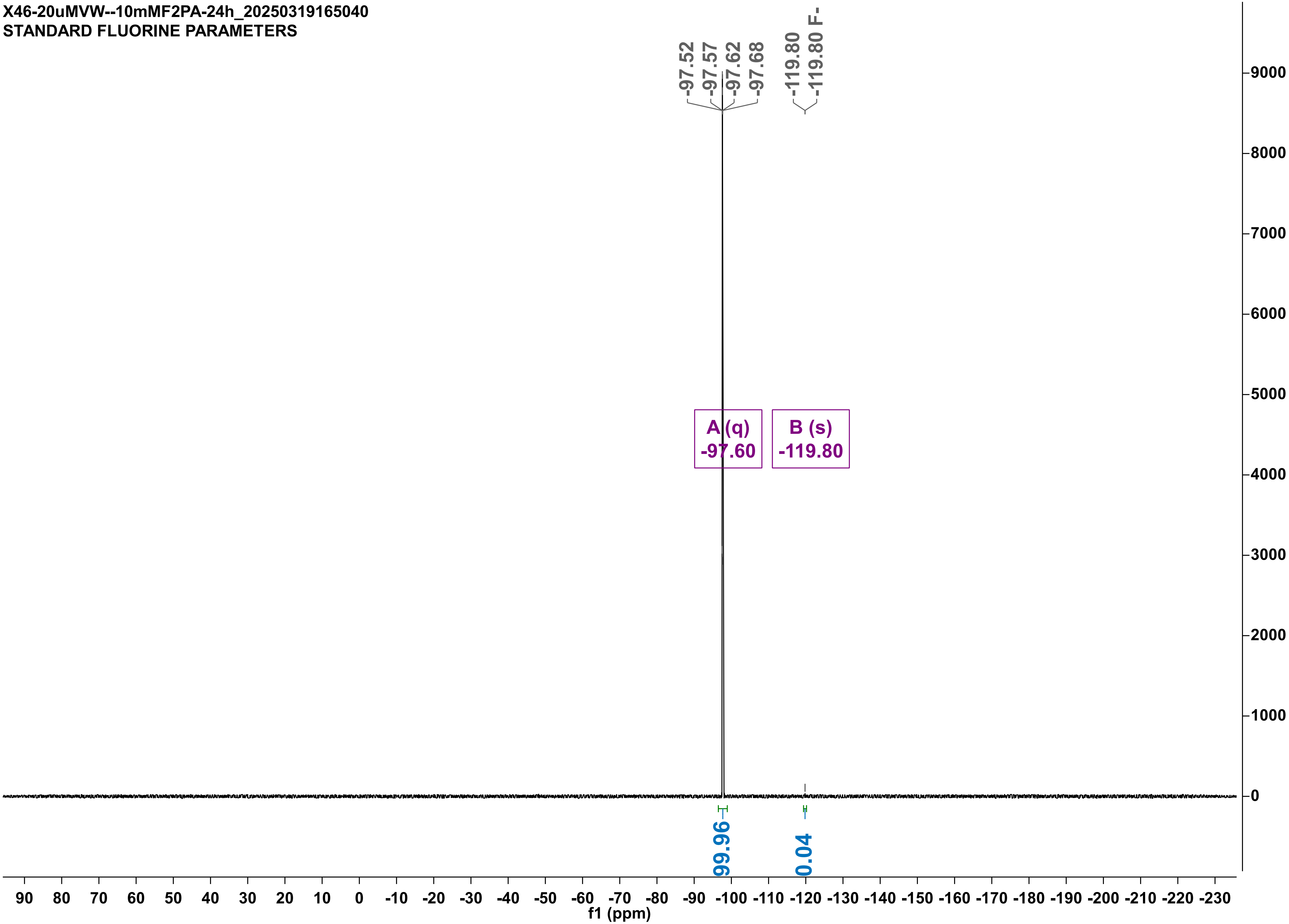

X49-17-4-20uMLW--10mM2FPA-time24h_20250417151555
STANDARD FLUORINE PARAMETERS

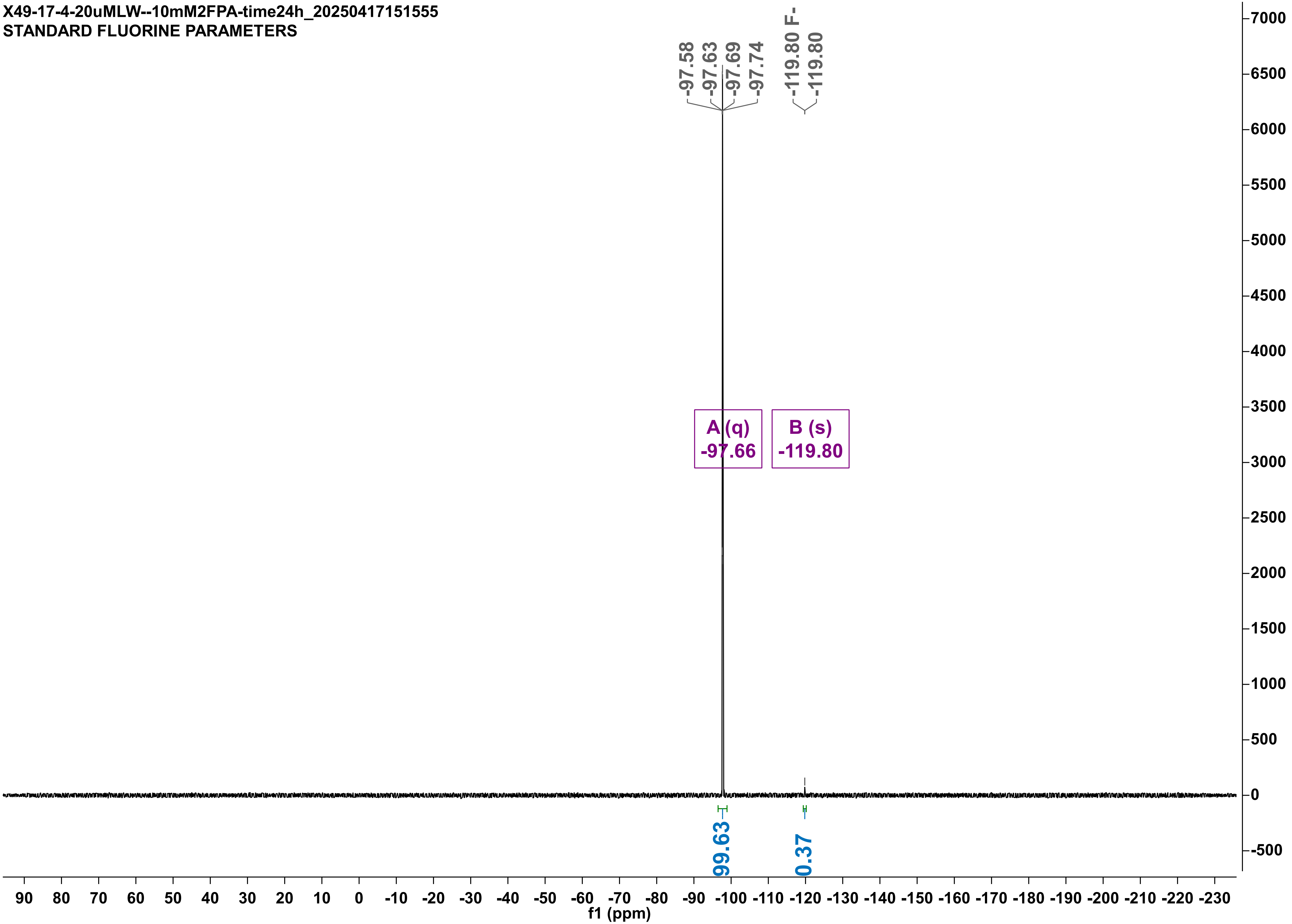

X49-17-4-20uM-ML-10mM2FPA-time24h_20250417151612
STANDARD FLUORINE PARAMETERS

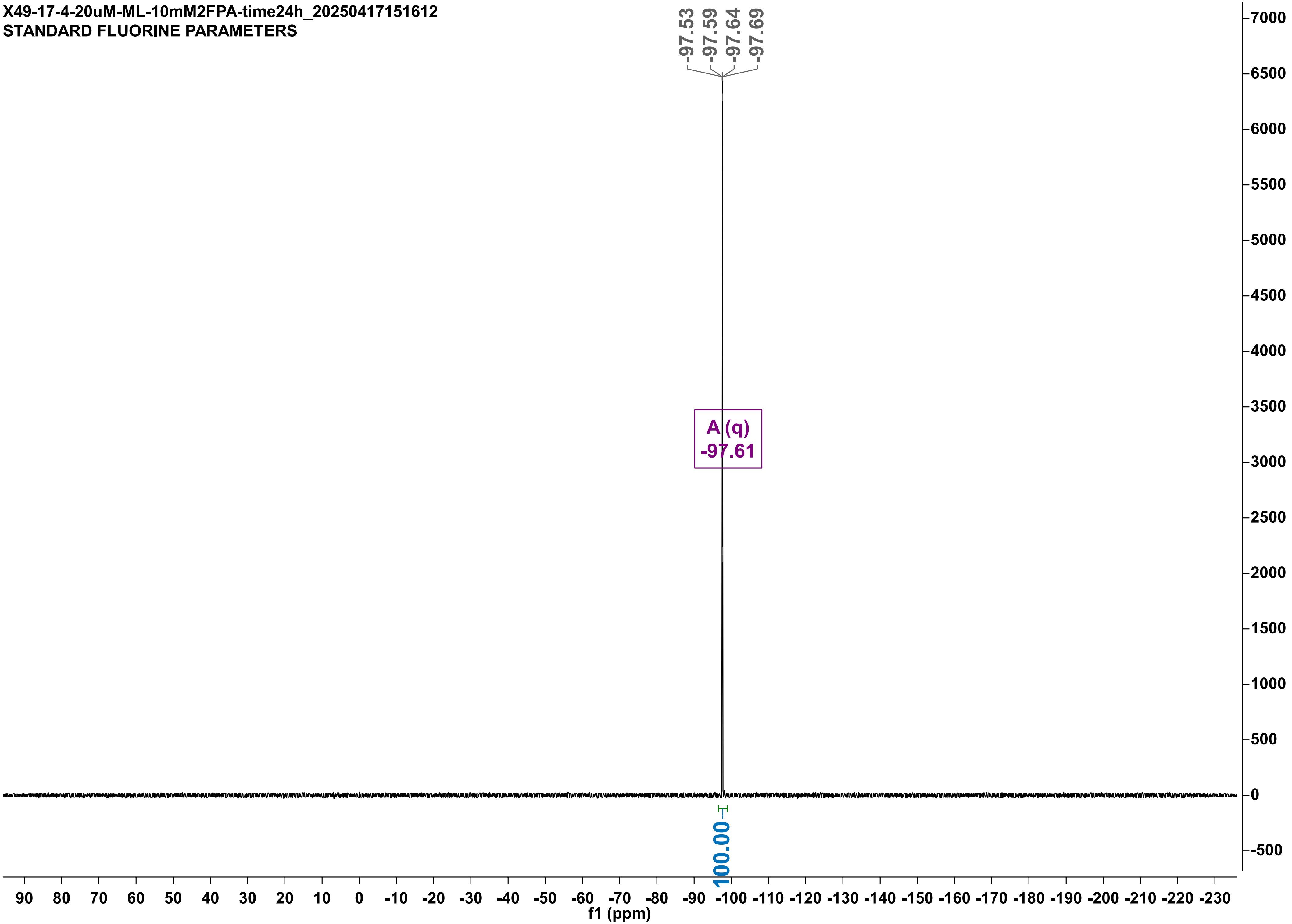

X49-17-4-20uMPLM-2-10mM2FPA-time24h_20250417151629
STANDARD FLUORINE PARAMETERS

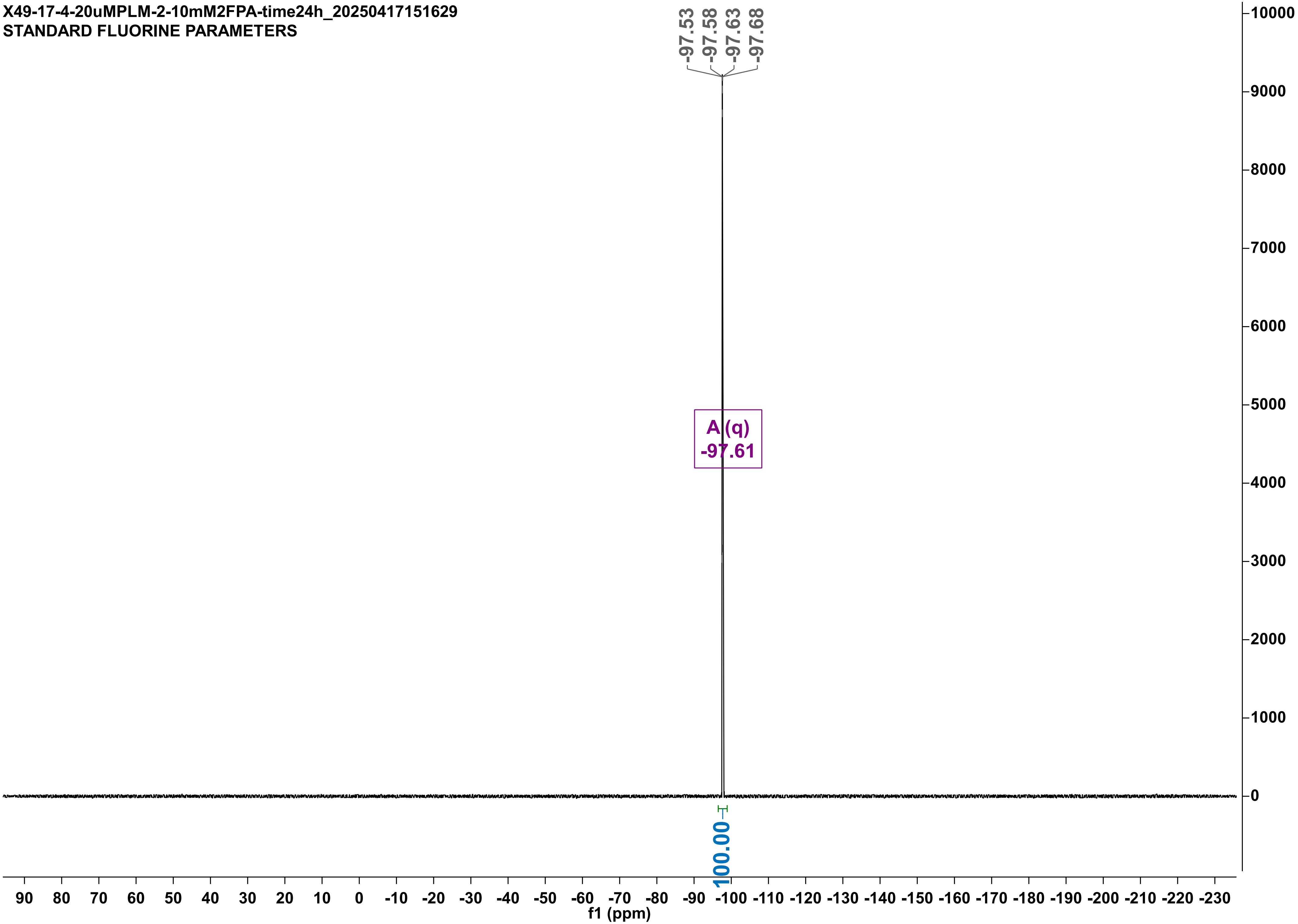

X49-17-4-20uMRHL-10mM2FPA-time24h_20250417161927
STANDARD FLUORINE PARAMETERS

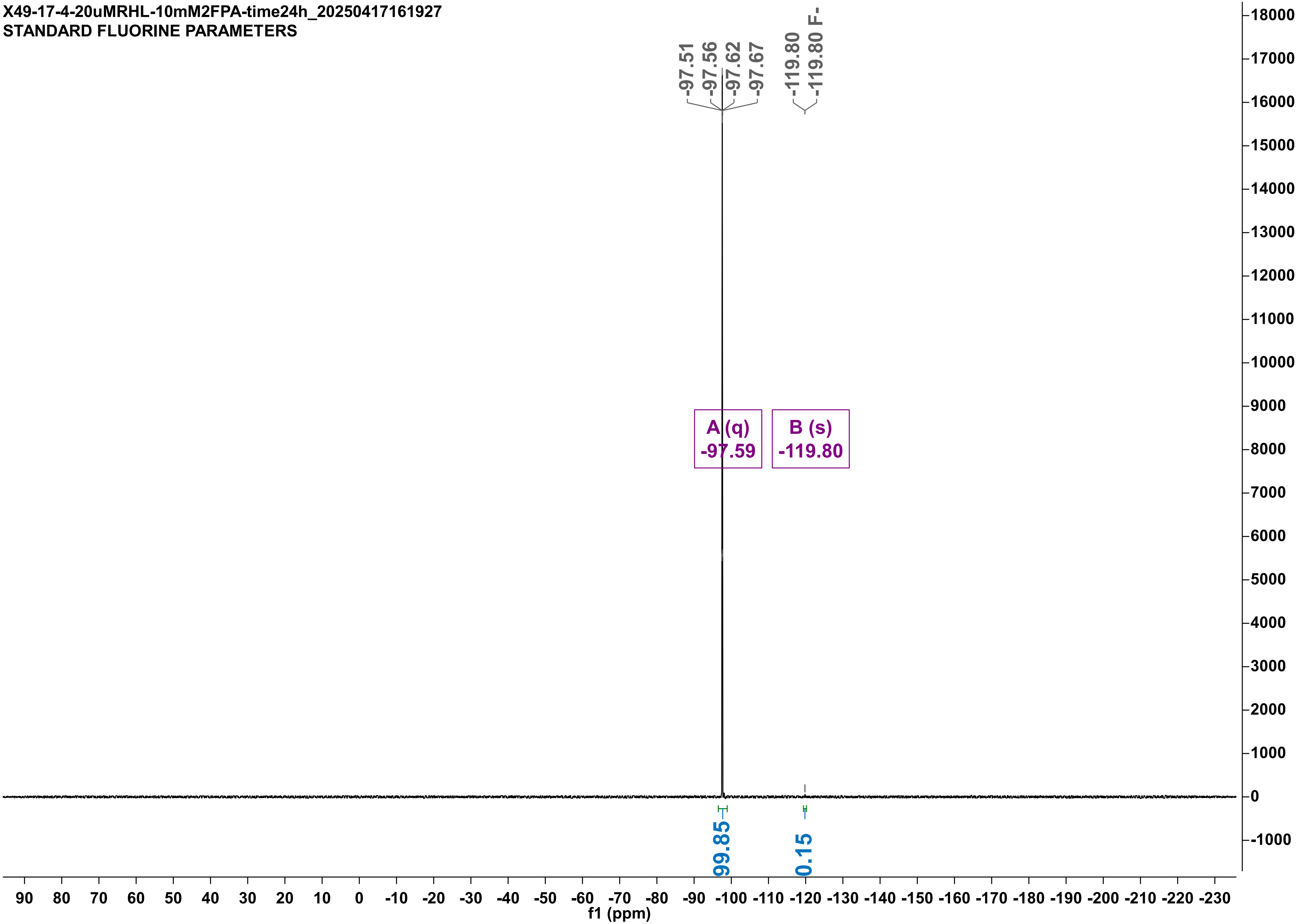

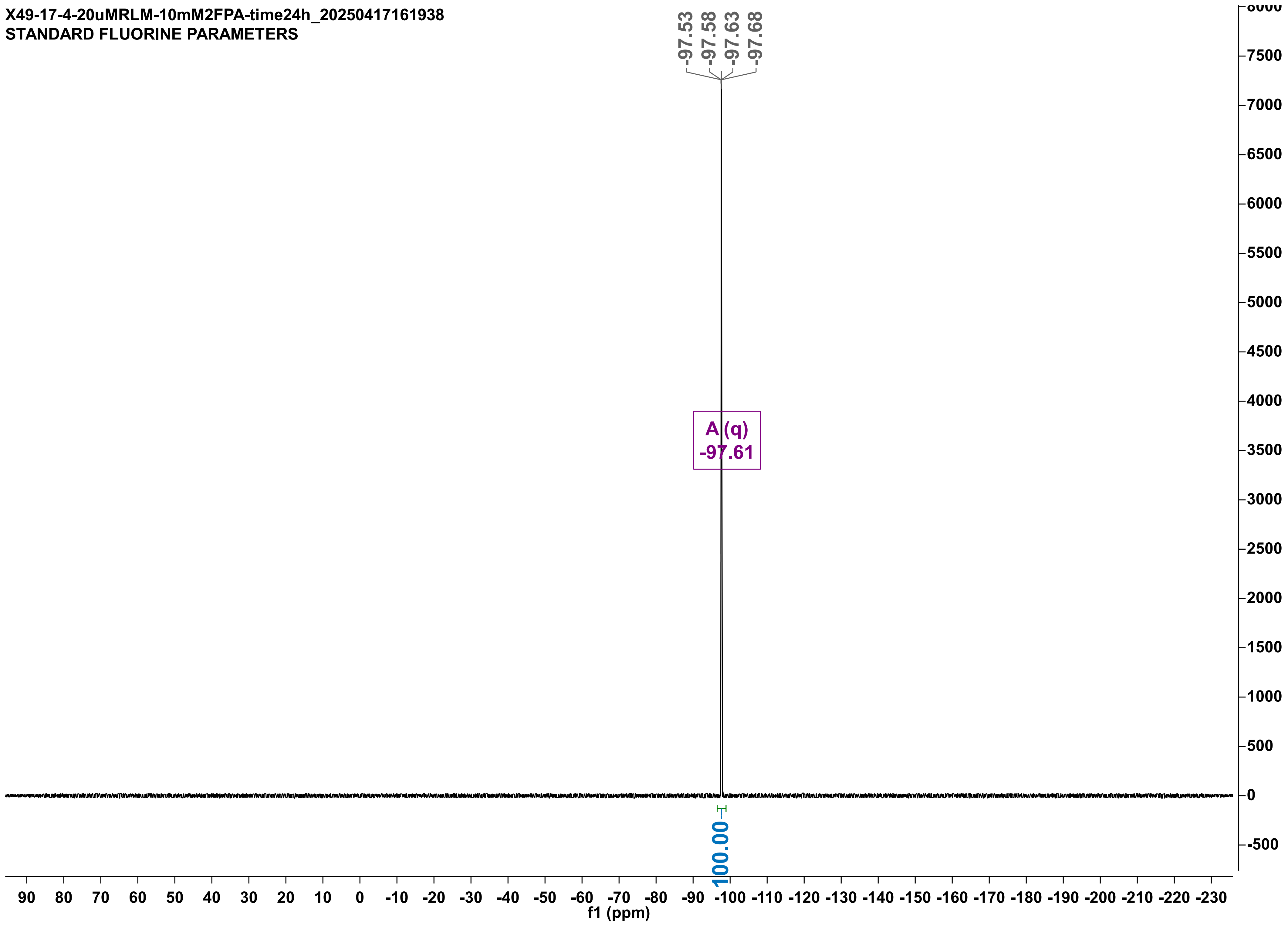

X49-17-4-20uMRLM-10mM2FPA-time24h_20250417161938
STANDARD FLUORINE PARAMETERS
-97.53
-97.58
-97.63
-97.68
A (q)
-97.61
100.00
90 80 70 60 50 40 30 20 10 0 -10 -20 -30 -40 -50 -60 -70 -80 -90 -100 -110 -120 -130 -140 -150 -160 -170 -180 -190 -200 -210 -220 -230
f1 (ppm)
8000 7500 7000 6500 6000 5500 5000 4500 4000 3500 3000 2500 2000 1500 1000 500 0 -500

X49-17-4-20uMRPL-10mM2FPA-time24h_20250417161907
STANDARD FLUORINE PARAMETERS

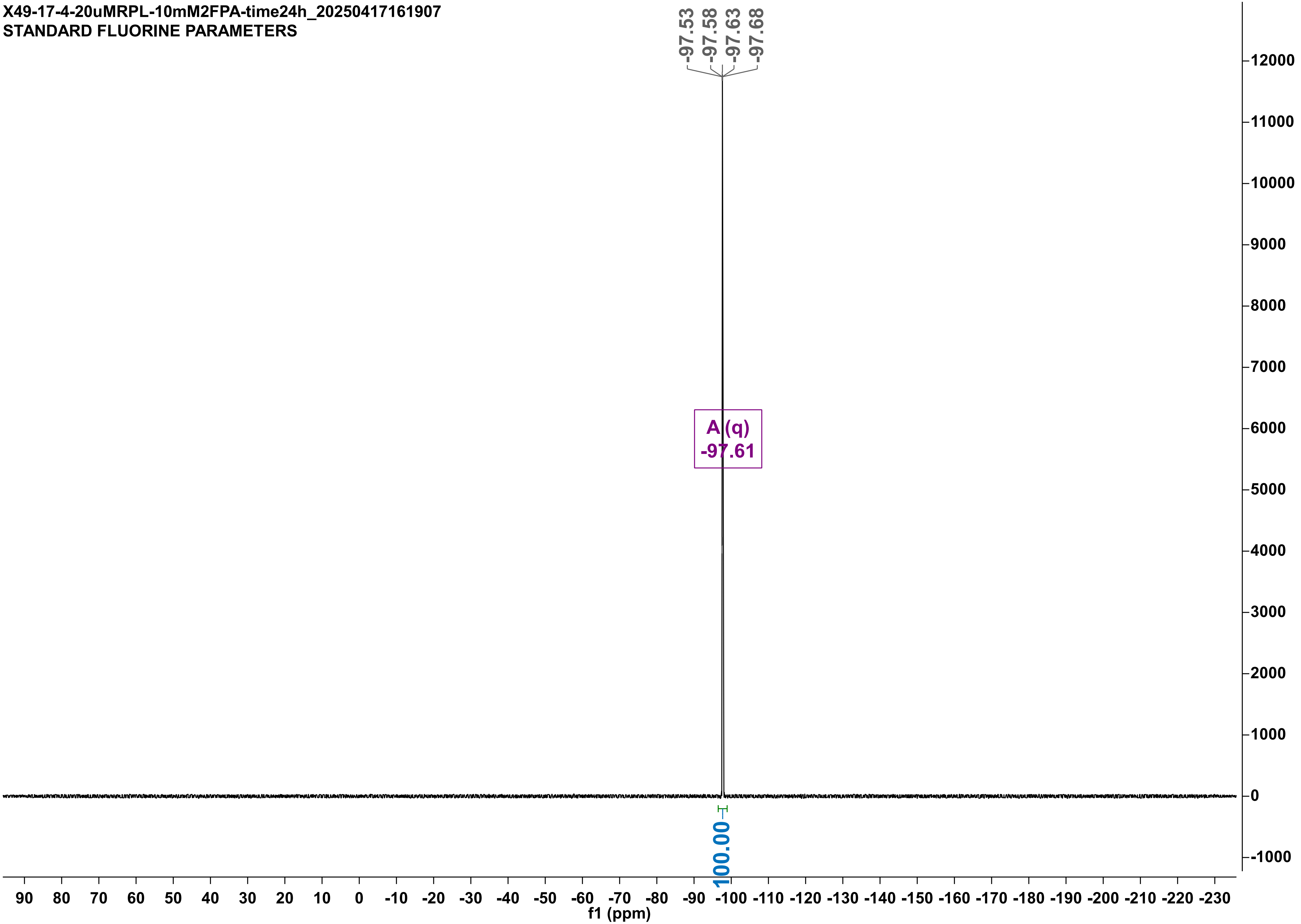

X49-30-5_20uM_-FI_10mM_2FPA_t24h_20250530133233
STANDARD FLUORINE PARAMETERS

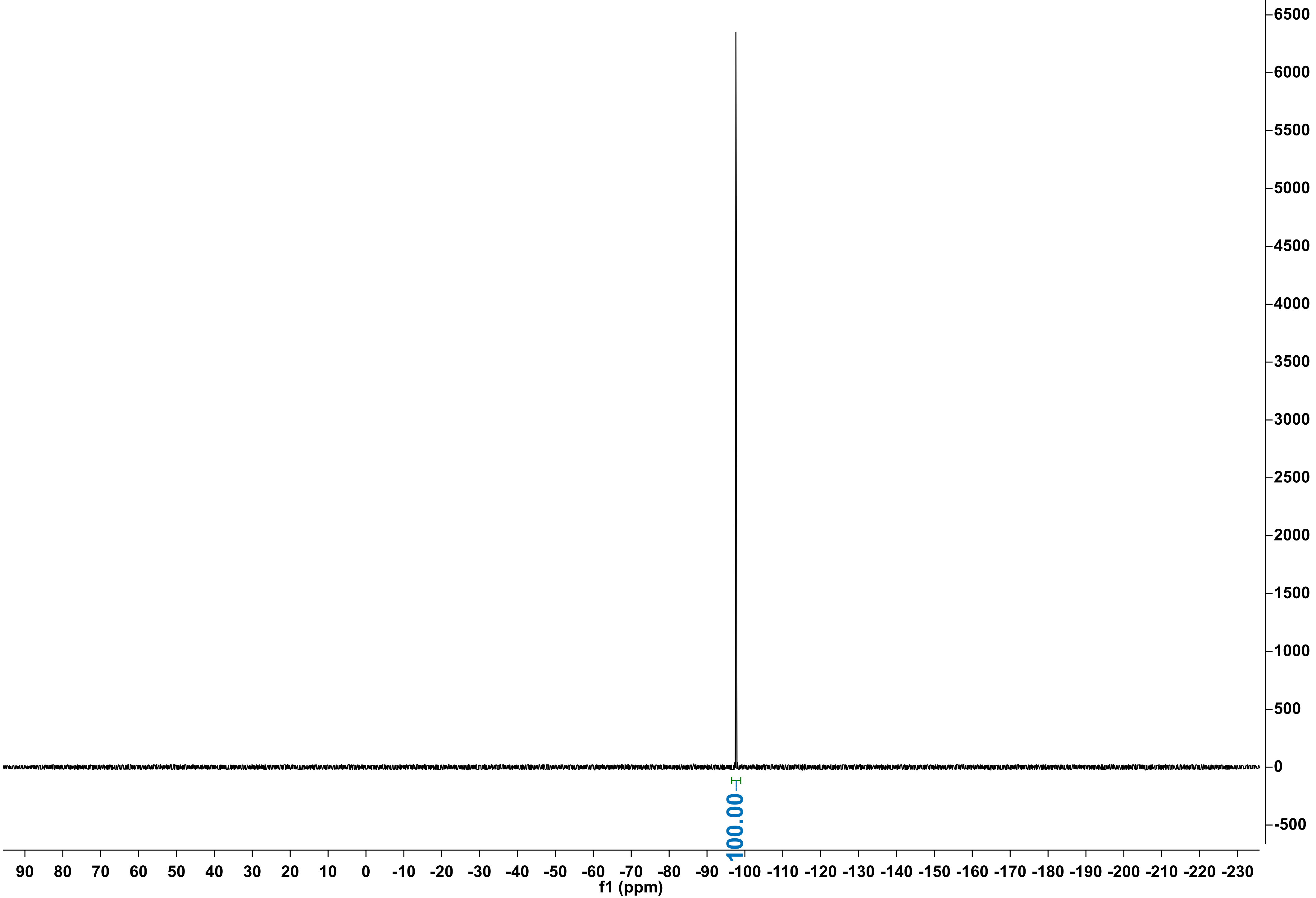

X49-30-5_20uM_LVM_10mM_2FPA_t24h_20250530124414
STANDARD FLUORINE PARAMETERS

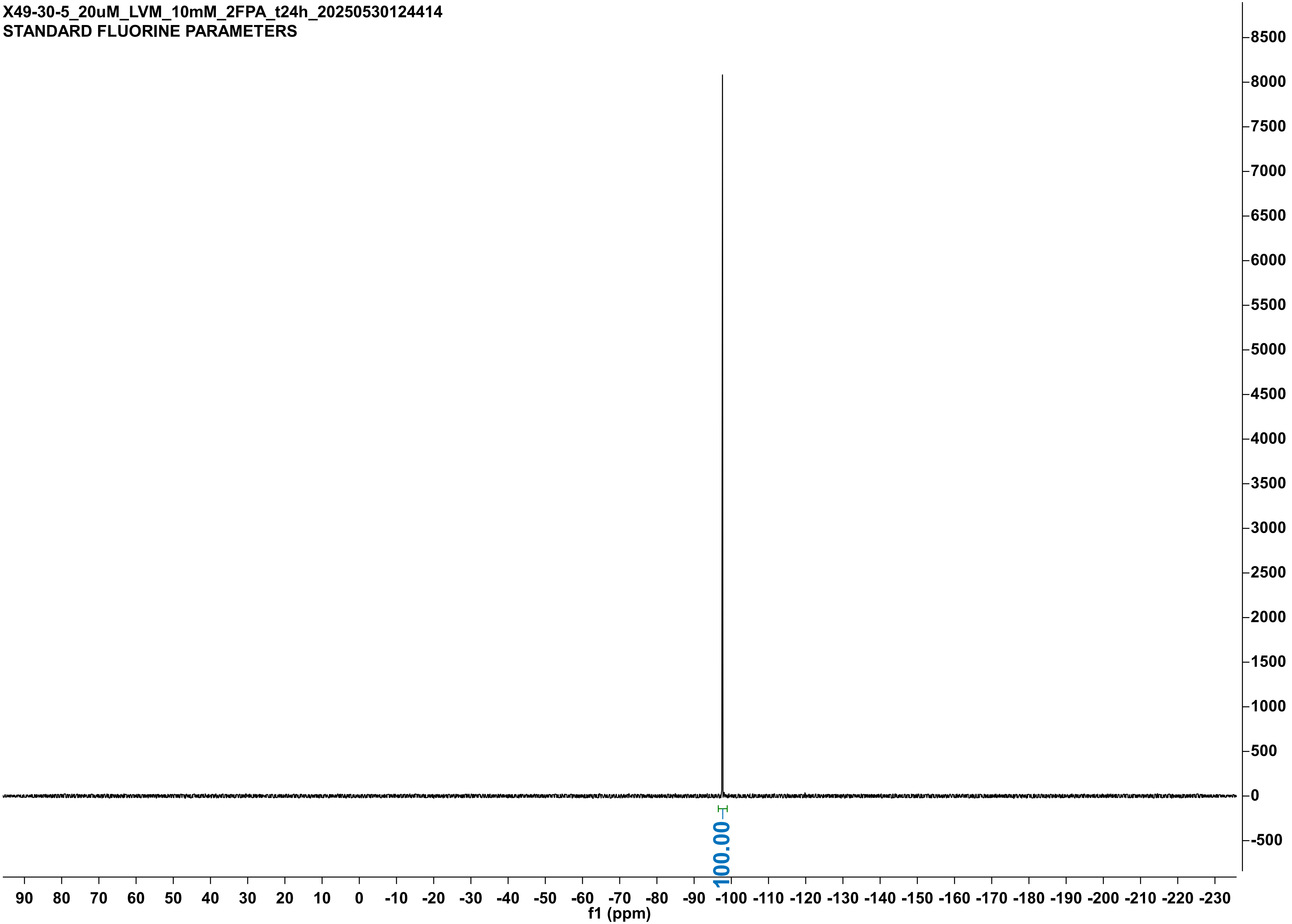

X49-30-5_20uM_Q245A_10mM_2FPA_t24h_20250530133729
STANDARD FLUORINE PARAMETERS

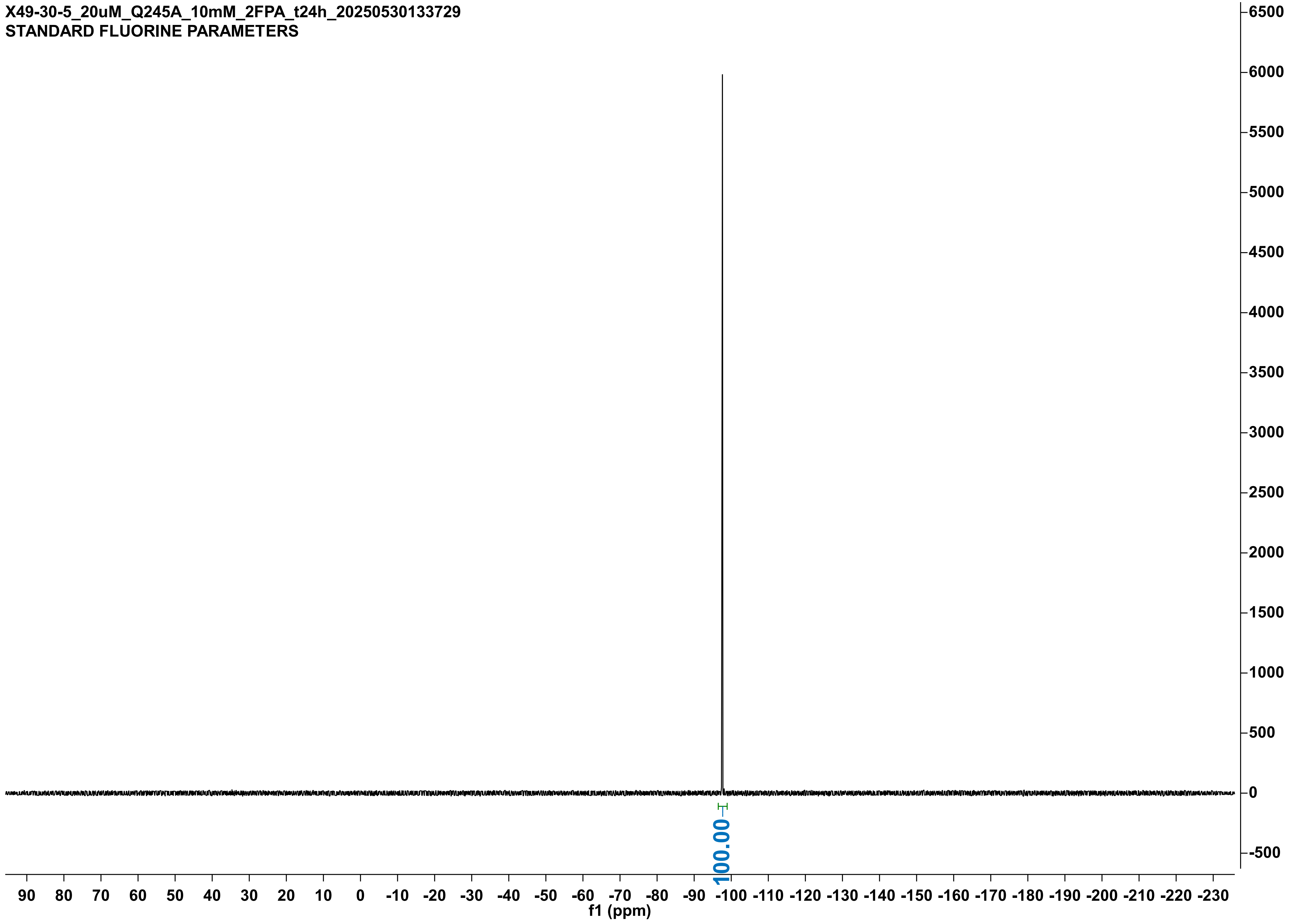

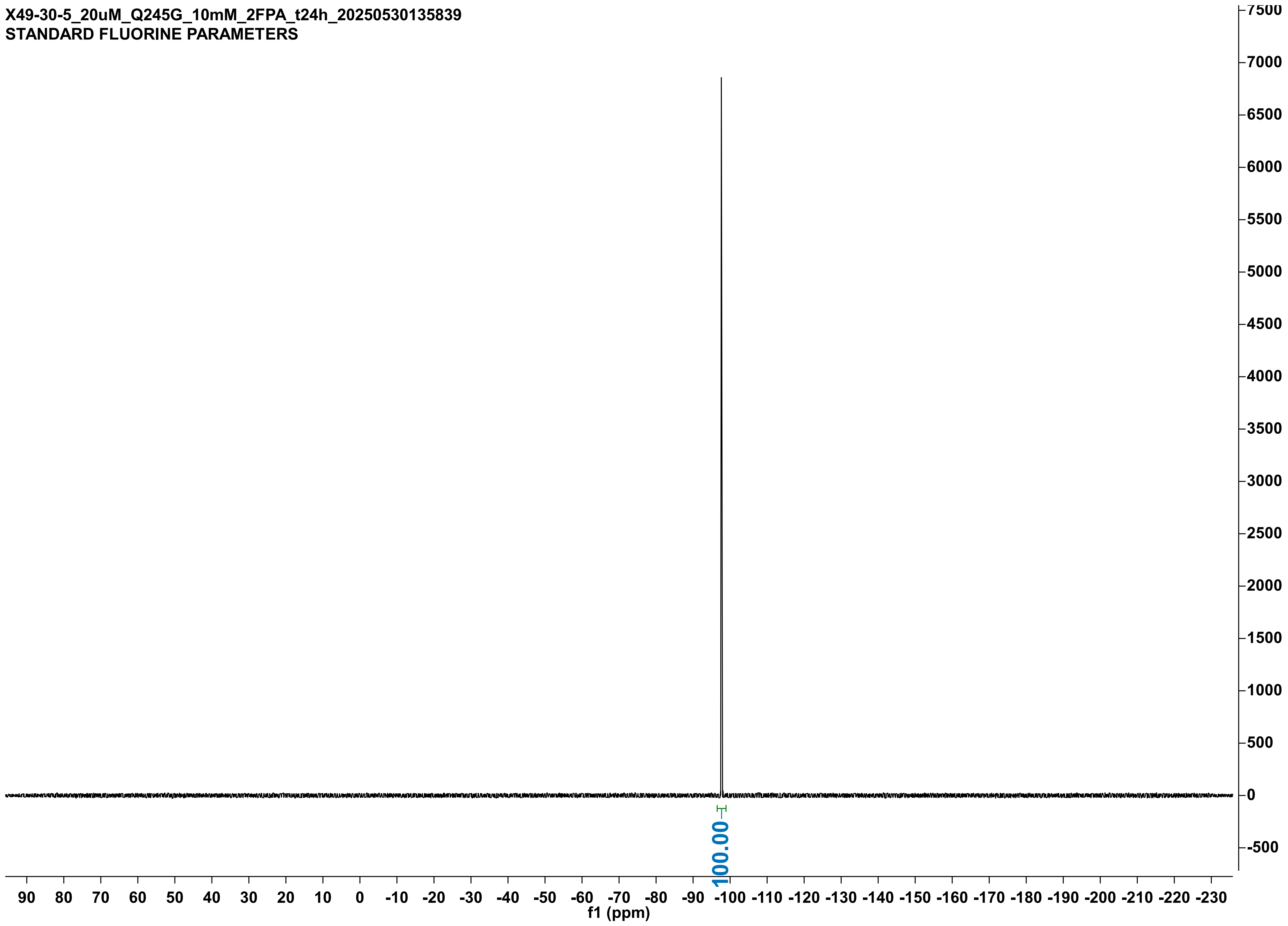
X49-30-5_20uM_Q245G_10mM_2FPA_t24h_20250530135839
STANDARD FLUORINE PARAMETERS
100.00
f1 (ppm)

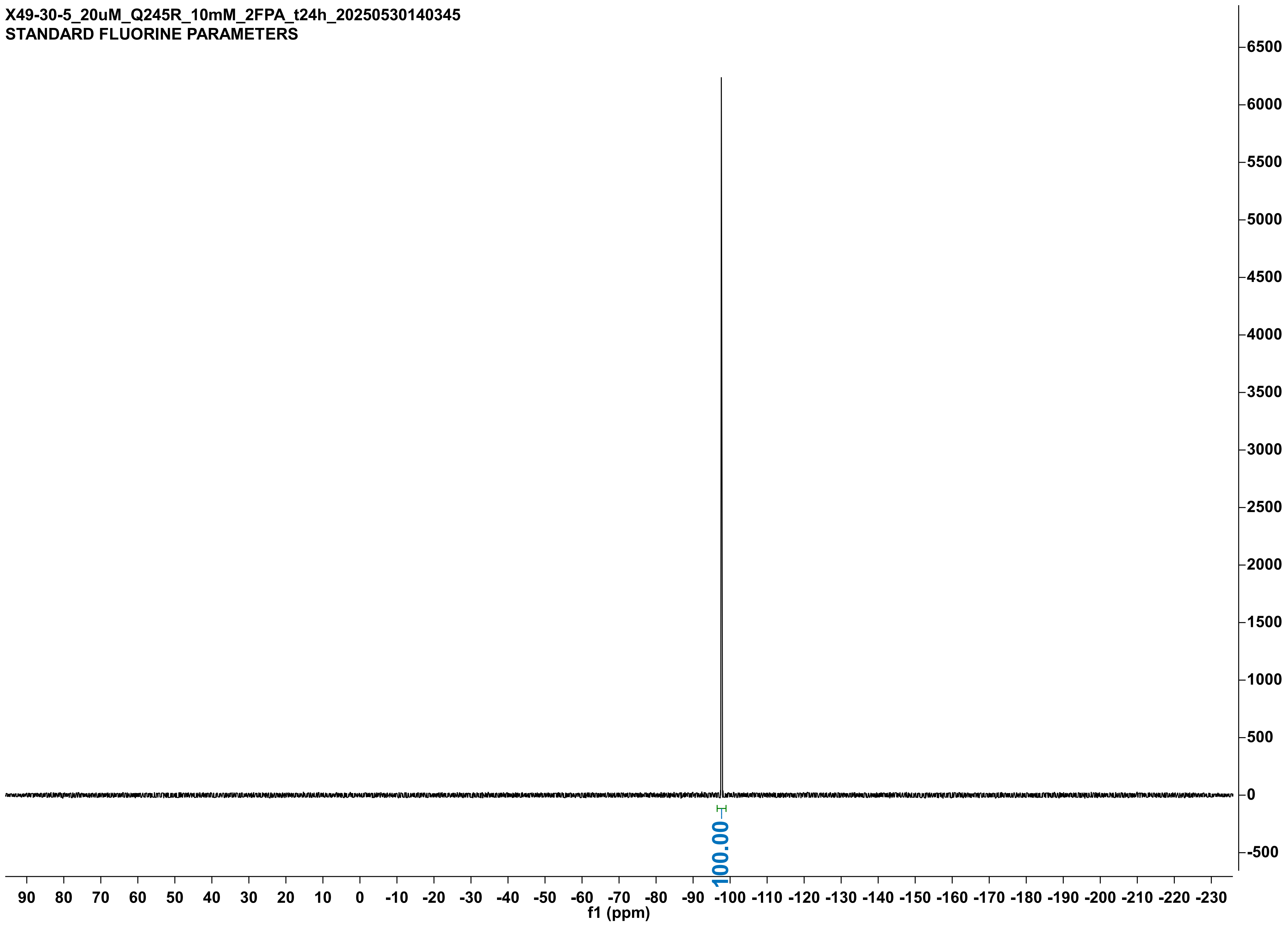
X49-30-5_20uM_Q245R_10mM_2FPA_t24h_20250530140345
STANDARD FLUORINE PARAMETERS
100.00
f1 (ppm)

X49-30-5_20uM_-VM_10mM_2FPA_t24h_20250530132959
STANDARD FLUORINE PARAMETERS

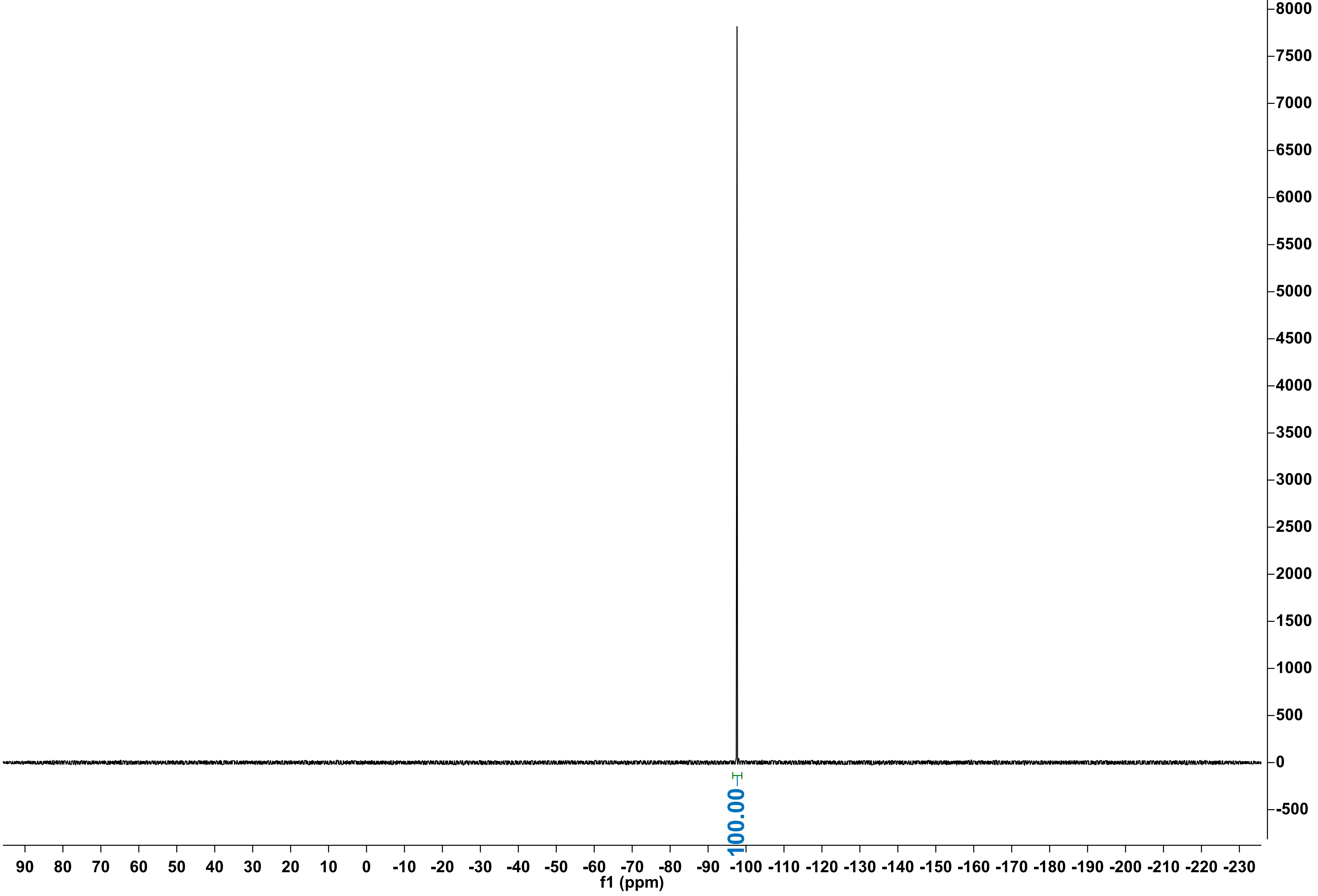
